# Photoaffinity capture compounds to profile the Magic Spot Nucleotide interactomes

**DOI:** 10.1101/2021.12.15.472736

**Authors:** Thomas M. Haas, Benoît-Joseph Laventie, Simon Lagies, Caroline Harter, Isabel Prucker, Danilo Ritz, Raspudin Saleem Batcha, Danye Qiu, Wolfgang Hüttel, Jennifer Andexer, Urs Jenal, Henning J. Jessen

## Abstract

Magic Spot Nucleotides (MSN) regulate the stringent response, a highly conserved bacterial stress adaptation mechanism, enabling survival when confronted with adverse external challenges. In times of antibiotic crisis, a detailed understanding of the stringent response is of critical importance, as potentially new targets for pharmacological intervention could be identified. In this study, we delineate the MSN interactome in *Escherichia coli* and *Salmonella typhimurium* cell lysates applying a family of trifunctional photoaffinity capture compounds. We introduce different MSN probes covering diverse phosphorylation patterns, such as pppGpp, ppGpp, and pGpp. Our chemical proteomics approach provides datasets of diverse putative MSN receptors both from cytosolic and membrane fractions that, upon validation, unveil new MSN targets. We find, for example, that the dinucleoside polyphosphate hydrolase activity of the non-Nudix hydrolase ApaH is potently inhibited by pppGpp, which itself is converted to pGpp by ApaH. The photoaffinity capture compounds described herein will be useful to identify MSN interactomes under varying conditions and across bacterial species.

**TOC:** ***Molecular fishing:*** a family of trifunctional photoaffinity capture compounds enables the identification of Magic Spot Nucleotide receptors by a chemoproteomics approach.

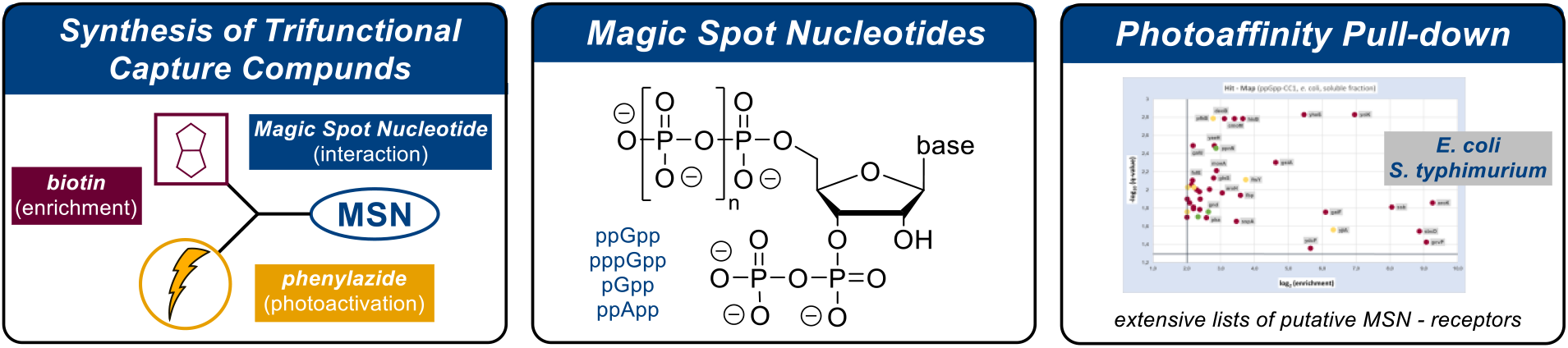

## INTRODUCTION

Magic Spot Nucleotides (MSN) constitute a class of 3’,5’-phosphorylated alarmones, featuring a characteristic 3’-diphosphate substructure (Figure 1, A). MSN are ubiquitous in bacteria, where they govern the stringent response, a fundamental stress adaptation mechanism.^[1-5]^ The stringent response enables bacteria to rapidly adjust metabolism under challenging conditions like nutrient-starvation (nitrogen, carbon, fatty-acids etc.),^[**Error! Bookmark not defined**.,6]^ extreme pH^[7]^, heat^[8]^ or antibiotics.^[9]^

**Figure 1:**
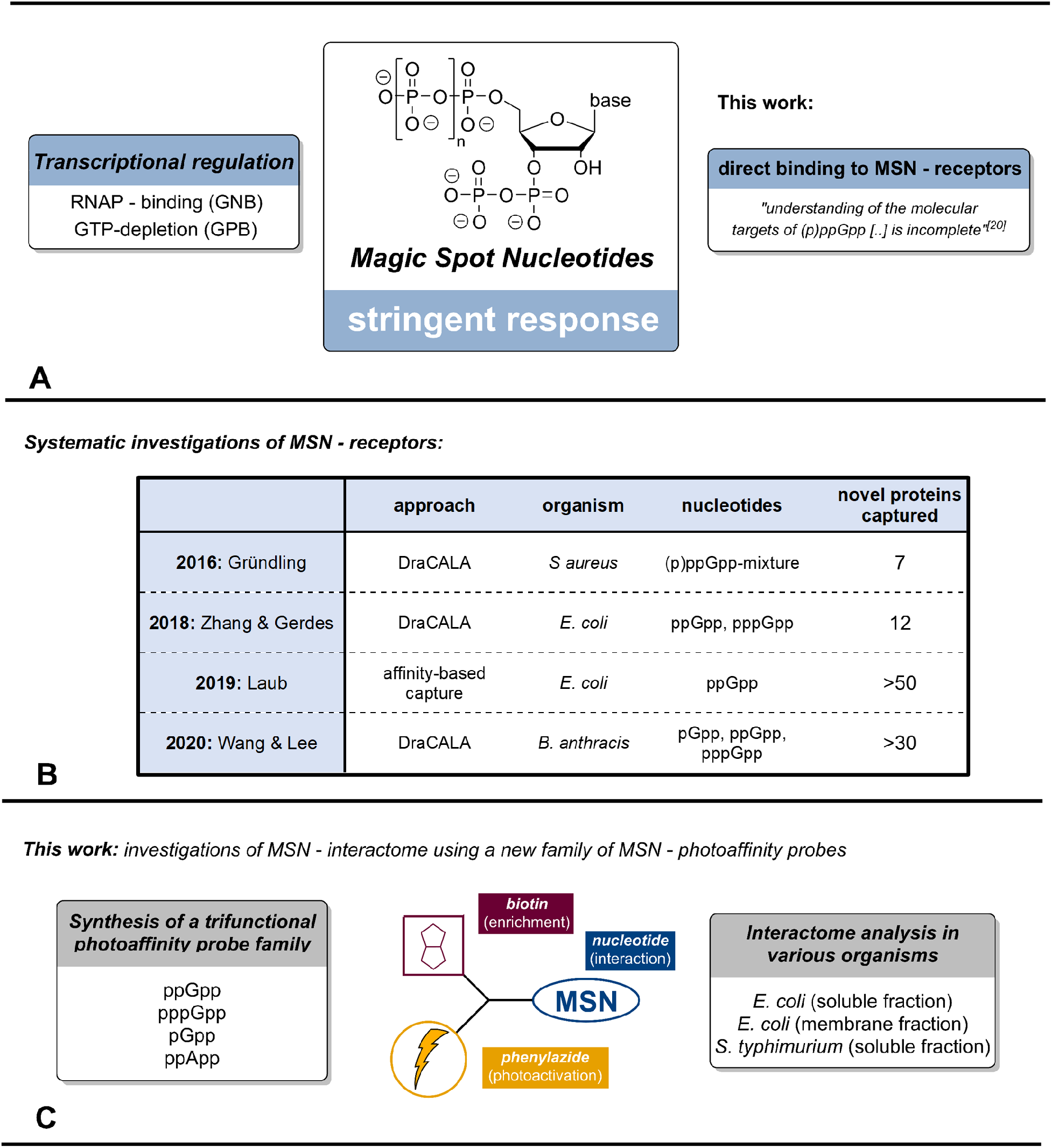
**A:** General structure of Magic Spot Nucleotides and two modes of action in the context of the bacterial stringent response. **B:** Summary of previous systematic investigations of MSN receptors in different bacteria. **C:** Graphical summary of this work including a simplified representation of the trifunctional capture compounds. RNAP: RNA polymerase.

Produced and degraded by RelA/SpoT Homologue (RSH)-enzymes and small alarmone synthetases (SAS), MSN such as ppGpp, pppGpp^[10]^ or pGpp^[11]^ operate through different modes of action. In gram-negative bacteria like *Escherichia coli*, (p)ppGpp directly binds to the RNA-polymerase in cooperation with DksA, altering the transcriptional profile and enabling rapid adaptation towards stresses.^[12]^ In gram-positive bacteria such as *Bacillus subtillis*, (p)ppGpp indirectly regulates transcription, through GTP-pool depletion during (p)ppGpp accumulation.^[13]^

In addition to these major transcriptional changes during stringent response, MSN directly bind and regulate several molecular targets such as proteins and riboswitches^[14]^ and modulate their function in a concentration dependent manner.^[**Error! Bookmark not defined**.,15]^ Through these direct interactions, (pp)pGpp modulates bacterial DNA replication, ribosome biogenesis and hibernation, translation, metabolism, and pathogenicity.^[16,17]^ In *Mycobacterium tuberculosis* (*Mtb*), it has recently been demonstrated that inhibition of Rel_Mtb_ -the enzyme responsible for MSN production-can directly kill nutrient starved *Mtb* and further enhances the killing potency of the known *Mtb* drug isoniazid in a mouse model. This highlights the importance to increase understanding of MSN function.^[18]^ The high conformational flexibility of the MSNs’ bisdiphosphate moiety enables various binding modes, also with regards to metal complexation, thus there is no consensus sequence of MSN-binding motives.^[19]^ Consequently, putative MSN-receptors are difficult to identify by bioinformatics. Recently, the systematic identification of new (p)ppGpp – effectors has become a focus of research by the application of different techniques, including radioactive labelling and mass spectrometry (Figure 1, B).

In 2016, Corrigan et al. presented a genome wide nucleotide-protein interaction screen in *Staphylococcus aureus* based on a differential radial capillary action of ligand assay (DRaCALA), radiolabelled [^32^P]-(p)ppGpp and an open reading frame library.^[20]^ This initial screening unveiled seven new MSN-targets. In 2018, Zhang et al. performed a DRaCALA screen, using [^32^P]-(p)ppGpp and *E. coli* proteins produced from the ASKA plasmid library.^[21]^ As a result, eight known and twelve unknown (p)ppGpp targets in *E. coli* were uncovered. In 2020, Yang et al. conducted a DRaCALA investigation using [^32^P]-(pp)pGpp, [^32^P]-pGpp, [^32^P]-pppGpp and an open reading frame library from *B. anthracis*.^[22]^ Notably, Yang et al. individually screened triphosphate, tetraphosphate and pentaphosphate modifications, generating specific interactome data sets, with substantial homology to analogue screens of *E. coli* and *S. aureus*.

In 2019, Wang et al. presented the chemoenzymatic synthesis and application of ppGpp-analogues substituted with biotin and diazirine-functionalities appended to the diphosphate moieties via phosphorothioates.^[23]^ These capture compounds were successfully applied in a unique photoaffinity-based SILAC mass spectrometry experiment using *E. coli* cell lysates. Here, more than 50 putative novel ppGpp – targets were captured.

Despite such impressive progress during the last five years, one can still consider bacterial MSN-interactomes to be understudied, e.g., regarding the unique targets of MSNs with different phosphorylation patterns and nucleobases (guanine and adenine), their variability across different species, aspects of compartmentalisation (membrane vs. cytosolic fractions), and different growth conditions. For example, Yang et al. stated in 2020 that “because binding targets differ between different species and most interactomes have not been characterized, the conserved and diversifying features of these interactomes remain incompletely understood.” Furthermore, “whether the (p)ppApp molecules exert growth control independent from (p)ppGpp via a separated protein target spectrum […] remains to be investigated”.^[5]^

The motivation of the present study is to overcome such limitations by introducing a new family of trifunctional MSN-photoaffinity capture compounds (Figure 1, C), accessible in milligram quantities by chemical synthesis. Equipped with biotin and photoreactive phenylazide functionalities,^[24]^ this new toolbox covers the most important MSN-representatives: ppGpp, pppGpp, pGpp and ppApp. Notably, in the case of the abundant ppGpp, modifications were introduced at the nucleobase as well as the 5’-diphosphate chain, enabling various binding modes, which should pull-down a more complete interactome. We applied these capture compounds in photoaffinity based pull-down experiments using the soluble and the membrane fraction of *E. coli* cell lysates.

Herein, we report extensive interactome datasets, covering numerous known and putative MSN-receptors. Despite substantial capture-overlap, our screening was indeed sensitive towards the phosphorylation pattern of the applied MSN and unveiled significant differences with regards to the nucleobase. In addition, we present the first systematic screen using a multiplexed mixture of two capture compounds for ppGpp-effectors in *S. typhimurium*, where (p)ppGpp and the stringent response already proved essential for the pathogen to invade host cells and express virulence genes.^[25]^

## RESULTS AND DISCUSSION

The synthesis of capture compounds was based on an extension of our recently presented synthetic methods in the context of MSN construction.^[26,27,28]^ Chemoselective P-amidite chemistry,^[29]^ regioselective RNase T_2_-catalysis, and NHS-ester based amidation reactions are synthetic key elements of this approach as shown in Scheme 1.

### Synthesis of amino - Magic Spot Nucleotides

MSN equipped with aliphatic amino-groups served as suitable intermediates towards diverse trifunctional MSN-capture compounds. Such amines can be chemoselectively functionalized in the presence of other nucleophilic groups as present in MSN.^[30]^ All syntheses started from commercially available ribonucleosides (guanosine (**1**), adenosine (**2**), Scheme 1). Treatment of **1** or **2** with pyrophosphoryl chloride followed by P-Cl-bond hydrolysis and RNase T_2_-induced regioselective 2’,3’-cyclophosphate-opening delivered the key intermediates pGp (**3**) and pAp (**4**) in 87 % and 43 % yield, respectively.^[26]^

To enable nucleobase modification (Scheme 1, A), pGp (**4**) was brominated selectively at the 8-position with a buffered solution of Br_2_. Subsequently, 8-bromo-pGp was aminated in a nucleophilic aromatic substitution process by 1,6-hexamethyldiamine. As the resulting amino-pGp **5** was not soluble, the amino-function was protected in 65% yield with an NHS-ester of Fmoc-glycine. The Fmoc-protected pGp derivative **6** was chemoselectively bisphosphitylated using a fluorenyl-modified P-amidite [(FmO)_2_P-N*i*Pr_2_] (**SI-1**). After oxidation, 1,8-diazabicyclo(5.4.0)undec-7-ene (DBU) induced global deprotection of Fm-groups led to amino – ppGpp **7** modified at the nucleobase in 33% yield.

**Scheme 1:**
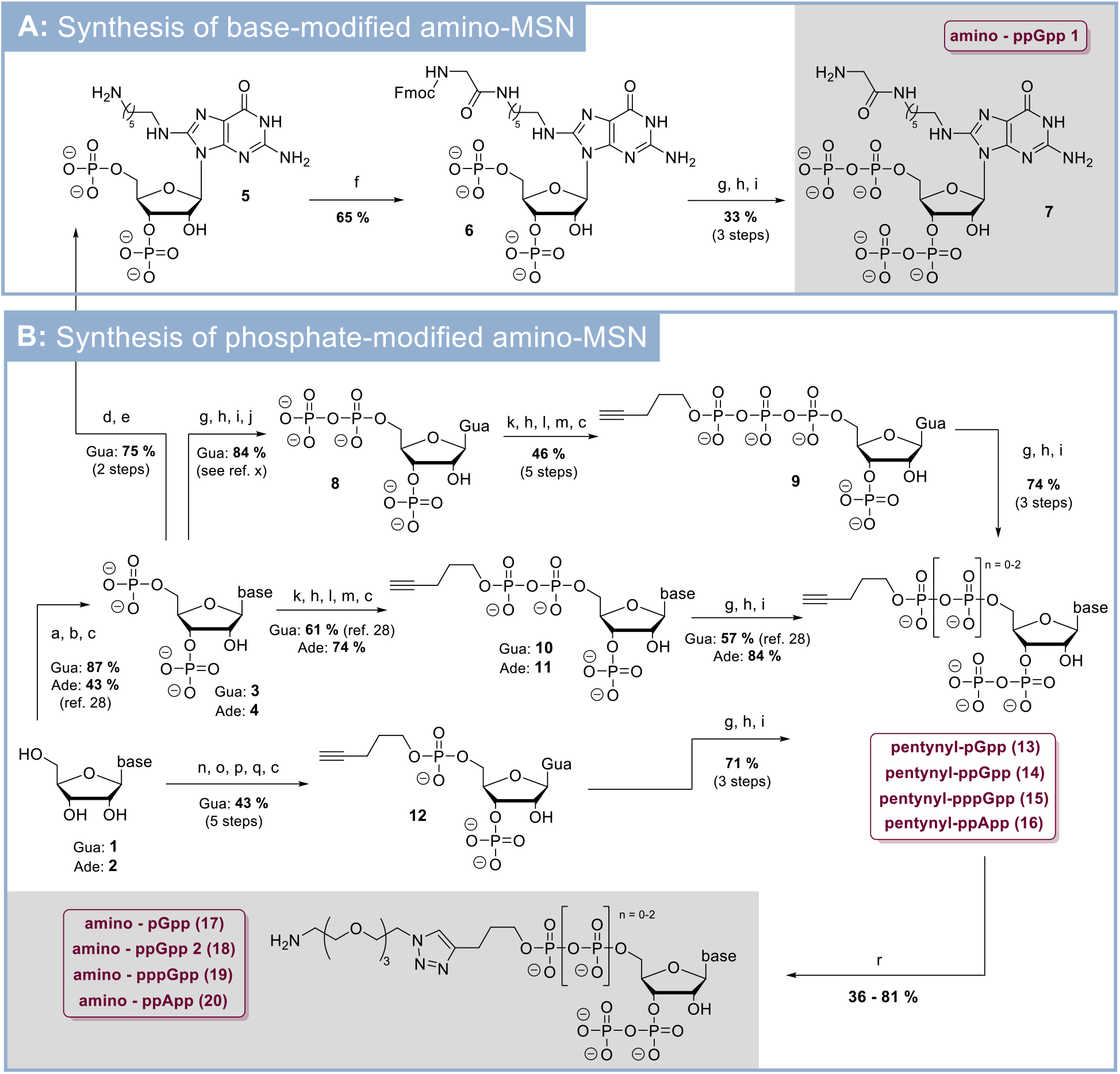
Synthesis of various amino – substituted MSN derivatives. **A:** Synthesis of amino-MSN modified at the nucleobase. **B:** Synthesis of amino-MSN modified at the 5’-phosphate chain as esters. **C:** Reaction conditions: (a) P_2_Cl_4_O_3_ (11 eq.), 0 °C, 3 h. (b) NaHCO_3_ (1 M), 0 °C. (c) RNase T_2_, pH 7.5, 37 °C, 12 h. (d) Br_2_ (3.5 eq.), H_2_O, NaOAc – buffer (200 mM, pH = 4), rt, 1 h. (e) 1,6-diaminohexane / H_2_O (pH = 9.8), 115 °C, 12 h. (f) Fmoc-Glu-OSu (3 eq.), DMSO/H_2_O, rt, 12 h. (g) (FmO)_2_P-N*i*Pr_2_ (3.0 eq.), ETT (5.0 eq.), DMF, rt, 15 min. (h) *m*CPBA (3.0 eq.), −20°C, 15 min. (i) DBU, rt, 30 min. (j) RNase T_2_, pH = 5.5, 37 °C, 12 h. (k) (FmO)(pentynylO)P-N*i*Pr_2_ (2.2 eq.), ETT (5.0 eq.), DMF, rt, 15 min. (l) MeOH, 37 °C, 4 h. (m) piperidine / DMF (1/4 v/v), rt, 30 min. (n) (FmO)(pentynylO)P-N*i*Pr_2_ (1.7 eq.), ETT (4.0 eq.), DMF/DMSO, 0 °C, 1 h. (o) (FmO)P-(N*i*Pr_2_)_2_ (1.3 eq.), rt, 45 min. (p) TBHP (3.3 eq.), rt, 1 h. (q) piperidine (20 vol%). (r) Amino-PEG_3_-azide (1.5 eq.), Na-ascorbate (1.8 eq.), CuSO_4_ x 5 H_2_O (0.35 eq.), TEAA-buffer (pH = 7, 200 mM), rt, 3 h. *Abbreviations:* Ade: adenine; DBU: diazabicycloundecene. DMF: dimethylformamide; DMSO: dimethylsulfoxide. ETT: 5-(ethylthio)-1*H*-tetrazole. Fm: fluorenylmethyl; Fmoc: fluorenylmethoxycarbonyl. Glu: glycine. Gua: guanine; *m*CPBA: *meta*-chloro perbenzoic acid. OSu: 1-hydroxy-2,5-pyrrolidindion. PEG: polyethylene glycol. TBHP: *tert*-butylhydroperoxide. TEAA: triethylammonium acetate.

pNp **3** and **4** were also crucial for the syntheses of MSN-derivatives modified at the 5’-phosphate chain with terminal esters. pGp (**3**) was transformed into ppGp (**8**) in a four-step telescoping sequence consisting of bisphosphitylation, oxidation, Fm-deprotection and RNase T_2_-catalysis, resulting in 87% yield.^[26]^ **8** was subsequently bisphosphorylated by (pentynylO)(FmO)P-N*i*Pr_2_ (**SI-3**). After oxidation, the intermediate was dissolved in MeOH, which induced the formation of a 2’,3’-cylcophosphate under concomitant phosphate group removal at the 3’-end. After Fm-deprotection on the 5’-end with piperidine, the 2’,3’-cyclophosphate was hydrolyzed regioselectively by RNase T_2_ leading to pentynyl-pppGp (**9**) in 46% yield after 5 steps from ppGp (**8**) without purification of any intermediate.

pGp (**3**) and pAp (**4**) were both bis-diphosphorylated using pentyne-substituted P-amidite **SI-3** followed by oxidation. The obtained bis-diphosphate intermediates were treated with MeOH, leading to the corresponding 2’,3’-cyclophosphates. The Fm-group on the 5’-end was removed by piperidine and the 2’,3’-cyclophosphates were regioselectively hydrolyzed by RNase T_2_–catalysis leading to pentynyl-ppGp (**10**) and pentynyl-ppAp (**11**) in yields of 61% and 74%, each without intermediate purification.

In addition, guanosine (**1**) was transformed to pentynyl-pGp (**12**) in 5 steps and 43% yield: First, **1** was regioselectively phosphitylated with P-amidite **SI-1** at the 5’-position.^[31]^ The intermediate was treated with the P-diamidite **SI-2** under formation of a 2’,3’-cyclophosphite. Oxidation and Fm-deprotection delivered the corresponding 2’,3’-cyclophosphate, that was ring-opened by RNase T_2_ to give pentynyl-pGp (**12**) in 43% yield after five telescoping steps.

**Scheme 2:**
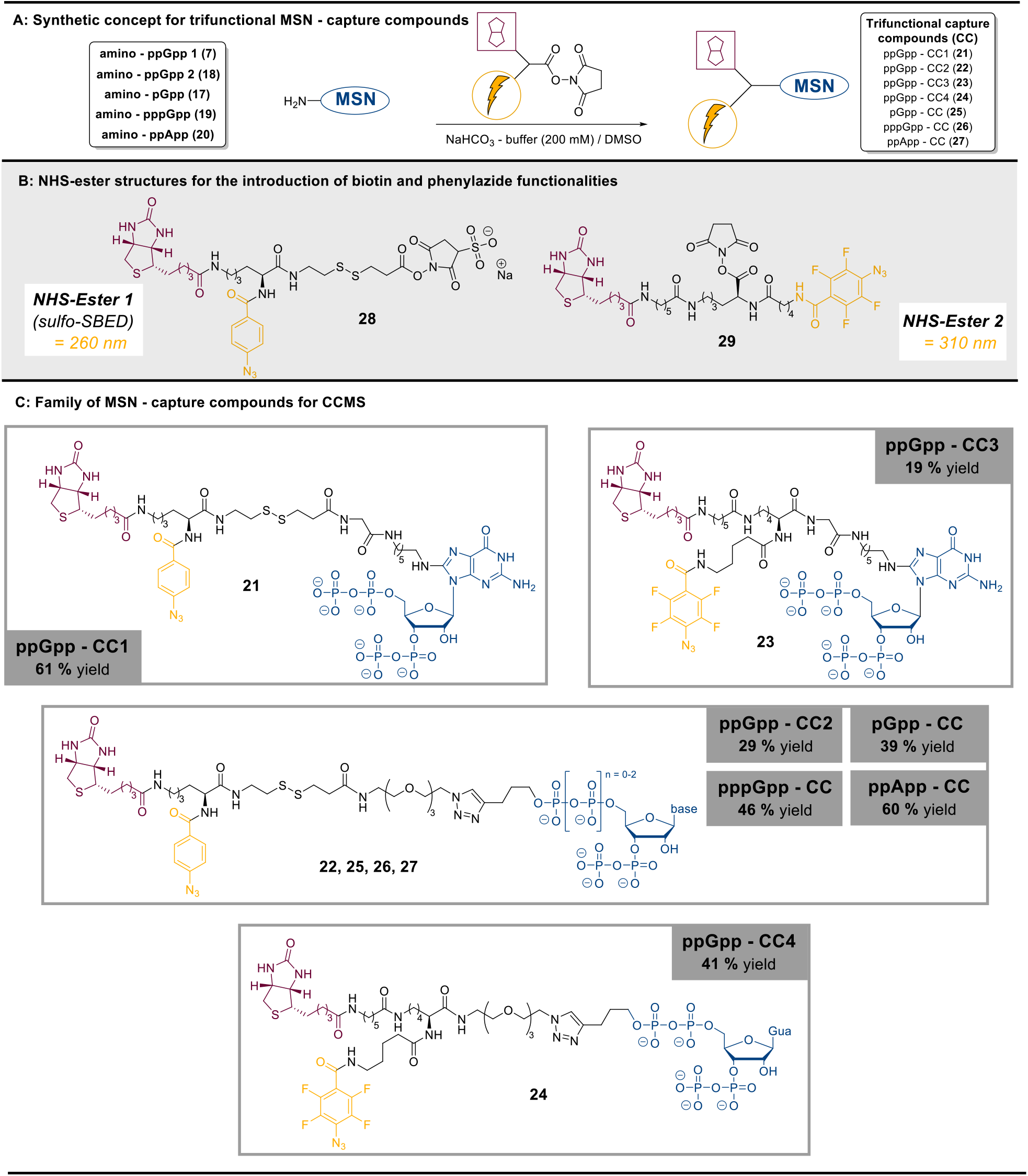
Overview of the trifunctional MSN capture compounds. **A:** NHS-ester modification as a synthetic concept towards trifunctional capture compounds. **B:** Structural representation of NHS-esters bearing biotin moieties and photoreactive phenylazide groups (optimal irradiation wavelength for nitrene generation indicated in orange). **C:** Structures and yields of various MSN capture compounds.

The four pentynylphosphates **9, 10, 11** and **12** were chemoselectively phosphorylated at the 3’-phosphate^[27]^ leading to a group of pentyne-substituted Magic Spot Nucleotides structures (**13**-**15**) in yields of 57-84 % in three steps without purification of intermediates. Alkyne tagged pentynyl-MSN **13, 14, 15**, and **16** are a versatile platform to obtain libraries of functionalized MSN for diverse applications via click chemistry.^[32]^ In this particular case, we used these intermediates in a Cu-catalyzed 1,3-dipolar cycloaddition reaction with amino-PEG_3_-azide. The corresponding triazole-containing click products (**17**-**20**) represent a family of phosphate modified amino-MSN and were isolated in yields of 36-81%. The amino functionality now enabled ready modification with commercially available NHS esters, such as e.g., photoaffinity labeled and biotinylated linkers (Scheme 2).

### Transformation of amino-MSN into trifunctional capture compounds

Biotin and a photoreactive phenylazide group were introduced to transform amino-MSN (**7, 17, 18, 19, 20**) into trifunctional capture compounds for photoaffinity pull-down. In this context, NHS-esters seemed particularly suitable, as they enable chemoselective amide bond formation with amines, presumably even in the context of highly modified MSN. Two different NHS-esters were applied in reactions with amino-MSN (Figure 2, A+B). NHS-ester 1 (**28**) is the commercially available *Sulfo-SBED*, containing biotin and a photoreactive phenylazide group. NHS-ester 2 (**29**) was designed and synthesized in our group (synthesis see SI) to include a second photoaffinity handle. Based on lysine as a trifunctional platform, **29** is functionalized with biotin and tetrafluorophenylazide. **28** and **29** differ in linker lengths, optimal wavelength for photoactivation and reactivity of the generated nitrene.^[33]^

**Figure 2:**
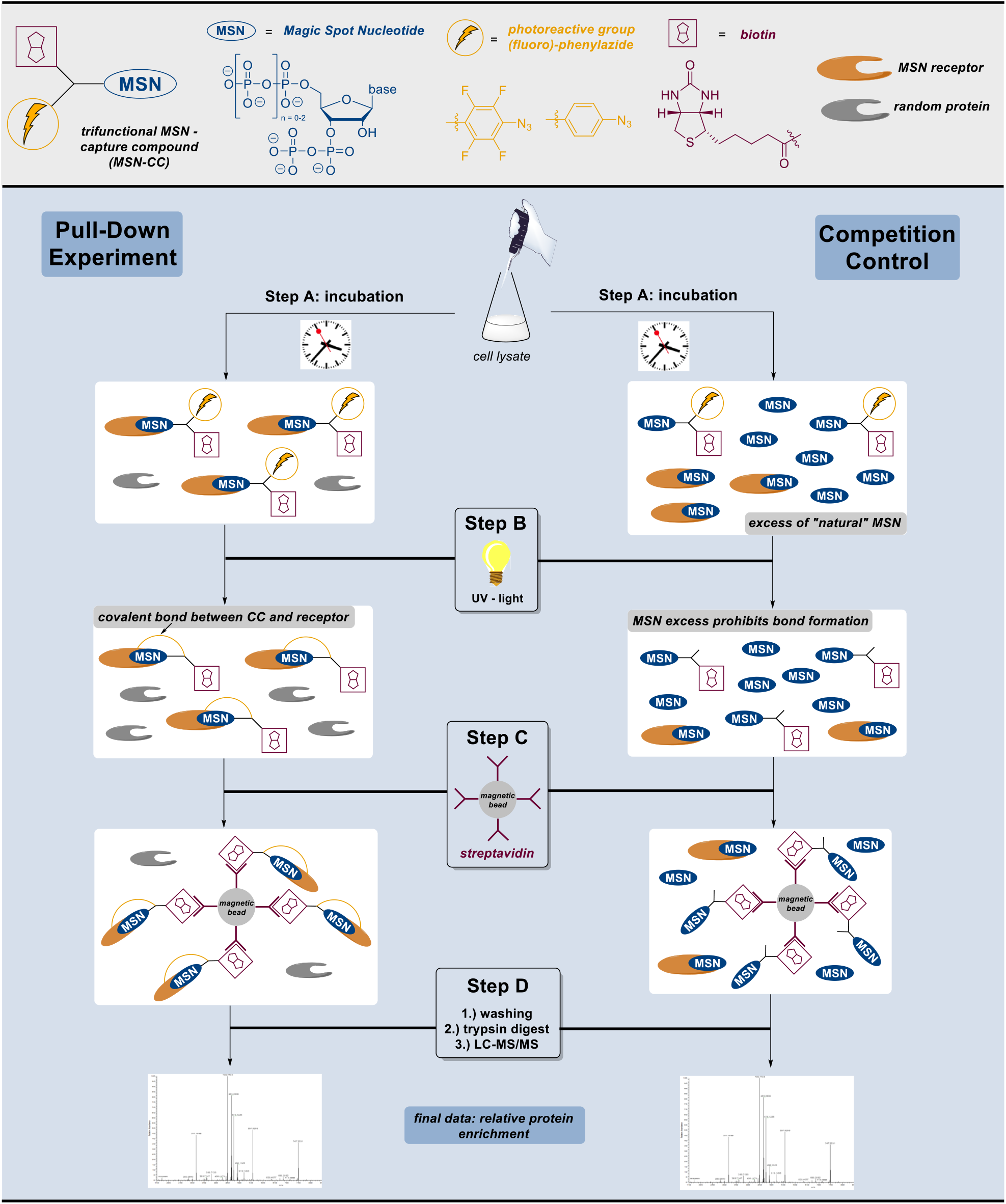
**Top (light grey):** explanation of simplified MSN – capture compound representation. **Bottom (light blue)**: Comparative workflow of pull-down experiment and the corresponding competition control experiment. **Step A:** incubation of capture compounds and MSN-competitors with cell lysates. **Step B:** UV-irradiation induces photo-crosslinking. **Step C:** Streptavidin coated magnetic beads allow the separation of captured proteins from the lysate. **Step D:** Trypsin digestion of captured proteins followed by LC-MS/MS-analysis.

In summary, various amino-MSN and NHS-esters were coupled successfully under slightly basic aqueous conditions in yields of 19–61% on highly functionalized, polar MSN-scaffolds. This resulted in a family of diverse trifunctional MSN-capture compounds, including four ppGpp-CC (**21**-**24**), pppGpp-CC (**25**), pGpp-CC (**26**) and ppApp-CC (**27**) (Scheme 2, C). The single capture compounds were synthesized on milligram scale, highlighting the effectiveness of our chemoenzymatic approach. Despite their structural complexity, we were able to thoroughly characterize the final products by ^1^H-NMR, ^31^P{^1^H}-NMR, ^19^F-NMR, HRMS and HPLC-UV (supporting information).

Notably, a capture compound (**SI-13**) based on ppGp (**8**), an understudied constitutional isomer of pGpp (or GTP), was synthesized and applied in pull-down experiments (see supporting information). Structurally, ppGp (**8**) is not part of the class of Magic Spot Nucleotides, but its presence and regulatory function in *E. coli* and *B. subtilis* has been suggested already in 1979.^[34,35]^ Furthermore, high ppGp levels were observed under N-starvation in *Streptomyces clavuligerus* and *S. coelicolor*, indicating a potential role as alarmone in these species.^[36,37]^

### The pull-down experiments

The capture compounds were applied in an experimental pull-down set-up developed by Laventie et al. 2017.^[38]^ A summary of the basic workflow is shown in Figure 2. Initially, bacterial cell lysates of *E. coli* and *S. typhimurium* were prepared (see the supporting information). Additionally, the membrane fraction of *E. coli* was resolubilized using *n*-dodecyl-B-D-maltoside as a detergent and processed separately as described in the supporting information.

The pull-down experiments (Figure 2, left) commenced with incubation of the bacterial cell-lysate with single MSN-capture compounds (step A). Subsequently, the mixture was irradiated with UV-light (310 nm), transforming the phenylazides into nitrene species (step B).^[24]^ The highly reactive nitrene undergoes follow-up reactions, such as CH insertions, leading to covalent bond-formation in its vicinity, which is ideally occupied by MSN receptors. The CC-protein complexes were pulled down by the addition of streptavidin coated magnetic beads. The residual lysate was removed by stringent washing, the captured proteins were trypsinized and analyzed by mass spectrometry.

In the competition control experiment (Figure 2, right), a 1000-fold excess of unmodified MSN competitor was present in the incubation mixture. The resulting preferential occupation of MSN-binding pockets by the MSN competitor prevents specific CC-protein interactions. At the same time, unspecific interactions not based on MSN-binding pockets are not suppressed, thereby reducing the background of false-positive hits. The competition control is processed identically concerning the subsequent steps. All experiments described were performed in triplicates. For evaluation, the relative protein enrichment of the pull-down samples compared to the competition control equates the final data set. The enriched proteins should derive from specific interaction with the MSN-moiety of the capture compounds and hence can be considered as putative MSN – receptors.

### Pull-down results: analysis and discussion

Figure 3, A summarizes and compares the results of the chemical proteomics experiment using different capture compounds and *E. coli extracts* (cytosolic vs membrane fraction). Furthermore, hit-overlaps between the different capture compounds are visualized and examples of enrichment lists as well as the corresponding hit-maps are shown (Figure 3, B+C). The thresholds of enrichment for incorporation of proteins into the lists were log_2_(enrichment) > 2, corresponding to a minimum 4-fold increase compared to the competition control. The q-value threshold was set to 0.05 to exclude most of the statistically not significant results. The detailed lists of putative MSN-receptors for all pull-down probes as well as their graphical summaries (hit-maps) are available in the supporting information (chapter 4). Given the large data sets, the following discussion is limited to a qualitative focused analysis of the pull-down results.

**Figure 3:**
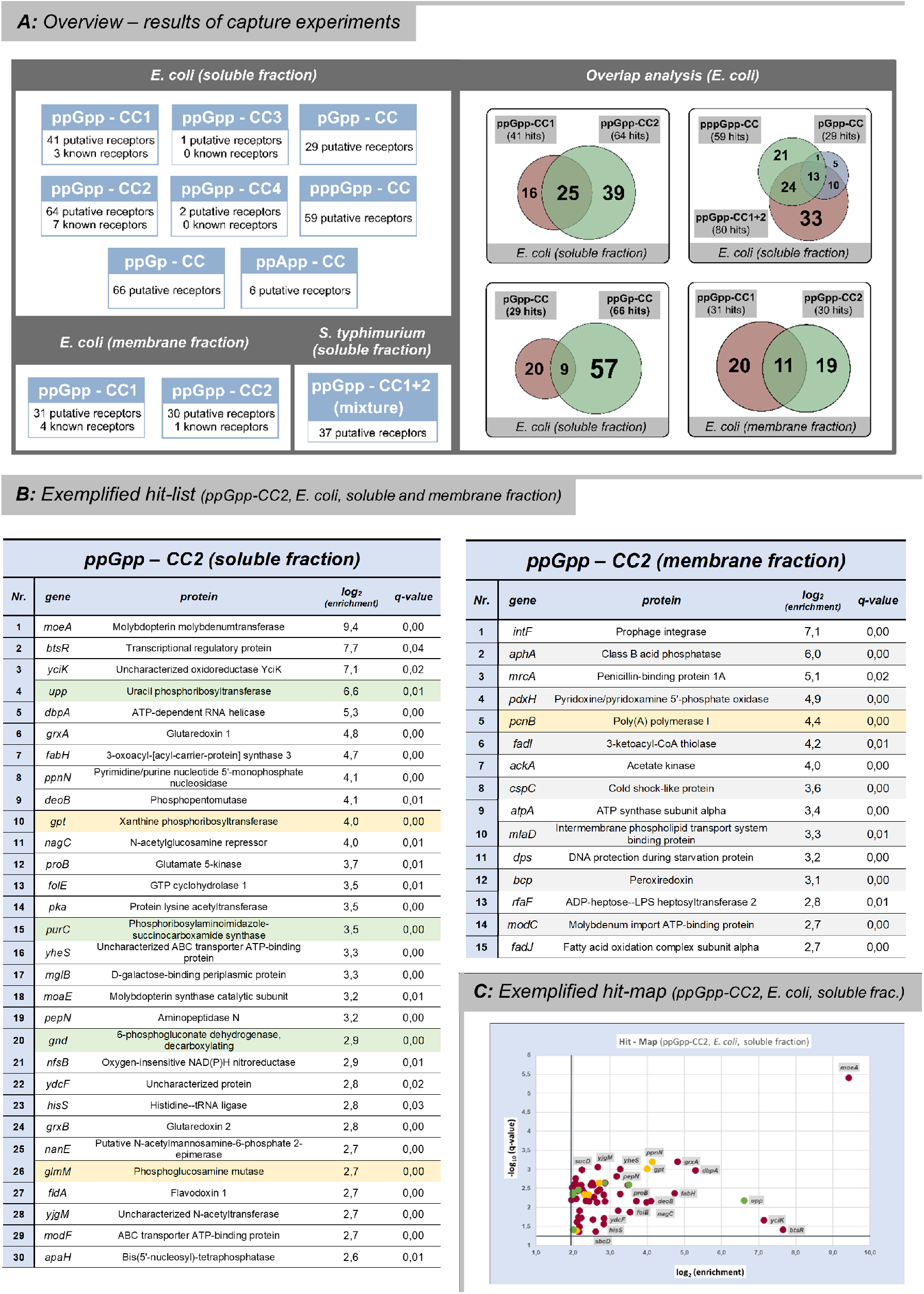
**A:** *Left:* Overview of datasets generated by various pull-down experiments including the numbers of hits. Thresholds (SI-data): log_2_(enrichment) > 2.0, q-value < 0.05. *Right:* Hit-overlap analysis. **B:** Exemplary hit-list extracts from ppGpp-CC2 (*E. coli*, soluble and membrane fraction). Known ppGpp – receptors are marked in green.^[5]^ Proteins already captured by Wang *et al*. are marked in yellow.^[23]^ **C:** Exemplary hit-map representation of ppGpp-CC2 (*E. coli* soluble fraction).

In a first round of pilot experiments, the efficiency of ppGpp-CC1+2 was compared to ppGpp-CC3+4 using the *E. coli* soluble lysate fraction. These capture compounds differ in their linker structure as well as the photoreactive moiety (phenylazide vs tetrafluorophenyl azide) and the attachment of the linker to either the 5’-diphosphate or the nucleobase. The results were clear-cut, as ppGpp-CC3 and -CC4 (same linker and tetrafluorophenyl azide) led to a combined enrichment of only three proteins. In contrast, ppGpp-CC1 and ppGpp-CC2 pulled-down 41 and 64 *E. coli* proteins, respectively, from the soluble fraction according to the threshold criteria described above (Figure 3, A). Despite an overlap of hits between ppGpp-CC1 and ppGpp-CC2, 68% of the combined hit count was specific to one of the CCs. (Figure 5, A, right). These results demonstrate that the point of attachment is of crucial importance for pull-down performance and that multiplexing of probes will likely provide superior coverage. Among the enriched proteins, three (ppGpp-CC1) or eight (ppGpp-CC2), respectively, were known ppGpp-receptors validating our approach.^[**Error! Bookmark not defined**.]^

Furthermore, several hits from our analysis were identical to the proteins found by Wang et al. 2019 (nine proteins from ppGpp-CC1 and CC2).^[23]^ Overall, we interpret these congruences as being supportive for the quality of our data, and conclude that our hits likely represent putative ppGpp-receptors. The vast majority (> 75 %) of the putative *E. coli* ppGpp-receptors described in our dataset have not been related to MSN before, and thus can serve as starting points for further biological investigations (vide infra).

As we found that the commercial *Sulfo-SBED*-linker (Scheme 2, B) gave good results consistently and irrespective of the attachment point (nucleobase vs. 5’ oligophosphate), further MSN-derivatives (pGpp-CC, pppGpp-CC and ppApp-CC) were then applied in pull-down experiments based on this linker, initially using the *E. coli* soluble fraction. In the case of pGpp-CC (29 hits) the hit-overlap with ppGpp-CC and pppGpp-CC was high (>80 %) but nevertheless, we obtained five unique hits: transcriptional regulator YjdC, D-mannonate reductase UxuB, tryptophane synthase beta chain TrpB, cold shock-like protein CspE, and glucose-phosphate-dehydrogenase Zwf. Interestingly, transcriptional control of the *zwf* operon by MSN was already observed by Shyp et al. in 2021 in *C. crescentus*.^[17]^ A subset of unique targets, despite a high overlap of pGpp with (p)ppGp regarding interactors also has recently been found by Wang using DRaCALA in Bacillus.^[22]^ The differentiation of MSN targets with regards to the degree of phosphorylation is underlined even more in the case of pppGpp-CC (59 hits), which enriched more than 35 % of proteins not captured by ppGpp-CCs or pGpp-CC (Figure 3, A, right).

In stark contrast to ppGpp-CC2, the structurally closely related ppApp-CC captured six proteins only (YhhA, PfkB, Eno, RibF, YeaG, Prs), with an overlap of ∼30 % compared to ppGpp-CCs. Until today, ppApp generation is under debate in *E. coli* wildtype.^[39]^ Nonetheless, (p)ppApp has recently attracted substantial attention also in combination with toxins-antitoxin systems.^[40,41]^ Therefore, while still unclear whether there is a relevance for this novel MSN representative in *E. coli*, the delineation of potential binding partners is very attractive. Our finding supports the idea, that the interactome of ppApp and ppGpp is not superimposable.^[42]^ Moreover, the low number of identified interactors using our approach could hint towards a more focused cellular function of ppApp.

In a next round of experiments, ppGpp-CC1 and -CC2 were applied in pull-downs using the resolubilized membrane fraction of *E. coli* cell lysates. The overlapping hits (∼50 %) between membrane and soluble lysate fraction probably arise from membrane-associated soluble proteins or incomplete removal of soluble proteins from the membrane pellet. Nevertheless, numerous membrane-associated proteins were now also identified as putative MSN interactors for example ArtP (arginine transport binding protein), ModC (molybdenum import ATP-binding protein), MrcA (penicillin-binding protein), MlaD (intermembrane phospholipid transport system), DtpA (dipeptide and tripeptide permease) and DamX (cell division protein).

Finally, a mixture of ppGpp-CC1+2 was applied in a pull-down experiment for a first systematic investigation of ppGpp-receptors in *S. typhimurium* (soluble fraction). Multiplexing of the different capture compounds enables broader coverage of targets in one single experiment. We identified 37 putative ppGpp-receptors in total. Among these hits, 9 proteins (e.g.: Ssb, HslV, FusA, GlmM and PfkB) represent homologues to proteins that have also been captured by ppGpp-CC in *E. coli*.

Although ppGp is no Magic Spot Nucleotide in a formal sense, it is structurally highly related to MSN and a potential alarmone as described above. Consequently, a pull-down experiment was performed using ppGp-CC in *E. coli* soluble lysate fraction. Here, 66 potential interactors related to various cellular processes were identified. Notably, the hit-overlap with its constitutional isomer pGpp was only 10% of the combined hit-count, underlining the approaches’ sensitivity towards specific phosphorylation patterns. These results should support further investigations on the potential role of ppGp in the bacterial stress response.

### Target Validation: Bis(5’-nucleosyl)-tetraphosphatase ApaH is regulated by Magic Spot Nucleotides *in vitro*

Recently, Yang et al. found that pGpp is produced from pppGpp by YvcI (also termed NaaH) in *Bacillus anthracis*.^.[22]^ YvcI is a Nudix hydrolase with MutT being a homologue in *E. coli*. Interestingly, MutT has already been identified as (p)ppGpp binding target in *E. coli* by Zhang and colleagues:^[21]^ Biochemical analysis revealed that ppGpp is cleaved to pGp, while no data regarding pppGpp cleavage was shown. Ndx8 from *Thermus thermophilus*, a homologous Nudix hydrolase, also converts ppGpp to pGp and other nucleoside diphosphates to nucleoside monophosphates.^[43]^

In our *E. coli* pulldowns, we consistently identified ApaH (bis(5’-nucleosyl)-tetraphosphatase) as binding target of pppGpp, ppGpp and pGpp. ApaH degrades dinucleoside polyphopshates, such as diadenosine tetraphosphate (Ap4A), a member of a second alarmone class in bacteria (Figure 4, A1). Interestingly, ApaH would be the first non-Nudix type hydrolase interacting with MSN, while several Nudix-type hydrolases have been described.^[21,44,45,46]^ We therefore investigated, if MSNs influence the activity of ApaH to uncover potential crosstalk between these different alarmones.

**Figure 4:**
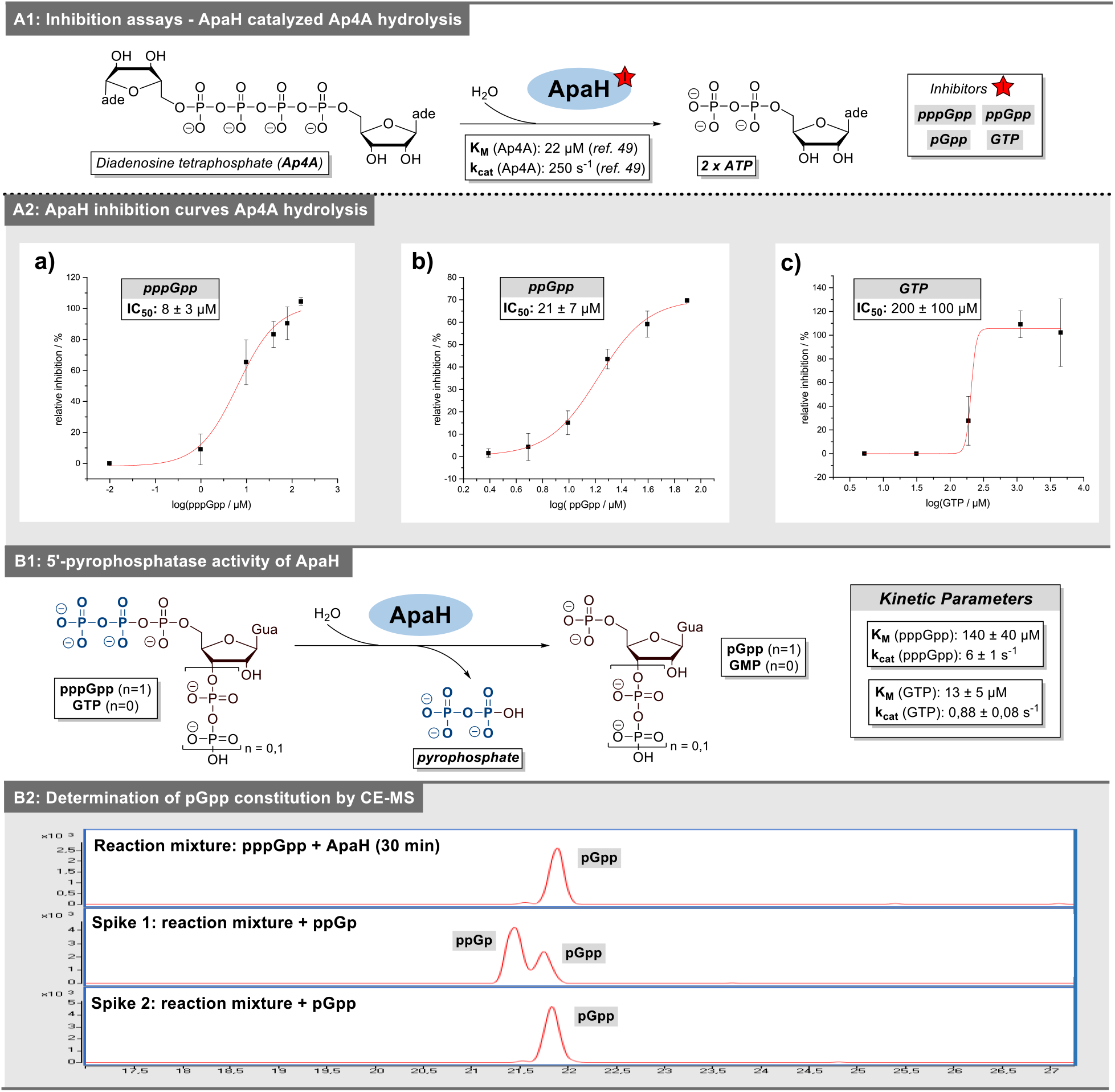
Biochemical characterization of MSN target ApaH: **A1:** Hydrolysis of Ap4A catalyzed by ApaH including overview of potential inhibitors. **A2:** Exemplary inhibition curves and IC50-values. **B1:** Graphic representation of ApaH pyrophosphatase activity including kinetic parameters. **B2:** Determination of pGpp-constitution by CE-MS (extracted ion-chromatogram for 522 m/z (triphosphorylated guanosine).

We used Ap4A as substrate. Co-incubation of either MSN with ApaH and Co^2+^ led to diminished degradation of Ap4A as determined by LC-MS (see supporting information). Consequently, IC_50_ values were determined for the MSNs as well as for GTP. pppGpp inhibited ApaH by 50% when present at 8 ± 3 μM (Figure 4, A2). The IC_50_ value of ppGpp was 21 ± 7 μM. Importantly, guanosine triphosphate (GTP) had a much higher IC_50_ value of 200 ± 100 μM. Our structural model of ApaH predicted by Alphafold readily accommodated pppGpp in the Ap4A binding pocket (supporting information, chapter 5.5).^[47]^

Since ApaH hydrolyzes a phospoanhydride bond with a nucleoside pyrophosphate as product, we examined, whether ApaH can also metabolize the MSNs pppGpp and ppGpp (Figure 4, B1). While no cleavage product was found when ApaH was incubated with ppGpp for 30 min, a GTP isomer was formed from pppGpp (supporting figure 15, A-F). This product had the same retention time as pGpp and ppGp but not as GTP. To further elucidate the cleavage product, capillary electrophoresis coupled to mass spectrometry was employed and confirmed that pGpp was produced by ApaH from pppGpp and not ppGp using synthetic references in spiking experiments (Figure 4, B2).

Next, we also determined the IC_50_ value of pGpp for Ap4A hydrolysis with 190 ± 30 μM (supporting figure 15, G). Incubation of ApaH with GTP resulted in GMP formation (Figure 4, B1), indicating ApaH requires a nucleoside 5’-triphosphate for hydrolysis. This is consistent with established ApaH activity against Ap3A, Ap4A, Ap5A, Ap6A, ATP and Ap_4_, whereas no activity is known towards Ap2A.^[48]^

Subsequently, we analyzed kinetic parameters of ApaH-catalyzed transformations. For pppGpp-hydrolysis, the Michaelis-Menten constant K_M_ and the catalytic rate k_cat_ were determined as 140 ± 40 μM and 6 ± 1 s^-1^, respectively (Figure 4, B1 and supporting figure 18). In contrast, K_M_ of GTP degradation was found to be 13 ± 5 μM and k_cat_ was 0.88 ± 0.08 s^-1^ (supporting figure 19). The latter value is consistent with the work by Plateau et al., with a k_cat_ < 1 s^-1^ for GTP. The hydrolytic reaction of Ap4A to ADP exhibits kinetic parameters of 22 μM for K_M_ and 250 s^-1^ for k_cat_ under similar conditions.^[49]^ In summary, the catalytic activity of ApaH towards pppGpp-hydrolysis is moderate compared to Ap4A-hydrolysis as primary reaction.

## CONCLUSION

In the present paper, we introduce a new family of trifunctional capture compounds based on biotin (enrichment), phenylazides (photoactivation) and various MSN structures such as ppGpp, pppGpp, pGpp and ppApp. In the case of ppGpp, modifications were installed at the phosphate chains as well as the nucleobase, facilitating different binding modes. We successfully applied this toolbox in pull-down experiments using cell lysates of *E. coli* and *S. typhimurium*. The chemical proteomics approach delivered extensive data sets of putative MSN-receptors, in part uniquely correlating with their distinct phosphorylation pattern. Several *bona fide* MSN-receptors were part of the hit-lists supporting the accuracy of the presented data. Interestingly, the majority of enriched proteins has not been correlated to MSN before. The ppGpp-pull-down was also performed using the membrane fraction of *E. coli* lysate, generating a list of putative membrane associated ppGpp – receptors.

We are confident, that the presented lists of putative Magic Spot Nucleotide receptors will be a starting point for the community to study MSN binding targets of different MSN in various organisms and more detail. Along these lines, we have validated ApaH as the first non-Nudix type hydrolase whose function is inhibited by MSN but that also degrades pppGpp to pGpp. While pGpp has not been described in *E. coli*, its recent identification in *Bacillus* points towards a potential biological relevance of our finding. We are confident, that the availability of this new class of capture compounds will pave the way for specific interactome analyses in other bacterial species under various conditions and that our probes can also be applied to plants or algae.

## ACKNOWLEDGMENT

This study was supported by the Deutsche Forschungsgemeinschaft (DFG) under Germany’s Excellence Strategy (CIBBS, EXC-2189, Project ID 390939984). This project has received funding from the European Research Council (ERC) under the European Union’s Horizon 2020 research and innovation program (grant agreement no. 864246, to H.J.J.). We gratefully acknowledge financial support from the Studienstiftung des deutschen Volkes. This work was supported by a grant of the Swiss National Science Foundation (310030B_185372) to U.J. We thank MagRes of the University of Freiburg for significant amount of time for NMR spectroscopy. We thank Prof. Dr. Bernd Kammerer for access to HPLC-MS equipment. We thank Dr. Pavel Salavei from the BIOSS protein laboratory (University of Freiburg) for protein expression and purification. We are grateful for the continuous support of the proteomics core facility at Biozentrum Basel: we especially thank Dr. Thomas K. C. Bock, Dr. Alexander Schmidt and Ulrike Lanner.

