## Supporting Info for "Photoaffinity capture compounds to profile the Magic Spot Nucleotide interactomes"

### Table of content

1. General synthetic remarks
2. Synthesis of MSN capture compounds
   2.1. Syntheses adapted from literature
   2.2. Synthesis of Pull-down linker 2
   2.3. Syntheses of base-modified amino-ppGpp
   2.4. Synthesis of pentynyl-substituted MSN
   2.5. Synthesis of phosphate-modified amino-MSN
   2.6. Syntheses of MSN-capture compounds
3. Procedural remarks, pull-down experiments
4. Capture results: enrichment tables and hit-maps
   4.1. *E. coli* (soluble fraction)
   4.2. *E. coli* (membrane fraction)
   4.3. *S. typhimurium* (soluble fraction)
5. Target validation: apaH is regulated by MSN *in-vitro*
   5.1. ApaH in-vitro assay
   5.2. LC/MS analysis of nucleotides
   5.3. $IC_{50}$ determination
   5.4. Determination of kinetic parameters ($K_M$ and $K_{cat}$)
   5.5. Molecular docking of ApaH substrates with AlphaFold
6. Supporting references
7. NMR-spectra
8. MS – spectra
9. HPLC-analyses

**Abbreviations**

| | |
|---|---|
| DBU | 1,8-Diazabicyclo[5.4.0]undec-7-ene |
| DCC | N,N’-Dicyclohexylcarbodiimide |
| DCM | Dichloromethane |
| DMF | Dimethylformamide |
| DMSO | Dimethyl sulfoxide |
| DTT | Dithiothreitol |
| EDC | 1-Ethyl-3-carbodiimide |
| $Et_2O$ | Diethyl ether |
| ETT | 5-(Ethylthio)-1*H*-tetrazole |
| Fm | Fluorenylmethyl |
| Fmoc | Fluorenylmethyloxycarbonyl |
| *m*CPBA | *meta*-Chloroperoxybenzoic acid |
| MeCN | Acetonitrile |
| HMDA | Hexamethylenediamine |
| HPLC | Reverse phase high-performance liquid chromatography |
| HRMS | High resolution mass spectrometry |
| MSN | Magic spot nucleotides |
| NHS | N-Hydroxysuccinimide |
| PEG | Polyethylene glycol |
| pGp | Guanosine-3’,5’-bisphosphate |
| pGpp | Guanosine-3’-diphosphate-5’-phosphate |
| ppApp | Adenosine-3’,5’-bisdiphosphate |
| ppGp | Guanosine-3’-phosphate-5’-diphosphate |
| ppGpp | Guanosine-3’,5’-bisdiphosphate |
| pppGpp | Guanosine-3’-diphosphate-5’-triphosphate |
| SAX | Strong anion exchange |
| TBA | Tetrabutylammonium |
| TEAA | Triethylammonium acetate |

### 1. General synthetic remarks

**Reactions** were carried out using glassware magnetically stirred, unless noted otherwise. Air- and moisture-sensitive liquids and solutions were transferred via syringe or stainless steel canula.

**Reagents** were purchased from commercial suppliers (Acros, Aldrich, Fluka, TCI) and used without further purification, unless noted otherwise.

**Solvents** were obtained in analytical grade and used as received for extractions, precipitation and solid washing.

**Dry solvents** for reactions were purchased in a dry form from Sigma and stored over molecular sieves as well as under the atmosphere of dry $N_2$.

**Deuterated solvents** for NMR and reactions were obtained from Armar Chemicals, Switzerland and euriso-top, Germany, in the indicated purity grade and used as received for NMR spectroscopy.

**Strong ion-exchange chromatography** was performed using an automated Äkta® – system. Q-Sepharose was purchased from Aldrich. Buffer solutions were produced manually using milliQ $H_2O$.

**Sulfo-SBED** biotin label transfer reagent was purchased from Thermofisher®.

**TBA-salt preparations** were performed by either using Dowex$H^+$ followed by TBA(OH) addition or Chelex®100 (preloaded with TBA). In both cases, the TBA salts were obtained after lyophilization.

**Lyophilizations** were done with Christ Freeze Dryer Alpha 1-4 LDplus and Christ Freeze Dryer Alpha 1-2 LDplus.

**Ribonuclease T2** from Aspergillus oryzae (50 ku) was purchased from Worthington Biochemical corporation as lyophilized powder and dissolved in a storage buffer [glycerol / $NaH_2PO_4$ (10 mM, pH 6.8), 1 / 1]. The stock solution was stored at -20 °C.

**$^1$H-NMR spectra** were recorded on Bruker 300 MHz spectrometers, Bruker 400 MHz (with cryoprobe) and Bruker 500 MHz spectrometers in the indicated deuterated solvent. Data are reported as follows: chemical shift (δ, ppm), multiplicity (s, singlet; d, doublet; t, triplet; q, quartet; m, multiplet; br. s, broad signal), coupling constant(s) (*J*, Hz), integration. All signals were referenced to the internal solvent signal as standard ($D_2O$, δ 4.79). After $NaClO_4$ – purification acetone residues were present in the products. These were also considered for yield determination.

**$^{13}$C{$^1$H}-NMR** spectra were recorded with $^1$H-decoupling on Bruker 126 MHz, Bruker 101 MHz (with cryoprobe) spectrometers at 298K in the indicated deuterated solvent. If possible, signals were referred to the internal solvent signal as standard.

**$^{31}$P{$^1$H}-NMR spectra and $^{31}$P-NMR spectra** were recorded with $^1$H-decoupling or $^1$H coupling, respectively, on Bruker 202 MHz , 162 MHz (with cryoprobe) and Bruker 122 MHz spectrometers in the indicated deuterated solvent. All signals were referenced to an internal standard (PPP).

**Mass spectra** were recorded by C. Warth (Mass spectrometry service of the University of Freiburg) on a Thermo LCQ Advantage [spray voltage: 2.5 – 4.0 kV, spray current: 5 µA, ion transfer tube: 250 (150) °C, evaporation temperature: 50 – 400°C.

**Capture compounds** were stored at -20 °C under light exclusion.

### 2. Synthesis of MSN capture compounds

#### 2.1. Syntheses adapted from literature

**Synthesis of (FmO)$_2$P-N(*i*Pr)$_2$ (SI-1)**

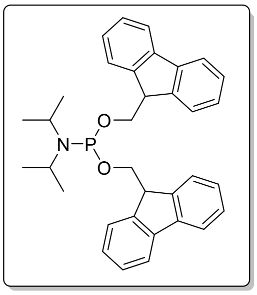

The compound was synthesized in two steps as reported previously starting from $PCl_3$. Analytical data were identical to literature.[1] The compound was stored at −20°C.

**Synthesis of (FmO)P-[N(*i*Pr)$_2$]$_2$ (SI-2)**

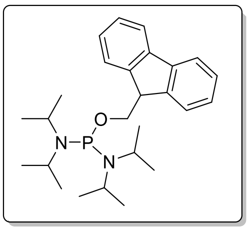

The compound was synthesized as reported previously. Analytical data were identical to literature.[2] The compound was stored at −20°C.

**Synthesis of (Pentynyl)(FmO)P-N(*i*Pr)$_2$ (SI-3)**

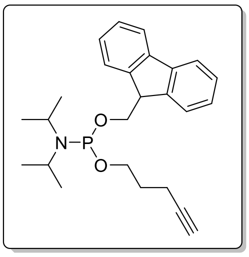

The compound was synthesized as reported previously Analytical data were identical to literature.[3] The compound was stored at −20°C.

**Synthesis of natural MSN**

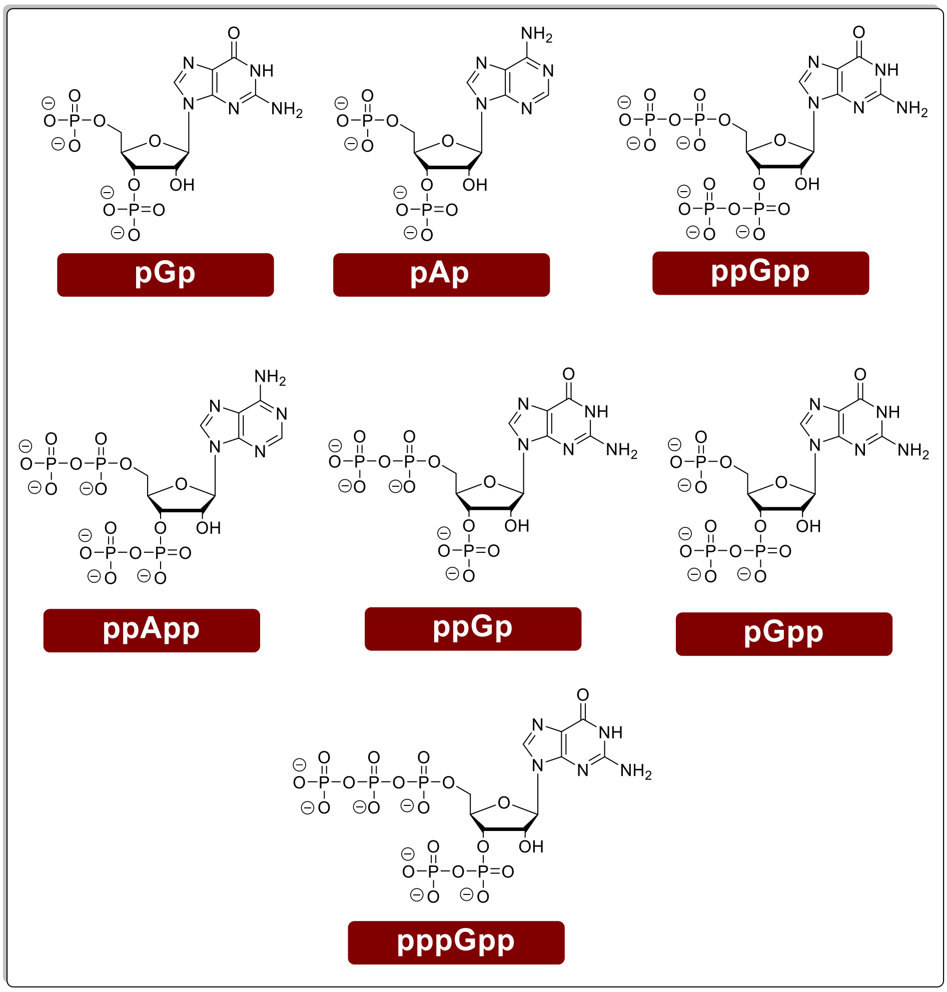


pGp, ppGpp, pppGpp, ppGp, pGpp and ppApp were synthesized according to Jessen et al.[3] The analytical data was in accordance with literature.

**Synthesis of amino-ppGpp (14)**

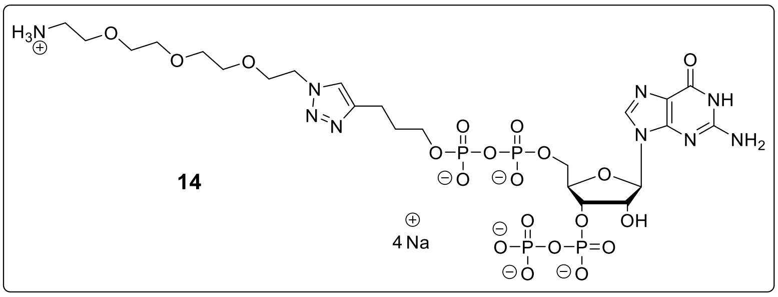


Amino-ppGpp **14** was synthesized according to Jessen et al.[3] The analytical data was in accordance with literature.

#### 2.2. Synthesis of Pull-down linker 2

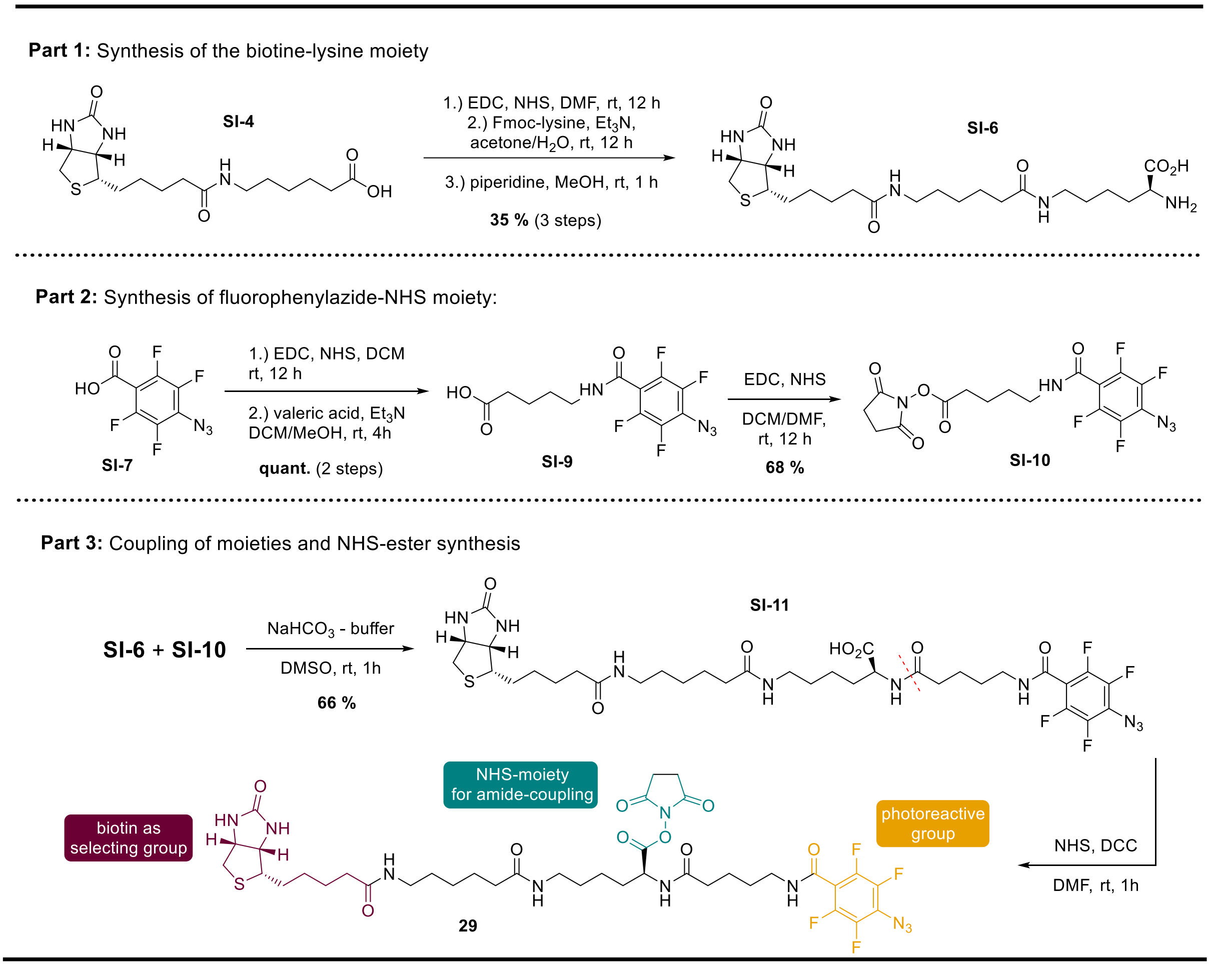


**Supporting figure 1:** Synthesis of fluorophenylazide linker **29** from commercial precursors **SI-4** and **SI-7** based on sequential, chemoselective NHS-ester couplings.

#### Synthesis of Biotin-NHS-ester SI-5

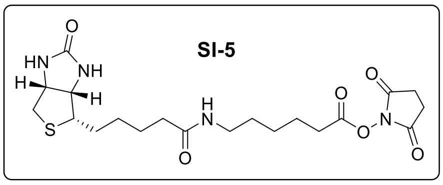


To a solution of N-(+)-Biotinyl-6-aminohexanoic acid (**SI-4**, 400 mg, 1.12 mmol) in dry DMF (20 ml), EDC·HCl (535 mg, 2.80 mmol, 2.50 eq.) and NHS (322 mg, 2.80 mmol, 2.50 eq.) were added. The solution was stirred overnight at 37 °C. Then the solvent was removed under reduced pressure and the crude product was purified by column chromatography on silica gel ($CH_2Cl_2$/MeOH, 15:1). The product **SI-5** was obtained as white solid (462 mg, 1.02 mmol, 90%).

**$^1$H-NMR** (400 MHz, DMSO-$d_6$, δ/ppm): 7.73 (t, $J$ = 5.6 Hz, 1H), 6.41 (s, 1H), 6.34 (s, 1H), 4.30 (ddd, $J$ = 7.4, 5.1, 0.9 Hz, 1H), 4.12 (ddd, $J$ = 7.8, 4.4, 1.9 Hz, 1H), 3.09 (ddd, $J$ = 8.6, 6.1, 4.4 Hz, 1H), 3.01 (ddd, $J$ = 6.5, 6.5, 6.5 Hz, 2H), 2.85 – 2.75 (m, 5H), 2.70 – 2.61 (m, 2H), 2.61 – 2.53 (m, 1H), 2.04 (t, $J$ = 7.4 Hz, 2H), 1.69 – 1.55 (m, 3H), 1.55 – 1.23 (m, 9H). **$^{13}$C{$^1$H}-NMR** (101 MHz, DMSO-$d_6$, δ/ppm): 171.81, 170.26, 168.94, 162.69, 61.04, 59.18, 55.44, 54.92, 38.06, 35.23, 30.15, 28.64, 28.23, 28.05, 25.45, 25.31, 23.96. **HRMS** (ESI) m/z for $C_{20}H_{31}O_6N_4S$ [M-H]$^+$: calcd. 455.1960, found 455.1959. **R*f*** ($SiO_2$, $CH_2Cl_2$/MeOH, 15:1): 0.31.

**Synthesis of Biotin-lysine derivative SI-6**

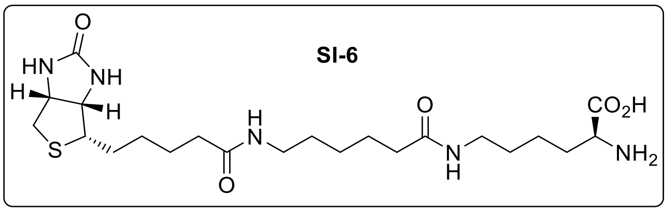


To a solution of compound **SI-5** (60.0 mg, 132 µmol) in acetone (6 ml) and $NaHCO_3$-solution (0.1 M, 3 ml) was added Fmoc-Lysine (0.146 mg, 356 µmol, 3.00 eq.). The solution was stirred overnight at room temperature. The reaction mixture was diluted with $CH_2Cl_2$ (10 ml) and extracted with $H_2O$ (2 x 10ml). The combined aqueous layers were diluted with $H_2O$ (10 ml) and the solution was acidified to pH 2.0 using HCl-solution (0.1 M). The precipitate was separated by centrifugation, washed with cold HCl-solution (0.1 M, 15 ml) and lyophilized. The resulting solid was dissolved in MeOH (2 ml) and piperidine (300 µl) was added. The solution was stirred at room temperature for 1 h. The crude product was precipitated by adding $Et_2O$ (10 ml), separated by centrifugation, washed with $Et_2O$ (10 ml) and dried under high vac. The solid was redissolved in DMSO (1 ml) reprecipitated with $Et_2O$, washed with $Et_2O$ (2 x 10 ml) and dried under high vac. The resulting solid was dissolved in $H_2O$ (2 ml) and lyophilized. The product **SI-6** was obtained as white solid (24.8 mg, 49.8 µmol, 39%).

**$^1$H-NMR** (400 MHz, $D_2O$, δ/ppm): 4.62 (ddd, $J$ = 8.0, 5.0, 0.9 Hz, 1H), 4.43 (dd, $J$ = 7.9, 4.4 Hz, 1H), 3.69 (dd, $J$ = 6.6, 5.7 Hz, 1H), 3.35 (dt, $J$ = 9.8, 5.1 Hz, 1H), 3.19 (dtd, $J$ = 6.7, 4.5, 2.3 Hz, 4H), 3.05 – 2.97 (m, 1H), 2.79 (d, $J$ = 13.1 Hz, 1H), 2.25 (td, $J$ = 7.3, 4.2 Hz, 4H), 1.91 – 1.80 (m, 2H), 1.79 – 1.27 (m, 16H). **$^{13}$C{$^1$H}-NMR** (101 MHz, $D_2O$, δ/ppm): 176.82, 176.63, 174.81, 165.35, 62.07, 60.24, 55.36, 54.68, 39.67, 39.05, 38.87, 35.66, 35.49, 30.12, 28.06, 27.99, 27.80, 27.64, 25.50, 25.17, 25.00, 21.80.**HRMS** (ESI) m/z for $C_{22}H_{38}O_5N_5S$ [M-H]$^-$: calcd. 484.2599, found 484.2598.

**Synthesis of Fluorophenylazide SI-10**

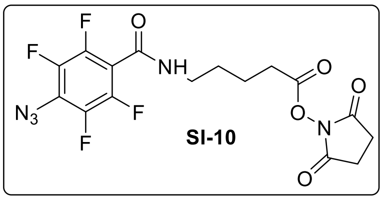

To a solution of 4-azidotetrafluorobenzoic acid (**SI-7**, 400 mg, 1.70 mmol) in dry $CH_2Cl_2$ (9.6 ml), was added EDC·HCl (390 mg, 2.04 mmol, 1.20 eq.) in one portion and the mixture was purged with argon for 5 min. Afterwards NHS (235 mg, 2.04 mmol, 1.20 eq.) was added and the solution was stirred overnight at room temperature in a foil covered flask. Afterwards the solution was washed with $H_2O$ (3 x 10 ml) and the aqueous layer was extracted with $CH_2Cl_2$ (20 ml). The combined organic layers were dried over $Na_2SO_4$ and the solvent was removed under reduced pressure.

The resulting solid (**SI-8**) was dissolved in a mixture of dry $CH_2Cl_2$ and MeOH (1/1, 20 ml). Subsequently, 5-aminovaleric acid (212 mg, 1.82 mmol, 1.10 eq.) and $NEt_3$ (228 µl, 167 mg, 1.65 mmol, 1.00 eq.) were added. The solution was stirred at room temperature in a foil covered flask. After 90 min further 5-Aminoverleric acid (39.0 mg, 0.330 mmol, 0.20 eq.) was added. The solution was stirred at room temperature for additional 3 h. The solution was diluted with $H_2O$ (15 ml) and the aqueous layer was extracted with $CH_2Cl_2$ (2 x 10 ml). The combined organics were washed with HCl (1 M, 20 ml) and sat.-aq. NaCl-solution (15 ml), dried over $Na_2SO_4$ and the solvent was removed under reduced pressure. The crude product was purified by column chromatography on silica gel (AcOEt/HOAc, 99:1).

The resulting intermediate product (**SI-9**) was dissolved in in dry $CH_2Cl_2$ (20 ml) was added EDC·HCl (441 mg, 2.31 mmol, 1.50 eq.) in one portion and the mixture was purged with argon for 5 min. Afterwards NHS (265 mg, 2.31 mmol, 1.50 eq.) and DMF (1.4 ml) were added. The solution was stirred overnight at room temperature in a foil covered flask. The solution was extracted with water (8 x 20 ml). The organic layer was dried over $Na_2SO_4$ and the solvent was removed under reduced pressure. The product (**SI-10**) was isolated as light brown, sticky oil (499 mg, 1.16 mmol, 68%, 3 steps).

**$^{1}$H-NMR** (400 MHz, $D_2O$, δ/ppm): 6.23 (s, 1H), 3.51 (td, $J$ = 6.5, 6.5 Hz, 2H), 2.84 (s, 4H), 2.69 (t, $J$ = 6.8 Hz, 2H), 1.93 – 1.82 (m, 2H), 1.81 – 1.71 (m, 2H). **$^{13}$C{$^{1}$H}-NMR** (101 MHz, $D_2O$, δ/ppm): 169.32, 168.56, 39.64, 30.80, 28.22, 25.73, 21.92.* **$^{19}$F-NMR** (377 MHz, $CDCl_3$, δ/ppm): –140.73 - –140.92 (m), –150.49 - –150.67 (m). **HRMS** (ESI) m/z for $C_{16}H_{14}O_5N_5F_4$

$[M-H]^+$: calcd. 432.0926, found 432.0926. **R*f*** ($SiO_2$, AcOEt/HOAc, 99:1): 0.88. *5 carbon signals are not detectable due to $^{13}C$-$^{19}F$-couplings.

**Synthesis of linker structure SI-11**

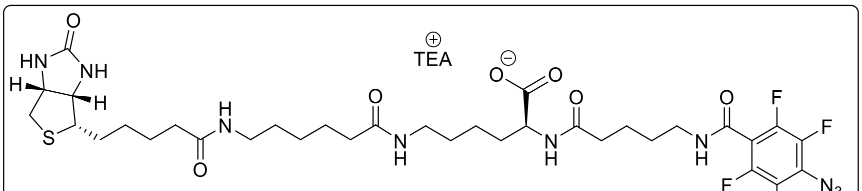


SI-11

Compound **SI-6** (18.2 mg, 0.038 µmol) was dissolved in $NaHCO_3$-solution (0.1 M, 2.75 ml) and compound **SI-10** (18.0 mg, 0.038 µmol, 1.00 eq.) was dissolved in DMSO (306 µl). The two solutions were combined and stirred for 1 h at room temperature. The solution was directly applied to an automated MPLC (Interchim) system (AQ-column). Elution was performed with a MeCN/$H_2O$ (10 mM TEAA) – gradient. The product containing fractions were combined and lyophilized. The product (**SI-11**, 20.0 mg, 0.022 µmol, 66%) was isolated as white solid.

**$^1$H-NMR** (400 MHz, DMSO-$d_6$, δ/ppm): 8.89 (t, $J$ = 5.6 Hz, 1H), 7.86 (s, 1H), 7.80 (s, 1H), 7.73 (t, $J$ = 5.6 Hz, 1H), 6.44 (s, 1H), 6.35 (s, 1H), 4.30 (dd, $J$ = 7.8, 5.2 Hz, 1H), 4.15 – 4.08 (m, 1H), 4.08 – 4.02 (m, 1H), 3.24 (q, $J$ = 6.4 Hz, 3H), 3.12 – 3.06 (m, 1H), 3.03 – 2.94 (m, 4H), 2.82 (dd, $J$ = 12.4, 5.1 Hz, 1H), 2.59 – 2.55 (m, 1H), 2.52 (d, $J$ = 1.9 Hz, 3H), 2.17 – 2.11 (m, 2H), 2.02 (dt, $J$ = 9.2, 7.5 Hz, 4H), 1.72 – 1.15 (m, 20H). **$^{19}$F-NMR** (377 MHz, DMSO-$d_6$, δ/ppm): –143.05 - –143.20 (m), –151.49 - –151.64 (m). **HRMS** (ESI) m/z for $C_{34}H_{48}O_7N_9F_4S$ $[M-H]^+$: calcd. 802.3328, found 802.3326.

**Activation of acid SI-11 as NHS-ester**

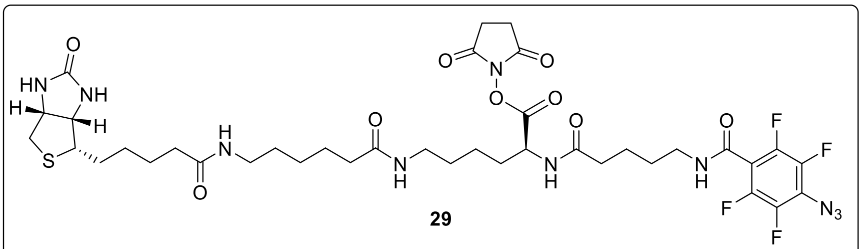


29

Carboxylic acid **SI-11** (20.0 mg, 24.1 µmol) and NHS (5.54 mg, 48.2 µmol, 2.0 eq.) were dissolved in DMF (1.0 ml) and DCC (14.9 mg, 72.3 µmol, 3.0 eq.) was added. The reaction mixture was stirred for 2 h and the product was isolated by precipitation with $Et_2O$ (10 ml) and drying under high vacuum. The isolated material was a mixture of starting material and product (29) as rapidly analyzed by HPLC. (see attachment) It was subjected to follow-up reactions immediately and without further purification.

#### 2.3. Syntheses of base-modified amino-ppGpp

**Synthesis of amino-pGp (5)**

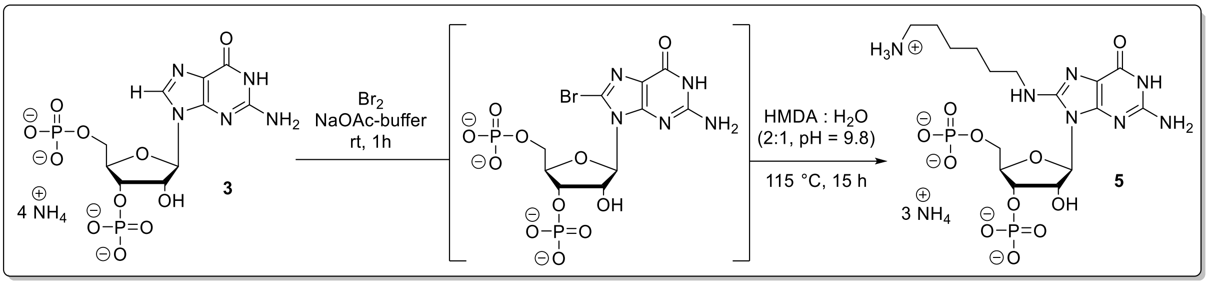


pGp x 4 $NH_4$ (**3**, 600 mg, 1.17 mmol) was dissolved in NaOAc – buffer (500 mM, pH 4.0, 22.0 ml). A solution of $Br_2$ (211 µl, 658 mg, 4.11 mmol, 3.5 eq.) in $H_2O$ (13.6 ml) was added. The resulting mixture was stirred for 1 h at rt. Afterwards $NaS_2O_3$ – buffer was added and the solution was stirred for 20 min at rt. The product was precipitated by adding EtOH (300 mHl). After incubating the suspension at 0°C for 30 min, the precipitate was separated by centrifugation and washed with EtOH (2 x 80 ml) and dried under high vacuum.

The resulting solid was dissolved in mixture of hexamethylenediamine and water (2:1, pH 9.8, 7.5 ml. The resulting solution was stirred in a pressure vessel at 115°C for 15 hours. The solution was cooled down to rt and the product was precipitated with EtOH (200 ml). After incubating the suspension at 0°C for 30 min, the precipitate was separated by centrifugation and washed with EtOH (2 x 40 ml). The crude product was dried under high vacuum and purified by automated SAX (Äkta pure system, Q-Sepharose®). Elution was performed with $NH_4HCO_3$ – buffer (300 mM). Product containing fractions were combined and lyophilized. The product (**5**, 534 mg, 876 µmol, 75 %) was isolated as white solid.

**$^1$H-NMR** (400 MHz, $D_2O$, δ/ppm): 5.91 (d, $J$ = 7.6 Hz, 1H), 4.88 – 4.83 (m, 2H), 4.54 – 4.44 (m, 1H), 4.22 (dd, $J$ = 11.8, 3.0 Hz, 1H), 4.11 (dd, $J$ = 11.8, 2.6 Hz, 1H), 3.51 – 3.34 (m, 2H), 2.99 (dd, $J$ = 7.4, 7.4 Hz, 2H), 1.76 – 1.61 (m, 5H), 1.49 – 1.35 (m, 5H). **$^{13}$C{$^1$H}-NMR** (101 MHz, $D_2O$, δ/ppm): 156.52, 152.64, 151.46, 150.36, 111.09, 86.17, 83.58 (dd, $J$ = 8.4, 3.7 Hz), 72.98 (d, $J$ = 4.8 Hz), 70.03 (d, $J$ = 4.4 Hz), 64.36 (d, $J$ = 4.6 Hz), 42.15, 39.33, 27.79,

26.43, 25.21, 25.04. **$^{31}$P{$^{1}$H}-NMR** (162 MHz, $D_2O$, δ/ppm): 0.35, -0.10. **HRMS** (ESI) m/z for $C_{16}H_{30}N_7O_{11}P_2$ [M+H]$^+$: calcd. 558.1473, found 558.1473.

**Synthesis of Fmoc-Gly-HDMA-pGp (6)**

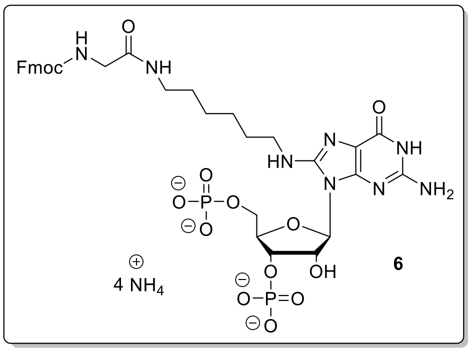


Fmoc-Gly-NHS (482 mg, 1.22 mmol, 2.0 equ.) was dissolved in DMSO (14 ml) before a solution of HMDA-pGp x 1.8 TBA (**5**, 605 mg, 612 µmol) in $H_2O$ (4.0 ml) was added. The resulting solution was stirred for 4 h at rtbefore further Fmoc-Gly-NHS (241 mg, 612 µmol, 1.0 eq.) was added. The resulting solution was stirred for 12 hours at rt. Afterwards $H_2O$ (65 ml) was added and the precipitate was separated by centrifugation. The supernatant was applied to automated SAX (Äkta, Q-Sepharose, $NH_4HCO_3$ – buffer). The product containing fractions were combined and lyophilized. The product (**6**, 357 mg, 395 µmol, 65 %) isolated as white solid.

**$^{1}$H-NMR** (400 MHz, $D_2O$, δ/ppm): 7.91 (d, *J* = 7.5 Hz, 1H), 7.77 (d, *J* = 7.7 Hz, 1H), 7.74 – 7.70 (m, 1H), 7.56 (dd, *J* = 8.2, 8.2 Hz, 1H), 7.50 (dd, *J* = 7.2, 7.2 Hz, 1H), 7.45 – 7.37 (m, 2H), 7.35 – 7.28 (m, 1H), 5.69 (d, *J* = 6.9 Hz, 1H), 4.55 (d, *J* = 5.8 Hz, 1H), 4.47 – 4.39 (m, 3H), 4.34 (t, *J* = 6.0 Hz, 1H), 4.27 – 4.14 (m, 2H), 4.11 (s, 1H), 3.72 (s, 1H), 3.61 (s, 1H), 3.36 – 3.04 (m, 4H), 1.62 – 1.43 (m, 4H), 1.39 – 1.24 (m, 4H). **$^{31}$P{$^{1}$H}-NMR** (162 MHz, $D_2O$, δ/ppm): 3.52 (s), 2.06 (d). **HRMS** (ESI) m/z for $C_{33}H_{43}N_8O_{14}P_2$ [M+H]$^+$: calcd. 837.2368, found 837.2367.

**Synthesis of Gly-HDMA-ppGpp (7)**

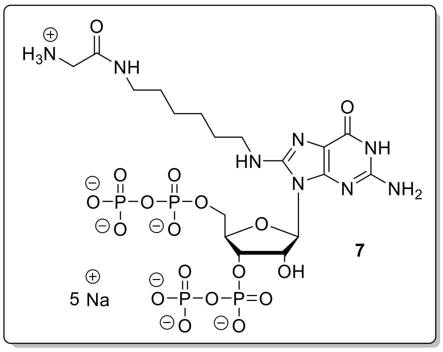

Fmoc-Gly-HDMA-pGp x 1.8 TBA (**6**, 100 mg, 78.9 µmol) was dissolved in DMF (2.0 mL) and ETT (51.3 mg, 394 µmol, 5.0 eq.) was added. Afterwards a solution of $(FmO)_2P$-N*i*$Pr_2$ (137 mg, 237 µmol, 3.0 eq.) in DMF (2.0 ml) was added. The resulting solution was stirred for 15 min at room temperature. The solution was cooled down to -20 °C and *m*CPBA (77 %, 52.9 mg, 237 µmol, 3.0 eq.) was added. The solution was stirred for 10 min at −20°C and warmed to 0°C before DBU (400 µl) was added. The resulting solution was stirred for 30 min at rt. before the product was precipitated by the addition of $Et_2O$ (40 ml). The precipitate was separated by centrifugation, washed with $Et_2O$ (2 x 10 ml) and dried under high vacuum. The crude product was purified by automated SAX (Äkta, Q-Sepharose, $NaClO_4$ – buffer). The product containing fractions were combined and precipitated using an 8-fold volume of $NaClO_4$- solution (−20 °C, 500 mM in acetone). The resulting solid was separated by centrifugation, washed with acetone (−20 °C, 2 x 3.0 ml) and dried under high vacuum for 2 h. The product (**7**, 23.8 mg, 26.9 µmol, 34 %) was isolated as white solid.

**$^1$H-NMR** (400 MHz, $D_2O$, δ/ppm): 5.89 (d, *J* = 6.8 Hz, 1H), 5.08 – 5.03 (m, 1H), 4.99 – 4.93 (m, 1H), 4.48 (ddd, *J* = 4.0, 3.8, 3.8 Hz, 1H), 4.28 – 4.22 (m, 2H), 3.65 (s, 2H), 3.47 – 3.34 (m, 2H), 3.24 (ddd, *J* = 6.5, 6.4, 2.9 Hz, 2H), 1.72 – 1.63 (m, 2H), 1.57 – 1.51 (m, 2H), 1.46 – 1.34 (m, 4H). **$^{13}$C{$^1$H}-NMR** (101 MHz, $D_2O$, δ/ppm): 156.96, 152.41, 151.87, 151.25, 151.16, 112.41, 86.38, 83.14 (m), 74.12 (m), 69.97 (m), 65.14 (m), 42.18, 41.23, 39.19, 27.99, 27.85, 25.53, 25.44.**$^{31}$P{$^1$H}-NMR** (162 MHz, $D_2O$, δ/ppm): -5.76 (d, *J* = 22.6 Hz, 1P), -6.26 (d, *J* = 21.3 Hz, 1P), -10.79 (d, *J* = 21.4 Hz, 2P). **HRMS** (ESI) m/z for $C_{18}H_{34}N_8O_{18}NaP_2$ $[M+Na]^+$: calcd. 797.0834, found 797.0835.

#### 2.4. Synthesis of pentynyl-substituted MSN

**Synthesis of pentynyl-pppGp (9)**

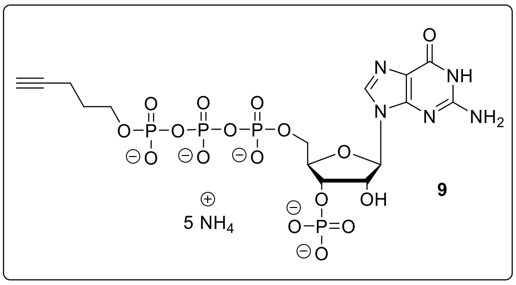


ppGp x 2.5 TBA (**8**, 200 mg, 178 µmol) was dissolved in DMF (2.5 ml) and ETT (82.9 mg, 623 µmol, 3.5 eq.) was added. Afterwards a solution of P-Amidite **SI-3** (90%, 202 mg, 445 µmol, 2.5 eq.) in DMF (2.5 ml) was added and the resulting solution was stirred for 15 min at rt. The solution was cooled to -20°C and *m*CPBA (77%, 99.5 mg, 445 µmol, 2.5 eq.) was added. After stirring for 10 min at -20°C, precipitation was induced by adding $Et_2O$ (40 ml). The resulting solid was washed with ether (2 x 15 ml) and dried under high vacuum before being dissolved in MeOH (8.0 ml). The resulting solution was stirred for 5 h at 37°C. The solvent was removed under reduced pressure and the residue was dissolved in DMF (5 ml) and piperidine (500 µl) was added at rt. After stirring the solution at rt for 30 min, precipitation was induced by the addition of $Et_2O$ (40 ml). The resulting precipitate was washed with $Et_2O$ (2 x 20 ml) and dried under high vacuum. The solid was redissolved in $H_2O$ (20 ml) and acidified with HCl to pH 5.3. Afterwards RNase T2 (50 µl) was added and the solution was incubated at 37 °C overnight. The solution was directly applied to automated SAX (Äkta pure system, Q-Sepharose®, $NH_4HCO_3$ - buffer). The product containing fractions were combined and lyophilized. The product (**9**, 64.0 mg, 82.2 µmol, 46 %) was isolated as white solid.

**$^1$H-NMR** (400 MHz, $D_2O$, δ/ppm): 8.15 (s, 1H), 5.97 (d, *J* = 7.1 Hz, 1H), 4.98 – 4.91 (m, 1H), 4.88 – 4.81 (m, 1H), 4.59 – 4.52 (m, 1H), 4.32 – 4.21 (m, 2H), 3.98 (ddd, *J* = 6.5, 6.5, 6.5 Hz, 2H), 2.26 (dd, *J* = 2.7 Hz, 1H), 2.21 (ddd, *J* = 7.3, 7.3, 2.7 Hz, 2H), 1.73 (dddd, *J* = 6.9, 6.8, 6.8 Hz, 2H). **$^{13}$C{$^1$H}-NMR** (101 MHz, $D_2O$, δ/ppm): 159.05, 153.93, 152.05, 137.89, 116.37, 86.34, 84.93, 83.41 (dd, *J* = 9.3, 3.6 Hz), 74.25 (d, *J* = 5.0 Hz), 72.74 (d, *J* = 4.9 Hz), 69.11, 65.45 (d, *J* = 5.4 Hz), 65.24 (d, *J* = 6.0 Hz), 28.79 (d, *J* = 7.3 Hz), 14.12. **$^{31}$P{$^1$H}-NMR** (162 MHz, $D_2O$, δ/ppm): 0.44, -11.05 (d, *J* = 19.3 Hz), -11.56 (d, *J* = 18.5 Hz), -23.25 (t, *J* = 19.1 Hz). **HRMS** (ESI) m/z for $C_{15}H_{21}O_{17}N_5P_4$ $[M-H_2]^{2-}$ calcd 333.4947 found 333.4947.

**Synthesis of pentynyl-ppAp (11)**

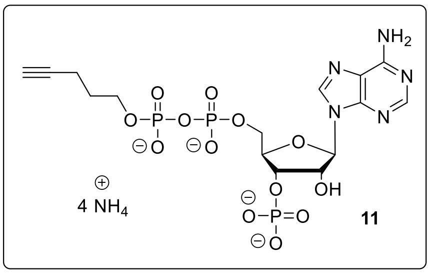

pAp x 1.87 TBA (**4**, 80.0 mg, 91.2 µmol) was dissolved in DMF (1.0 ml) and ETT (47.4 mg, 365 µmol, 4.0 eq.) was added. Afterwards a solution of P-Amidite **SI-3** (90%, 104 mg, 228 µmol, 2.5 eq.) in DMF (2.0 ml) was added and the resulting solution was stirred for 15 min at rt. The solution was cooled to -20°C and *m*CPBA (77%, 61.1 mg, 273 µmol, 3.0 eq.) was added. After stirring for 10 min at -20 °C, precipitation was induced by $Et_2O$ (40 ml). The resulting solid was washed with $Et_2O$ (2 x 15 ml) and dried under high vacuum before being dissolved in MeOH (3.0 ml). The resulting solution was stirred for 3 h at 37 °C. The solvent was removed under reduced pressure and the residue was dissolved in DMF (2.5 ml) before piperidine (250 µl) was added at rt. After stirring the solution at rt for 30 min, precipitation was induced by the addition of $Et_2O$ (20 ml). The resulting precipitate was washed with ether (2 x 10 ml) and dried under high vacuum. The solid was redissolved in $H_2O$ (10 ml) and acidified with HCl to pH 5.3. Afterwards RNase T2 (50 µl) was added and the solution was incubated at 37 °C overnight. The solution was directly applied to automated SAX (Äkta pure system, Q-Sepharose®, $NH_4HCO_3$ - buffer). The product containing fractions were combined and lyophilized. The product (**11**, 43.0 mg, 67.1 µmol, 74 %) was isolated as white solid.

**$^1$H-NMR** (400 MHz, $D_2O$, δ/ppm): 8.61 (s, 1H), 8.33 (s, 1H), 6.21 (d, *J* = 6.6 Hz, 1H), 4.94 – 4.88 (m, 1H), 4.88 – 4.83 (m, 1H), 4.61 (dddd, *J* = 2.7, 2.7, 2.7 Hz, 1H), 4.28 – 4.22 (m, 2H), 3.95 (ddd, *J* = 6.5, 6.5, 6.5 Hz, 2H), 2.25 – 2.16 (m, 3H), 1.73 (dddd, *J* = 6.8, 6.7, 6.7 Hz, 2H). **$^{13}$C{$^1$H}-NMR** (101 MHz, $D_2O$, δ/ppm): 153.95, 150.46, 149.12, 140.63, 118.63, 86.52, 84.82, 83.51 (dd, J = 9.2, 3.6 Hz), 74.23 (d, J = 5.1 Hz), 73.67 (d, J = 5.1 Hz), 69.15, 65.27 – 65.07 (m, 2C), 28.75 (d, J = 7.3 Hz), 14.07. **$^{31}$P{$^1$H}-NMR** (162 MHz, $D_2O$, δ/ppm): -0.14, -10.92 (d, J = 21.1 Hz), -11.55 (d, J = 21.1 Hz). **HRMS** (ESI) m/z for $C_{15}H_{21}O_{13}N_5P_3$ $[M-H]^-$ calcd 572.0354 found 572.0350.

**Synthesis of pentynyl-pGp (12)**

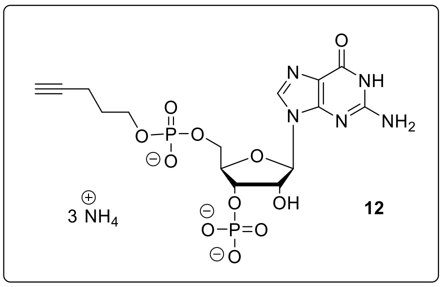

Guanosine dihydrate (**1**, 50.0 mg, 157 µmol), ETT (81.5 mg, 627 µmol, 4.0 eq.) and P-Amidite **SI-3** (121 mg, 266 µmol, 1.7 eq.) were coevaporated separately with dry MeCN (3 x 3.0 ml). Afterwards, a solution of ETT in DMSO (400 µl) and DMF (100 µl) was added to Guanosine and the resulting solution was cooled to 3 °C. Subsequently, a solution of P-Diamidite **SI-2** in DMF (3°C, 500 µl) was added. The reaction mixture was stirred for 1 h at 0 °C before (FmO)P-(N*i*Pr$_2$)$_2$ was added and the solution was stirred for 45 min at rt. The solution was cooled to -20°C and TBHP (5.5 M in decanes, 94.0 µl, 517 µmol, 3.3 eq.) was added dropwise. The solution was warmed to rt and stirred for 1 h. Afterwards, piperidine (90.0 µl) was added and the solution was stirred for 30 min at rt. The intermediate product was precipitated by the addition of $Et_2O$ (10 ml). The precipitate was separated by centrifugation, $Et_2O$ (3 x 5 ml) and dried under high vacuum. The solid was dissolved in $H_2O$ (5 ml) and RNase T2 (50 µl) was added. The solution was incubated for 12 h at 37 °C and directly applied to automated SAX (Äkta pure system, Q-Sepharose®, $NH_4HCO_3$ - buffer). The product containing fractions were combined and lyophilized. The product (**12**, 38.0 mg, 67.8 µmol, 43 %) was isolated as white solid.

**$^1$H-NMR** (400 MHz, $D_2O$, δ/ppm): 8.09 (s, 1H), 5.92 (d, $J$ = 6.2 Hz, 1H), 4.90 (dd, $J$ = 5.7, 5.7 Hz, 1H), 4.86 – 4.79 (m, 1H), 4.48 – 4.42 (m, 1H), 4.11 – 4.00 (m, 2H), 3.81 – 3.68 (m, 2H), 2.16 (dd, $J$ = 2.7, 2.7 Hz, 1H), 2.11 – 2.01 (m, 2H), 1.61 (dddd, $J$ = 6.6 Hz, 2H). **$^{13}$C{$^1$H}-NMR** (101 MHz, $D_2O$, δ/ppm): 158.84, 153.98, 151.91, 137.57, 116.10, 86.78, 84.48, 83.13 (dd, J = 8.8, 3.7 Hz), 73.93 (d, J = 5.1 Hz), 72.58 (d, J = 4.9 Hz), 69.18, 64.86 (d, J = 5.3 Hz), 64.66 (d, J = 5.7 Hz), 28.60 (d, J = 7.6 Hz), 13.99. **$^{31}$P{$^1$H}-NMR** (162 MHz, $D_2O$, δ/ppm): 0.24, -0.16. **HRMS** (ESI) m/z for $C_{15}H_{20}O_{11}N_5P_2$ [M-H]$^-$ calcd 508.0640 found 508.0644.

**Synthesis of pentynyl-pGpp (13)**

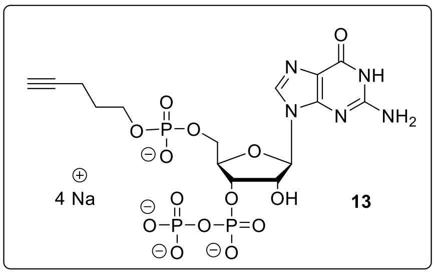

Pentynyl-pGp x 1.75 TBA (**12**, 35.0 mg, 37.6 µmol) was dissolved in DMF (1.0 ml) and ETT (14.6 mg, 113 µmol, 3.0 eq.) was added. Afterwards a solution of $(FmO)_2P$-N*i*$Pr_2$ (**SI-1**, 90%, 32.6 mg, 56.3 µmol, 1.5 eq.) in DMF (2.5 ml) was added and the resulting solution was stirred for 15 min at rt. The solution was cooled to -20°C and *m*CPBA (77%, 12.6 mg, 56.3 µmol, 1.5 eq.) was added. After stirring for 10 min at -20°C, precipitation was induced by adding $Et_2O$ (40 ml). The resulting solid was washed with $Et_2O$ (2 x 15 ml) and dried under high vacuum. The crude product was purified by automated SAX (Äkta pure system, Q-Sepharose®, $NaClO_4$ - buffer). The product containing fractions were precipitated by $NaClO_4$ – solution (0.5 M in acetone, -20 °C, 40 ml), and the precipitate was separated by centrifugation, washed with acetone (-20°C, 3 x 10 ml) and dried under high vacuum. The product (**13**, 18.0 mg, 26.7 µmol, 71%) was isolated as white solid.

**$^1$H-NMR** (400 MHz, $D_2O$, δ/ppm): 8.10 (s, 1H), 5.99 (d, *J* = 5.4 Hz, 1H), 5.03 (ddq, *J* = 28.3, 9.0, 4.6 Hz, 1H), 4.95 (dd, *J* = 5.3, 5.3 Hz, 1H), 4.49 (br.s , *J* = 4.1 Hz, 1H), 4.18 – 4.08 (m, 2H), 3.83 – 3.71 (m, 2H), 2.12 – 2.05 (m, 2H), 1.69 – 1.59 (m, 2H). **$^{13}$C{$^1$H}-NMR** (101 MHz, $D_2O$, δ/ppm): 159.10, 153.91, 151.97, 137.91, 116.48, 87.35, 83.09 – 82.87 (m), 74.43 (d, *J* = 5.2 Hz), 72.65 (d, *J* = 3.6 Hz), 65.11 (d, *J* = 5.1 Hz), 64.63 (d, *J* = 5.6 Hz), 28.60 (d, *J* = 7.4 Hz), 13.94. **$^{31}$P{$^1$H}-NMR** (162 MHz, $D_2O$, δ/ppm): 0.24, -5.90 (d, *J* = 22.7 Hz), -11.08 (d, *J* = 22.8 Hz). **HRMS** (ESI) m/z for $C_{15}H_{21}O_{14}N_5P_3$ $[M-H]^-$ calcd 588.0303 found 588.0300.

**Synthesis of pentynyl-pppGpp (15)**

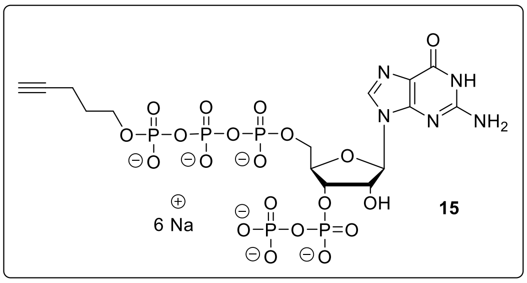

Pentynyl-pppGp x 3.2 TBA (**9**, 90.0 mg, 62.6 µmol) was dissolved in DMF (1.0 ml) and ETT (24.4 mg, 187 µmol, 3.0 eq.) was added. Afterwards a solution of $(FmO)_2P$-N*i*$Pr_2$ (**SI-1**, 90%, 54.3 mg, 93.9 µmol, 1.5 eq.) in DMF (2.5 ml) was added and the resulting solution was stirred for 15 min at rt. The solution was cooled to -20°C and *m*CPBA (21.0 mg, 93.9 µmol, 1.5 eq.) was added. After stirring for 10 min at -20°C, precipitation was induced by $Et_2O$ (40 ml). The resulting solid was washed with ether (2 x 15 ml) and dried under high vacuum. The crude product was purified by automated SAX (Äkta pure system, Q-Sepharose®, $NaClO_4$ - buffer). The product containing fractions were precipitated by $NaClO_4$ – solution (0.5 M in acetone, -20°C, 40 ml), and the precipitate was separated by centrifugation, washed with acetone (-20°C, 3 x 10 ml) and dried under high vacuum. The product (**15**, 41.0 mg, 46.5 µmol, 74%) was isolated as white solid.

**$^1$H-NMR** (400 MHz, $D_2O$, δ/ppm): 8.14 (s, 1H), 6.00 (d, *J* = 6.3 Hz, 1H), 5.01 – 4.93 (m, 1H), 4.89 (dd, *J* = 5.7, 5.7 Hz, 1H), 4.58 – 4.49 (m, 1H), 4.36 – 4.18 (m, 2H), 3.95 (ddd, *J* = 6.5, 6.5, 6.5 Hz, 2H), 2.26 (dd, *J* = 2.7, 2.7 Hz, 1H), 2.18 (ddd, *J* = 7.3, 7.3, 2.7 Hz, 2H), 1.69 (dddd, *J* = 7.9, 7.9, 7.4, 7.4 Hz, 2H). **$^{13}$C{$^1$H}-NMR** (101 MHz, $D_2O$, δ/ppm): 159.14, 153.95, 151.98, 137.98, 116.38, 86.88, 83.27 (dd, *J* = 9.2, 4.3 Hz), 74.86 (d, *J* = 5.2 Hz), 72.99 (d, *J* = 4.2 Hz), 65.57 (d, *J* = 5.3 Hz), 65.22 (d, *J* = 6.0 Hz), 28.78 (d, *J* = 7.4 Hz), 14.07. **$^{31}$P{$^1$H}-NMR** (162 MHz, $D_2O$, δ/ppm): -5.72 (d, *J* = 22.2 Hz), -10.73 – -11.00 (m, 2P), -11.34 (d, *J* = 17.9 Hz), -22.84 (dd, *J* = 18.5 Hz). **HRMS** (ESI) for $C_{15}H_{23}O_{20}N_5P_5$ [M-H]$^-$ calcd 747.9630 found 747.9636.

**Synthesis of pentynyl-ppApp (16)**

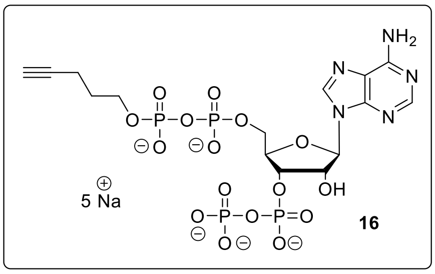


Pentynyl-ppAp x 1.89 TBA (**11**, 45.0 mg, 43.8 µmol) was dissolved in DMF (1.0 ml) and ETT (20.0 mg, 154 µmol, 3.5 eq.) was added. Afterwards a solution of $(FmO)_2P$-N*i*$Pr_2$ (**SI-1**, 80%, 42.8 mg, 65.8 µmol, 1.5 eq.) in DMF (2.0 ml) was added and the resulting solution was stirred for 15 min at rt. The solution was cooled to -20°C and *m*CPBA (77%, 19.6 mg, 87.7 µmol, 2.0 eq.) was added. After stirring for 10 min at -20°C, precipitation was induced by $Et_2O$ (40 ml). The resulting solid was washed with ether (2 x 15 ml) and dried under high vacuum. The crude product was purified by automated SAX (Äkta pure system, Q-Sepharose®, $NaClO_4$ - buffer). The product containing fractions were precipitated by $NaClO_4$ – solution (0.5 M in acetone, -20°C, 40 ml) and the precipitate was separated by centrifugation, washed with acetone (-20°C, 3 x 10 ml) and dried under high vacuum. The product (**16**, 28.3 mg, 37.1 µmol, 84%) was isolated as white solid.

**$^1$H-NMR** (400 MHz, $D_2O$, δ/ppm): 8.56 (s, 1H), 8.28 (s, 1H), 6.21 (d, *J* = 6.2 Hz, 1H), 4.99 – 4.92 (m, 1H), 4.88 (dd, *J* = 5.6, 5.6 Hz, 1H), 4.61 – 4.54 (m, 1H), 4.30 – 4.18 (m, 2H), 3.86 (ddd, *J* = 6.5, 6.3, 6.3 Hz, 2H), 2.19 (dd, *J* = 2.7, 2.7 Hz, 1H), 2.15 – 2.09 (m, 2H), 1.63 (dddd, *J* = 6.8, 6.7, 6.7 Hz, 3H). **$^{13}$C{$^1$H}-NMR** (101 MHz, $D_2O$, δ/ppm): δ 155.68, 152.93, 149.35, 139.97, 118.68, 86.66, 84.75, 83.36 (dd, *J* = 9.4, 4.1 Hz), 74.72 (d, *J* = 5.1 Hz), 73.56 (d, *J* = 4.3 Hz), 69.04, 65.31 (d, *J* = 5.7 Hz), 65.03 (d, *J* = 6.0 Hz), 28.72 (d, *J* = 7.4 Hz), 14.01. **$^{31}$P{$^1$H}-NMR** (162 MHz, $D_2O$, δ/ppm): -5.85 (d, *J* = 22.6 Hz), -10.58 – -11.14 (m, 2P), -11.55 (d, *J* = 20.5 Hz). **HRMS** (ESI, m/z) for $C_{15}H_{21}O_{16}N_5P_4$ $[M-H_2]^{2-}$ calcd 325.4972 found 325.4973.

#### 2.5. Synthesis of phosphate-modified amino-MSN

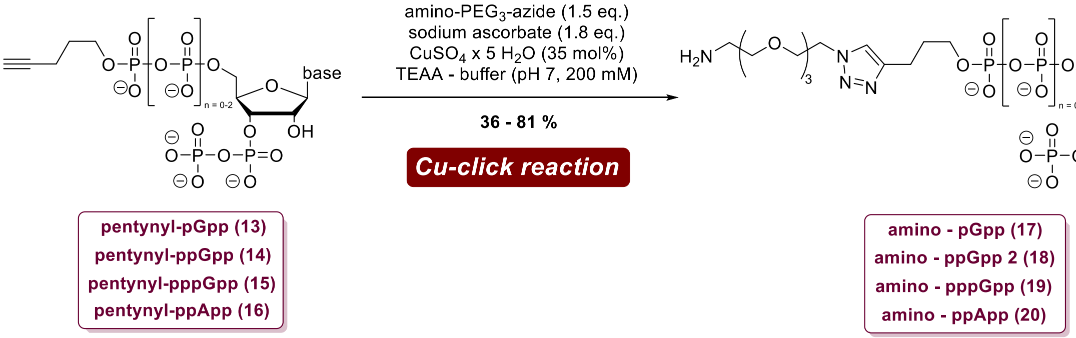


**Supporting figure 2:** Synthesis of phosphate-modified amino-MSN from pentynyl-MSN by Cu-catalyzed cycloaddition reactions.

**General procedure A** for Cu-click reactions towards amino-MSN **17-20**:

Pentynyl-MSN (10 – 25 µmol) was dissolved in TEAA-buffer (200 mM, 1.5 ml). Amino-PEG3-azide (1.5 eq.) and sodium ascorbate (1.8 eq.) were added and argon was flown through the solution for 20 min at rt. Afterwards $CuSO_4$ x 5 $H_2O$ (0.35 eq.) was added and the solution was stirred over argon atmosphere for 2-3 h. Afterwards $H_2O$ (17 ml) was added and the solution was directly applied to automated SAX (Äkta pure system, Q-Sepharose®, $NaClO_4$ - buffer). The product containing fractions were precipitated by NaClO4 – solution (0.5 M in acetone, -20°C, 40 ml) and the solid was separated by centrifugation, washed with acetone (-20°C, 3 x 10 ml) and dried under high vacuum.

As the products still contained residual amounts of $Cu^{II}$-ions, NMR-analysis was not possible at this stage. The $Cu^{II}$-impurities were removed in subsequent steps allowing NMR-characterization of the final capture compounds.

##### Synthesis of amino-pGpp (17)

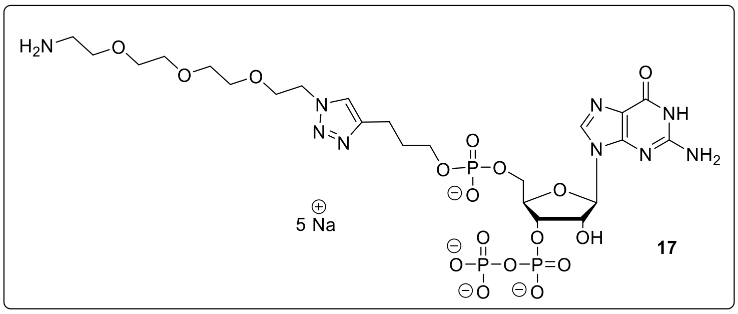


Pentynyl-pGpp (**13**, 10.0 mg, 14.7 µmol) was subjected to the general **procedure A**. The product (**17**, 4.80 mg, 5.36 µmol, 36%) was isolated as white solid.

**HRMS** (ESI) m/z for $C_{23}H_{41}N_9O_{17}P_3$ $[M+H]^+$: calcd 808.1828, found 808.1816. **HPLC-UV** analysis (see SI, chapter 9) was performed.

**Synthesis of amino-pppGpp (19)**

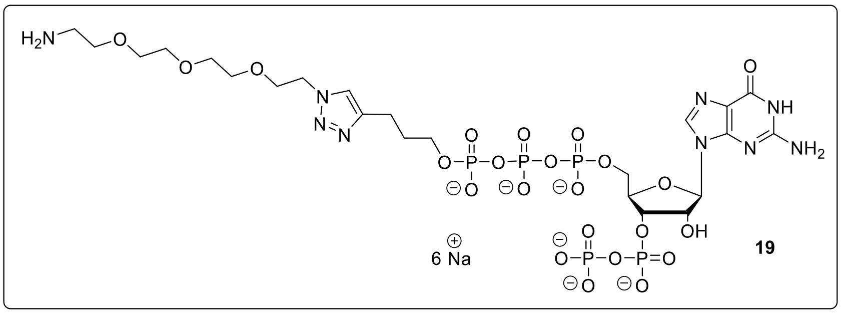


Pentynyl-pppGpp (**15**, 15.0 mg, 17.0 µmol) was subjected to the general **procedure A**. The product (**19**, 15.2 mg, 13.8 µmol, 81%) was isolated as white solid.

**HRMS** (ESI, m/z) m/z for $C_{23}H_{40}N_9O_{23}P_5$ $[M\text{-}H_2]^{2-}$: calcd 482.5468, found 482.5468. **HPLC-UV** analysis (see appendix) was performed.

**Synthesis of amino-ppApp (20)**

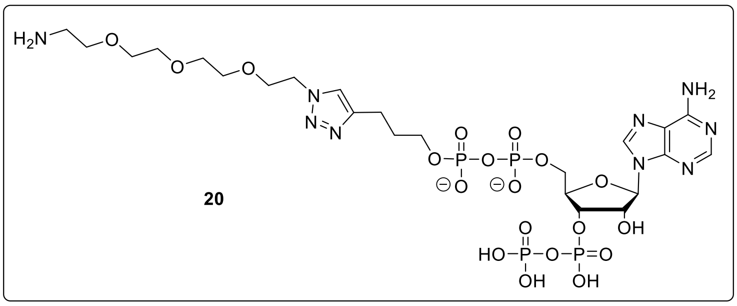


Pentynyl-ppApp (**16**, 10.0 mg, 13.1 µmol) was subjected to the general **procedure A**. The product (**20**, 5.30 mg, 5.40 µmol, 41%) was isolated as white solid.

**HRMS** (ESI) m/z for $C_{23}H_{39}N_9O_{19}P_4$ $[M\text{-}H_2]^{2-}$: calcd 434.5662, found 434.5671. **HPLC-UV** analysis (see SI, chapter 9) was performed.

**Synthesis of amino-ppGp (SI-12)**

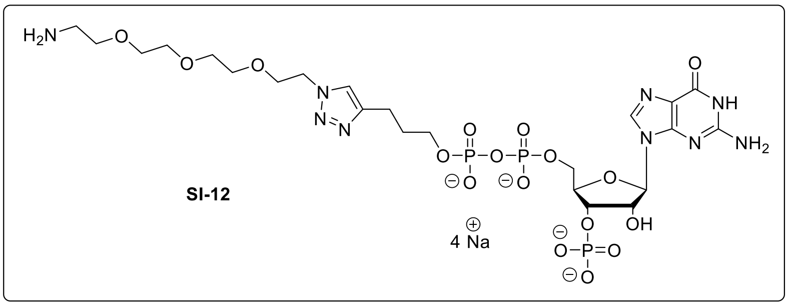


Pentynyl-ppGp (**10**, 15.0 mg, 22.2 µmol) was subjected to the general **procedure A**. The product (**SI-12**, 15.3 mg, 17.0 µmol, 77%) was isolated as white solid.

**HRMS** (ESI) m/z for $C_{23}H_{39}N_{9}O_{17}P_{3}$ $[M-H]^{-}$: calcd 806.1682, found 806.1686. **HPLC-UV** analysis (see SI, chapter 9) was performed.

#### 2.6. Syntheses of MSN-capture compounds

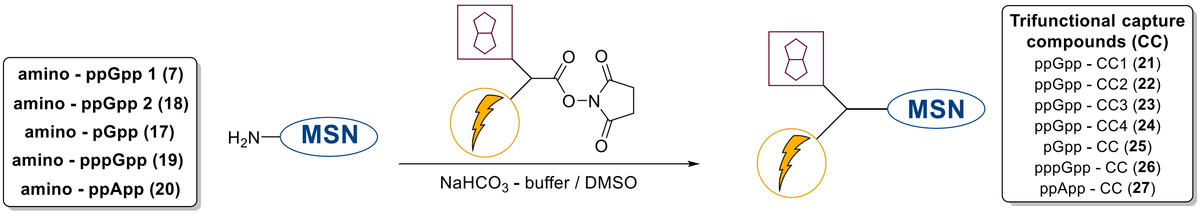


**Supporting figure 3:** Synthesis of trifunctional MSN-capture compounds from amino-MSN and NHS-esters by chemoselective amide forming reactions.

**General procedure B** for NHS – Ester coupling towards MSN-capture compounds **21-27**:

Amino – MSN (0.7 – 3.4 µmol) were dissolved in $NaHCO_3$ – buffer (200 mM) in a concentration of 10 µg/µl. Sulfo-NHS – Ester (1.2 – 1.5 eq.) were dissolved in $H_2O$ in a concentration of 10 µg/µl. The solutions were mixed and the resulting reaction mixture was incubated for 30 min at rt under light exclusion. Reaction progress was monitored by HPLC. Afterwards the mixture was diluted with $H_2O$ (6.0 ml) and directly applied to automated SAX (Äkta pure system, Q-Sepharose®, $NaClO_4$ – buffer, VIS-detection at 600 nm). The product containing fractions were determined by HPLC and precipitated by $NaClO_4$ – solution (0.5 M in acetone, -20°C, 40 ml). The solid was separated by centrifugation, washed with acetone (-20°C, 3 x 10 ml) and dried under high vacuum. In contrast to Sulfo-NHS esters, NHS-esters were dissolved in DMSO before mixing.

**Synthesis of ppGpp – CC1 (21)**

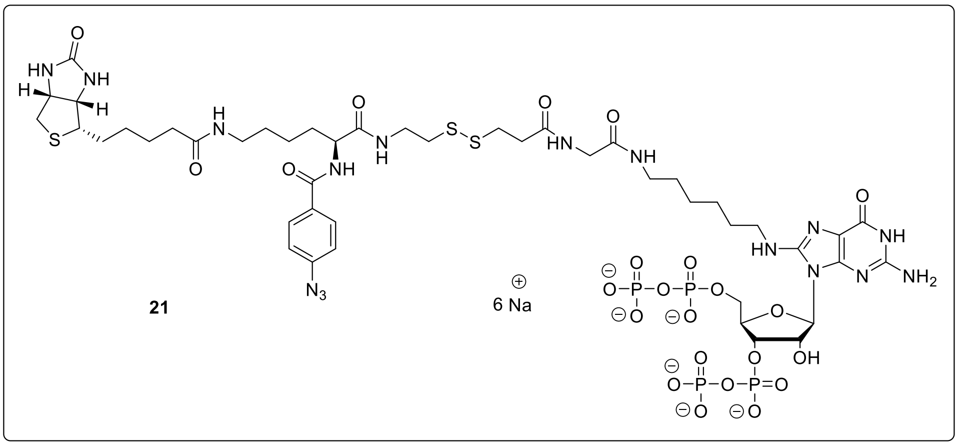

CC2 was synthesized from 8‘-amino-ppGpp (**7**, 1.21 mg, 1.36 µmol) and Sulfo-SBED according to the general **procedure B**. The product (**21**, 1.30 mg, 829 nmol, 61%) was isolated as white solid.

**$^{1}$H-NMR** (400 MHz, $D_2O$, δ/ppm): 7.99 – 7.92 (m, 2H), 7.33 – 7.27 (m, 2H), 5.96 (d, *J* = 6.5 Hz, 1H), 4.86 – 4.80 (m, 1H), 4.58 (dd, *J* = 9.3, 5.8 Hz, 1H), 4.52 – 4.30 (m, 2H), 3.97 (s, 2H), 3.69 (dd, *J* = 12.9, 6.9 Hz, 2H), 3.47 (d, *J* = 11.3 Hz, 2H), 3.41 – 3.23 (m, 6H), 3.06 (ddd, *J* = 14.1, 10.4, 5.3 Hz, 6H), 2.94 – 2.77 (m, 3H), 2.35 (dd, *J* = 4.4 Hz, 2H), 2.30 (dd, *J* = 7.3, 7.3 Hz, 2H), 2.11 – 1.91 (m, 2H), 1.89 – 1.35 (m, 17H). **$^{31}$P{$^{1}$H}-NMR** (162 MHz, $D_2O$, δ/ppm): -5.75 (br. s), -10.61 (br.s). **HRMS** (ESI) m/z for $C_{46}H_{73}N_{16}O_{23}P_4S_3$ $[M+H]^+$: calcd 1437.3142, found 1437.3147. **HPLC-UV** analysis was performed (see SI, chapter 9).

**Synthesis of ppGpp – CC2 (22)**

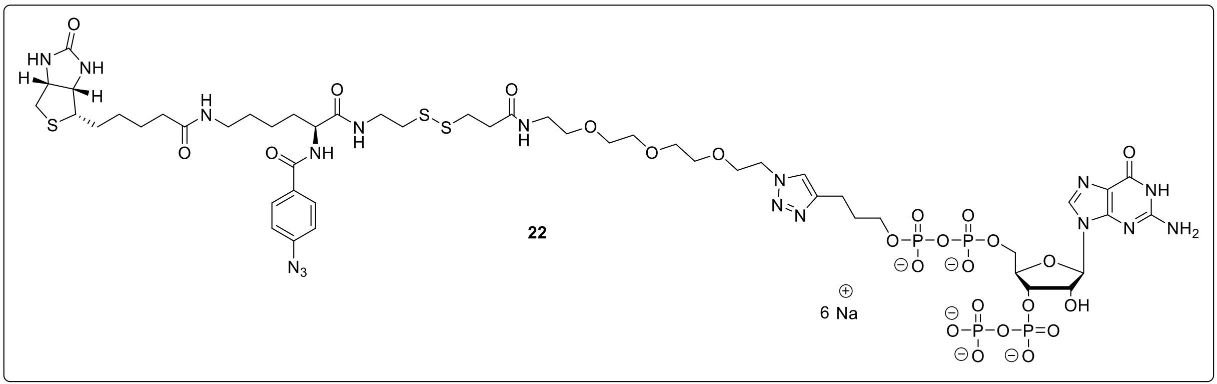


CC2 was synthesized from amino-ppGpp (**18**, 770 µg, 772 nmol) and Sulfo-SBED according to the general **procedure B**. The product (**22**, 370 µg, 223 nmol, 29%) was isolated as white solid.

**$^{1}$H-NMR** (400 MHz, $D_2O$, δ/ppm): 8.10 (s, 1H), 7.81 (d, *J* = 8.5 Hz, 2H), 7.67 (s, 1H), 7.17 (d, *J* = 8.5 Hz, 2H), 5.94 (d, *J* = 6.1 Hz, 1H), 5.05 – 4.93 (m, 3H), 4.61 – 4.49 (m, 4H), 4.45 (dd, *J* = 8.9, 5.8 Hz, 1H), 4.37 – 4.19 (m, 3H), 3.92 (dd, *J* = 5.1, 5.1 Hz, 2H), 3.87 (ddd, *J* = 6.4, 6.4, 6.4 Hz, 2H), 3.65 – 3.51 (m, 12H), 3.36 (dd, *J* = 5.3, 5.3 Hz, 2H), 3.21 (q, *J* = 4.8 Hz, 3H), 2.97 – 2.84 (m, 5H), 2.73 (d, *J* = 13.0 Hz, 1H), 2.66 – 2.52 (m, 4H), 2.18 (t, *J* = 7.2 Hz, 2H), 1.98 – 1.84 (m, 2H), 1.78 (p, *J* = 7.2 Hz, 2H), 1.70 – 1.37 (m, 8H), 1.36 – 1.20 (m, 2H). **$^{31}$P{$^{1}$H}-NMR** (162 MHz, $D_2O$, δ/ppm): -5.80 (d, *J* = 20.8 Hz), -10.71 – -11.11 (m), -11.57 (d, *J* = 20.9 Hz). **HRMS** (ESI) m/z for $C_{51}H_{77}N_{17}O_{25}P_4S_3$ $[M-H_2]^{2-}$: calcd 773.6700, found 773.6704. **HPLC-UV** analysis was performed (see attachment).

**Synthesis of ppGpp – CC3 (23)**

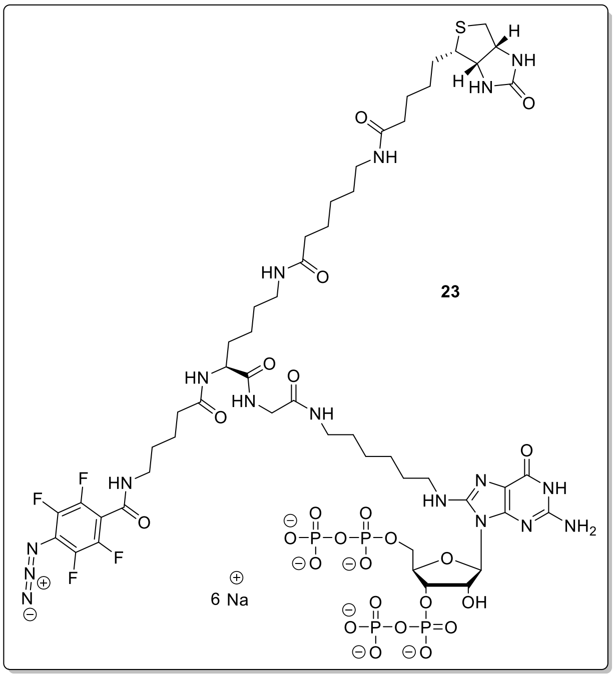

ppGpp – CC5 was synthesized from 8-amino-ppGpp (**7**, 3.00 mg, 3.39 µmol) and the synthesized Fluorophenylazide-NHS - linker **29** according to the general **procedure B**. The product (**23**, 1.10 mg, 650 nmol, 19%) was isolated as white solid.

**$^1$H-NMR** (400 MHz, $D_2O$, δ/ppm): 5.85 (d, *J* = 6.8 Hz, 1H), 5.05 (s, 1H), 4.59 (dd, *J* = 8.0, 4.8 Hz, 1H), 4.47 (t, *J* = 3.6 Hz, 1H), 4.39 (dd, *J* = 8.0, 4.4 Hz, 1H), 4.33 – 4.15 (m, 3H), 3.87 (d, *J* = 3.7 Hz, 2H), 3.48 – 3.08 (m, 12H), 2.98 (dd, *J* = 13.1, 5.0 Hz, 1H), 2.77 (d, *J* = 13.0 Hz, 1H), 2.37 (dd, *J* = 7.3, 7.3 Hz, 2H), 2.27 – 2.18 (m, 2H), 1.80 – 1.23 (m, 32H). **$^{19}$F{$^1$H}-NMR** (377 MHz, δ/ppm): -142.95 – -143.21 (m, 2F), -150.98 – -151.26 (m, 2F). **$^{31}$P{$^1$H}-NMR** (162 MHz, $D_2O$, δ/ppm): -5.79 (d, J = 22.2 Hz, 2P), -10.81 (d, J = 20.4 Hz, 2P). **HRMS** (ESI) m/z for $C_{52}H_{80}N_{17}O_{24}F_4P_4S$ [M+H]$^+$: calcd 1558.4164, found 1558.4139. **HPLC-UV** analysis was performed (see SI, chapter 9).

**Synthesis of ppGpp - CC4 (24)**

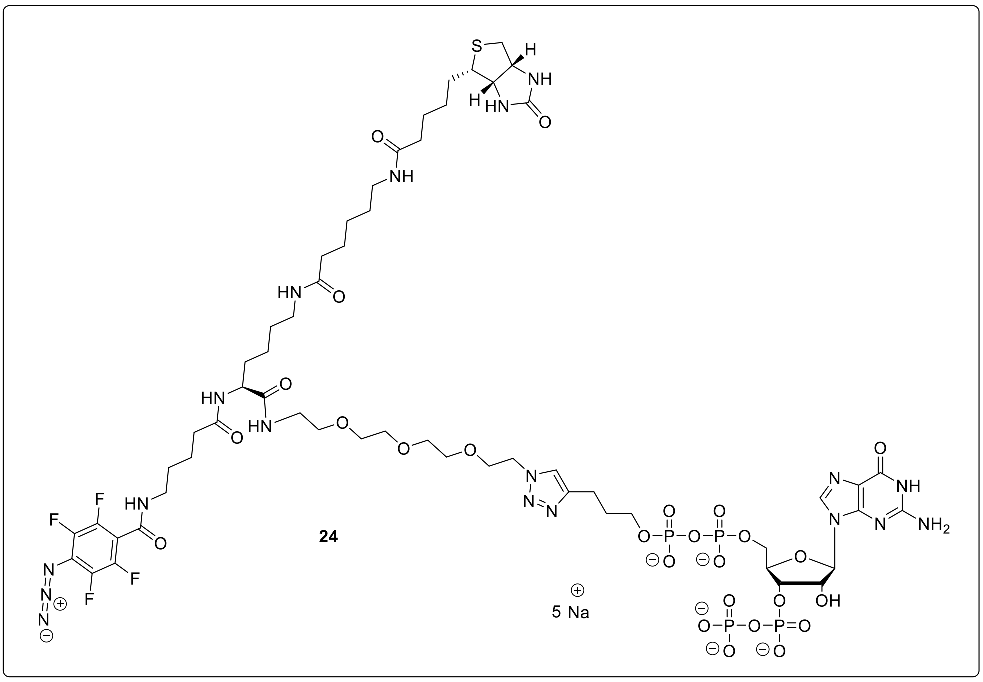


ppGpp – CC3 was synthesized from amino-ppGpp (**18**, 3.00 mg, 3.00 µmol) and the synthesized Fluorophenylazide-NHS - linker **29** according to the general **procedure B**. The product (**24**, 2.20 mg, 1.24 µmol, 41%) was isolated as white solid.

**$^{1}$H-NMR** (400 MHz, $D_2O$, δ/ppm): $^{1}$H NMR (400 MHz, $D_2O$) δ 8.11 (s, 1H), 7.70 (s, 1H), 5.95 (d, *J* = 6.2 Hz, 1H), 4.63 – 4.58 (m, 1H), 4.59 – 4.48 (m, 2H), 4.46 – 4.34 (m, 1H), 4.33 – 4.15 (m, 3H), 4.00 – 3.79 (m, 4H), 3.74 – 3.48 (m, 12H), 3.48 – 3.35 (m, 4H), 3.35 – 3.26 (m, 1H), 3.24 – 3.06 (m, 4H), 2.98 (dd, *J* = 13.0, 5.0 Hz, 1H), 2.77 (d, *J* = 13.0 Hz, 1H), 2.59 (dd, *J* = 8.0, 8.0 Hz, 2H), 2.45 – 2.30 (m, 3H), 1.92 – 1.19 (m, 28H). **$^{19}$F{$^{1}$H}-NMR** (377 MHz, δ/ppm): -143.09 – -143.27 (m), -151.02 – -151.18 (m). **$^{31}$P{$^{1}$H}-NMR** (162 MHz, $D_2O$, δ/ppm): -6.06 (br.s), -10.66 – -11.24 (m), -11.61 (d, J = 21.0 Hz). **HRMS** (ESI) m/z for $C_{57}H_{83}N_{18}O_{26}$ $F_4P_4S$ $[M\text{-}H_3]^{3-}$: calcd 555.8117, found 555.8113. **HPLC-UV** analysis was performed (see SI, chapter 9).

**Synthesis of pGpp – CC (25)**

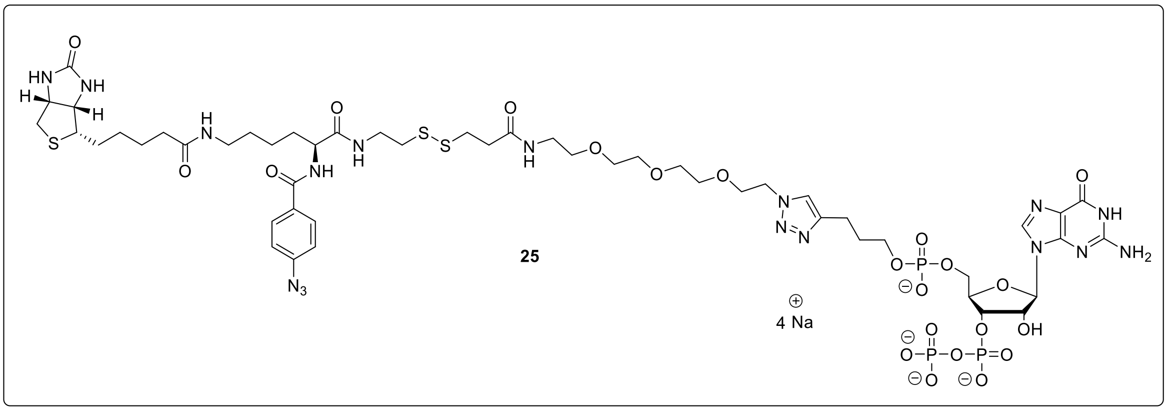

pGpp – CC5 was synthesized from amino-pGpp (**17**, 2.00 mg, 2.23 µmol) and Sulfo-SBED according to the general **procedure B**. The product (**25**, 1.35 mg, 867 nmol, 39%) was isolated as white solid.

**$^1$H-NMR** (400 MHz, $D_2O$, δ/ppm): 8.03 (s, 1H), 7.77 (d, *J* = 8.3 Hz, 2H), 7.61 (s, 1H), 7.12 (d, *J* = 8.5 Hz, 2H), 5.91 (d, *J* = 5.4 Hz, 1H), 4.98 (s, 1H), 4.57 – 4.37 (m, 4H), 4.34 – 4.24 (m, 1H), 4.18 – 4.04 (m, 2H), 3.89 (dd, *J* = 5.1, 5.1 Hz, 2H), 3.80 – 3.65 (m, 2H), 3.63 – 3.46 (m, 13H), 3.32 (dd, *J* = 5.3, 5.3 Hz, 2H), 3.23 – 3.12 (m, 3H), 2.94 – 2.80 (m, 5H), 2.70 (d, *J* = 13.1 Hz, 1H), 2.58 (dd, *J* = 6.7, 6.7 Hz, 2H), 2.56 – 2.43 (m, 2H), 2.15 (dd, *J* = 7.2, 7.2 Hz, 2H), 1.96 – 1.79 (m, 2H), 1.74 (dd, *J* = 7.8, 7.8 Hz, 2H), 1.66 – 1.34 (m, 9H), 1.34 – 1.15 (m, 2H). **$^{31}$P{$^1$H}-NMR** (162 MHz, $D_2O$, δ/ppm): 0.28, -5.90 (br. s), -11.08 (br. s). **HRMS** (ESI) m/z for $C_{51}H_{75}N_{17}O_{22}P_3S_3$ [M-H]$^{3-}$: calcd 488.7888, found 488.7890. **HPLC-UV** analysis was performed (see SI, chapter 9).

**Synthesis of pppGpp – CC (26)**

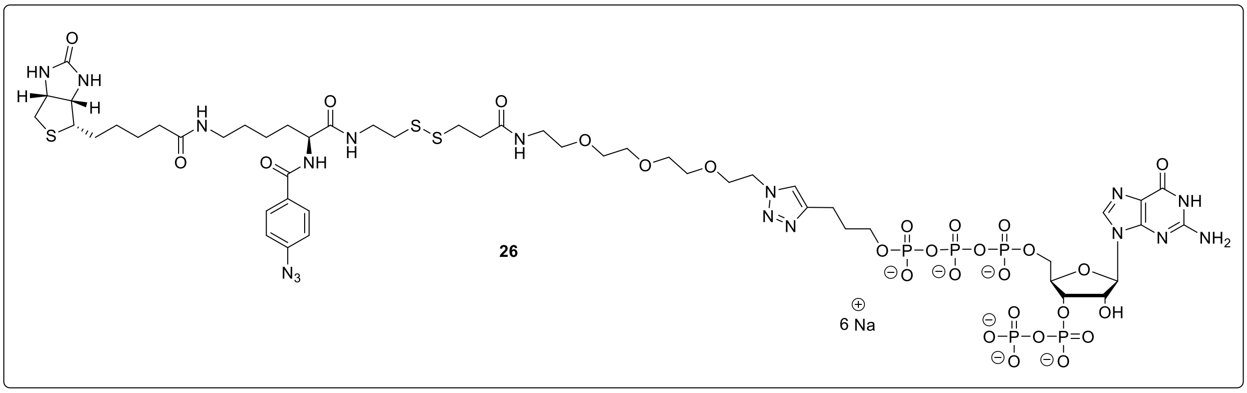


pppGpp – CC7 was synthesized from amino-pppGpp (**19**, 2.10 mg, 1.91 µmol) and Sulfo-SBED according to the general **procedure B**. The product (**26**, 1.56 mg, 886 nmol, 46%) was isolated as white solid.

**$^1$H-NMR** (400 MHz, $D_2O$, δ/ppm): 8.10 (s, 1H), 7.82 (d, *J* = 8.4 Hz, 2H), 7.71 (s, 1H), 7.18 (d, *J* = 8.4 Hz, 2H), 5.94 (d, *J* = 6.4 Hz, 1H), 4.96 (t, *J* = 6.7 Hz, 1H), 4.61 – 4.49 (m, 4H), 4.45 (dd, *J* = 8.8, 5.8 Hz, 1H), 4.38 – 4.21 (m, 3H), 3.99 – 3.86 (m, 4H), 3.69 – 3.49 (m, 13H), 3.36 (dd, *J* = 5.3, 5.3 Hz, 2H), 3.28 – 3.17 (m, 3H), 3.00 – 2.83 (m, 5H), 2.73 (d, *J* = 13.0 Hz, 1H), 2.68 – 2.57 (m, 4H), 2.18 (dd, *J* = 7.2, 7.2 Hz, 2H), 1.99 – 1.85 (m, 2H), 1.85 – 1.74 (m, 2H), 1.69 – 1.37 (m, 8H), 1.36 – 1.21 (m, 2H). **$^{31}$P{$^1$H}-NMR** (162 MHz, $D_2O$, δ/ppm): -5.72 (d, *J* = 22.4 Hz), -10.90 (d, *J* = 22.3 Hz), -10.97 (d, *J* = 19.2 Hz), -11.37 (d, *J* = 17.4 Hz), -22.89 (dd, *J* = 18.5 Hz). **HRMS** (ESI) m/z for $C_{51}H_{77}N_{17}O_{28}P_5S_3$ $[M-H]^{3-}$: calcd 542.0997, found 542.1000. **HPLC-UV** analysis was performed (see SI, chapter 9).

**Synthesis of ppApp – CC (27)**

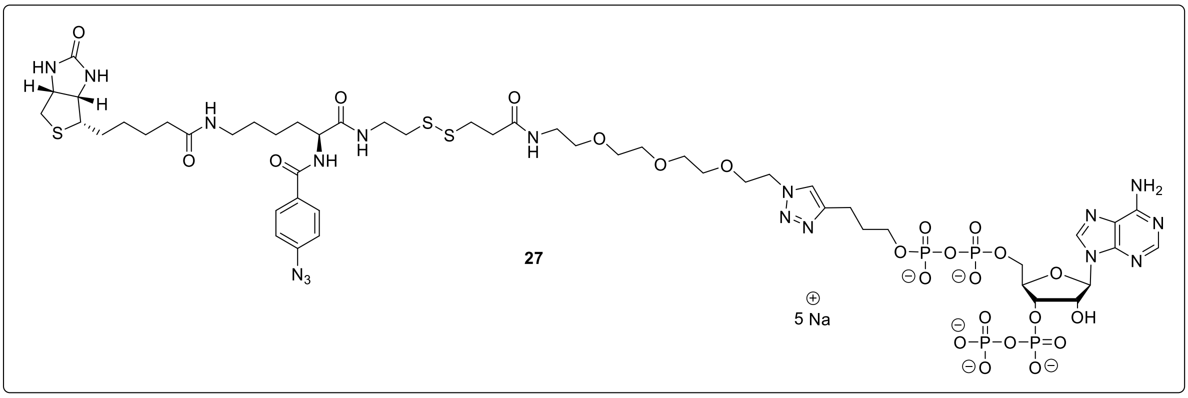


ppApp – CC8 was synthesized from amino-ppApp (**20**, 2.00 mg, 2.04 µmol) and Sulfo-SBED according to the general **procedure B**. The product (**27**, 2.02 mg, 1.21 µmol, 60%) was isolated as white solid.

**$^{1}$H-NMR** (400 MHz, $D_2O$, δ/ppm): 8.52 (s, 1H), 8.18 (d, *J* = 2.8 Hz, 1H), 7.80 (d, *J* = 8.4 Hz, 2H), 7.59 (s, 1H), 7.15 (dd, *J* = 8.7, 2.8 Hz, 2H), 6.14 (d, *J* = 6.1 Hz, 1H), 4.96 (br.s , 1H), 4.78 – 4.75 (m, 1H), 4.62 – 4.51 (m, 2H), 4.51 – 4.38 (m, 2H), 4.38 – 4.17 (m, 3H), 3.95 – 3.79 (m, 4H), 3.66 – 3.47 (m, 13H), 3.35 (dd, *J* = 5.3, 5.3 Hz, 2H), 3.25 – 3.15 (m, 3H), 2.98 – 2.82 (m, 5H), 2.73 (d, *J* = 13.0 Hz, 1H), 2.61 (dd, *J* = 6.7, 6.7 Hz, 2H), 2.57 – 2.44 (m, 2H), 2.17 (dd, *J* = 7.2, 7.2 Hz, 2H), 1.98 – 1.80 (m, 2H), 1.78 – 1.67 (m, 2H), 1.67 – 1.35 (m, 8H), 1.34 – 1.20 (m, 2H). **$^{31}$P{$^{1}$H}-NMR** (162 MHz, $D_2O$, δ/ppm): -5.79 (d, *J* = 22.2 Hz), -10.90 (d, *J* = 19.7 Hz), -11.56 (d, *J* = 20.4 Hz). **HRMS** (ESI) m/z for $C_{51}H_{80}N_{17}O_{24}NaP_4S_3$ $[M+H]^{2+}$: calcd 778.6781, found 778.6774. **HPLC-UV** analysis was performed (see SI, chapter 9).

**Synthesis of ppGp – CC (SI-13)**

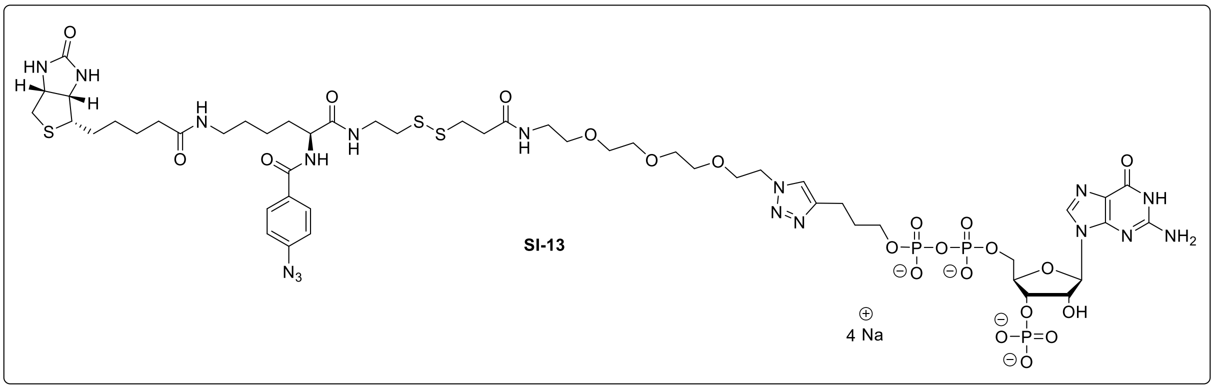


ppGp – CC6 was synthesized from amino-ppGp (**SI-12**, 2.00 mg, 2.23 µmol) and Sulfo-SBED according to the general **procedure B**. The product (**SI-13**, 1.45 mg, 931 nmol, 42%) was isolated as white solid.

**$^{1}$H-NMR** (400 MHz, $D_2O$, δ/ppm): 8.13 (s, 1H), 7.80 (d, *J* = 8.4 Hz, 2H), 7.67 (s, 1H), 7.15 (d, *J* = 8.5 Hz, 2H), 5.91 (d, *J* = 7.0 Hz, 1H), 4.76 (s, 1H), 4.60 – 4.50 (m, 3H), 4.45 (dd, *J* = 8.9, 5.8 Hz, 1H), 4.37 – 4.28 (m, 1H), 4.23 (br. s, *J* = 4.2 Hz, 2H), 3.99 – 3.82 (m, 4H), 3.68 – 3.48 (m, 13H), 3.36 (dd, *J* = 5.3, 5.3 Hz, 2H), 3.28 – 3.14 (m, 3H), 2.99 – 2.83 (m, 5H), 2.73 (d, *J* = 13.1 Hz, 1H), 2.69 – 2.51 (m, 4H), 2.18 (dd, *J* = 7.2, 7.2 Hz, 2H), 1.99 – 1.84 (m, 2H), 1.84 – 1.74 (m, 2H), 1.68 – 1.35 (m, 8H), 1.34 – 1.22 (m, 2H). **$^{31}$P{$^{1}$H}-NMR** (162 MHz, $D_2O$, δ/ppm): 3.89 (s), -10.97 (d, J = 21.0 Hz), -11.52 (d, J = 20.7 Hz). **HRMS** (ESI) m/z for $C_{51}H_{75}N_{17}O_{22}P_3S_3$ $[M-H]^{3-}$: calcd 488.7888, found 488.7889. **HPLC-UV** analysis was performed (see SI, chapter 9).

#### 3. Procedural remarks, pull-down experiments

The pull-down experiments were performed exactly according to the method published by Laventie, Glatter and Jenal in 2017 (“Pull-Down with a c-di-GMP-Specific Capture Compound Coupled to Mass Spectromety as a Powerful Tool to Identify Novel Effector Proteins”).[4,5] In contrast to the example described by Laventie et al. *E. coli* and *S. typhimurium* were used as organisms.

The following methodical steps were performed exactly according to Laventie et al. including the application of identical materials (e.g.: buffers, solvents, reagents, magnetic beads, UV-cross linker…):

1.) Lysate Preparation
2.) Removal of Free Nucleotides (soluble fraction only)
3.) Membrane Resuspension and Solubilization (membrane fraction only)
4.) Protein concentration measurement
5.) Capture
6.) Washing steps
7.) MS Sample Preparation
8.) LC-MS/MS analysis
9.) Database Search
10.) Label-Free Quantification

**Important:** no reducing agents (e.g.: DTT) were used in buffers before the trypsine digest, as some of the capture compounds beard a disulfide bridge.

Notably, typical reaction set-ups differed slightly from Laventie et al. and are summarized in the following tables:

#### MSN – pull-down experiments in *E. coli*:

***Supporting table 1:*** MSN – pull down experiment set-up using *E. coli* extracts. Usually, the protein concentration was above 10 mg/ml so the extract was diluted accordingly before the capture experiment. Both the capture experiment as well as the competition control were performed in triplicates. MSN – competitors were unmodified ppGpp, pppGpp, pGpp, ppApp or ppGp.

| | **Capture Experiment** | **Competition Control** |
|---|---|---|
| ***E. coli* extract** ($c_{protein}$ = 10 mg/mL) | 30 µl | 30 µl |
| **MSN – competitor** (c = 20 mM) | - | 10 µl |
| **Capture buffer** (5x) | 20 µl | 20 µl |
| **dd$H_2O$** | 40 µl | 30 µl |
| *30 min incubation at 4 °C* | | |
| **MSN – capture compound** (c = 200 µM) | 10 µl | 10 µl |
| *2 h incubation at 4 °C, then UV-crosslinking* | | |
| **final volume** | 100 µl | 100 µl |

#### MSN – pull-down experiments in *S. typhimurium*:

***Supporting table 2:*** MSN – pull down experiment set-up using *S. typhimurium* extracts. In this case the protein concentration of the soluble fraction was below 10 mg/ml. Consequently, the applied volume was increased to keep the protein roughly constant compared to the *E. coli* experiments. Both the capture experiment as well as the competition control were performed in triplicates. In the case of *S. typhimurium*, ppGpp-CC1&2 were applied as mixture.

| | **Capture Experiment** | **Competition Control** |
|---|---|---|
| ***S. typhimurium* extract** ($c_{protein}$ = 4.7 mg/mL) | 60 µl | 60 µl |
| **ppGpp** (c = 20 mM) | - | 10 µl |
| **Capture buffer** (5x) | 20 µl | 20 µl |
| **dd$H_2O$** | 10 µl | - |
| *30 min incubation at 4 °C* | | |
| **ppGpp – CC1+2 - mix** (c = 100 µM each) | 10 µl | 10 µl |
| *2 h incubation at 4 °C, then UV-crosslinking* | | |
| **final volume** | 100 µl | 100 µl |

### 4. Pull-down results: enrichment tables and hit-maps

***Legend:***

- Table entries marked in **green** are known ppGpp – receptors.[6]
- Table entries marked in **yellow** were already captured by Laub et al.[7]
- *Threshold 1:* $\log_2$ (enrichment) ≥ 2.0
- *Threshold 2:* q-value ≤ 0.05

#### 4.1. *E. coli* (soluble fraction)

##### 4.1.1. ppGpp-CC1 (soluble fraction)

***Supporting table 3:*** *putative ppGpp receptors captured by ppGpp-CC1 from the soluble fraction of E. coli cell lysate.*

| Nr. | ac | gene | protein | $\log_2$ (enrichment) | q-value |
|---|---|---|---|---|---|
| 1 | P0A6D7 | *aroK* | Shikimate kinase 1 | 9,3 | 0,01 |
| 2 | P33195 | *gcvP* | Glycine dehydrogenase | 9,1 | 0,04 |
| 3 | P0AG76 | *sbcD* | Nuclease SbcCD subunit D | 8,9 | 0,03 |
| 4 | P0AGE0 | *ssb* | Single-stranded DNA-binding protein | 8,0 | 0,02 |
| 5 | P31808 | *yciK* | Uncharacterized oxidoreductase | 6,9 | 0,00 |
| 6 | P24203 | *yjiA* | P-loop guanosine triphosphatase | 6,3 | 0,03 |
| 7 | P0AAB6 | *galF* | UTP--glucose-1-phosphate uridylyltransferase | 6,1 | 0,02 |
| 8 | P34209 | *ydcF* | uncharacterized protein | 5,6 | 0,04 |
| 9 | P63389 | *yheS* | Uncharacterized ABC transporter ATP-binding protein | 5,5 | 0,00 |
| 10 | P0A9D2 | *gstA* | Glutathione S-transferase | 4,6 | 0,00 |
| 11 | P10121 | *ftsY* | Signal recognition particle receptor | 3,7 | 0,01 |
| 12 | P06987 | *hisB* | Histidine biosynthesis bifunctional protein | 3,7 | 0,00 |
| 13 | P0A993 | *fbp* | Fructose-1,6-bisphosphatase class 1 | 3,6 | 0,01 |
| 14 | P0ACA3 | *sspA* | Stringent starvation protein A | 3,5 | 0,02 |
| 15 | P36566 | *cmoM* | tRNA 5-carboxymethoxyuridine methyltransferase | 3,4 | 0,00 |
| 16 | P0A6K6 | *deoB* | Phosphopentomutase | 3,1 | 0,00 |
| 17 | P00887 | *aroH* | Phospho-2-dehydro-3-deoxyheptonate aldolase | 3,0 | 0,01 |
| 18 | P12281 | *moeA* | Molybdopterin molybdenumtransferase | 2,9 | 0,01 |
| 19 | P0ADR8 | *ppnN* | Pyrimidine/purine nucleotide 5'-monophosphate nucleosidase | 2,9 | 0,00 |
| 20 | P62768 | *yaeH* | uncharacterized protein | 2,8 | 0,00 |
| 21 | P00962 | *glnS* | Glutamine--tRNA ligase | 2,8 | 0,01 |

| | | | | | |
|---|---|---|---|---|---|
| **22** | P06999 | *pfkB* | ATP-dependent 6-phosphofructokinase isozyme 2 | 2,8 | 0,00 |
| **23** | P0AF20 | *nagC* | N-acetylglucosamine repressor | 2,7 | 0,01 |
| **24** | P00350 | *gnd* | 6-phosphogluconate dehydrogenase, decarboxylating | 2,6 | 0,02 |
| **25** | P76594 | *pka* | Protein lysine acetyltransferase | 2,6 | 0,02 |
| **26** | P0A7B8 | *hslV* | ATP-dependent protease subunit | 2,4 | 0,01 |
| **27** | P32132 | *typA* | GTP-binding protein TypA/BipA | 2,4 | 0,02 |
| **28** | P60906 | *hisS* | Histidine--tRNA ligase | 2,4 | 0,01 |
| **29** | P0A9M5 | *gpt* | Xanthine phosphoribosyltransferase | 2,3 | 0,02 |
| **30** | P0AEE5 | *mglB* | D-galactose-binding periplasmic protein | 2,3 | 0,01 |
| **31** | P14081 | *selB* | Selenocysteine-specific elongation factor | 2,2 | 0,01 |
| **32** | P08203 | *araD* | L-ribulose-5-phosphate 4-epimerase | 2,2 | 0,02 |
| **33** | P0A850 | *tig* | Trigger factor | 2,2 | 0,02 |
| **34** | P0AEP3 | *galU* | UTP--glucose-1-phosphate uridylyltransferase | 2,2 | 0,00 |
| **35** | P0A6T5 | *folE* | GTP cyclohydrolase 1 | 2,2 | 0,01 |
| **36** | P0A6M8 | *fusA* | Elongation factor G | 2,1 | 0,01 |
| **37** | P21165 | *pepQ* | Xaa-Pro dipeptidase | 2,1 | 0,01 |
| **38** | P0A796 | *pfkA* | ATP-dependent 6-phosphofructokinase isozyme 1 | 2,0 | 0,01 |
| **39** | P0A7B5 | *proB* | Glutamate 5-kinase | 2,0 | 0,01 |
| **40** | P17117 | *nfsA* | Oxygen-insensitive NADPH nitroreductase | 2,0 | 0,02 |
| **41** | P0A7A5 | *pcm* | Protein-L-isoaspartate O-methyltransferase | 2,0 | 0,02 |

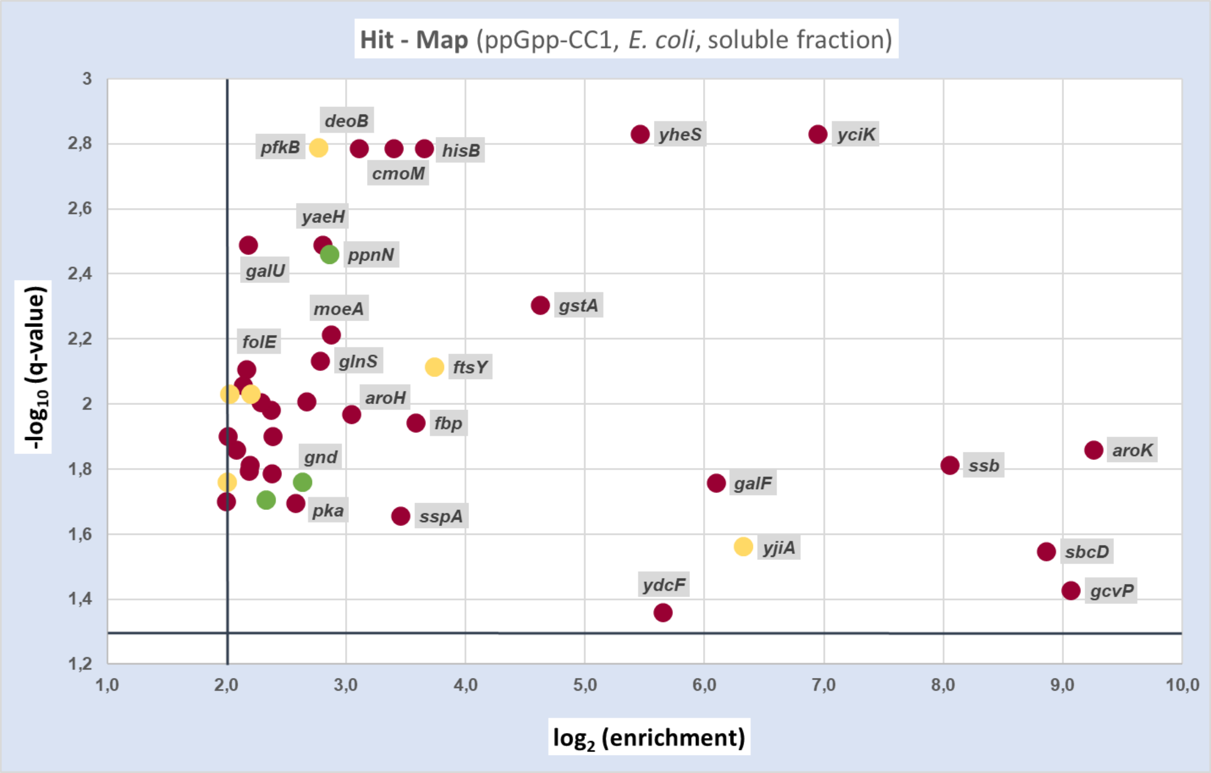


***Supporting figure 4:*** *graphic representation of proteins captured by ppGpp-CC1 from the E. coli soluble fraction. Only hits complying the threshold criteria are shown in the plot.*

##### 4.1.2. ppGpp-CC2 (soluble fraction)

***Supporting table 4:*** *putative ppGpp receptors captured by ppGpp-CC2 from the soluble fraction of E. coli cell lysate. An extract of this table was already shown in the main part of the manuscript.*

| Nr. | ac | gene | protein | $log_2$ (enrichment) | q-value |
|---|---|---|---|---|---|
| 1 | P12281 | *moeA* | Molybdopterin molybdenumtransferase | 9,4 | 0,00 |
| 2 | P0AFT5 | *btsR* | Transcriptional regulatory protein | 7,7 | 0,04 |
| 3 | P31808 | *yciK* | Uncharacterized oxidoreductase YciK | 7,1 | 0,02 |
| 4 | P0A8F0 | *upp* | Uracil phosphoribosyltransferase | 6,6 | 0,01 |
| 5 | P21693 | *dbpA* | ATP-dependent RNA helicase | 5,3 | 0,00 |
| 6 | P68688 | *grxA* | Glutaredoxin 1 | 4,8 | 0,00 |
| 7 | P0A6R0 | *fabH* | 3-oxoacyl-[acyl-carrier-protein] synthase 3 | 4,7 | 0,00 |
| 8 | P0ADR8 | *ppnN* | Pyrimidine/purine nucleotide 5'-monophosphate nucleosidase | 4,1 | 0,00 |
| 9 | P0A6K6 | *deoB* | Phosphopentomutase | 4,1 | 0,01 |
| 10 | P0A9M5 | *gpt* | Xanthine phosphoribosyltransferase | 4,0 | 0,00 |
| 11 | P0AF20 | *nagC* | N-acetylglucosamine repressor | 4,0 | 0,01 |
| 12 | P0A7B5 | *proB* | Glutamate 5-kinase | 3,7 | 0,01 |
| 13 | P0A6T5 | *folE* | GTP cyclohydrolase 1 | 3,5 | 0,01 |

| | | | | | |
|---|---|---|---|---|---|
| **14** | P76594 | *pka* | Protein lysine acetyltransferase | 3,5 | 0,00 |
| **15** | P0A7D7 | *purC* | Phosphoribosylaminoimidazole-succinocarboxamide synthase | 3,5 | 0,00 |
| **16** | P63389 | *yheS* | Uncharacterized ABC transporter ATP-binding protein | 3,3 | 0,00 |
| **17** | P0AEE5 | *mglB* | D-galactose-binding periplasmic protein | 3,3 | 0,00 |
| **18** | P30749 | *moaE* | Molybdopterin synthase catalytic subunit | 3,2 | 0,01 |
| **19** | P04825 | *pepN* | Aminopeptidase N | 3,2 | 0,00 |
| **20** | P00350 | *gnd* | 6-phosphogluconate dehydrogenase, decarboxylating | 2,9 | 0,00 |
| **21** | P38489 | *nfsB* | Oxygen-insensitive NAD(P)H nitroreductase | 2,9 | 0,01 |
| **22** | P34209 | *ydcF* | Uncharacterized protein | 2,8 | 0,02 |
| **23** | P60906 | *hisS* | Histidine--tRNA ligase | 2,8 | 0,03 |
| **24** | P0AC59 | *grxB* | Glutaredoxin 2 | 2,8 | 0,00 |
| **25** | P0A761 | *nanE* | Putative N-acetylmannosamine-6-phosphate 2-epimerase | 2,7 | 0,00 |
| **26** | P31120 | *glmM* | Phosphoglucosamine mutase | 2,7 | 0,00 |
| **27** | P61949 | *fldA* | Flavodoxin 1 | 2,7 | 0,00 |
| **28** | P39337 | *yjgM* | Uncharacterized N-acetyltransferase | 2,7 | 0,00 |
| **29** | P31060 | *modF* | ABC transporter ATP-binding protein | 2,7 | 0,00 |
| **30** | P05637 | *apaH* | Bis(5'-nucleosyl)-tetraphosphatase | 2,6 | 0,01 |
| **31** | P0AG76 | *sbcD* | Nuclease | 2,6 | 0,04 |
| **32** | P00887 | *aroH* | Phospho-2-dehydro-3-deoxyheptonate aldolase, Trp-sensitive | 2,6 | 0,01 |
| **33** | P0AEP3 | *galU* | UTP--glucose-1-phosphate uridylyltransferase | 2,6 | 0,00 |
| **34** | P36566 | *cmoM* | tRNA 5-carboxymethoxyuridine methyltransferase | 2,5 | 0,02 |
| **35** | P68206 | *yjbJ* | Uncharacterized protein | 2,5 | 0,00 |
| **36** | P52061 | *rdgB* | dITP/XTP pyrophosphatase | 2,5 | 0,00 |
| **37** | P0A870 | *talB* | Transaldolase B | 2,4 | 0,01 |
| **38** | P14081 | *selB* | Selenocysteine-specific elongation factor | 2,4 | 0,00 |
| **39** | P0A796 | *pfkA* | ATP-dependent 6-phosphofructokinase | 2,4 | 0,00 |
| **40** | P0A7B8 | *hslV* | ATP-dependent protease subunit | 2,3 | 0,01 |
| **41** | P25665 | *metE* | 5-methyltetrahydropteroyltriglutamate--homocysteine methyltransferase | 2,3 | 0,00 |
| **42** | P0AEZ9 | *moaB* | Molybdenum cofactor biosynthesis protein B | 2,3 | 0,00 |
| **43** | P00934 | *thrC* | Threonine synthase | 2,2 | 0,00 |
| **44** | P0AGE9 | *sucD* | Succinate--CoA ligase subunit alpha | 2,2 | 0,00 |
| **45** | P07118 | *valS* | Valine--tRNA ligase | 2,2 | 0,02 |
| **46** | P37330 | *glcB* | Malate synthase G | 2,2 | 0,00 |
| **47** | P37744 | *rfbA* | Glucose-1-phosphate thymidylyltransferase | 2,2 | 0,01 |
| **48** | P0A6T3 | *galK* | Galactokinase | 2,2 | 0,03 |

| | | | | | |
|---|---|---|---|---|---|
| **49** | P0A6N8 | *yeiP* | Elongation factor P-like protein | 2,2 | 0,04 |
| **50** | P0A799 | *pgk* | Phosphoglycerate kinase | 2,1 | 0,00 |
| **51** | P0A6M8 | *fusA* | Elongation factor G | 2,1 | 0,02 |
| **52** | Q46829 | *bglA* | 6-phospho-beta-glucosidase | 2,1 | 0,00 |
| **53** | P0ACW6 | *ydcH* | Uncharacterized protein | 2,1 | 0,02 |
| **54** | P0A9J0 | *rng* | Ribonuclease G | 2,1 | 0,04 |
| **55** | P0A6P9 | *eno* | Enolase | 2,1 | 0,01 |
| **56** | P08997 | *aceB* | Malate synthase A | 2,1 | 0,00 |
| **57** | P0A7E5 | *pyrG* | CTP synthase | 2,1 | 0,00 |
| **58** | P32132 | *typA* | GTP-binding protein TypA/BipA | 2,1 | 0,00 |
| **59** | P06987 | *hisB* | Histidine biosynthesis bifunctional protein | 2,0 | 0,00 |
| **60** | P69503 | *apt* | Adenine phosphoribosyltransferase | 2,0 | 0,04 |
| **61** | P62768 | *yaeH* | UPF0325 protein YaeH | 2,0 | 0,00 |
| **62** | P0A7D4 | *purA* | Adenylosuccinate synthetase | 2,0 | 0,00 |
| **63** | P0AB89 | *purB* | Adenylosuccinate lyase | 2,0 | 0,00 |
| **64** | P0AE12 | *amn* | AMP nucleosidase | 2,0 | 0,01 |

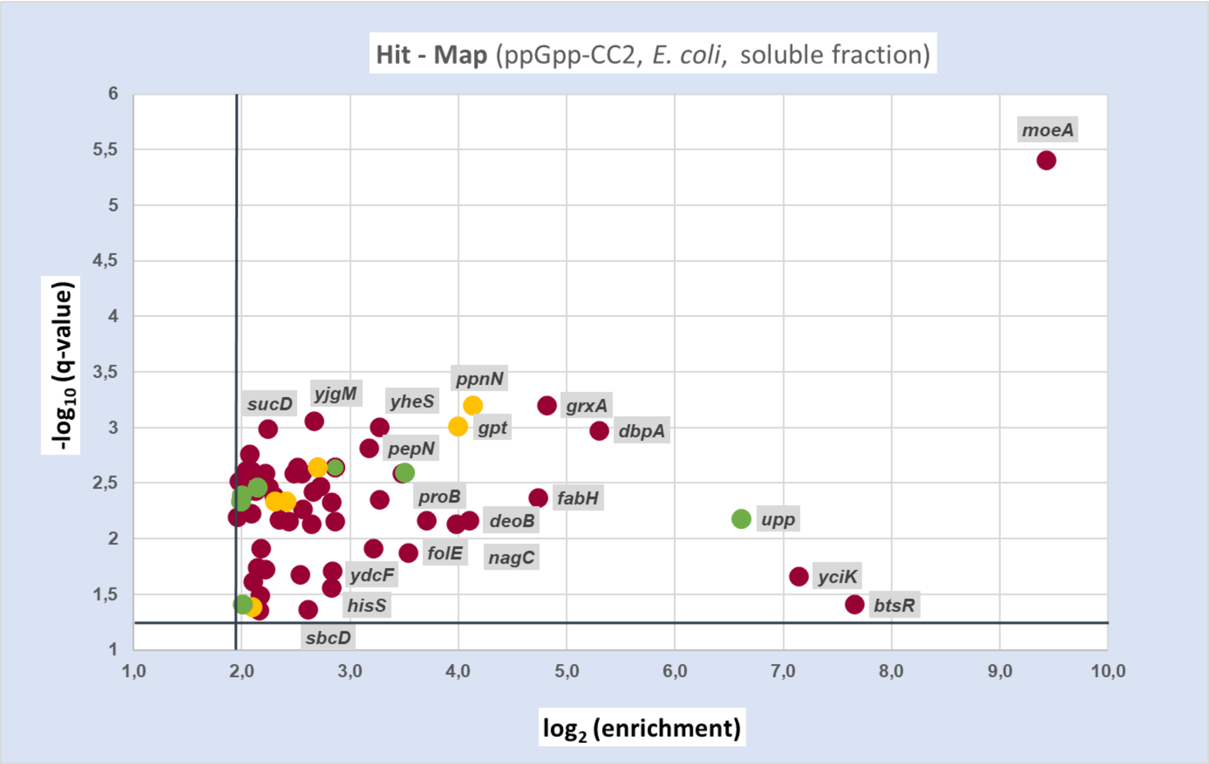


***Supporting figure 5:*** *graphic representation of proteins captured by ppGpp-CC2 from the E. coli soluble fraction. Only hits complying the threshold criteria are shown in the plot.*

##### 4.1.3. ppGpp-CC3 (soluble fraction)

***Supporting table 5:*** *putative ppGpp receptors captured by ppGpp-CC3 from the soluble fraction of E. coli cell lysate.*

| Nr. | ac | gene | protein | $log_2$ (enrichment) | q-value |
|---|---|---|---|---|---|
| 1 | P0ACY1 | *ydjA* | Putative NAD(P)H nitroreductase | 4,0 | 0,00 |

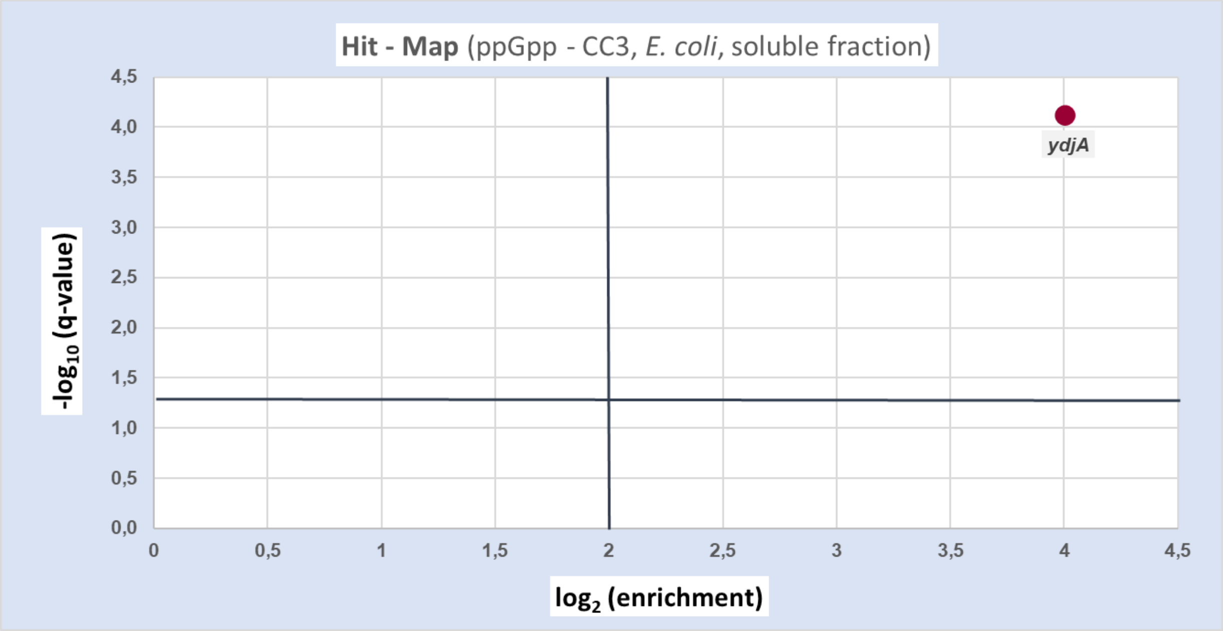


***Supporting figure 6:*** *graphic representation of proteins captured by ppGpp-CC3 from the E. coli soluble fraction. Only hits complying the threshold criteria are shown in the plot.*

##### 4.1.4. ppGpp-CC4 (soluble fraction)

***Supporting table 6:*** *putative ppGpp receptors captured by ppGpp-CC4 from the soluble fraction of E. coli cell lysate.*

| Nr. | ac | gene | protein | $log_2$ (enrichment) | q-value |
|---|---|---|---|---|---|
| 1 | P62707 | *gpmA* | 2,3-bisphosphoglycerate-dependent phosphoglycerate mutase | 2,6 | 0,00 |
| 2 | P0A6R0 | *fabH* | 3-oxoacyl-[acyl-carrier-protein] synthase 3 | 2,5 | 0,00 |

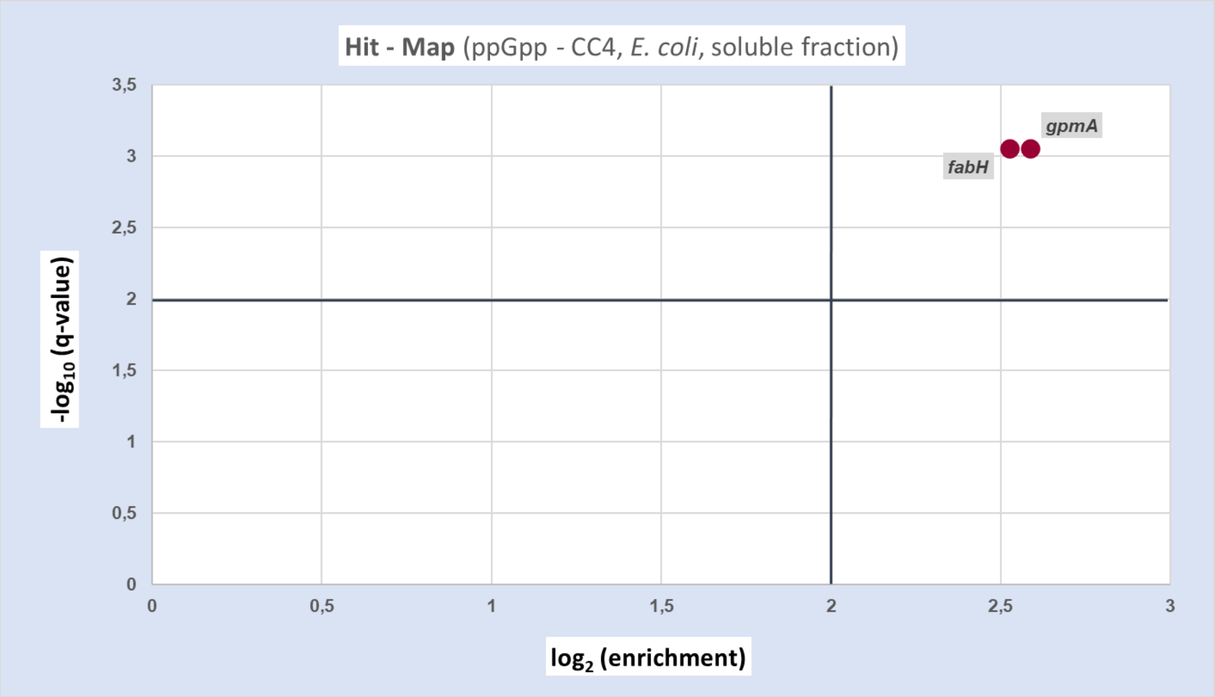


***Supporting figure 7:*** *graphic representation of proteins captured by ppGpp-CC4 from the E. coli soluble fraction. Only hits complying the threshold criteria are shown in the plot.*

##### 4.1.5. pGpp-CC (soluble fraction)

***Supporting table 7:*** *putative pGpp receptors captured by pGpp-CC from the soluble fraction of E. coli cell lysate.*

| Nr. | ac | gene | protein | $log_2$ (enrichment) | q-value |
|---|---|---|---|---|---|
| 1 | P0AEE5 | *mglB* | D-galactose-binding periplasmic protein | 3,8 | 0,00 |
| 2 | P0ACU7 | *yjdC* | HTH-type transcriptional regulator | 3,6 | 0,02 |
| 3 | P0A6R0 | *fabH* | 3-oxoacyl-[acyl-carrier-protein] synthase 3 | 3,5 | 0,00 |
| 4 | P39160 | *uxuB* | D-mannonate oxidoreductase | 3,4 | 0,04 |
| 5 | P34209 | *ydcF* | Uncharacterized protein | 3,2 | 0,02 |
| 6 | P0A6K6 | *deoB* | Phosphopentomutase | 3,1 | 0,00 |
| 7 | P00962 | *glnS* | Glutamine--tRNA ligase | 3,1 | 0,01 |
| 8 | P52061 | *rdgB* | dITP/XTP pyrophosphatase | 3,0 | 0,00 |
| 9 | P00934 | *thrC* | Threonine synthase | 3,0 | 0,02 |
| 10 | P04825 | *pepN* | Aminopeptidase N | 3,0 | 0,01 |
| 11 | P0A7B8 | *hslV* | ATP-dependent protease subunit | 2,9 | 0,03 |
| 12 | P0A879 | *trpB* | Tryptophan synthase beta chain | 2,9 | 0,00 |
| 13 | P0A8F0 | *upp* | Uracil phosphoribosyltransferase | 2,8 | 0,03 |
| 14 | P76594 | *pka* | Protein lysine acetyltransferase | 2,8 | 0,03 |
| 15 | P00864 | *ppc* | Phosphoenolpyruvate carboxylase | 2,7 | 0,01 |
| 16 | P21165 | *pepQ* | Xaa-Pro dipeptidase | 2,6 | 0,00 |
| 17 | P0AGE0 | *ssb* | Single-stranded DNA-binding protein | 2,5 | 0,00 |
| 18 | P0ADR8 | *ppnN* | Pyrimidine/purine nucleotide 5'-monophosphate nucleosidase | 2,3 | 0,00 |
| 19 | P0AC59 | *grxB* | Glutaredoxin | 2,3 | 0,01 |
| 20 | P0A9D2 | *gstA* | Glutathione S-transferase GstA | 2,2 | 0,02 |
| 21 | P0A972 | *cspE* | Cold shock-like protein CspE | 2,2 | 0,01 |
| 22 | P06999 | *pfkB* | ATP-dependent 6-phosphofructokinase isozyme 2 | 2,2 | 0,01 |
| 23 | P30749 | *moaE* | Molybdopterin synthase catalytic subunit | 2,1 | 0,01 |
| 24 | P37744 | *rfbA* | Glucose-1-phosphate thymidylyltransferase 1 | 2,1 | 0,00 |
| 25 | P0A799 | *pgk* | Phosphoglycerate kinase | 2,1 | 0,01 |
| 26 | P05637 | *apaH* | Bis(5'-nucleosyl)-tetraphosphatase | 2,1 | 0,03 |
| 27 | P0AC53 | *zwf* | Glucose-6-phosphate 1-dehydrogenase | 2,1 | 0,00 |
| 28 | P14081 | *selB* | Selenocysteine-specific elongation factor | 2,0 | 0,01 |
| 29 | P06987 | *hisB* | Histidine biosynthesis bifunctional protein | 2,0 | 0,02 |

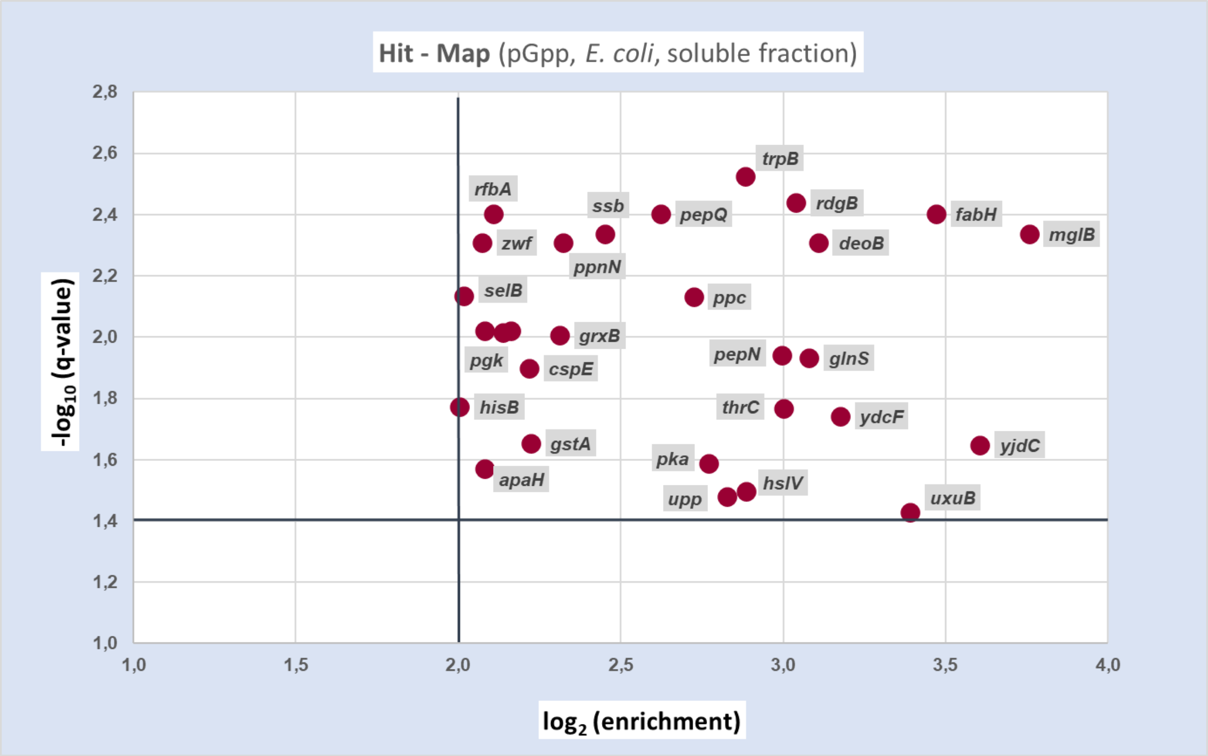


***Supporting figure 8:*** *graphic representation of proteins captured by pGpp-CC from the E. coli soluble fraction. Only hits complying the threshold criteria are shown in the plot.*

##### 4.1.6. pppGpp-CC (soluble fraction)

***Supporting table 8:*** *putative pppGpp receptors captured by pppGpp-CC from the soluble fraction of E. coli cell lysate.*

| Nr. | ac | gene | protein | $log_2$ (enrichment) | q-value |
|---|---|---|---|---|---|
| 1 | P0A6N4 | *efp* | Elongation factor P | 8,4 | 0,02 |
| 2 | P0A6T5 | *folE* | GTP cyclohydrolase 1 | 7,4 | 0,00 |
| 3 | P0A6N8 | *yeiP* | Elongation factor P-like protein | 6,4 | 0,01 |
| 4 | P68688 | *grxA* | Glutaredoxin 1 | 6,0 | 0,01 |
| 5 | P34209 | *ydcF* | Protein YdcF | 5,6 | 0,00 |
| 6 | P0AEE5 | *mglB* | D-galactose-binding periplasmic protein | 4,9 | 0,00 |
| 7 | P0ADR8 | *ppnN* | Pyrimidine/purine nucleotide 5'-monophosphate nucleosidase | 4,4 | 0,01 |
| 8 | P00864 | *ppc* | Phosphoenolpyruvate carboxylase | 3,8 | 0,01 |
| 9 | P61949 | *fldA* | Flavodoxin 1 OS=Escherichia coli | 3,7 | 0,01 |
| 10 | P0A7D7 | *purC* | Phosphoribosylaminoimidazole-succinocarboxamide synthase | 3,7 | 0,00 |
| 11 | P0AC59 | *grxB* | Glutaredoxin 2 | 3,5 | 0,00 |
| 12 | P0A7B5 | *proB* | Glutamate 5-kinase | 3,3 | 0,01 |

| | | | | | |
|---|---|---|---|---|---|
| **13** | P0A870 | *talB* | Transaldolase B | 3,3 | 0,01 |
| **14** | P14081 | *selB* | Selenocysteine-specific elongation factor | 3,3 | 0,00 |
| **15** | P37330 | *glcB* | Malate synthase G | 3,3 | 0,01 |
| **16** | P60906 | *hisS* | Histidine--tRNA ligase | 3,3 | 0,03 |
| **17** | P0A6R0 | *fabH* | 3-oxoacyl-[acyl-carrier-protein] synthase 3 | 3,3 | 0,01 |
| **18** | P55135 | *rlmD* | 23S rRNA (uracil(1939)-C(5))-methyltransferase RlmD | 3,2 | 0,01 |
| **19** | P38489 | *nfsB* | Oxygen-insensitive NAD(P)H nitroreductase | 2,9 | 0,01 |
| **20** | P0ABT2 | *dps* | DNA protection during starvation protein | 2,9 | 0,00 |
| **21** | P21165 | *pepQ* | Xaa-Pro dipeptidase | 2,8 | 0,00 |
| **22** | P14294 | *topB* | DNA topoisomerase 3 | 2,6 | 0,00 |
| **23** | P00962 | *glnS* | Glutamine--tRNA ligase | 2,6 | 0,01 |
| **24** | P42641 | *obgE* | GTPase ObgE/CgtA | 2,6 | 0,00 |
| **25** | P0A761 | *nanE* | Putative N-acetylmannosamine-6-phosphate 2-epimerase | 2,6 | 0,00 |
| **26** | P0A6P7 | *engB* | Probable GTP-binding protein | 2,6 | 0,03 |
| **27** | P07118 | *valS* | Valine--tRNA ligase | 2,6 | 0,01 |
| **28** | P0A6P9 | *eno* | Enolase | 2,6 | 0,00 |
| **29** | P68206 | *yjbJ* | UPF0337 protein YjbJ | 2,5 | 0,01 |
| **30** | P0AG40 | *ribF* | Bifunctional riboflavin kinase/FMN adenylyltransferase | 2,5 | 0,00 |
| **31** | P00934 | *thrC* | Threonine synthase | 2,4 | 0,01 |
| **32** | P24203 | *yjiA* | P-loop guanosine triphosphatase | 2,4 | 0,03 |
| **33** | P0A7B8 | *hslV* | ATP-dependent protease subunit | 2,4 | 0,01 |
| **34** | P0AED5 | *uvrY* | Response regulator UvrY | 2,3 | 0,02 |
| **35** | P77735 | *yajO* | 1-deoxyxylulose-5-phosphate synthase | 2,3 | 0,00 |
| **36** | P0AEH3 | *elaA* | Protein ElaA | 2,3 | 0,01 |
| **37** | P0AFG8 | *aceE* | Pyruvate dehydrogenase E1 component | 2,3 | 0,00 |
| **38** | P25665 | *metE* | 5-methyltetrahydropteroyltriglutamate – homocysteine methyltransferase | 2,3 | 0,00 |
| **39** | P31120 | *glmM* | Phosphoglucosamine mutase | 2,3 | 0,01 |
| **40** | P05637 | *apaH* | Bis(5'-nucleosyl)-tetraphosphatase | 2,3 | 0,01 |
| **41** | P21599 | *pykA* | Pyruvate kinase II | 2,3 | 0,00 |
| **42** | P52061 | *rdgB* | dITP/XTP pyrophosphatase | 2,3 | 0,01 |
| **43** | Q46920 | *queF* | NADPH-dependent 7-cyano-7-deazaguanine reductase | 2,2 | 0,00 |
| **44** | P0A6F9 | *groS* | 10 kDa chaperonin | 2,2 | 0,00 |
| **45** | P76291 | *cmoB* | tRNA U34 carboxymethyltransferase | 2,2 | 0,00 |
| **46** | P0A796 | *pfkA* | ATP-dependent 6-phosphofructokinase isozyme 1 | 2,2 | 0,01 |
| **47** | P0AE12 | *amn* | AMP nucleosidase | 2,2 | 0,00 |
| **48** | P08956 | *hsdR* | Type I restriction enzyme EcoKI R | 2,2 | 0,03 |

| | | | | | |
|---|---|---|---|---|---|
| **49** | P06987 | *hisB* | Histidine biosynthesis bifunctional protein | 2,2 | 0,02 |
| **50** | P00350 | *gnd* | 6-phosphogluconate dehydrogenase, decarboxylating | 2,1 | 0,01 |
| **51** | P0AEZ9 | *moaB* | Molybdenum cofactor biosynthesis protein B | 2,1 | 0,01 |
| **52** | P30850 | *rnb* | Exoribonuclease 2 | 2,1 | 0,01 |
| **53** | P62768 | *yaeH* | UPF0325 protein YaeH | 2,1 | 0,00 |
| **54** | P0AB89 | *purB* | Adenylosuccinate lyase | 2,1 | 0,00 |
| **55** | P0A6E4 | *argG* | Argininosuccinate synthase | 2,1 | 0,01 |
| **56** | P0A805 | *frr* | Ribosome-recycling factor | 2,1 | 0,00 |
| **57** | P77570 | *anmK* | Anhydro-N-acetylmuramic acid kinase | 2,1 | 0,01 |
| **58** | P06612 | *topA* | DNA topoisomerase 1 | 2,0 | 0,01 |
| **59** | P0A7E5 | *pyrG* | CTP synthase | 2,0 | 0,00 |

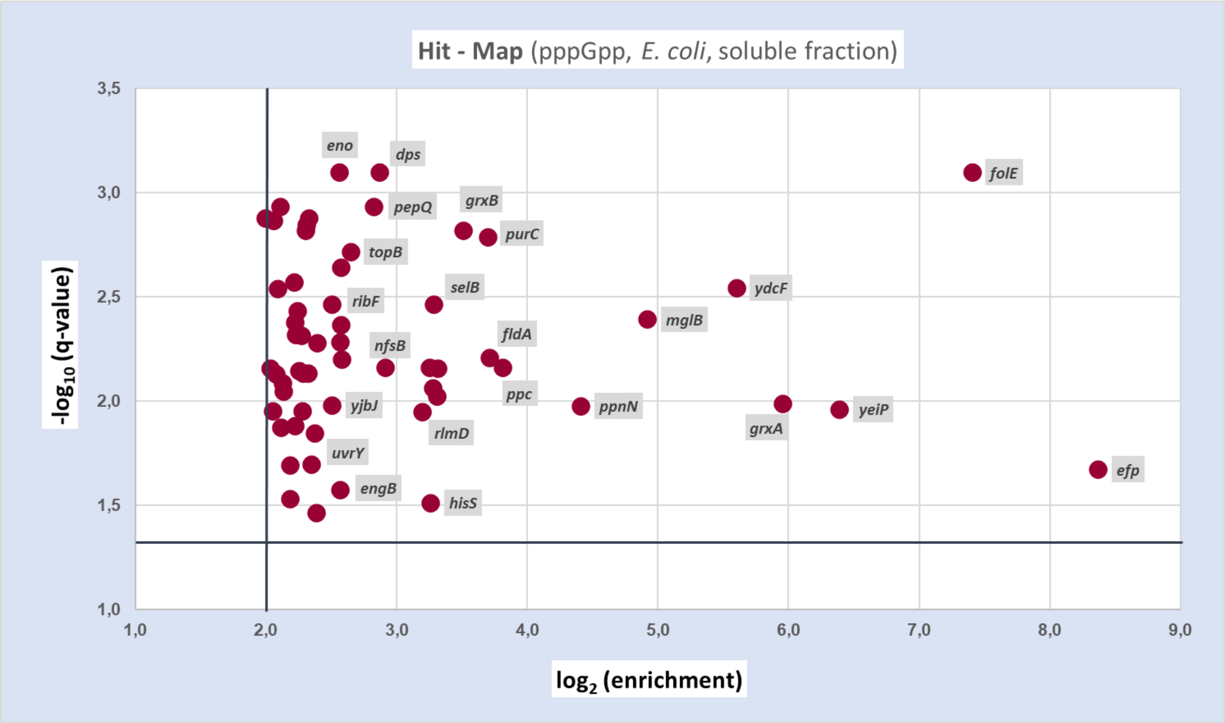


***Supporting figure 9:*** *graphic representation of proteins captured by pppGpp-CC from the E. coli soluble fraction. Only hits complying the threshold criteria are shown in the plot.*

##### 4.1.7. ppApp-CC (soluble fraction)

***Supporting table 9:*** *putative ppApp receptors captured by ppApp-CC from the soluble fraction of E. coli cell lysate.*

| ***Nr.*** | ***ac*** | ***gene*** | ***protein*** | ***log₂ (enrichment)*** | ***q-value*** |
|---|---|---|---|---|---|
| **1** | P0ADX7 | *yhhA* | Uncharacterized protein YhhA | 7,3 | 0,00 |
| **2** | P06999 | *pfkB* | ATP-dependent 6-phosphofructokinase isozyme 2 | 3,1 | 0,01 |
| **3** | P0A6P9 | *eno* | Enolase | 2,4 | 0,01 |
| **4** | P0AG40 | *ribF* | Bifunctional riboflavin kinase/FMN adenylyltransferase | 2,4 | 0,03 |
| **5** | P0ACY3 | *yeaG* | Uncharacterized protein | 2,0 | 0,02 |
| **6** | P0A717 | *prs* | Ribose-phosphate pyrophosphokinase | 2,0 | 0,02 |

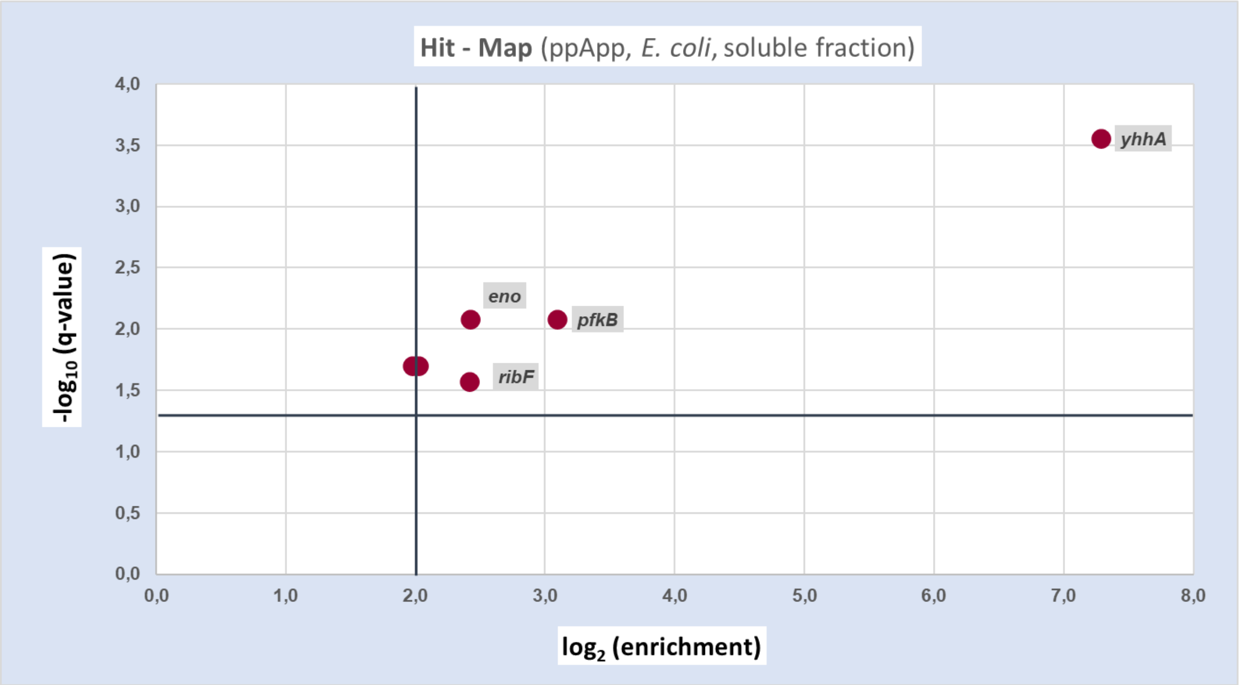


***Supporting figure 10:*** *graphic representation of proteins captured by ppApp-CC from the E. coli soluble fraction. Only hits complying the threshold criteria are shown in the plot.*

##### 4.1.8. ppGp-CC (soluble fraction)

***Supporting table 10:** putative ppGp receptors captured by ppGp-CC from the soluble fraction of E. coli cell lysate.*

| Nr. | ac | gene | protein | $log_2$ (enrichment) | q-value |
|---|---|---|---|---|---|
| 1 | P0ACF4 | *hupB* | DNA-binding protein HU-beta | 8,3 | 0,01 |
| 2 | P0A6T5 | *folE* | GTP cyclohydrolase 1 | 6,7 | 0,01 |
| 3 | P39831 | *ydfG* | NADP-dependent 3-hydroxy acid dehydrogenase | 6,7 | 0,00 |
| 4 | P77374 | *ynfE* | Putative dimethyl sulfoxide reductase chain | 5,1 | 0,00 |
| 5 | P0A6K3 | *def* | Peptide deformylase | 4,9 | 0,00 |
| 6 | P34209 | *ydcF* | Protein YdcF | 4,7 | 0,00 |
| 7 | P33218 | *yebE* | Inner membrane protein YebE | 4,7 | 0,00 |
| 8 | P17117 | *nfsA* | Oxygen-insensitive NADPH nitroreductase | 4,6 | 0,00 |
| 9 | P0AGG8 | *tldD* | Metalloprotease TldD | 4,2 | 0,00 |
| 10 | P10121 | *ftsY* | Signal recognition particle receptor | 4,1 | 0,00 |
| 11 | P0ACF0 | *hupA* | DNA-binding protein | 4,0 | 0,00 |
| 12 | P12281 | *moeA* | Molybdopterin molybdenumtransferase | 3,9 | 0,00 |
| 13 | P0A8F0 | *upp* | Uracil phosphoribosyltransferase | 3,9 | 0,00 |
| 14 | P0AEE5 | *mglB* | D-galactose-binding periplasmic protein | 3,8 | 0,00 |
| 15 | P0A8X0 | *yjgA* | UPF0307 protein YjgA | 3,8 | 0,00 |
| 16 | P0AFI7 | *pdxH* | Pyridoxine/pyridoxamine 5'-phosphate oxidase | 3,8 | 0,00 |
| 17 | P38489 | *nfsB* | Oxygen-insensitive NAD(P)H nitroreductase | 3,7 | 0,00 |
| 18 | P31120 | *glmM* | Phosphoglucosamine mutase | 3,6 | 0,00 |
| 19 | P23871 | *hemH* | Ferrochelatase | 3,6 | 0,00 |
| 20 | P75960 | *cobB* | NAD-dependent protein deacylase | 3,5 | 0,00 |
| 21 | P14294 | *topB* | DNA topoisomerase 3 | 3,5 | 0,00 |
| 22 | P39160 | *uxuB* | D-mannonate oxidoreductase | 3,3 | 0,00 |
| 23 | P37747 | *glf* | UDP-galactopyranose mutase | 3,1 | 0,00 |
| 24 | P05055 | *pnp* | Polyribonucleotide nucleotidyltransferase | 2,9 | 0,00 |
| 25 | P75821 | *ybjS* | Uncharacterized protein YbjS | 2,9 | 0,00 |
| 26 | P26616 | *maeA* | NAD-dependent malic enzyme | 2,9 | 0,00 |
| 27 | P0AE22 | *aphA* | Class B acid phosphatase | 2,8 | 0,00 |
| 28 | P08201 | *nirB* | Nitrite reductase (NADH) large subunit | 2,8 | 0,00 |
| 29 | P0AAG8 | *mglA* | Galactose/methyl galactoside import ATP-binding protein MglA | 2,7 | 0,00 |
| 30 | P0A6R0 | *fabH* | 3-oxoacyl-[acyl-carrier-protein] synthase 3 | 2,7 | 0,00 |

| | | | | | |
|---|---|---|---|---|---|
| **31** | P0A761 | *nanE* | Putative N-acetylmannosamine-6-phosphate 2-epimerase | 2,7 | 0,00 |
| **32** | P37744 | *rfbA* | Glucose-1-phosphate thymidylyltransferase 1 | 2,7 | 0,00 |
| **33** | P37765 | *rluB* | Ribosomal large subunit pseudouridine synthase B | 2,7 | 0,00 |
| **34** | P21363 | *yciE* | Protein YciE | 2,6 | 0,00 |
| **35** | P03014 | *pinE* | Serine recombinase PinE | 2,5 | 0,00 |
| **36** | P0A799 | *pgk* | Phosphoglycerate kinase | 2,5 | 0,00 |
| **37** | P21177 | *fadB* | Fatty acid oxidation complex subunit alpha | 2,5 | 0,00 |
| **38** | P0A993 | *fbp* | Fructose-1,6-bisphosphatase class 1 | 2,5 | 0,00 |
| **39** | P0AFD1 | *nuoE* | NADH-quinone oxidoreductase subunit E | 2,5 | 0,01 |
| **40** | P0A9E0 | *araC* | Arabinose operon regulatory protein | 2,4 | 0,02 |
| **41** | P08956 | *hsdR* | Type I restriction enzyme EcoKI R protein | 2,4 | 0,00 |
| **42** | P0A6Z3 | *htpG* | Chaperone protein HtpG | 2,4 | 0,00 |
| **43** | P0A9L3 | *fklB* | FKBP-type 22 kDa peptidyl-prolyl cis-trans isomerase | 2,4 | 0,00 |
| **44** | P0A9G6 | *aceA* | Isocitrate lyase | 2,4 | 0,00 |
| **45** | P69801 | *manY* | PTS system mannose-specific EIIC component | 2,4 | 0,00 |
| **46** | P0C0V0 | *degP* | Periplasmic serine endoprotease DegP | 2,4 | 0,00 |
| **47** | P0A9B2 | *gapA* | Glyceraldehyde-3-phosphate dehydrogenase A | 2,3 | 0,00 |
| **48** | P0A862 | *tpx* | Thiol peroxidase | 2,3 | 0,00 |
| **49** | P0AGD3 | *sodB* | Superoxide dismutase [Fe] | 2,2 | 0,03 |
| **50** | P0A8M6 | *yeeX* | UPF0265 protein YeeX | 2,2 | 0,00 |
| **51** | P23830 | *pssA* | CDP-diacylglycerol--serine O-phosphatidyltransferase | 2,2 | 0,00 |
| **52** | P31979 | *nuoF* | NADH-quinone oxidoreductase subunit F | 2,2 | 0,00 |
| **53** | P64610 | *yrbL* | Uncharacterized protein YrbL | 2,1 | 0,00 |
| **54** | P75913 | *ghrA* | Glyoxylate/hydroxypyruvate reductase A | 2,1 | 0,00 |
| **55** | P0A7A5 | *pcm* | Protein-L-isoaspartate O-methyltransferase | 2,1 | 0,00 |
| **56** | P76010 | *ycgR* | Flagellar brake protein | 2,1 | 0,00 |
| **57** | P0AEZ9 | *moaB* | Molybdenum cofactor biosynthesis protein B | 2,1 | 0,00 |
| **58** | P09372 | *grpE* | HSP-70 cofactor | 2,1 | 0,00 |
| **59** | P10441 | *lpxB* | Lipid-A-disaccharide synthase | 2,1 | 0,00 |
| **60** | P0A870 | *talB* | Transaldolase B | 2,1 | 0,00 |
| **61** | P0AF26 | *narJ* | Nitrate reductase molybdenum cofactor assembly chaperone | 2,1 | 0,03 |
| **62** | P0AC59 | *grxB* | Glutaredoxin 2 | 2,0 | 0,00 |
| **63** | P0AF24 | *nagD* | Ribonucleotide monophosphatase NagD | 2,0 | 0,00 |
| **64** | P0A6H1 | *clpX* | ATP-dependent Clp protease ATP-binding subunit | 2,0 | 0,00 |
| **65** | P0AC53 | *zwf* | Glucose-6-phosphate 1-dehydrogenase | 2,0 | 0,00 |

| | | | | | |
|---|---|---|---|---|---|
| **66** | P11868 | *tdcD* | Propionate kinase | 2,0 | 0,00 |

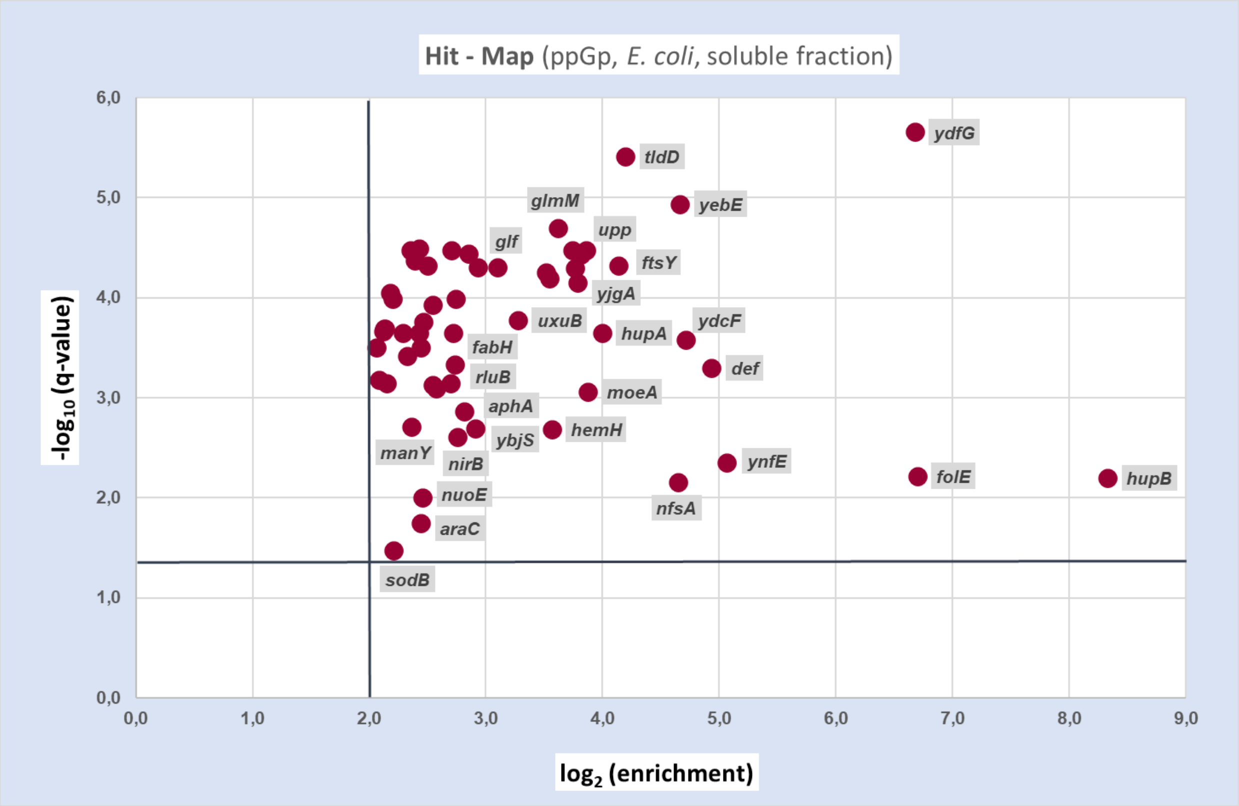


***Supporting figure 11:*** *graphic representation of proteins captured by ppGp-CC from the E. coli soluble fraction. Only hits complying the threshold criteria are shown in the plot.*

#### 4.2. *E.* coli (membrane fraction)

##### 4.2.1. ppGpp-CC1 (membrane fraction)

***Supporting table 11:*** *putative ppGpp receptors captured by ppGpp-CC1 from the membrane fraction of E. coli cell lysate.*

| Nr. | ac | gene | protein | $log_2$ (enrichment) | q-value |
|---|---|---|---|---|---|
| 1 | P0A9M5 | *gpt* | Xanthine phosphoribosyltransferase | 8,1 | 0,00 |
| 2 | P0ACY1 | *ydjA* | Putative NAD(P)H nitroreductase | 7,3 | 0,00 |
| 3 | P69451 | *fadD* | Long-chain-fatty-acid--CoA ligase | 6,8 | 0,02 |
| 4 | P0ABS1 | *dksA* | RNA polymerase-binding transcription factor | 6,7 | 0,04 |
| 5 | P0A6B7 | *iscS* | Cysteine desulfurase | 6,3 | 0,04 |
| 6 | P62707 | *gpmA* | 2,3-bisphosphoglycerate-dependent phosphoglycerate mutase | 5,4 | 0,00 |
| 7 | P75824 | *hcr* | NADH oxidoreductase | 5,3 | 0,00 |
| 8 | P0A6A3 | *ackA* | Acetate kinase | 4,4 | 0,01 |
| 9 | P0ABF1 | *pcnB* | Poly(A) polymerase I | 4,3 | 0,03 |
| 10 | P0AAF6 | *artP* | Arginine transport ATP-binding protein | 4,2 | 0,00 |
| 11 | P0AC23 | *focA* | Probable formate transporter 1 | 3,8 | 0,00 |
| 12 | P0AG67 | *rpsA* | 30S ribosomal protein S1 | 3,6 | 0,02 |
| 13 | P0A744 | *msrA* | Peptide methionine sulfoxide reductase | 3,3 | 0,01 |
| 14 | P0A862 | *tpx* | Thiol peroxidase | 3,2 | 0,00 |
| 15 | P76346 | *mtfA* | uncharacterized protein | 3,1 | 0,04 |
| 16 | P09833 | *modC* | Molybdenum import ATP-binding protein | 3,1 | 0,00 |
| 17 | P0A9U1 | *ybhF* | probable multidrug ABC transporter ATP-binding protein | 2,9 | 0,00 |
| 18 | P23871 | *hemH* | Ferrochelatase | 2,7 | 0,00 |
| 19 | P00579 | *rpoD* | RNA polymerase sigma factor | 2,7 | 0,02 |
| 20 | P0ABB0 | *atpA* | ATP synthase subunit alpha | 2,7 | 0,00 |
| 21 | P0A6K6 | *deoB* | Phosphopentomutase | 2,6 | 0,02 |
| 22 | P0ABT2 | *dps* | DNA protection during starvation protein | 2,5 | 0,01 |
| 23 | P64604 | *mlaD* | Intermembrane phospholipid transport system binding protein | 2,5 | 0,00 |
| 24 | P08200 | *icd* | Isocitrate dehydrogenase | 2,5 | 0,00 |
| 25 | P0ADU5 | *ygiW* | uncharacterized protein | 2,3 | 0,01 |
| 26 | P63386 | *mlaF* | Intermembrane phospholipid transport system ATP-binding protein | 2,3 | 0,01 |
| 27 | P37624 | *rbbA* | Ribosome-associated ATPase | 2,3 | 0,01 |
| 28 | P0AE52 | *bcp* | Peroxiredoxin | 2,2 | 0,00 |

| | | | | | |
|---|---|---|---|---|---|
| **29** | P0AAG8 | *mglA* | Galactose/methyl galactoside import ATP-binding protein | 2,2 | 0,01 |
| **30** | P21645 | *lpxD* | UDP-3-O-(3-hydroxymyristoyl)glucosamine N-acyltransferase | 2,1 | 0,01 |
| **31** | P37330 | *glcB* | Malate synthase G | 2,1 | 0,00 |

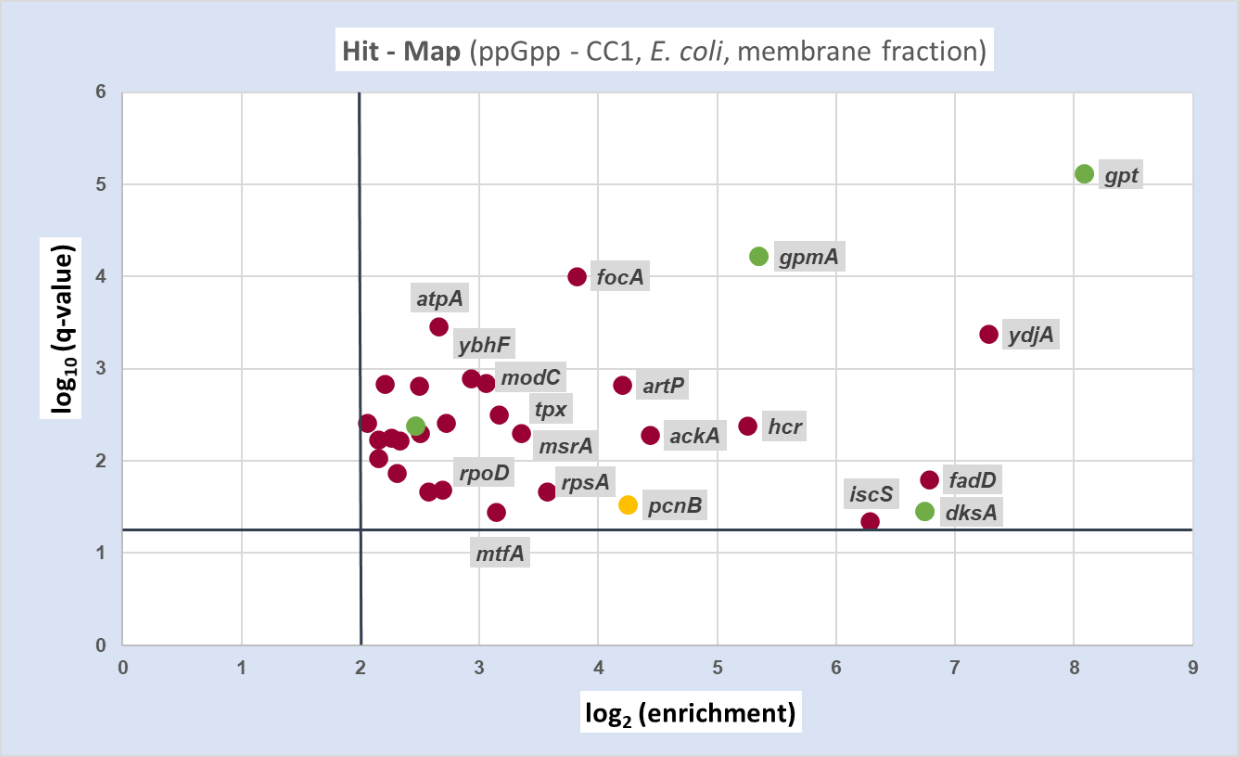


***Supporting figure 12:*** *graphic representation of proteins captured by ppGpp-CC1 from the E. coli membrane fraction. Only hits complying the threshold criteria are shown in the plot.*

##### 4.2.2. ppGpp-CC2 (membrane fraction)

***Supporting table 12:*** *putative ppGpp receptors captured by ppGpp-CC2 from the membrane fraction of E. coli cell lysate. An extract of this table was already shown in the main part of the manuscript.*

| **Nr.** | **ac** | **gene** | **protein** | **$log_2$ (enrichment)** | **q-value** |
|---|---|---|---|---|---|
| 1 | P71298 | *intF* | Prophage integrase | 7,1 | 0,00 |
| 2 | P0AE22 | *aphA* | Class B acid phosphatase | 6,0 | 0,00 |
| 3 | P02918 | *mrcA* | Penicillin-binding protein 1A | 5,1 | 0,02 |
| 4 | P0AFI7 | *pdxH* | Pyridoxine/pyridoxamine 5'-phosphate oxidase | 4,9 | 0,00 |
| 5 | P0ABF1 | *pcnB* | Poly(A) polymerase I | 4,4 | 0,00 |
| 6 | P76503 | *fadI* | 3-ketoacyl-CoA thiolase | 4,2 | 0,01 |
| 7 | P0A6A3 | *ackA* | Acetate kinase | 4,0 | 0,00 |
| 8 | P0A9Y6 | *cspC* | Cold shock-like protein | 3,6 | 0,00 |
| 9 | P0ABB0 | *atpA* | ATP synthase subunit alpha | 3,4 | 0,00 |
| 10 | P64604 | *mlaD* | Intermembrane phospholipid transport system binding protein | 3,3 | 0,01 |
| 11 | P0ABT2 | *dps* | DNA protection during starvation protein | 3,2 | 0,00 |
| 12 | P0AE52 | *bcp* | Peroxiredoxin | 3,1 | 0,00 |
| 13 | P37692 | *rfaF* | ADP-heptose--LPS heptosyltransferase 2 | 2,8 | 0,01 |
| 14 | P09833 | *modC* | Molybdenum import ATP-binding protein | 2,7 | 0,00 |
| 15 | P77399 | *fadJ* | Fatty acid oxidation complex subunit alpha | 2,7 | 0,00 |
| 16 | P77304 | *dtpA* | Dipeptide and tripeptide permease A | 2,4 | 0,01 |
| 17 | P0A6M8 | *fusA* | Elongation factor G | 2,3 | 0,00 |
| 18 | P63386 | *mlaF* | Intermembrane phospholipid transport system ATP-binding protein | 2,3 | 0,01 |
| 19 | P0ADU5 | *ygiW* | uncharacterized protein | 2,2 | 0,04 |
| 20 | P23173 | *tnaB* | Low affinity tryptophan permease | 2,2 | 0,00 |
| 21 | P0AFP4 | *ybbO* | Uncharacterized oxidoreductase | 2,2 | 0,00 |
| 22 | P62517 | *mdoH* | Glucans biosynthesis glucosyltransferase H | 2,2 | 0,00 |
| 23 | P11557 | *damX* | Cell division protein | 2,2 | 0,00 |
| 24 | P0A862 | *tpx* | Thiol peroxidase | 2,2 | 0,00 |
| 25 | P37765 | *rluB* | Ribosomal large subunit pseudouridine synthase B | 2,2 | 0,00 |
| 26 | P08200 | *icd* | Isocitrate dehydrogenase [NADP] | 2,1 | 0,00 |
| 27 | P11880 | *murF* | UDP-N-acetylmuramoyl-tripeptide--D-alanyl-D-alanine ligase | 2,1 | 0,00 |
| 28 | P27836 | *wecG* | UDP-N-acetyl-D-mannosaminuronic acid transferase | 2,1 | 0,00 |
| 29 | P45537 | *yhfK* | uncharacterized protein YhfK | 2,0 | 0,01 |
| 30 | P0A972 | *cspE* | cold shock-like protein | 2,0 | 0,00 |

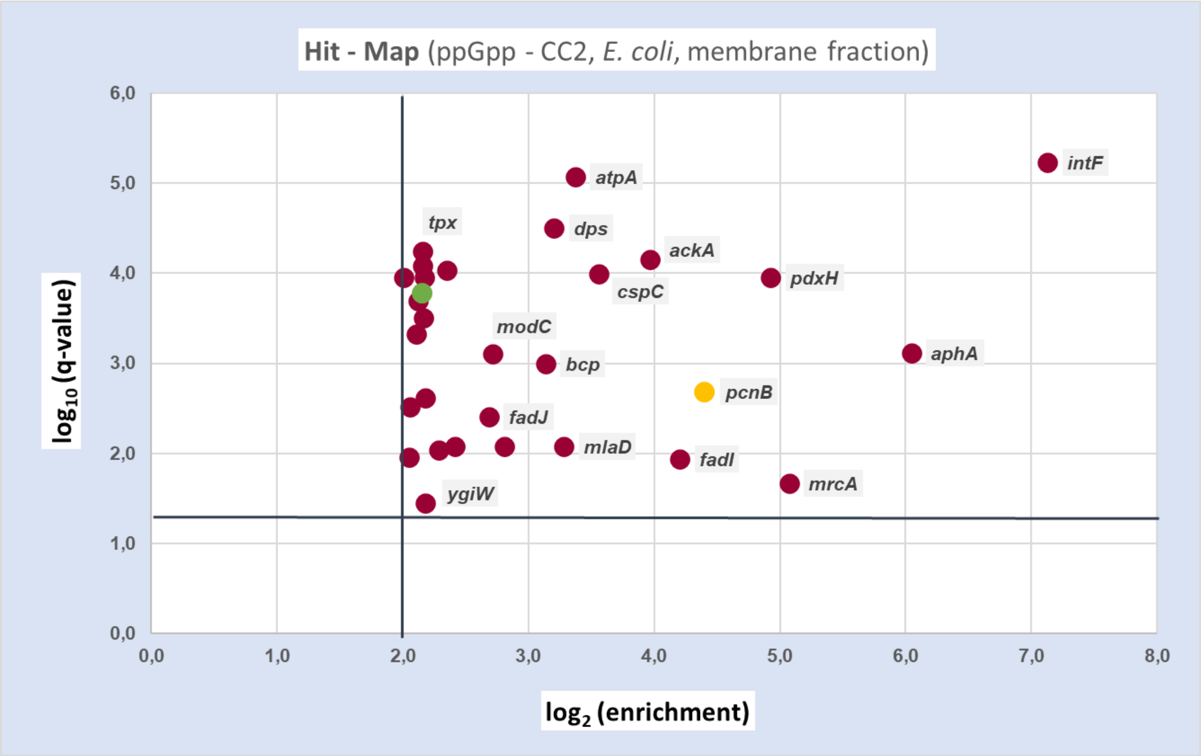


***Supporting figure 13:*** *graphic representation of proteins captured by ppGpp-CC2 from the E. coli membrane fraction. Only hits complying the threshold criteria are shown in the plot.*

#### 4.3. *S. typhimurium* (soluble fraction)

***Supporting table 13:*** *putative ppGpp receptors captured by a mixture of ppGpp-CC1 and ppGpp-CC2 from the soluble fraction of S. typhimurium cell lysate.*

| Nr. | ac | gene | protein | $log_2$ (enrichment) | q-value |
|---|---|---|---|---|---|
| 1 | E1WH24 | *rfbG* | CDP-glucose 4,6-dehydratase | 4,5 | 0,00 |
| 2 | E1WEU6 | *ssb* | Single-stranded DNA-binding protein | 4,0 | 0,00 |
| 3 | E1WHL6 | *tktA* | Transketolase | 3,6 | 0,00 |
| 4 | E1WEE7 | *hslV* | ATP-dependent protease subunit | 3,5 | 0,00 |
| 5 | E1WET0 | *SL1344_4176* | Putative uncharacterized protein | 3,5 | 0,00 |
| 6 | E1W7A8 | *mgsA* | Methylglyoxal synthase | 3,4 | 0,00 |
| 7 | E1W9S6 | *hutG* | Formimidoylglutamase | 3,1 | 0,00 |
| 8 | E1W883 | *yaeH* | UPF0325 protein yaeH | 3,0 | 0,00 |
| 9 | E1WIM3 | *fusA* | Elongation factor G | 2,9 | 0,00 |
| 10 | E1WFA8 | *icdA* | Isocitrate dehydrogenase [NADP] | 2,8 | 0,00 |
| 11 | E1WBB1 | *SL1344_4508* | Conserved hypothetical ABC transporter | 2,8 | 0,00 |
| 12 | E1W9J2 | *pgm* | Phosphoglucomutase | 2,8 | 0,02 |
| 13 | E1WAK6 | *ygcX* | Probable glucarate dehydratase 1 | 2,7 | 0,00 |
| 14 | E1WIE2 | *mdh* | Malate dehydrogenase OS=Salmonella typhimurium (strain SL1344) GN=mdh | 2,7 | 0,01 |
| 15 | E1WFF4 | *SL1344_1223* | Putative oxidoreductase | 2,6 | 0,00 |
| 16 | E1WFF1 | *SL1344_1220* | Putative uncharacterized protein | 2,5 | 0,00 |
| 17 | E1W9W7 | *dps* | DNA protection during starvation protein | 2,5 | 0,00 |
| 18 | E1WI76 | *glmM* | Phosphoglucosamine mutase | 2,5 | 0,00 |
| 19 | E1WE88 | *fdhE* | Protein FdhE | 2,5 | 0,00 |
| 20 | E1WDY2 | *trxA* | Thioredoxin | 2,5 | 0,00 |
| 21 | E1W7M8 | *yaaA* | UPF0246 protein yaaA | 2,4 | 0,00 |
| 22 | E1W713 | *himD* | Integration host factor subunit beta | 2,4 | 0,00 |
| 23 | E1W8R3 | *SL1344_0397* | Probable peroxidase | 2,4 | 0,00 |
| 24 | E1WF16 | *aspA* | Aspartate ammonia-lyase | 2,4 | 0,00 |
| 25 | E1WF24 | *efp* | Elongation factor P | 2,3 | 0,00 |
| 26 | E1W731 | *asnS* | Asparaginyl-tRNA synthetase | 2,3 | 0,00 |
| 27 | E1W9M3 | *gltA* | Citrate synthase | 2,3 | 0,00 |
| 28 | E1WFJ3 | *pfkB* | 6-phosphofructokinase isozyme | 2,2 | 0,00 |
| 29 | E1WHK9 | *pgk* | Phosphoglycerate kinase | 2,2 | 0,00 |

| | | | | | |
|---|---|---|---|---|---|
| **30** | E1WFJ1 | *SL1344_1259* | Putative uncharacterized protein | 2,1 | 0,00 |
| **31** | E1WGW6 | *cbiC* | Precorrin-8X methylmutase | 2,1 | 0,00 |
| **32** | E1WHI0 | *lysS* | Lysyl-tRNA synthetase | 2,1 | 0,00 |
| **33** | E1WHV3 | *dkgA* | 2,5-diketo-D-gluconic acid reductase A | 2,0 | 0,00 |
| **34** | E1WCL8 | *fabB* | 3-oxoacyl-[acyl-carrier-protein] synthase I | 2,0 | 0,00 |
| **35** | E1WAJ7 | *pyrG* | CTP synthase | 2,0 | 0,00 |
| **36** | E1W8Q3 | *rdgC* | Recombination-associated protein rdgC | 2,0 | 0,00 |
| **37** | E1W7J4 | *grxB* | Glutaredoxin 2 | 2,0 | 0,00 |

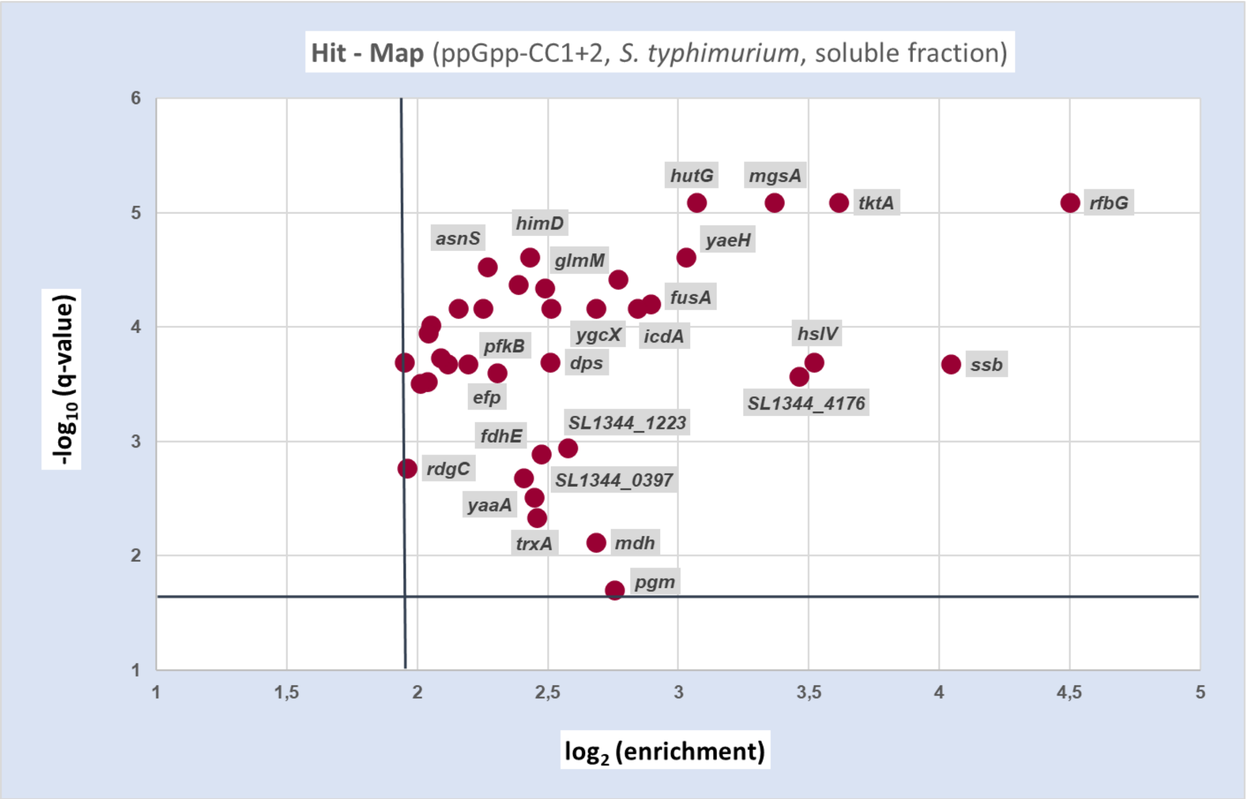


***Supporting figure 14:*** *graphic representation of proteins captured by a mixture of ppGpp-CC1 and ppGpp-CC2 from the S. typhimurium soluble fraction. Only hits complying the threshold criteria are shown in the plot.*

### 5. Target validation: Bis(5’-nucleosyl)-tetraphosphatase ApaH is regulated by Magic Spot Nucleotides in vitro

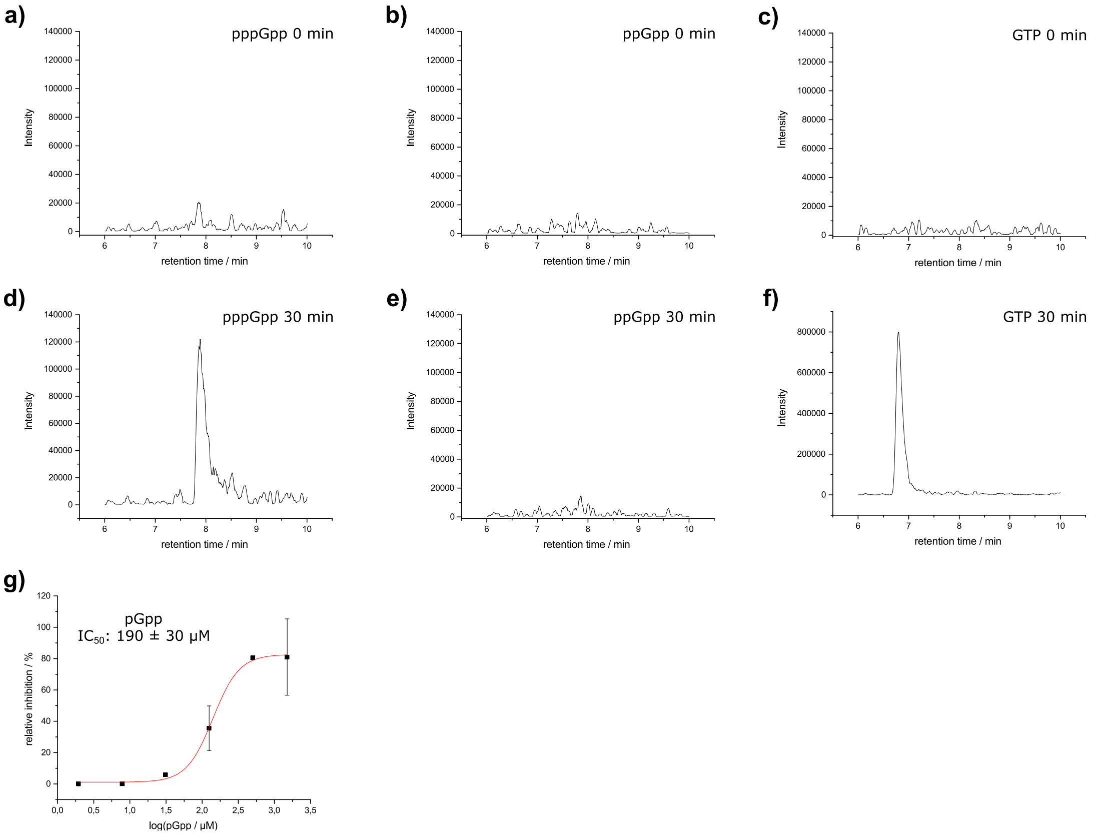


***Supporting Figure 15:** Biochemical characterization of MSN target ApaH: **a) and d)**: extracted ion chromatogram for 522 m/z (triphosphorylated guanosine) of ApaH treated pppGpp for given reaction time. **b) and e)**: extracted ion chromatogram for 442 m/z (diphosphorylated guanosine) of ApaH treated ppGpp for given reaction time. **c) and f)**: extracted ion chromatogram for 362 m/z (monophosphorylated guanosine) of ApaH treated GTP for given reaction time. Representative chromatogram of three experiments. **g)** representative $IC_{50}$ curve of pGpp on diadenosine tetraphosphate hydrolysis by ApaH. n=2. Experiments were independently repeated with n=2. Error bars indicate standard deviation. Insert: average and standard deviation of independent experiments.*

#### 5.1. ApaH in vitro assay

The protocol was derived from Guranowski et al.[8] In brief, for $IC_{50}$ experiments, 20 µl reaction buffer (62.5 mM Hepes [4-(2-hydroxyethyl)-1-piperazineethanesulfonic acid] pH 7.5, 150 µM $CoCl_2$, 1.25 mM 2-mercaptoethanol), 3 µl 243 µM diadenosine tetraphosphate (Ap4A) and 8 µl $H_2O$ or inhibitor dissolved in $H_2O$ were mixed. 2 µl ApaH solution (4.3 µg/ml in 10% glycerol, 5 mM 2-mercaptoethanol, 2 mg/ml bovine serum albumin) were added and the reaction mixture were incubated for 60 s at 37 °C and 600 rpm on a thermocycler. For degradation experiments using other nucleotides than Ap4A, 6 µl of 304 µM substrate were used with 125 µM CoCl2 in the reaction buffer and incubated for 30 min. The reaction was quenched by addition of 3 µl 3 mM alkalized ethylenediaminetetraacetic acid solution.

#### 5.2. LC/MS analysis of nucleotides

30 µl acetonitrile was added to the quenched reaction mixture and after vortexing, the samples were centrifuged for 15 min at 4 °C and 20,000 g. The supernatants were transferred to a vial with glass insert.

4 µl of the samples were injected on a Hilic-Z, PEEK-lined column (Agilent technologies, 100 x 2.1 mm) operated at 35 °C. The gradient was: 90 % B (acetonitrile) for 0.5 min, to 40 % B within 11.5 min, hold for 2 min, to 90 % B within 0.5 min, hold for 5.5 min. Solvent A was 20 mM ammonium acetate pH 9. The flow rate was set to 0.4 ml/min. The LC was coupled with a QqQ-MS (Agilent Technologies: G4220A, G4226A, G1316A, G6460A) with the following parameters: Gas temperature 310 °C at 10.7 L/min, sheath gas temperature 310 °C with 10 L/min flow, nebulizer pressure of 30 psi, capillary voltage of 4 kV in negative mode, delta EMV of 400 and 2 kV nozzle voltage. Data were acquired by Agilent MassHunter Data Acquisition (version B.08.02) and analyzed with Agilent MassHunter Qualitative Analysis (version B.07.00, SP1).

#### 5.3. $IC_{50}$ - determination

Each $IC_{50}$ assay was performed with two reactions per condition and reproduced at least once (total n=4). Relative intensity of Ap4A to EDTA treated reactions were determined and the average of each condition was plotted against the $log_{10}$ concentration of the inhibitor. OriginPro 2017 was used to fit the sigmoidal dose/response curves. The derived functions were used to determine the absolute $IC_{50}$ values. Averaged $IC_{50}$ values are given with corresponding standard deviation.

#### 5.4. Determination of kinetic parameters ($K_M$ and $K_{cat}$)

For $K_M$ and $k_{cat}$ determination, 6 µl of GTP- or pppGpp-solutions were added at different concentrations to 20 µl reaction buffer (62.5 mM Hepes [4-(2-hydroxyethyl)-1-piperazineethanesulfonic acid] pH 7.5, 125 µM CoCl2, 1.25 mM 2-mercaptoethanol) and 2 µl ApaH. The reaction was incubated at 37 °C and 650 rpm and terminated by addition of EDTA after 30 min. Product concentration was determined by multiple reaction monitoring LC/MS using a standard curve. $K_M$ and $v_{max}$ was determined by Origin 9 using the Michaelis-Menten function. $K_{cat}$ was calculated by dividing $v_{max}$ by the enzyme concentration (9.8 nM).

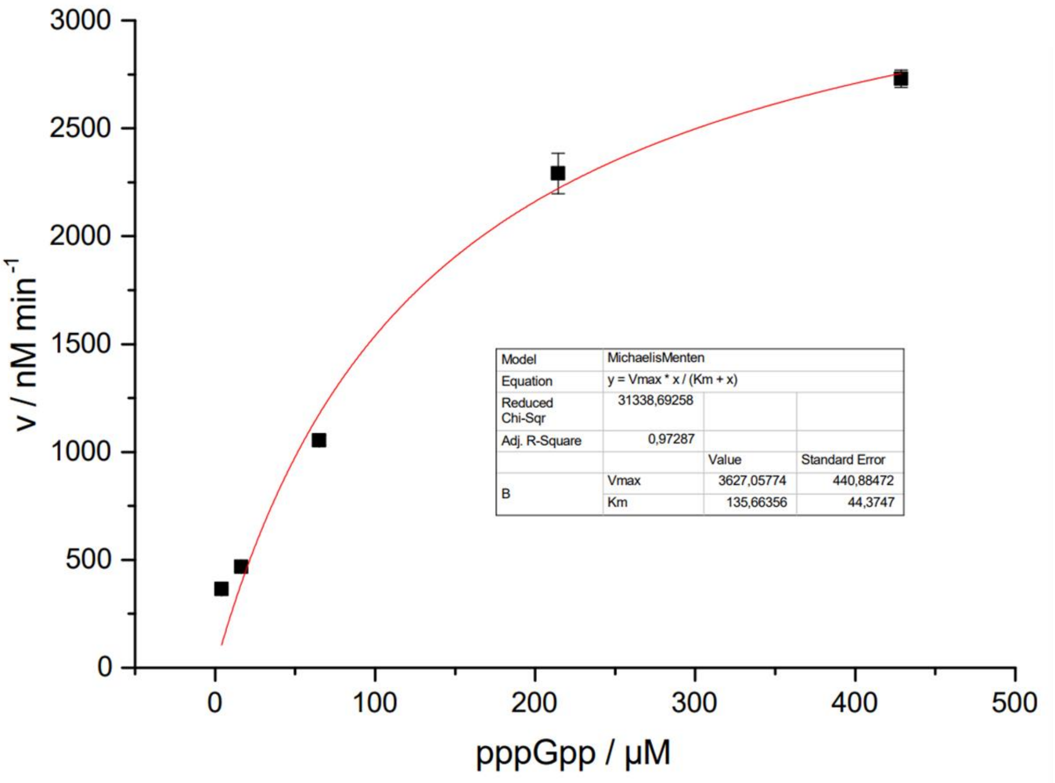


***Supporting Figure 16:*** *Michaelis-Menten model for ApaH-catalyzed hydrolysis of pppGpp towards pGpp.*

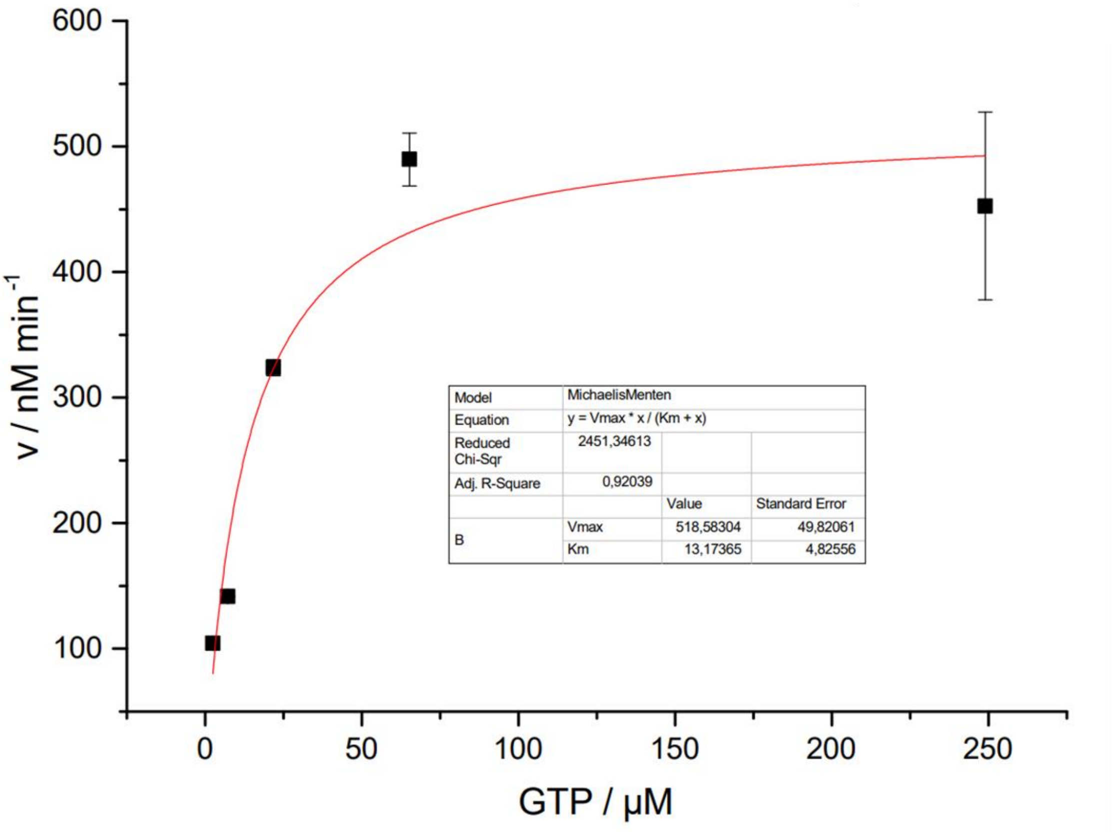


***Supporting Figure 17:*** *Michaelis-Menten Fit for ApaH-catalyzed hydrolysis of GTP towards GMP.*

#### 5.5. Molecular docking of ApaH substrates with AlphaFold

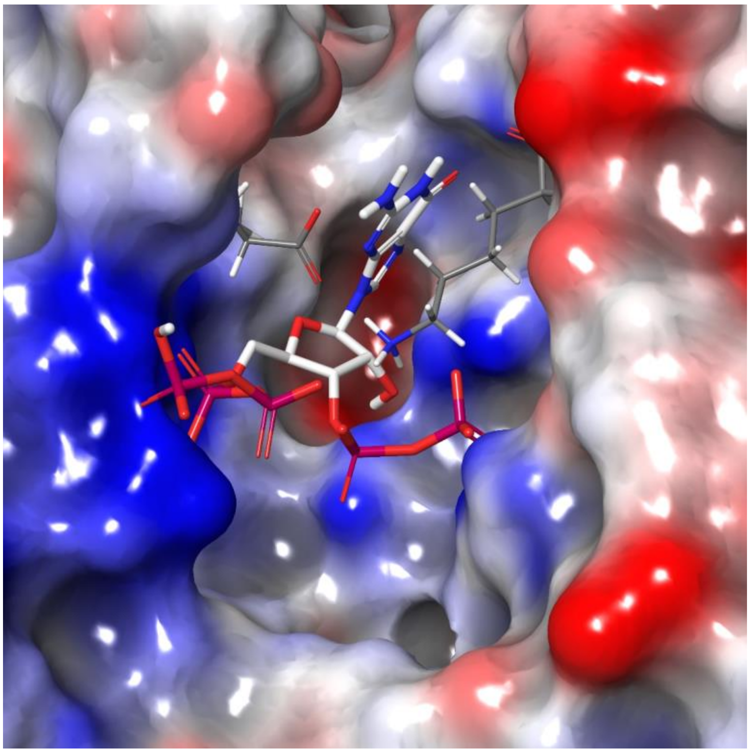

***Supporting Figure 18:*** *Docking of pppGpp into the active site of the AlphaFold model of E. coli ApaH (AF-P05637-F1).[9] The surface of the covering amino acids Lys 197 and Ser280 (C-terminus) has been removed to increase visibility. The surface is colored according to the electrostatic potential, with negative charge in red and positive chare in blue. The model shown is the best scored pose obtained from the Induced Fit protocol of the Schrödinger Platform using default parameters.[10] The following residues of ApaH have been included: No. 8, 10, 41, 65, 66, 69, 81, 83, 84, 155, 156, 158, 182, 184, 193, 194, 196, 197, 198, 227, 228, 229, 230, 232, 244, 245, 246, 247, 248, 249, 251, 272, 274, 277, 278, 280.*

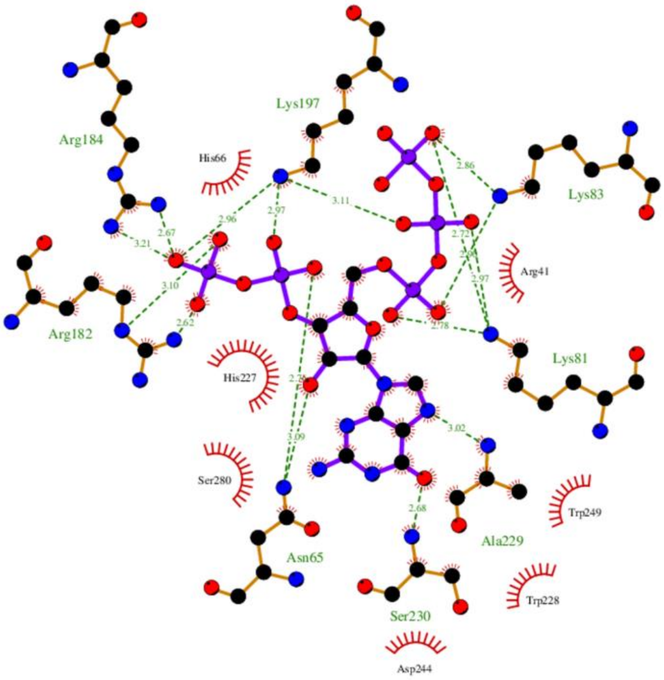


***Supporting Figure 19:*** *Two-dimensional representation of pppGpp coordination in the ApaH active site created with with LigPlot.[11]*

Induced fit docking of pppGpp into the AlphaFold model of *E. coli* ApaH (AF-P05637-F1) resulted in a model in which the pppGpp is deeply buried in the active site (Supporting Figure 16). The diphosphate moiety binds to Arg182 and Arg184, while the triphosphate chain coordinates to Lys81 and Lys83 (Supporting Figure 19). In contrast to ADP, equipped with a single pyrophosphate group, pppGpp can bind in a bidentate manner, strongly increasing complex stability. The guanine residue is coordinated to Ser230 (via carbonyl oxygen) and Ala229. However, this binding is not essential, as other strongly binding poses exist, where guanine is located in the active site's 'lower' part (Supporting Figure 18).

### *7. NMR - spectra*

***(sorted according to molecule numbering)***

Compound 5 (HMDA-pGp), $^{1}H$ - NMR ($D_2O$, 400 MHz)



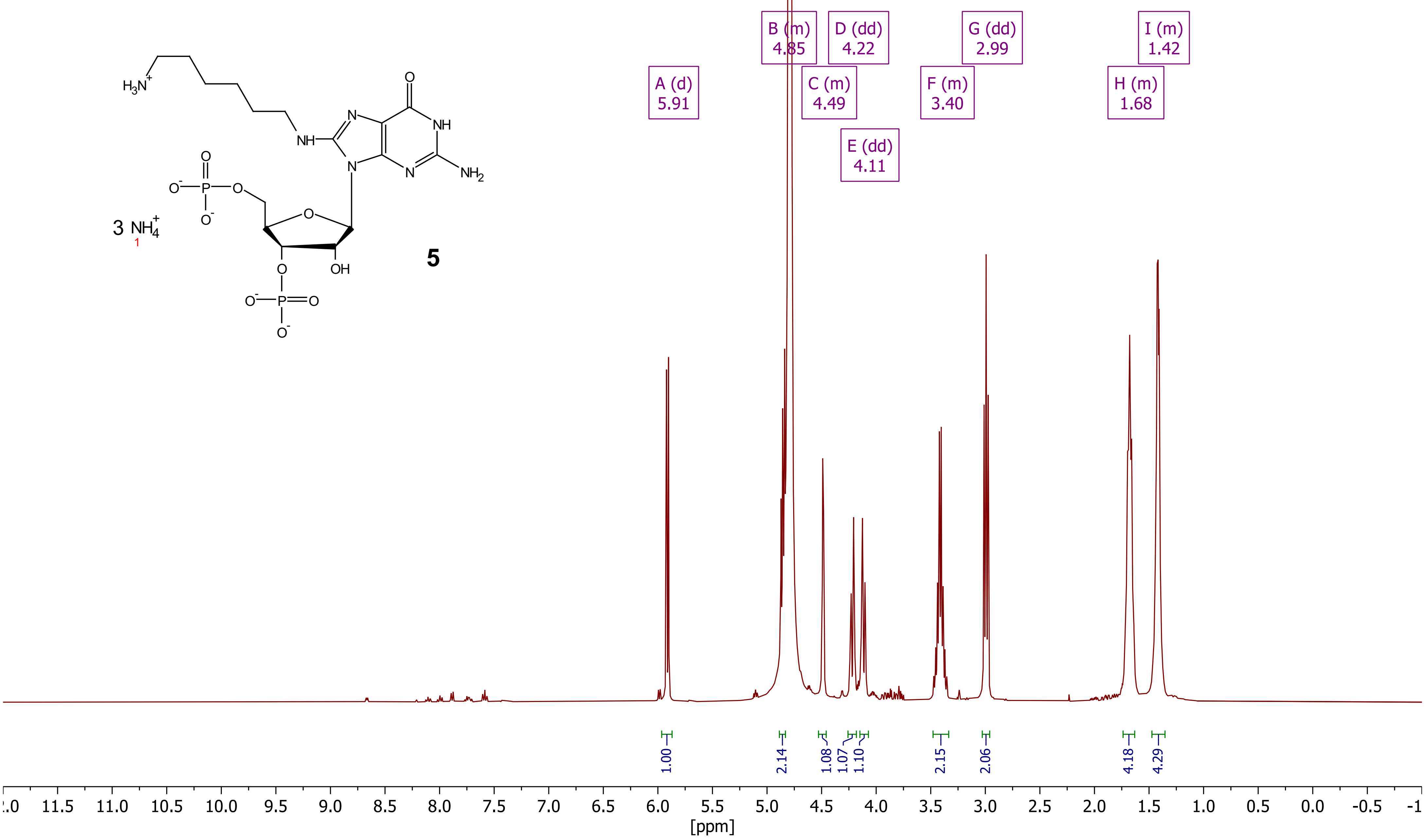

***Compound 5 (HMDA-pGp), $^{31}P[^{1}H]$ - NMR ($D_2O$, 162 MHz)***

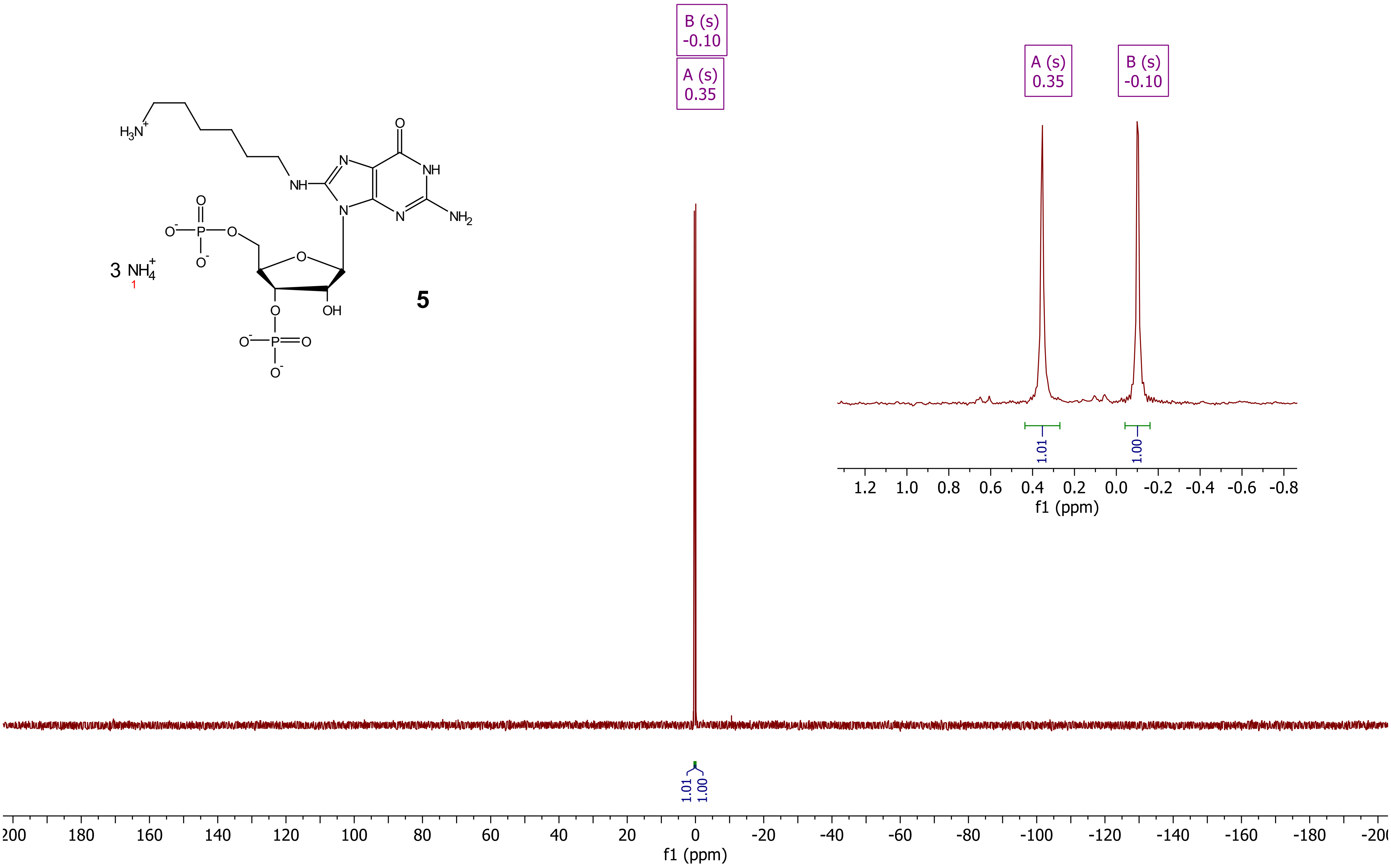

***Compound 5 (HMDA-pGp), $^{13}C[^{1}H]$ - NMR ($D_2O$, 101 MHz)***

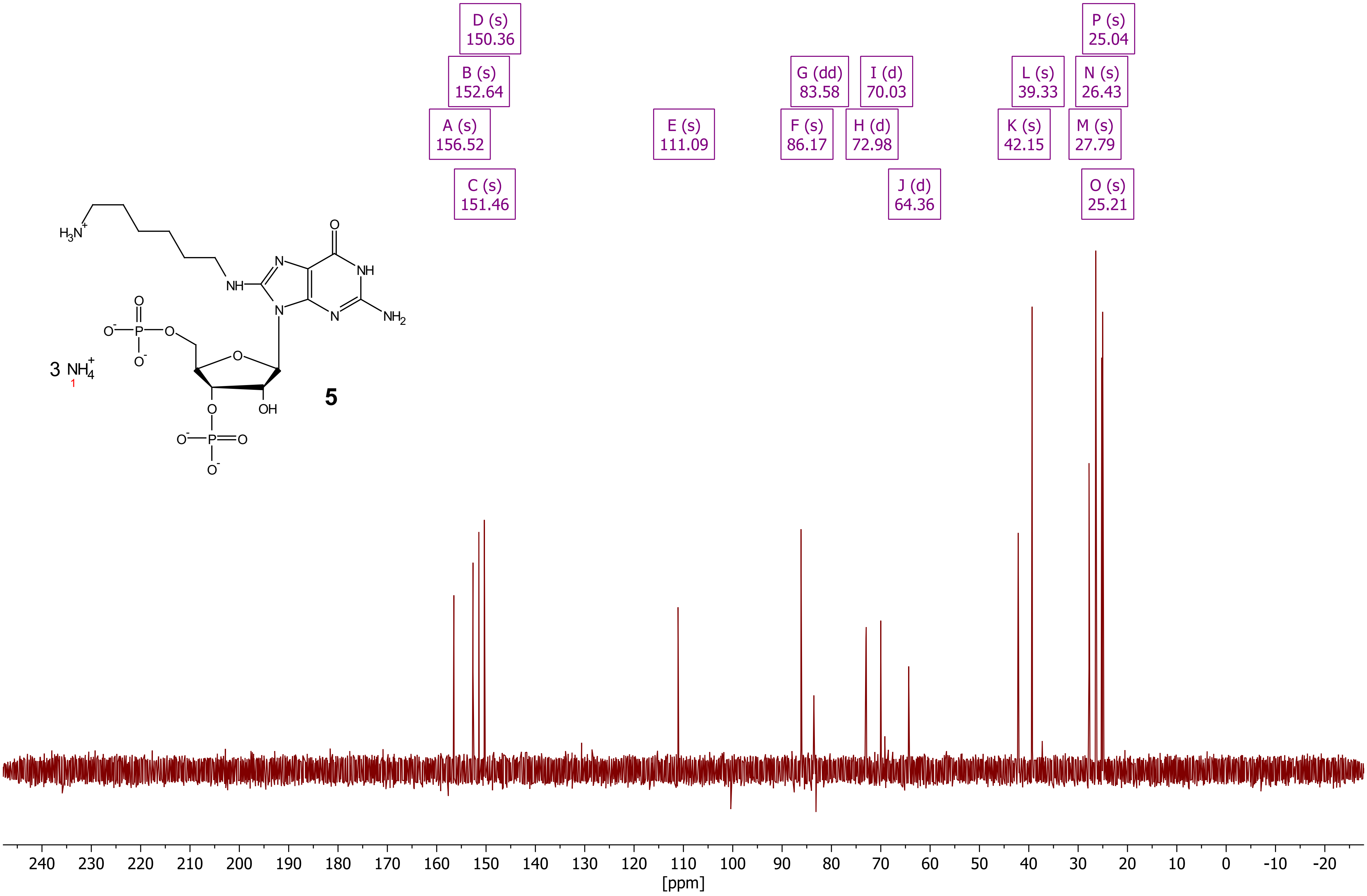


**5**

*Compound 5 (HMDA-pGp), DQF-COSY ($D_2O$)*

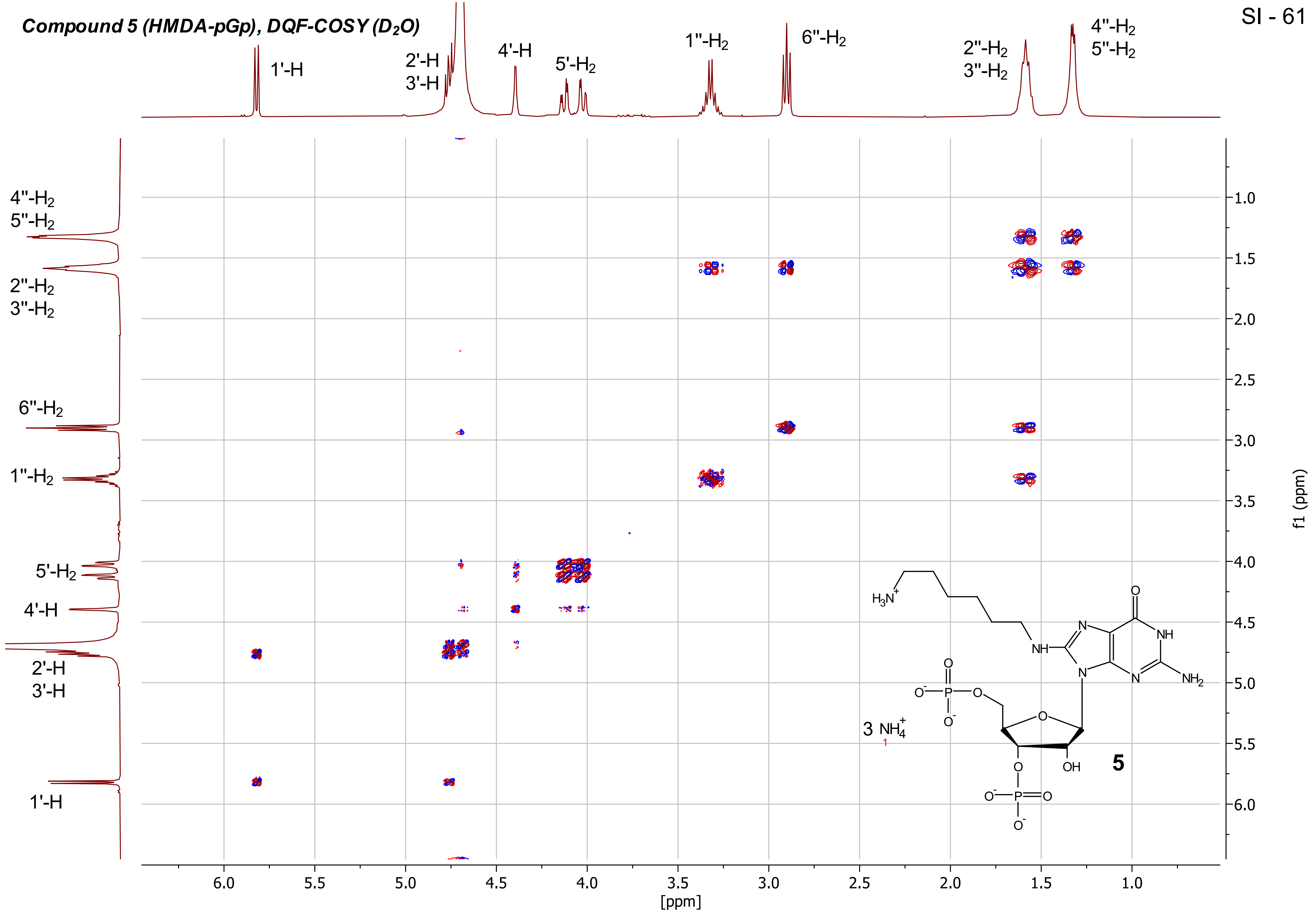

***Compound 5 (HMDA-pGp), HSQC ($D_2O$)***

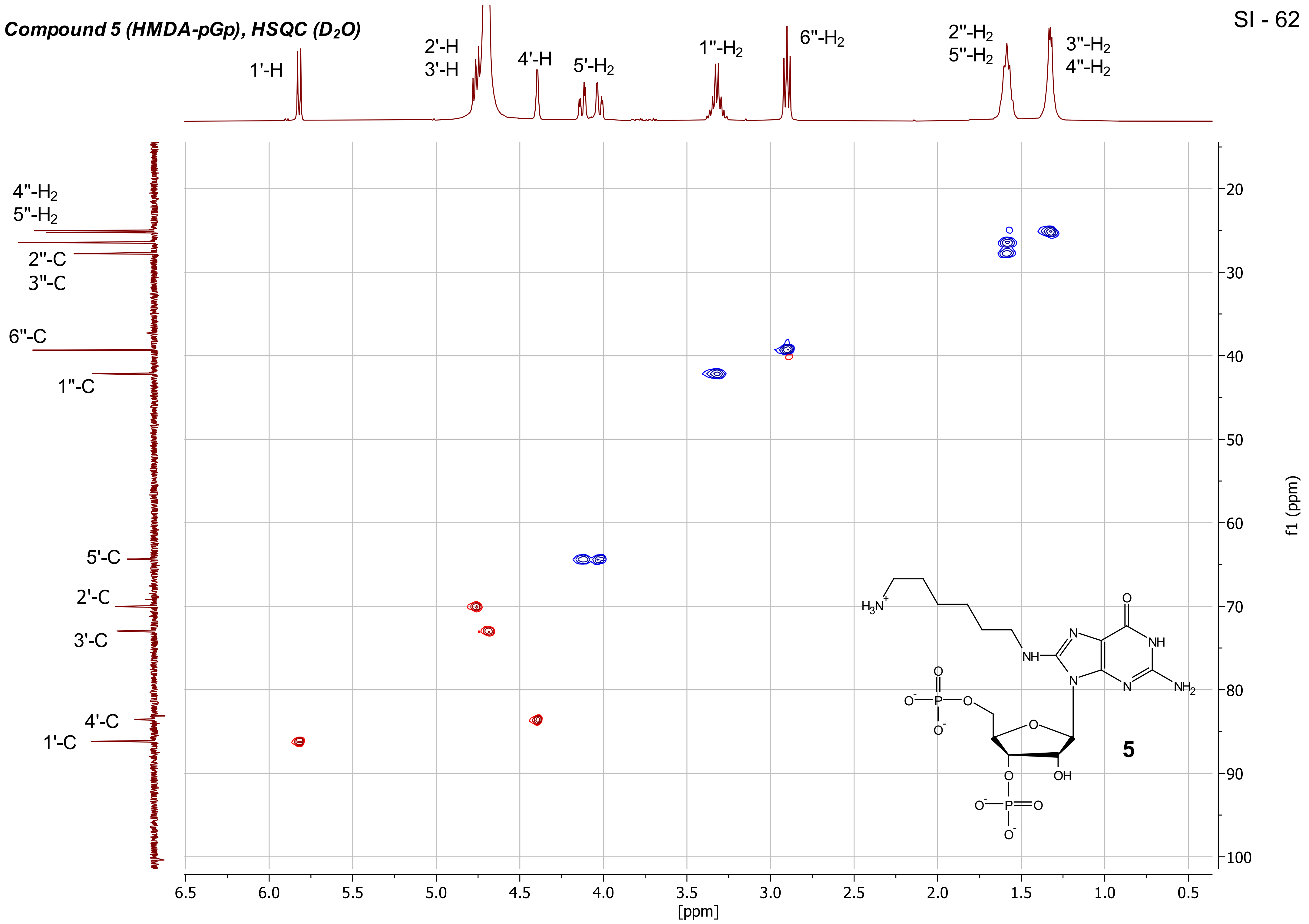

***Compound 5 (HMDA-pGp), $^{31}P$ - $^{1}H$ - HMBC ($D_2O$)***

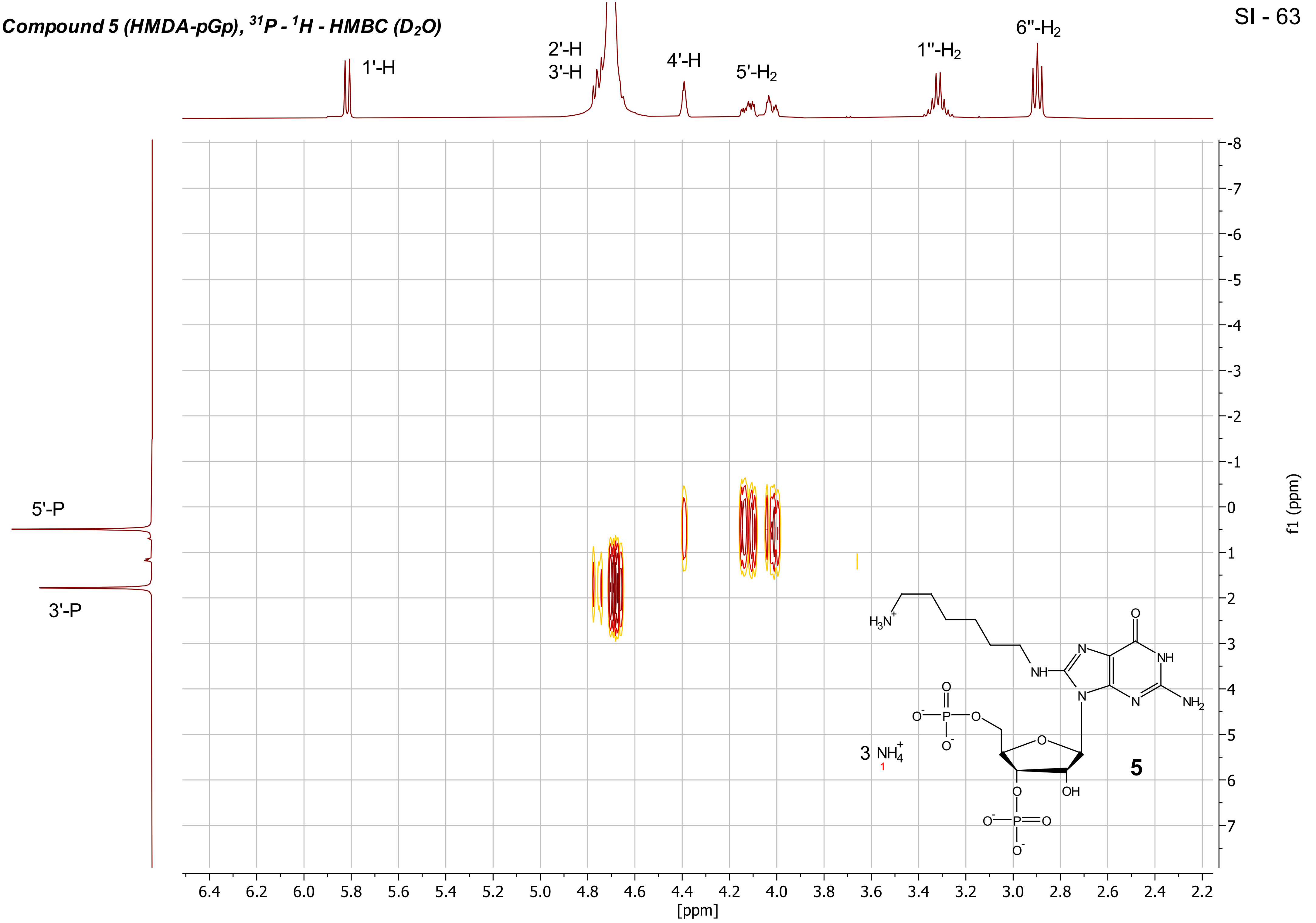

***Compound 6 (Fmoc-Gly-HMDA-pGp), $^{1}H$ - NMR ($D_2O$, 400 MHz)***

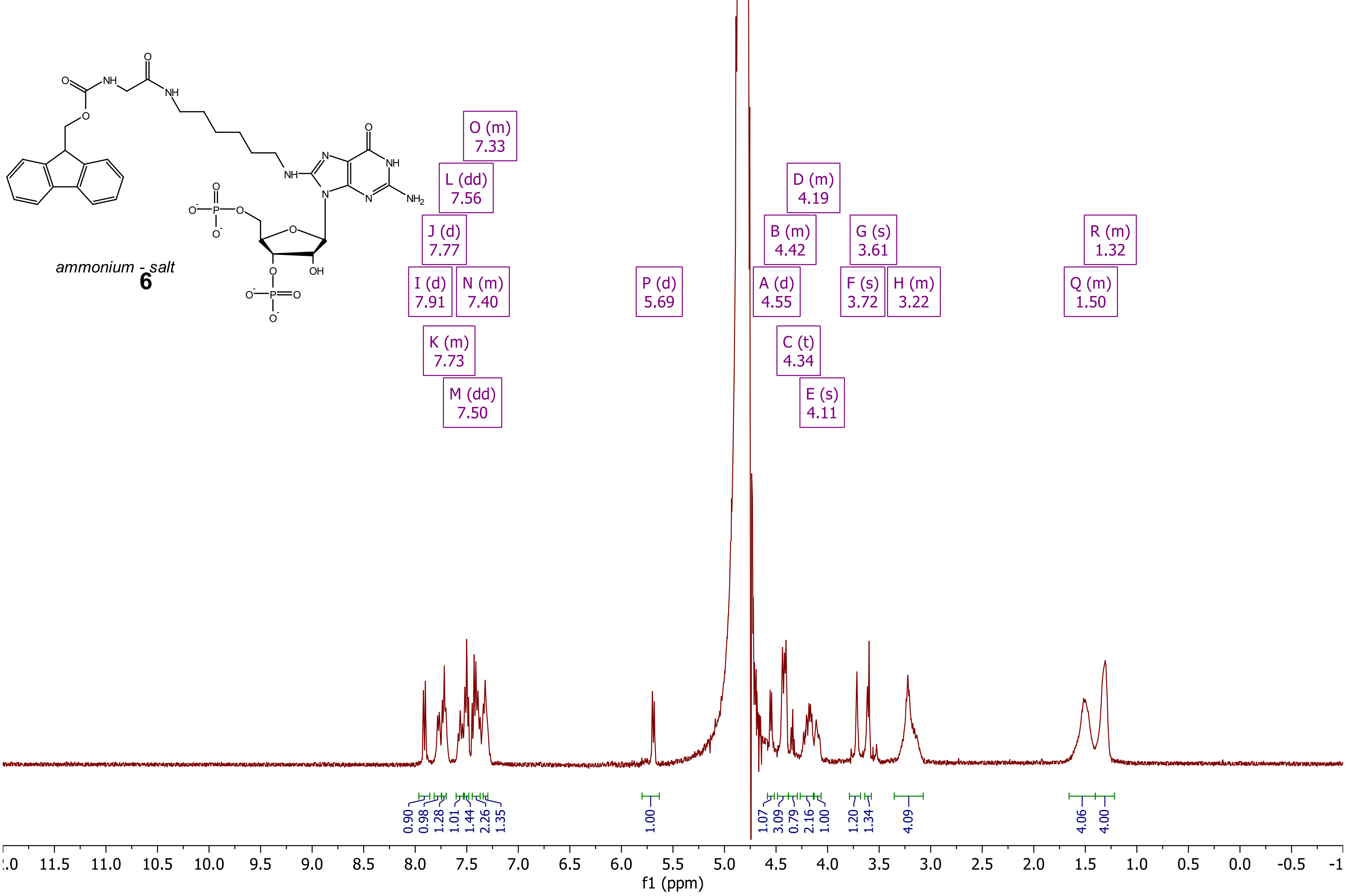

##### *Compound 6 (Fmoc-Gly-HMDA-pGp), $^{31}P\{^{1}H\}$ - NMR ($D_2O$, 400 MHz)*

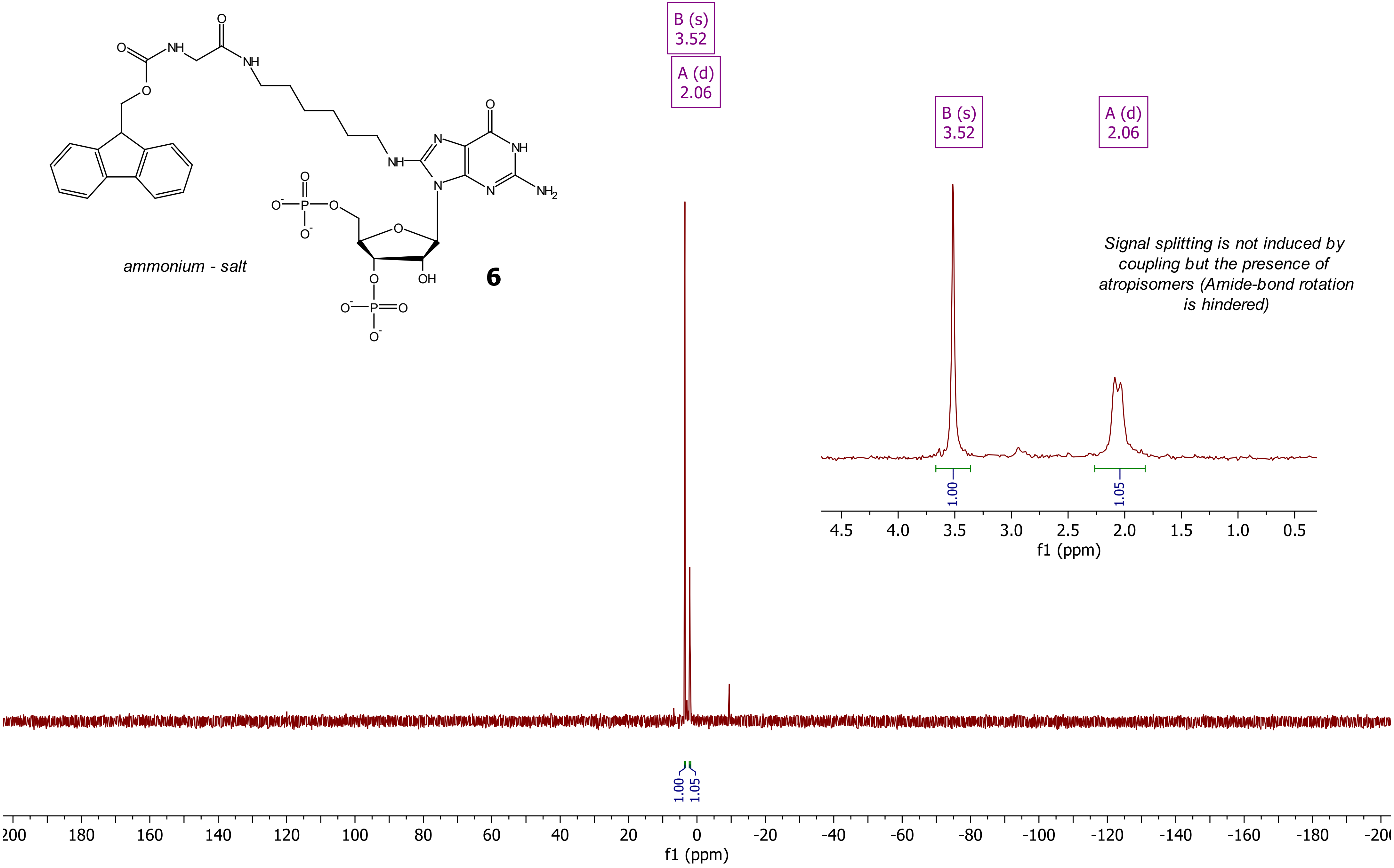

***Compound 7 (Aminoacyl-HMDA-ppGpp), $^1H$ - NMR (400 MHz, $D_2O$)***

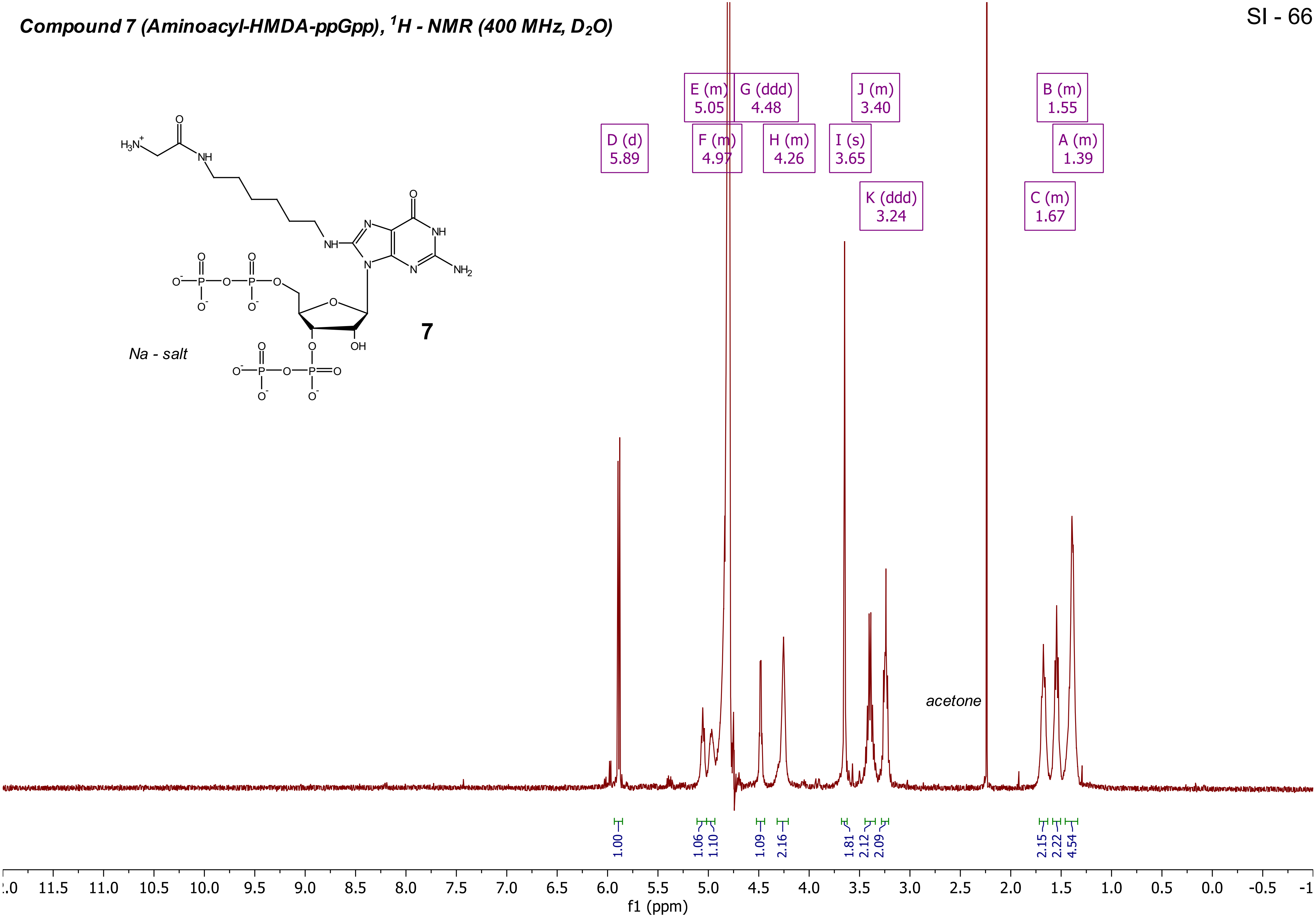


**7**

***Compound 7 (Aminoacyl-HMDA-ppGpp), $^{31}P\{^{1}H\}$ - NMR (162 MHz, $D_2O$)***

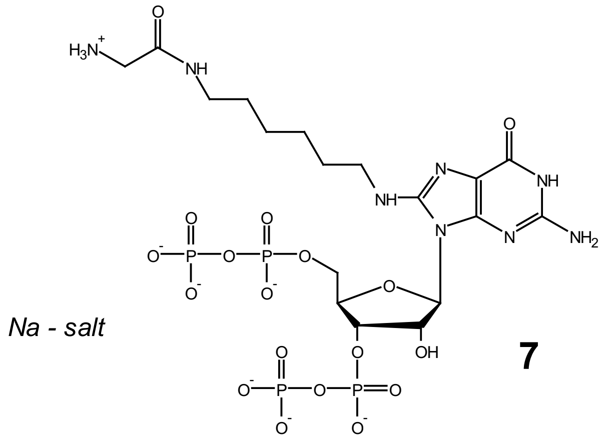


**7**

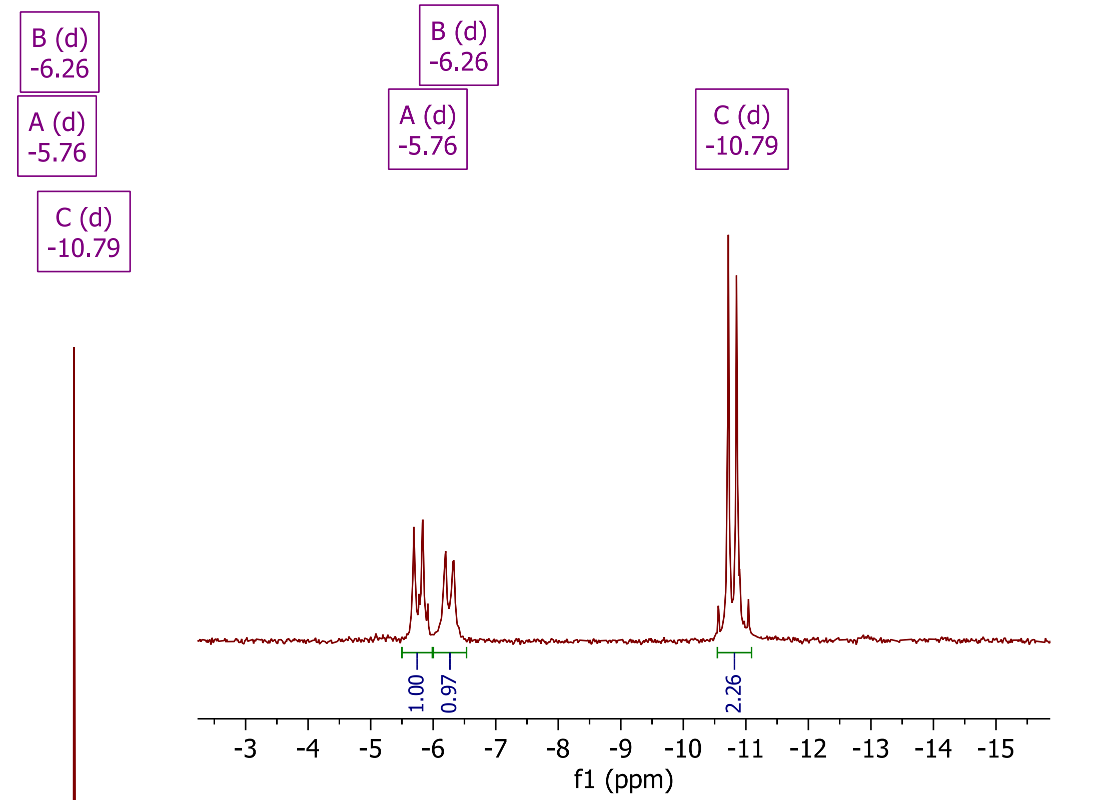


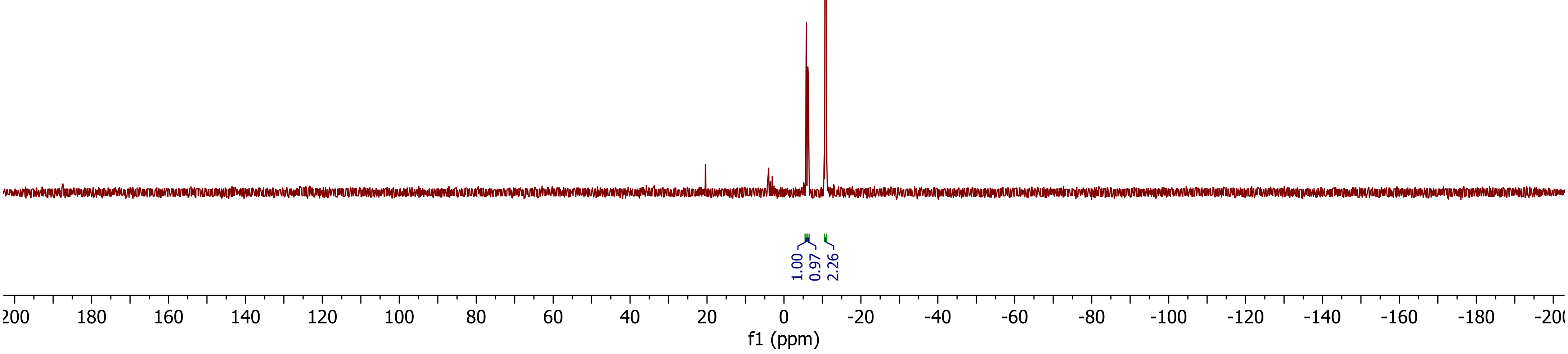

***Compound 7 (Aminoacyl-HMDA-ppGpp), DQF-COSY ($D_2O$)***

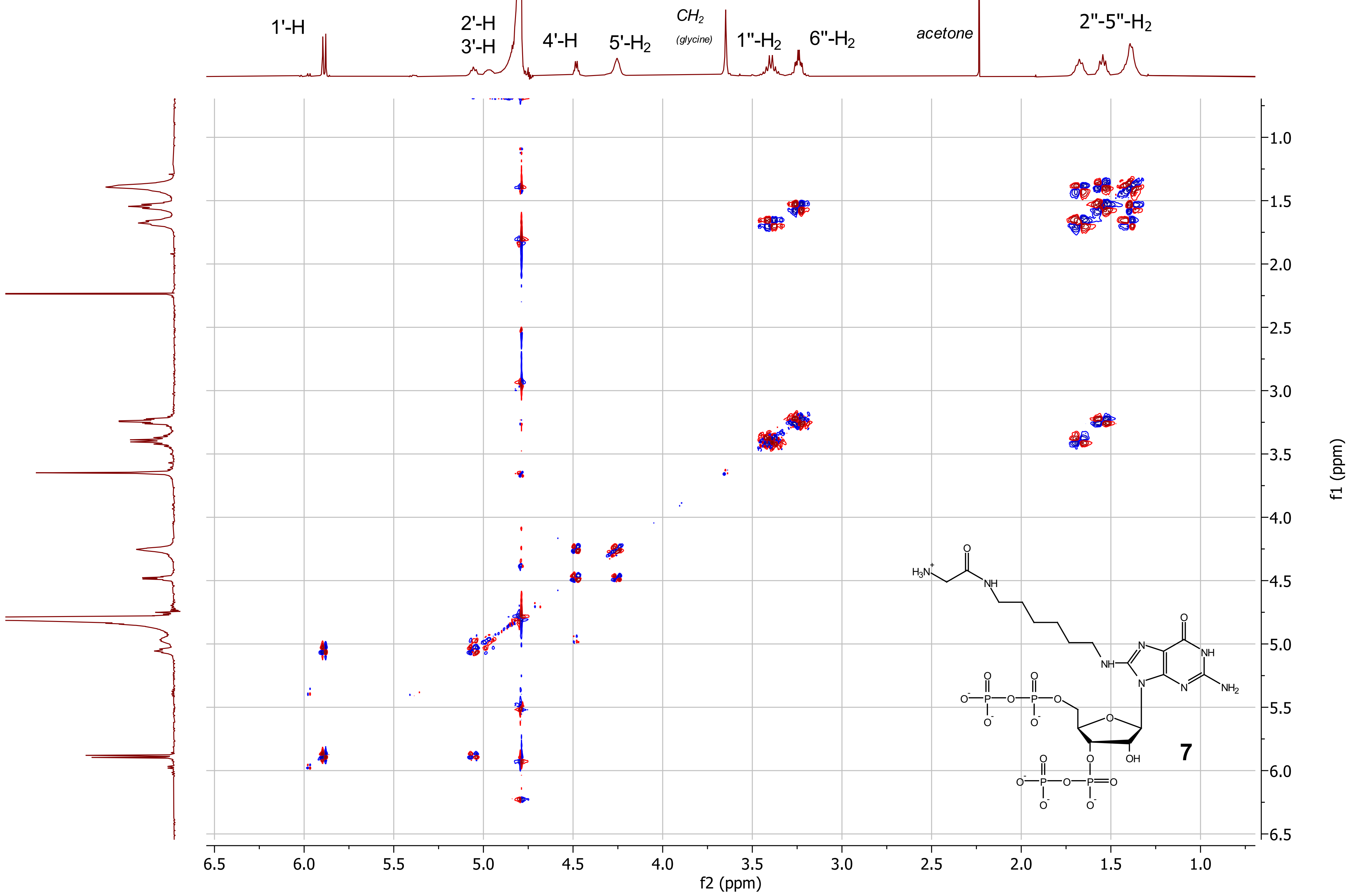

**Compound 7 (Aminoacyl-HMDA-ppGpp), HSQC ($D_2O$)**

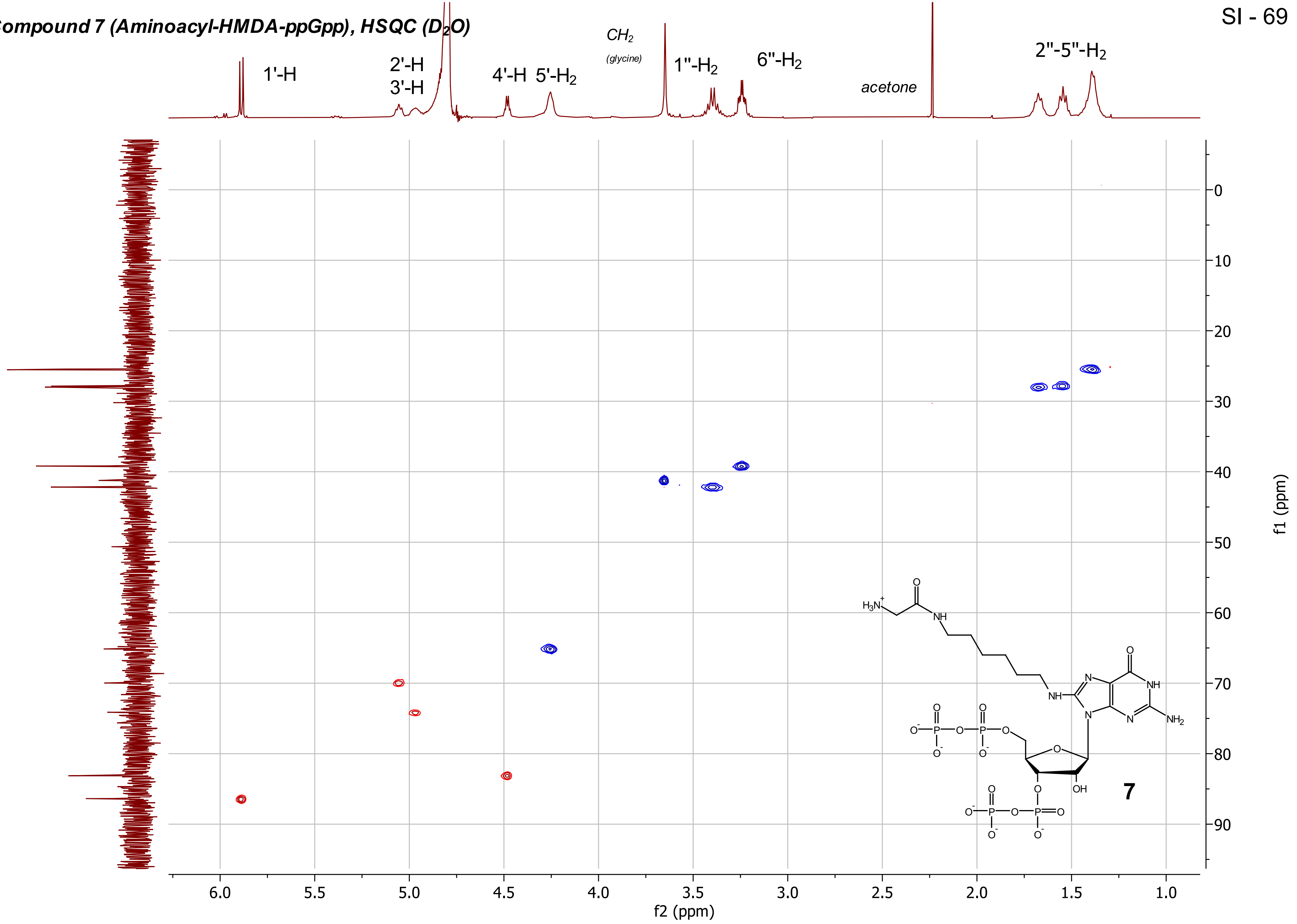

***Compound 7 (Aminoacyl-HMDA-ppGpp), ¹H-³¹P-HMBC***

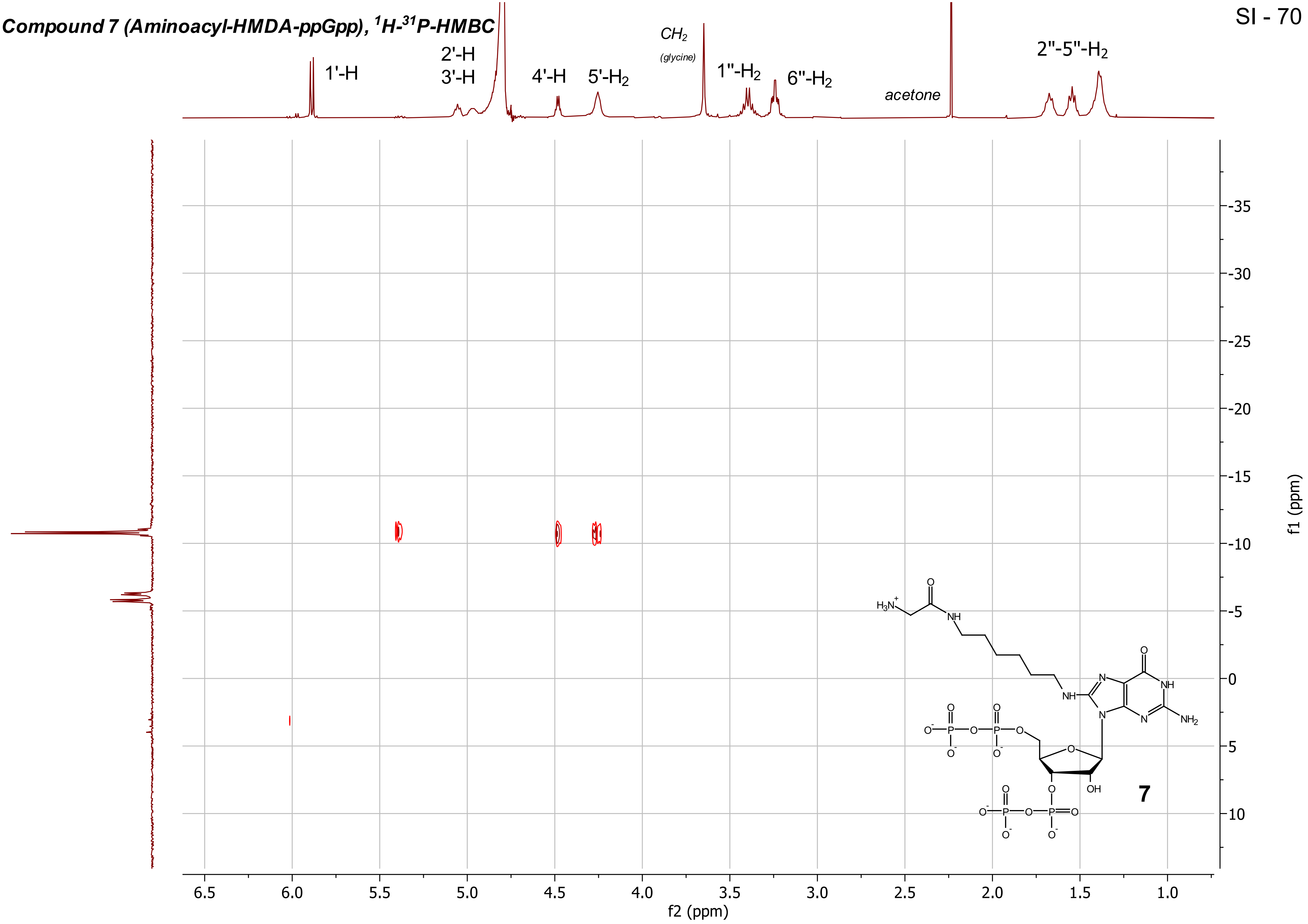

***Compound 7 (Aminoacyl-HMDA-ppGpp), PP - COSY***

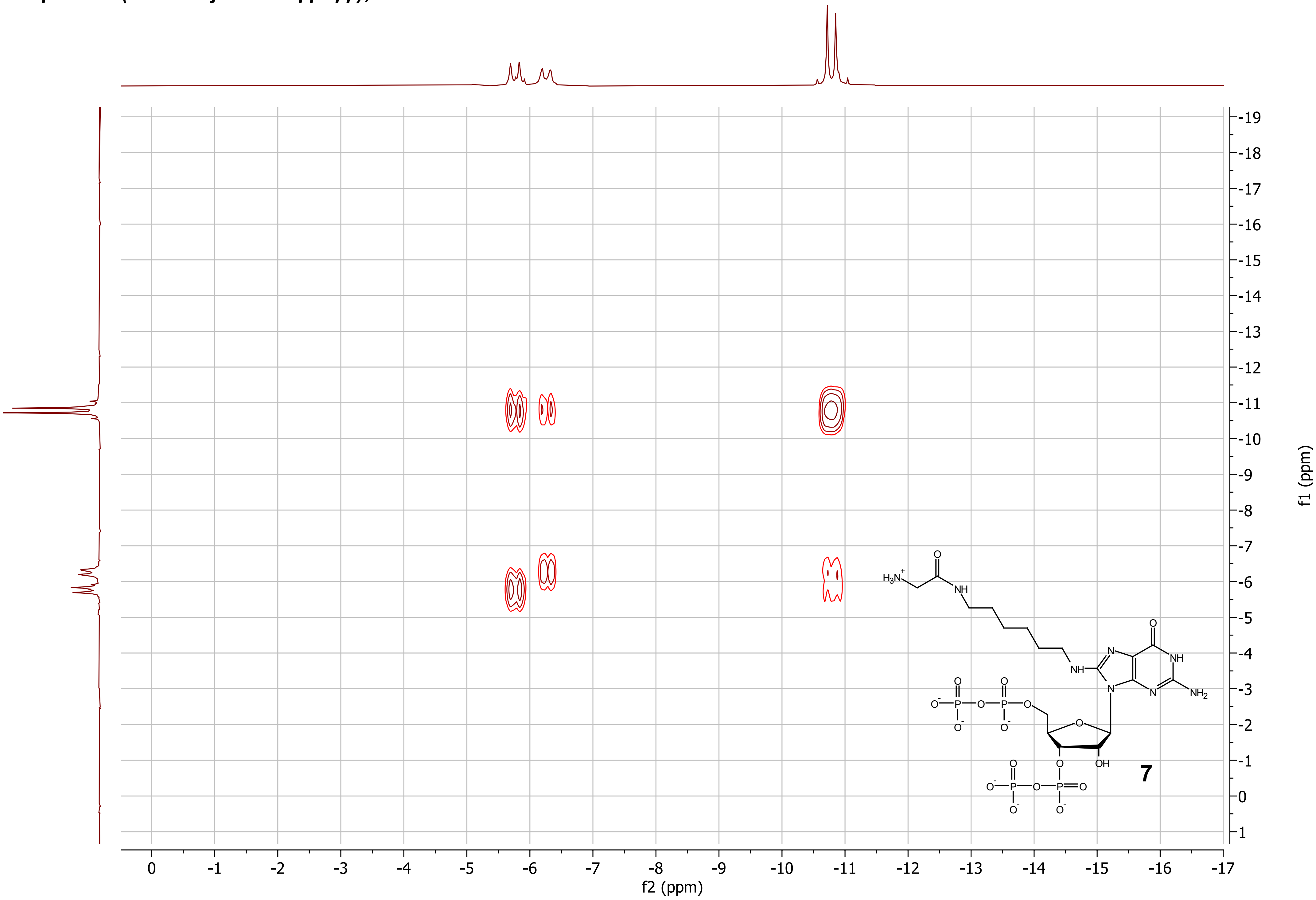

***Compound 7 (Aminoacyl-HMDA-ppGpp), $^{13}C\{^{1}H\}$ - NMR ($D_2O$, 101 MHz)***

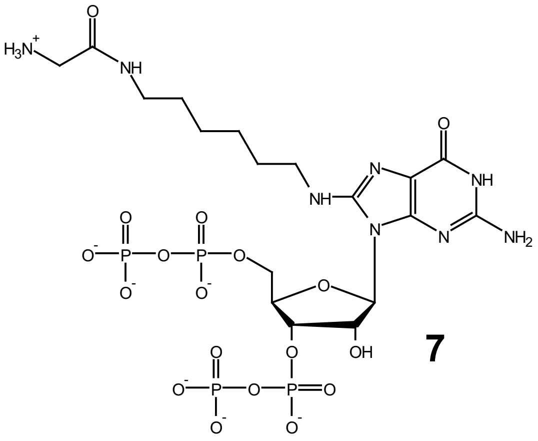

**7**

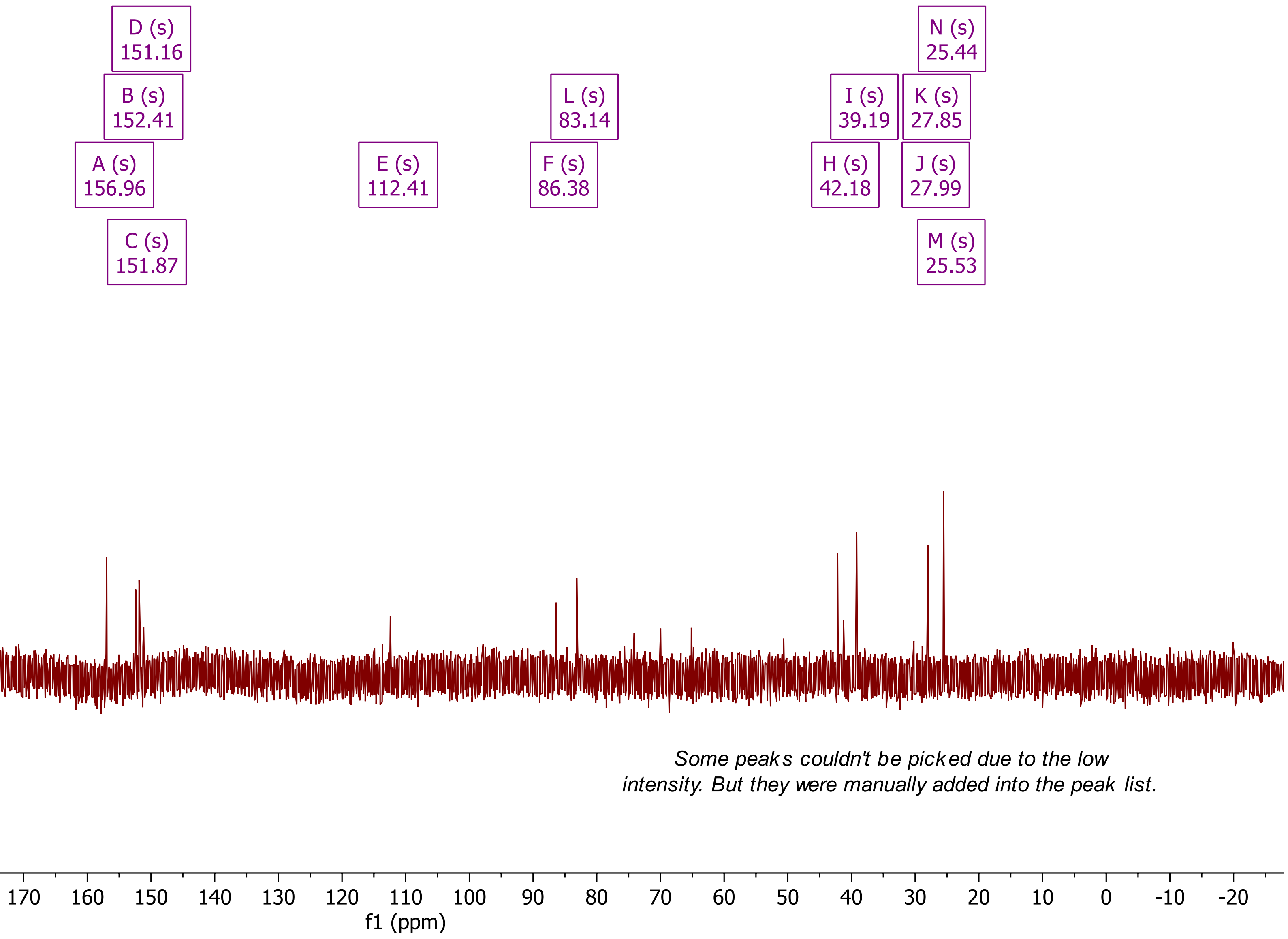


*Some peaks couldn't be picked due to the low intensity. But they were manually added into the peak list.*

*Compound 9 (pentynyl-pppGp), [1]H - NMR ($D_2O$, 400 MHz)*

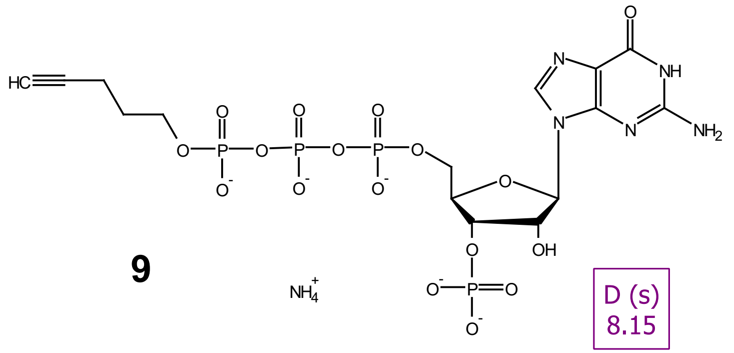

**9**

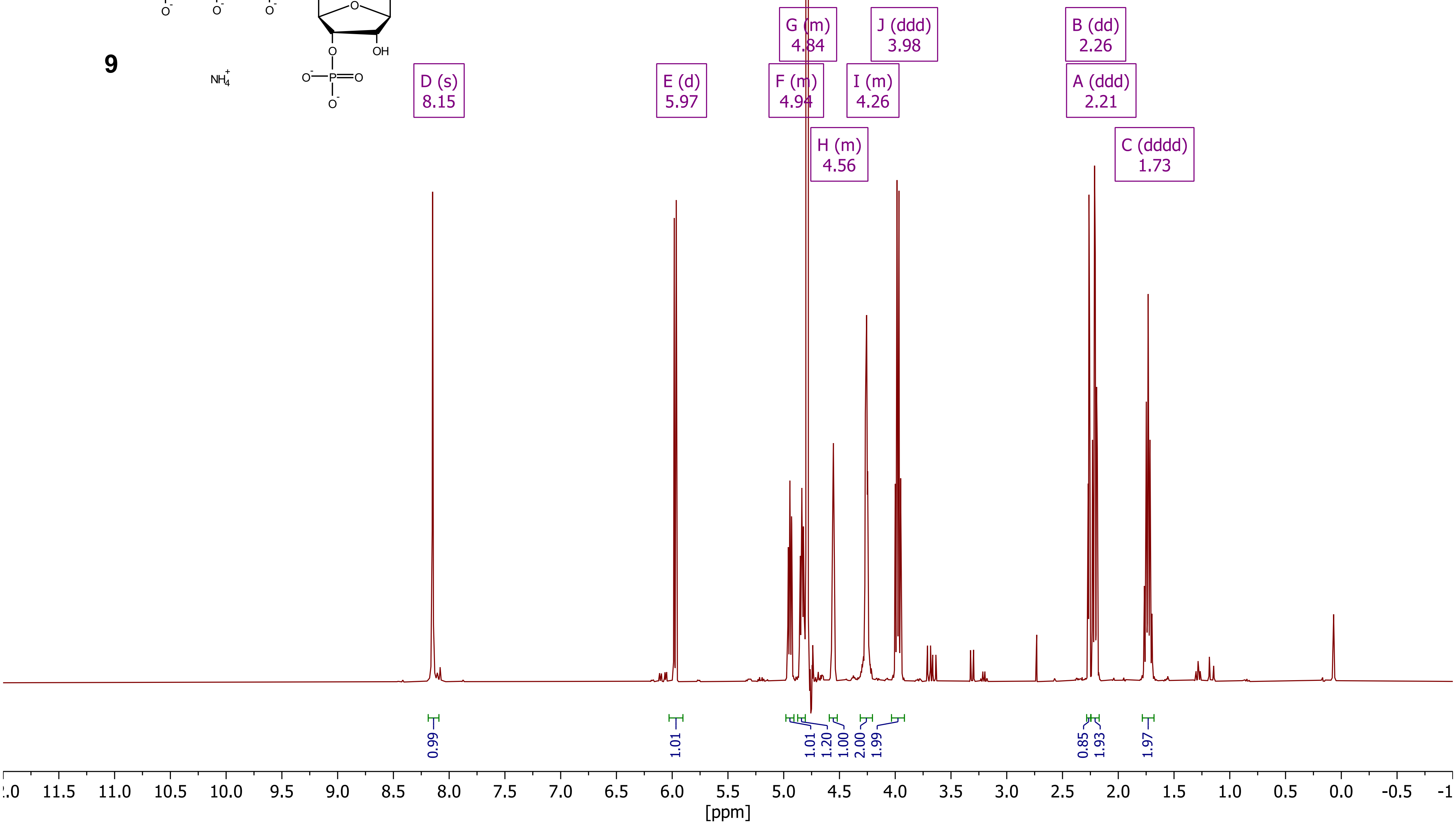

*Compound 9 (pentynyl-pppGp), $^{31}P\{^1H\}$ - NMR ($D_2O$, 162 MHz)*



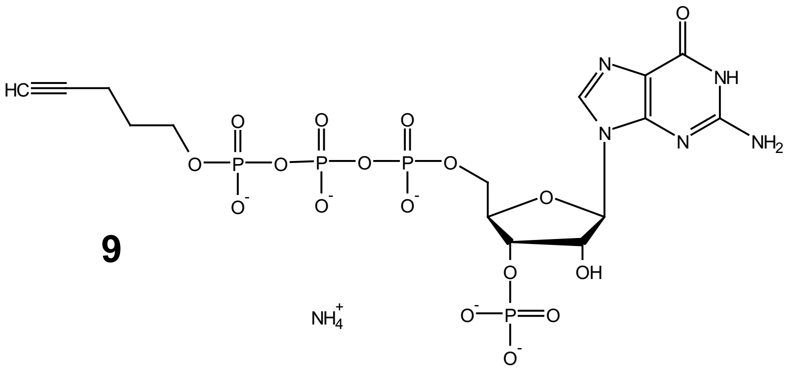

**9**

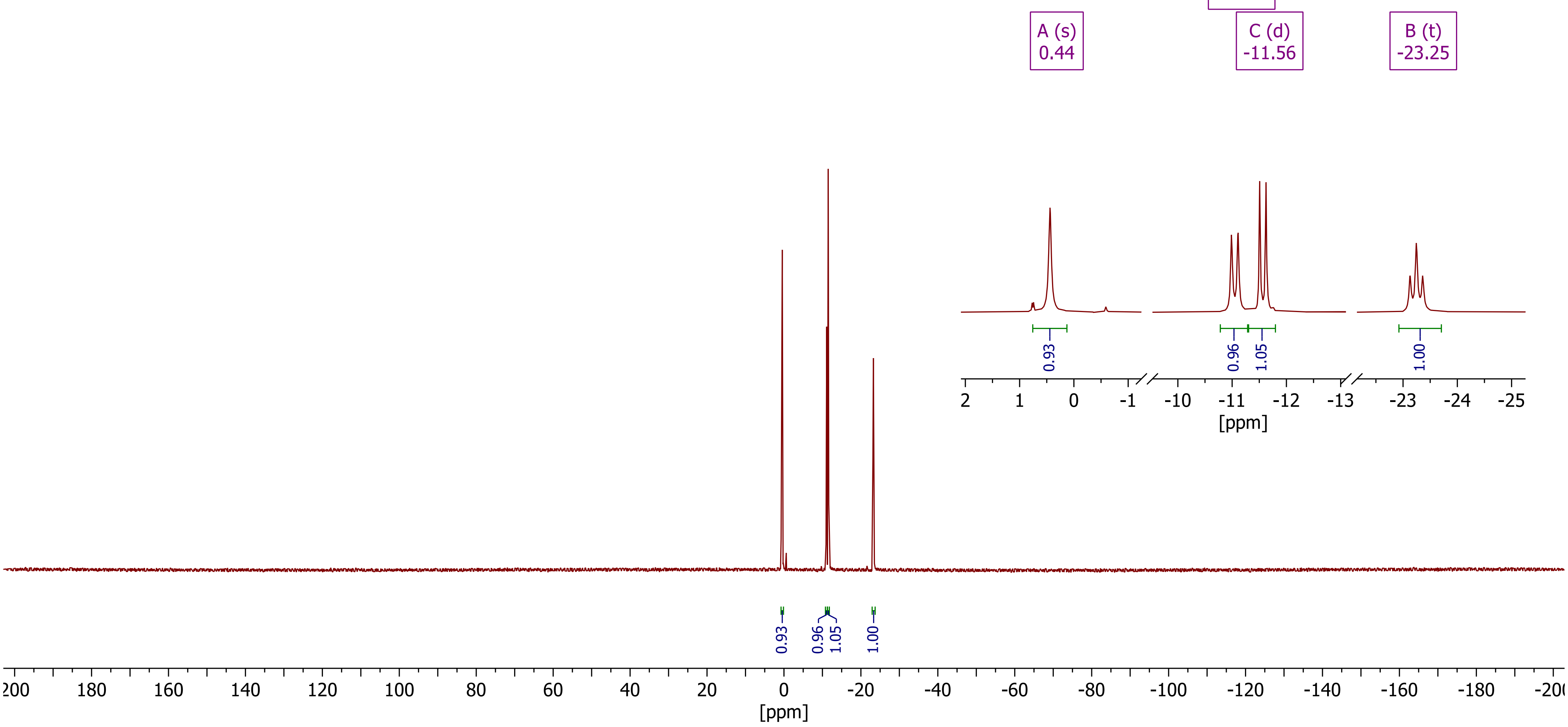

*Compound 9 (pentynyl-pppGp), DQF-COSY*



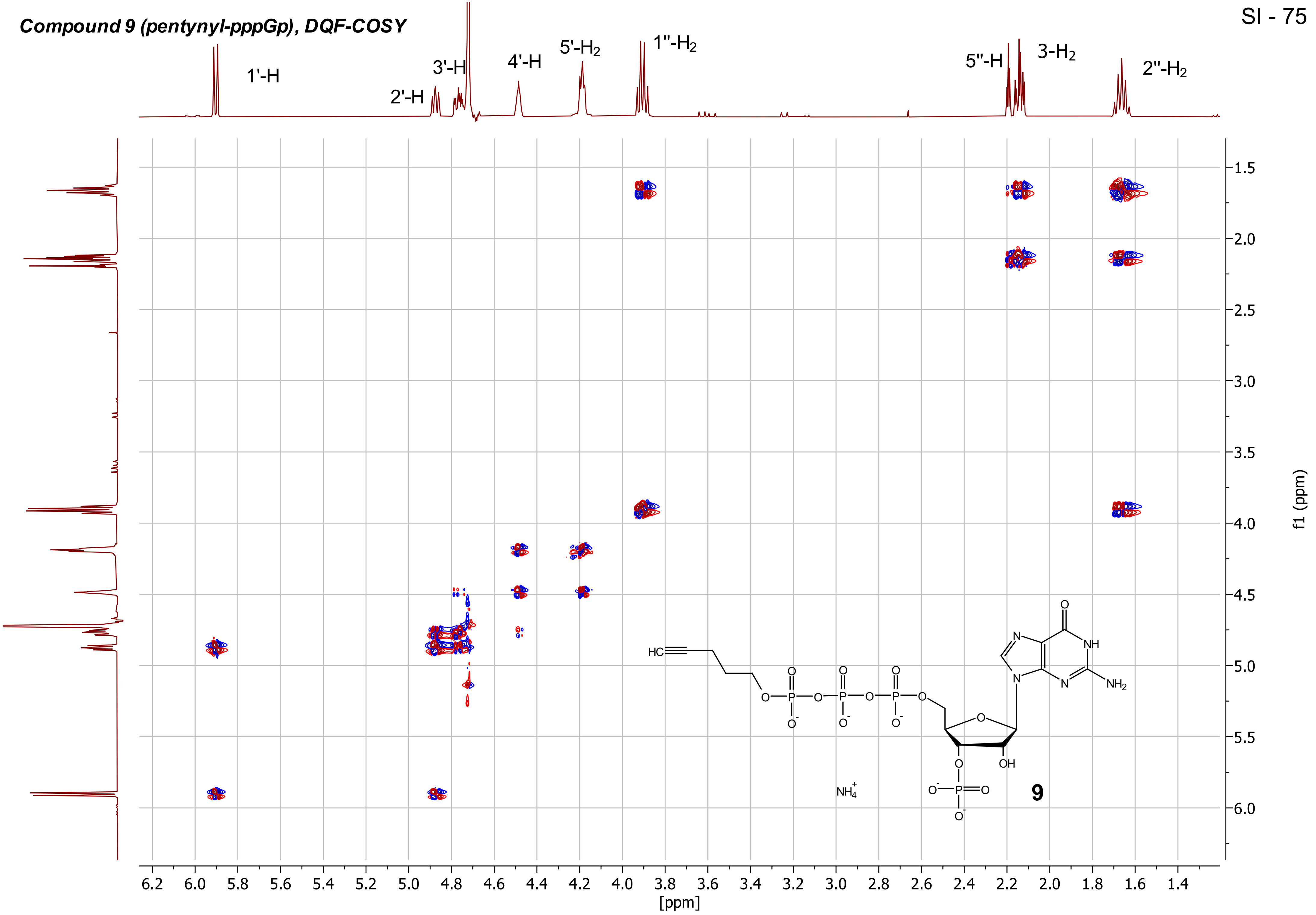

**_Compound 9 (pentynyl-pppGp), HSQC_**



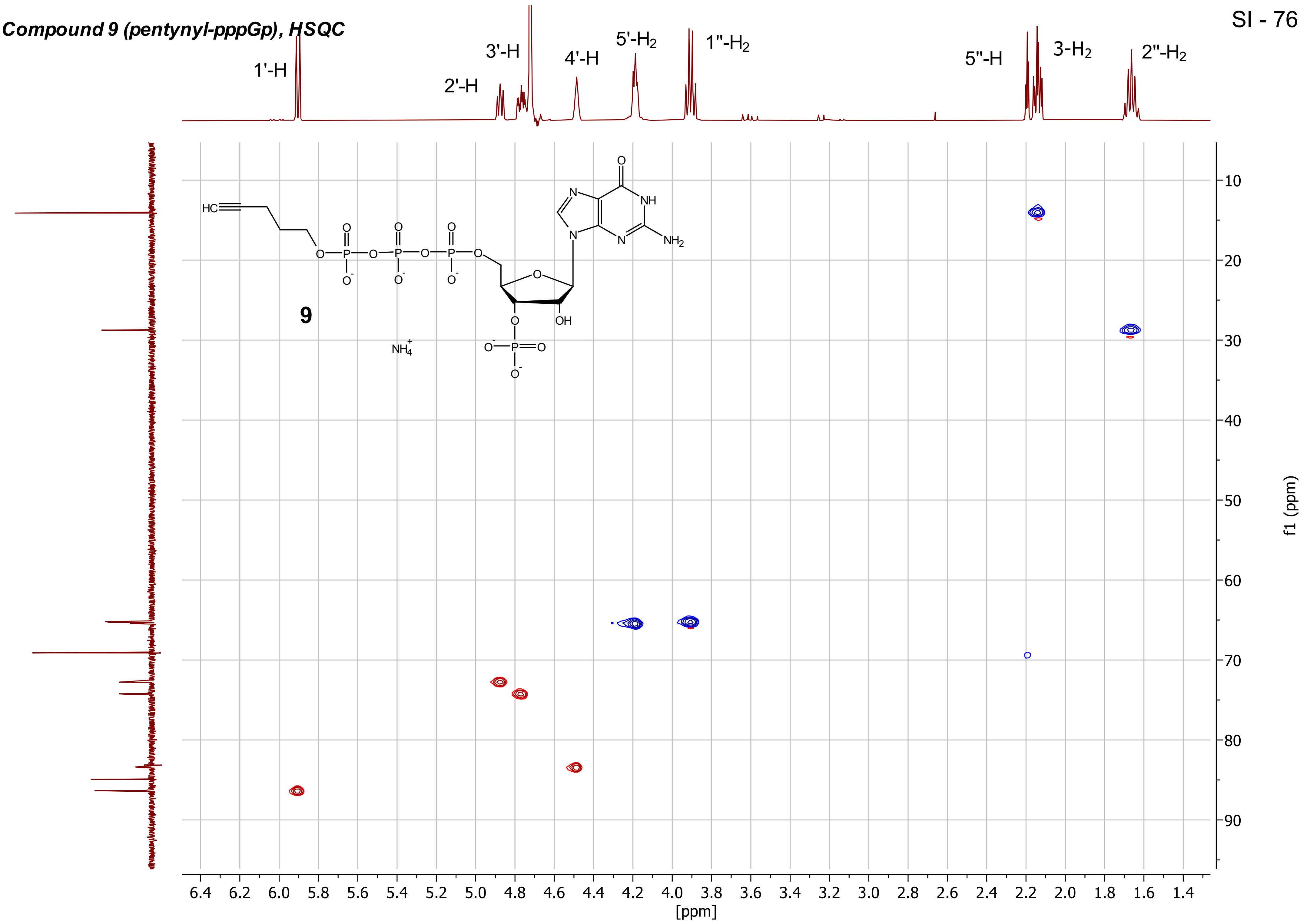

***Compound 9 (pentynyl-pppGp), 1H-31P-HMBC***

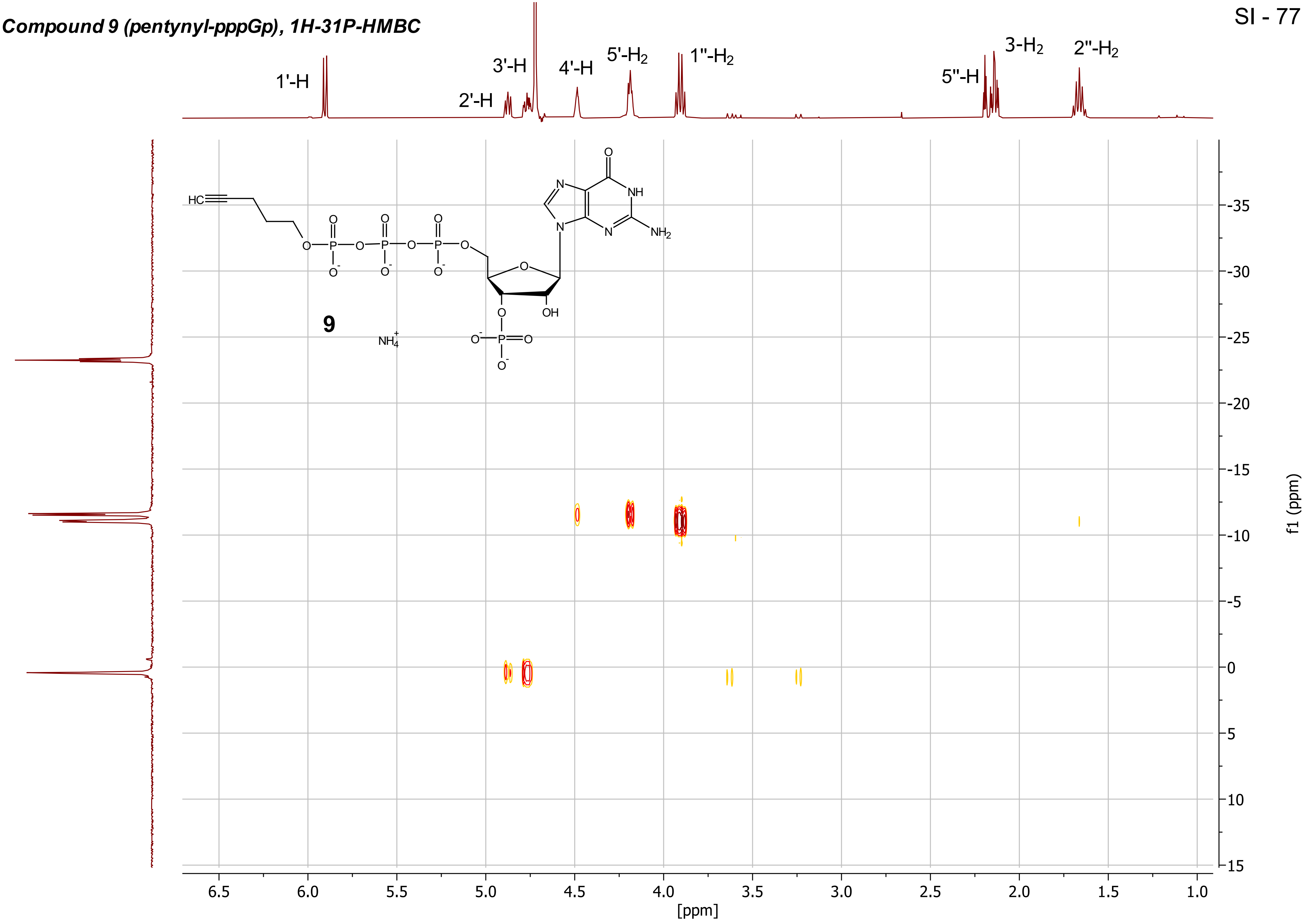

***Compound 9 (pentynyl-pppGp), $^{13}C\{^1H\}$ - NMR ($D_2O$, 101 MHz)***

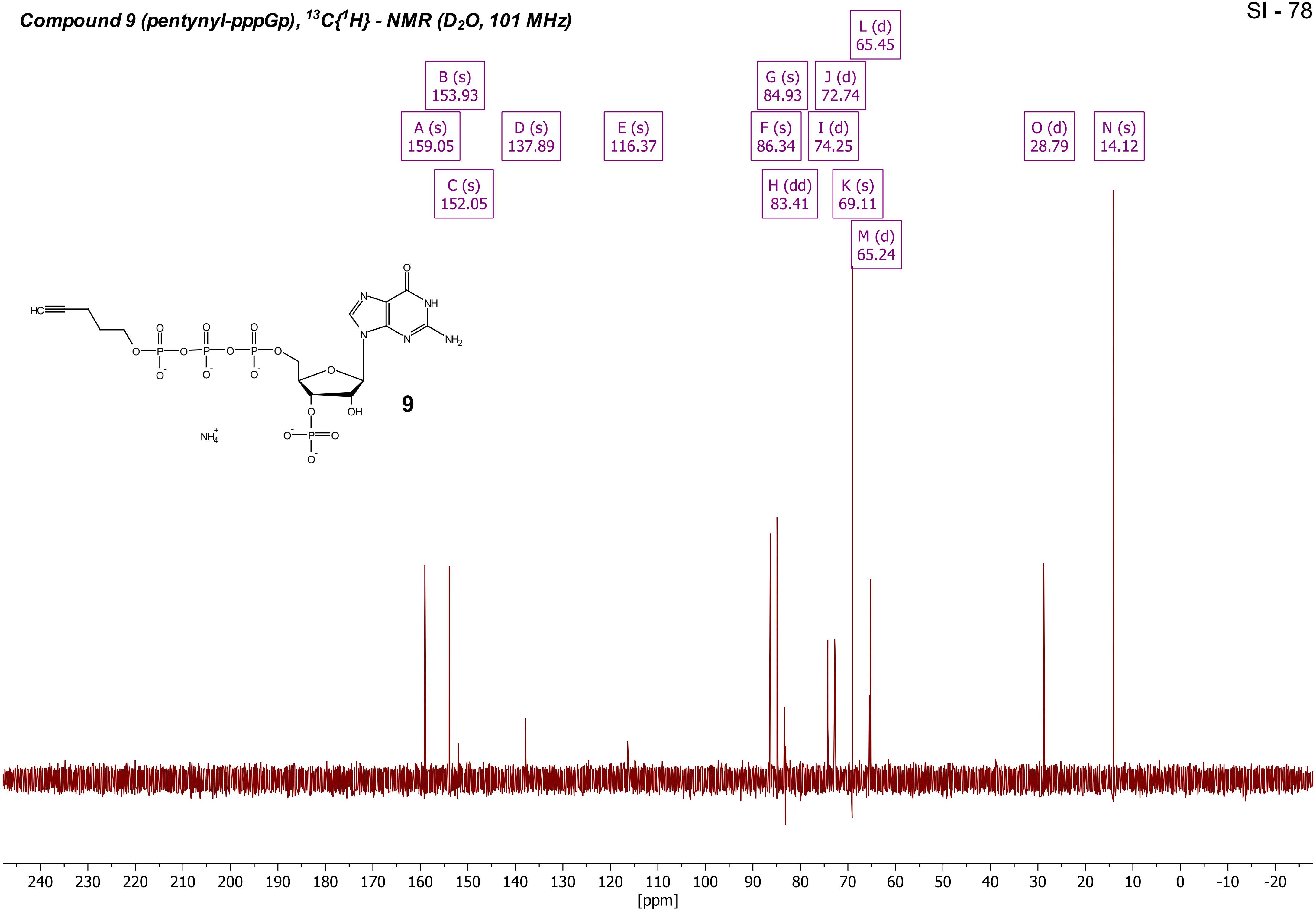

***Compound 11 (pentynyl-ppAp), $^{1}H$ - NMR ($D_2O$, 400 MHz)***

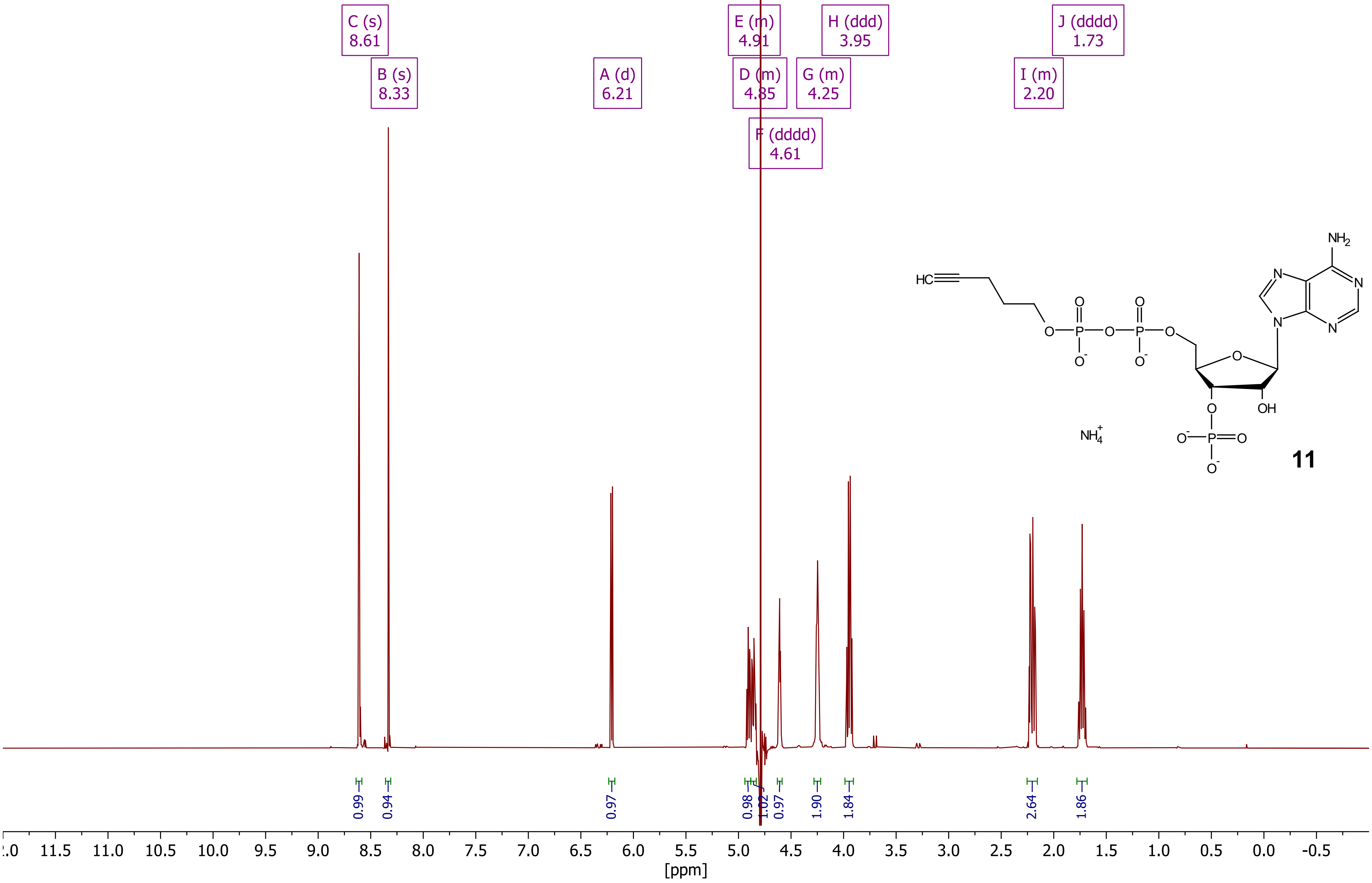

***Compound 11 (pentynyl-ppAp), $^{31}P\{^1H\}$ - NMR ($D_2O$, 162 MHz)***

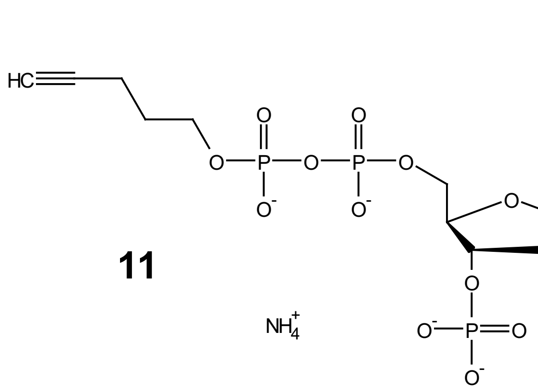

**11**

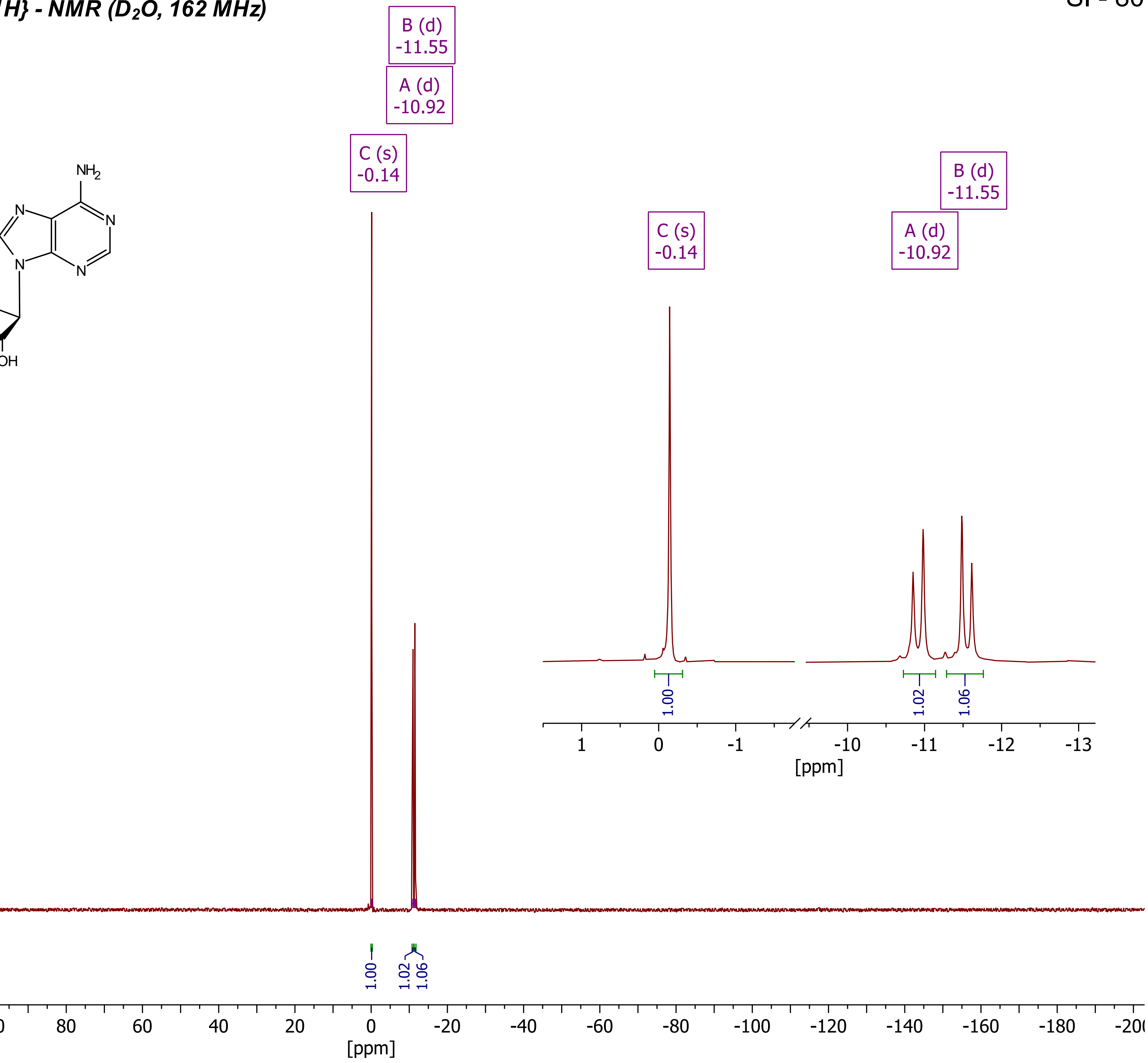

***Compound 11 (pentynyl-ppAp), DQF-COSY ($D_2O$)***



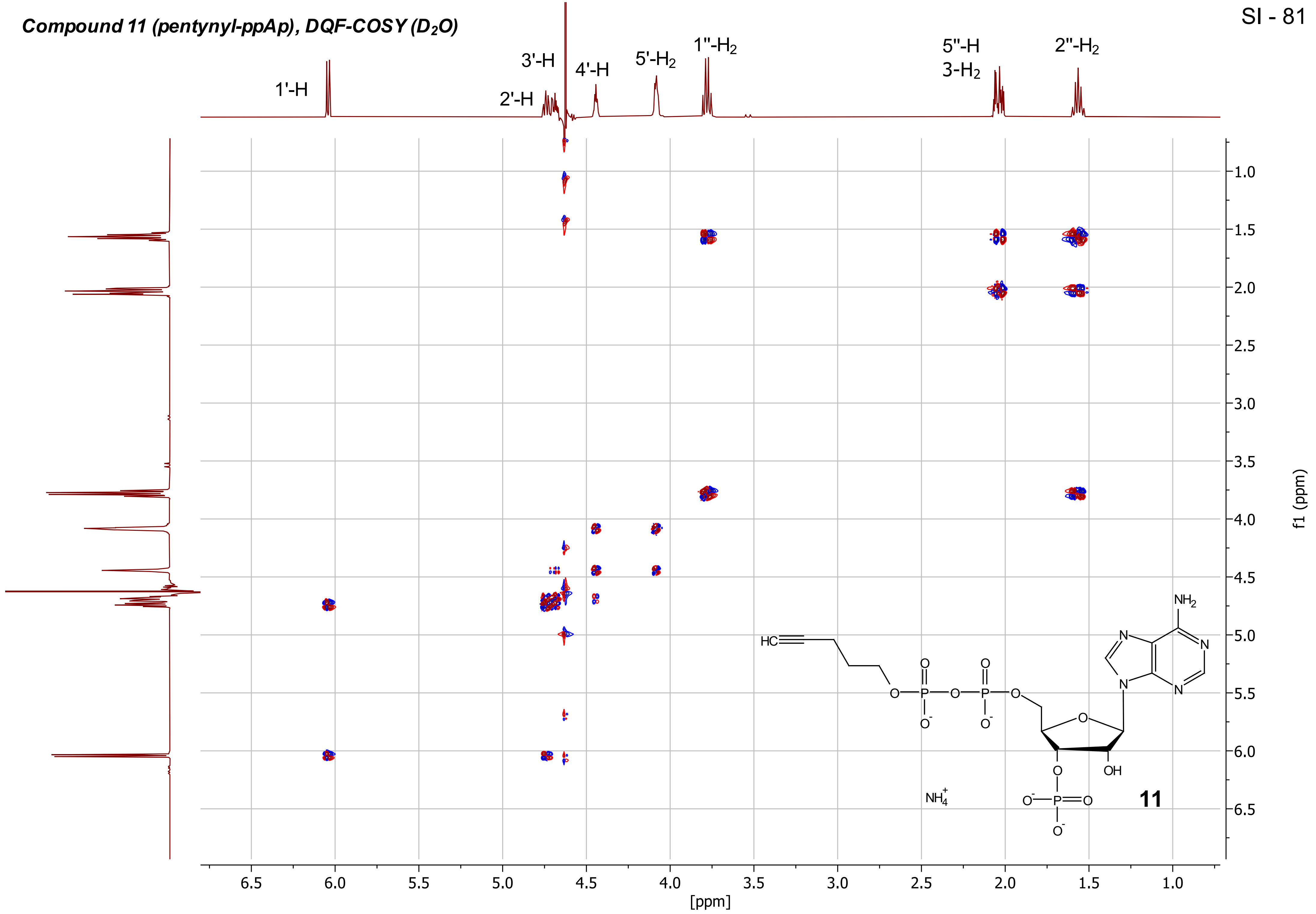

***Compound 11 (pentynyl-ppAp), HSQC ($D_2O$)***

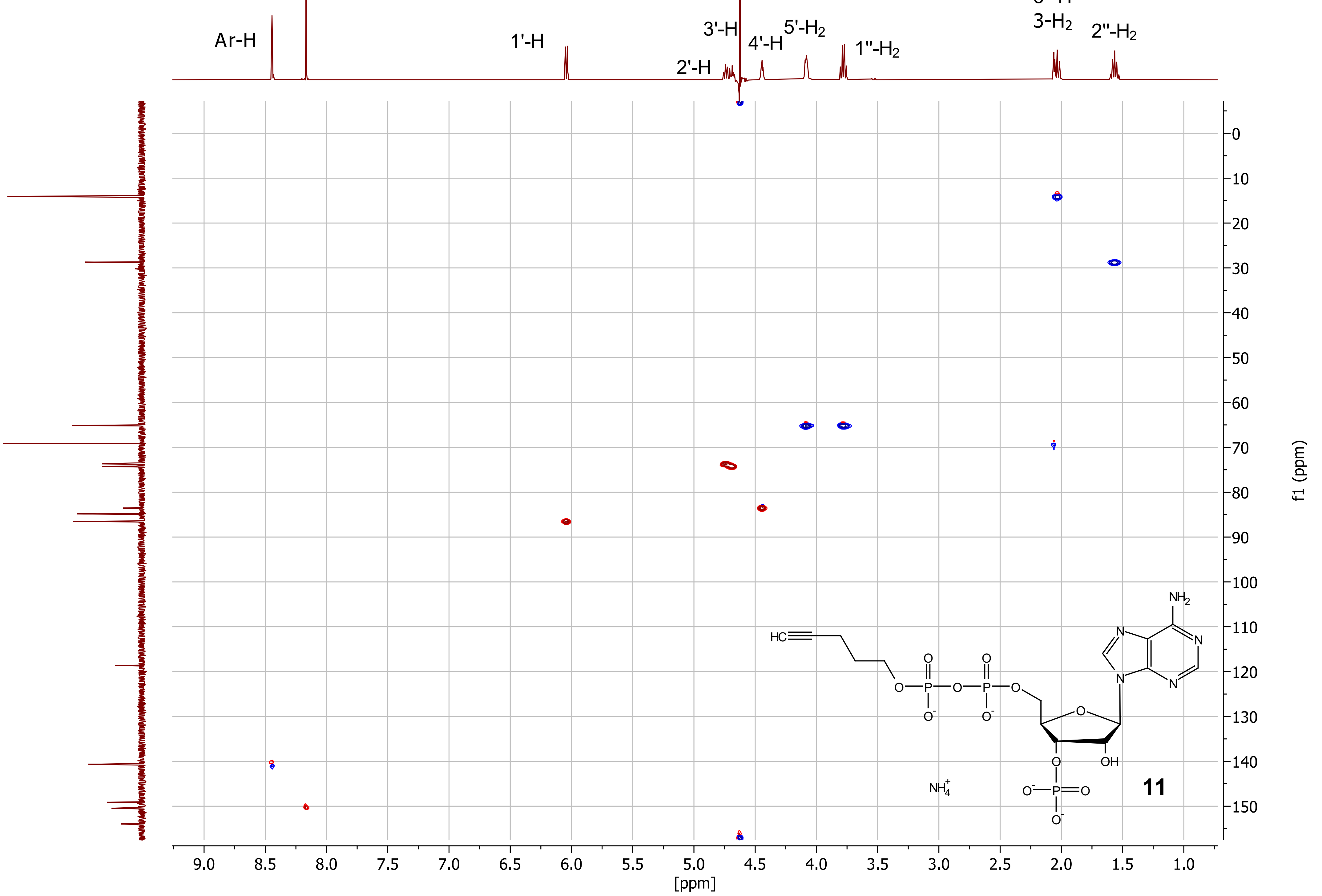

***Compound 11 (pentynyl-ppAp), $^{1}H$-$^{31}P$-HMBC ($D_2O$)***

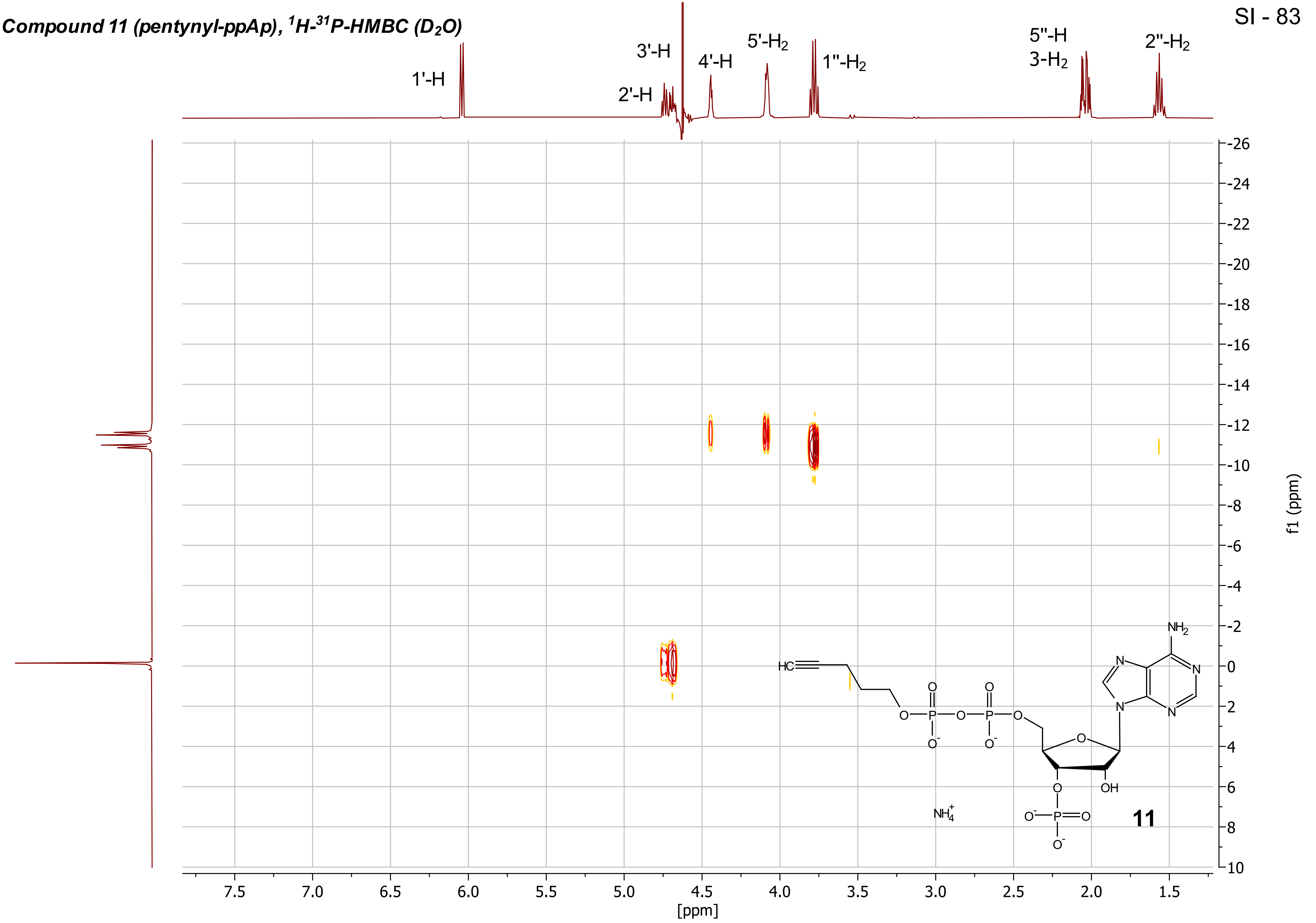

***Compound 11 (pentynyl-ppAp), $^{13}C\{^{1}H\}$ - NMR ($D_2O$, 101 MHz)***

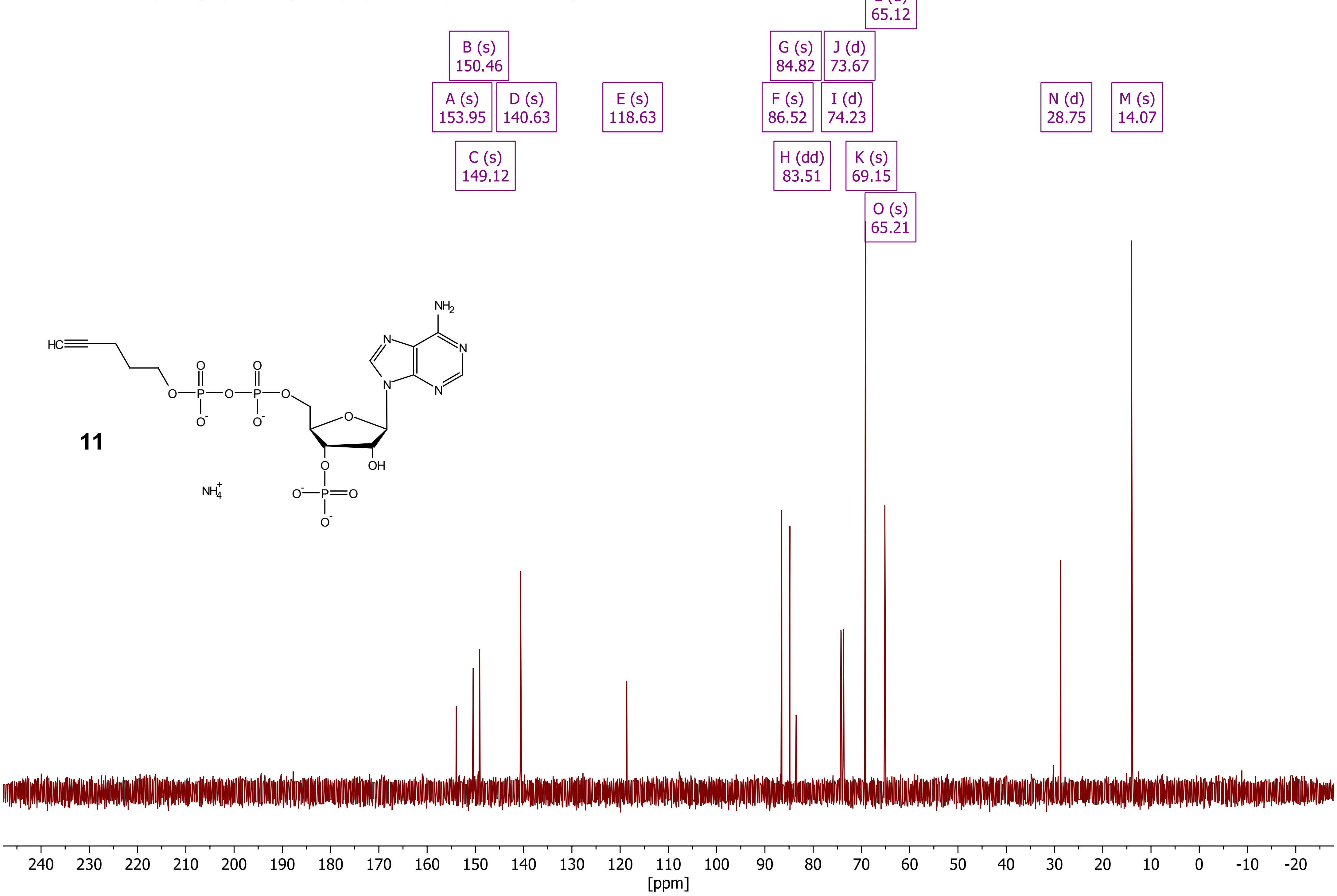

***Compound 12 (pentynyl-pGp), $^{1}H$ - NMR ($D_2O$, 400 MHz)***

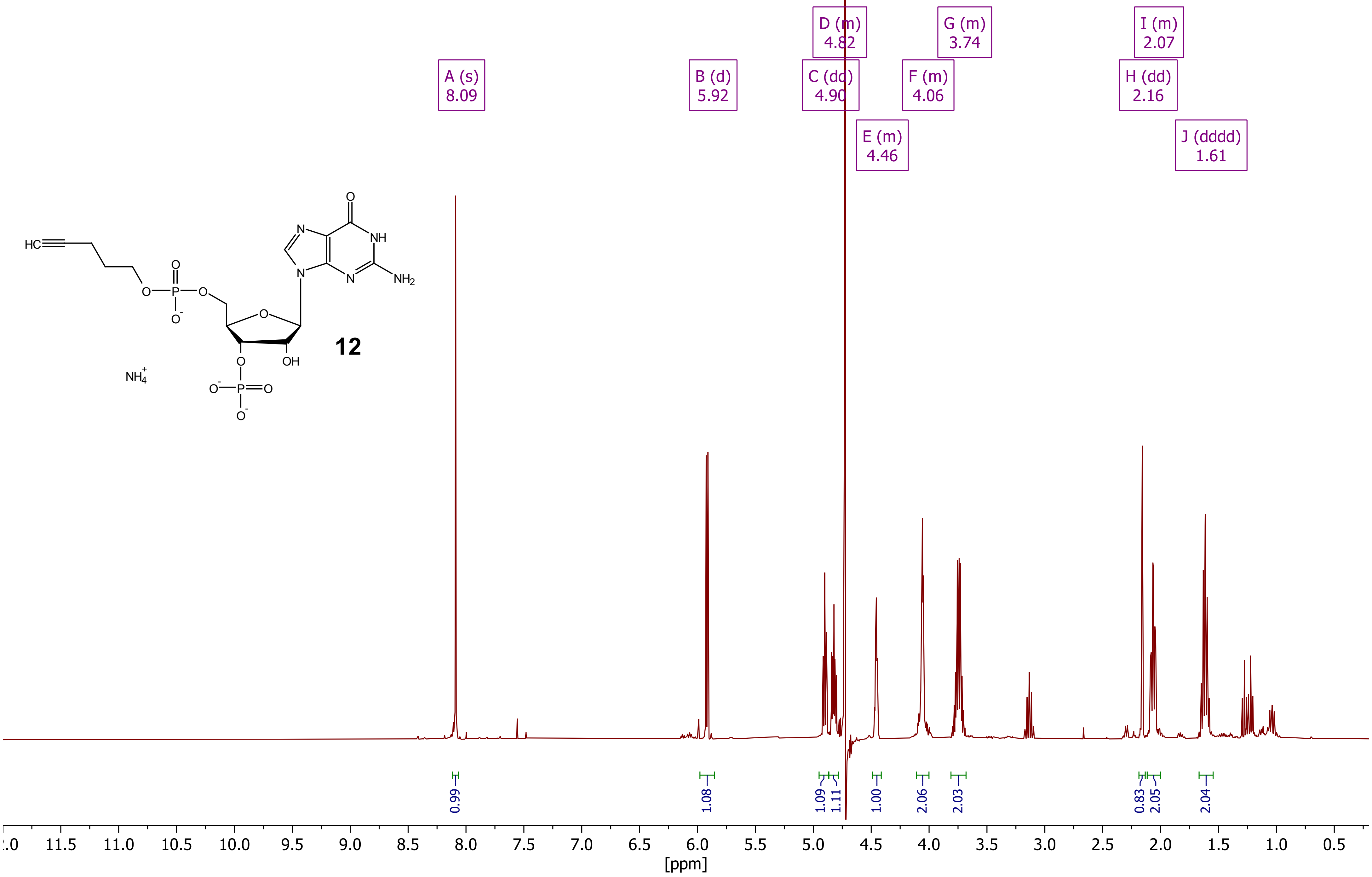

***Compound 12 (pentynyl-pGp), $^{31}P\{^1H\}$ - NMR ($D_2O$, 162 MHz)***

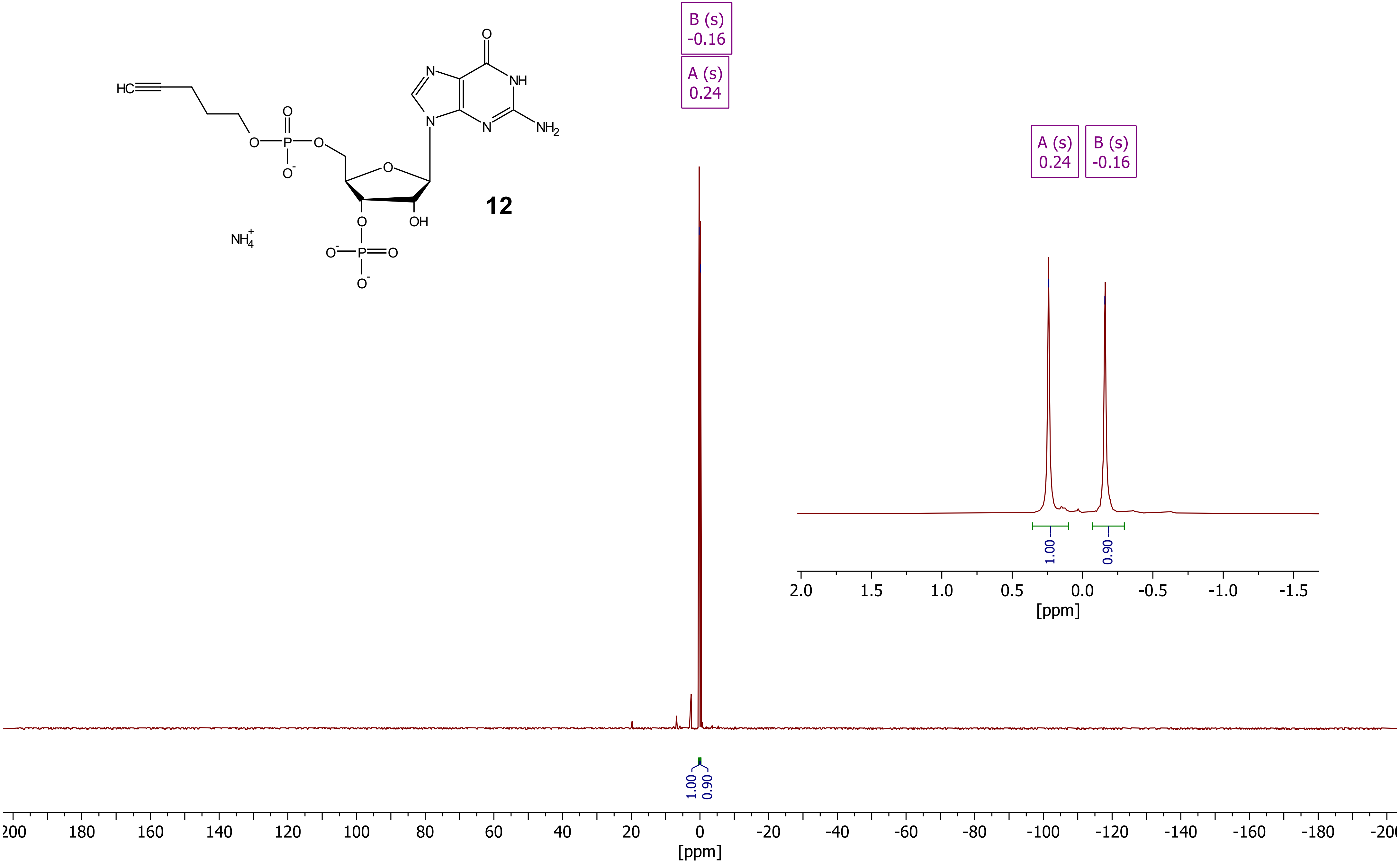

***Compound 12 (pentynyl-pGp), DQF-COSY ($D_2O$)***

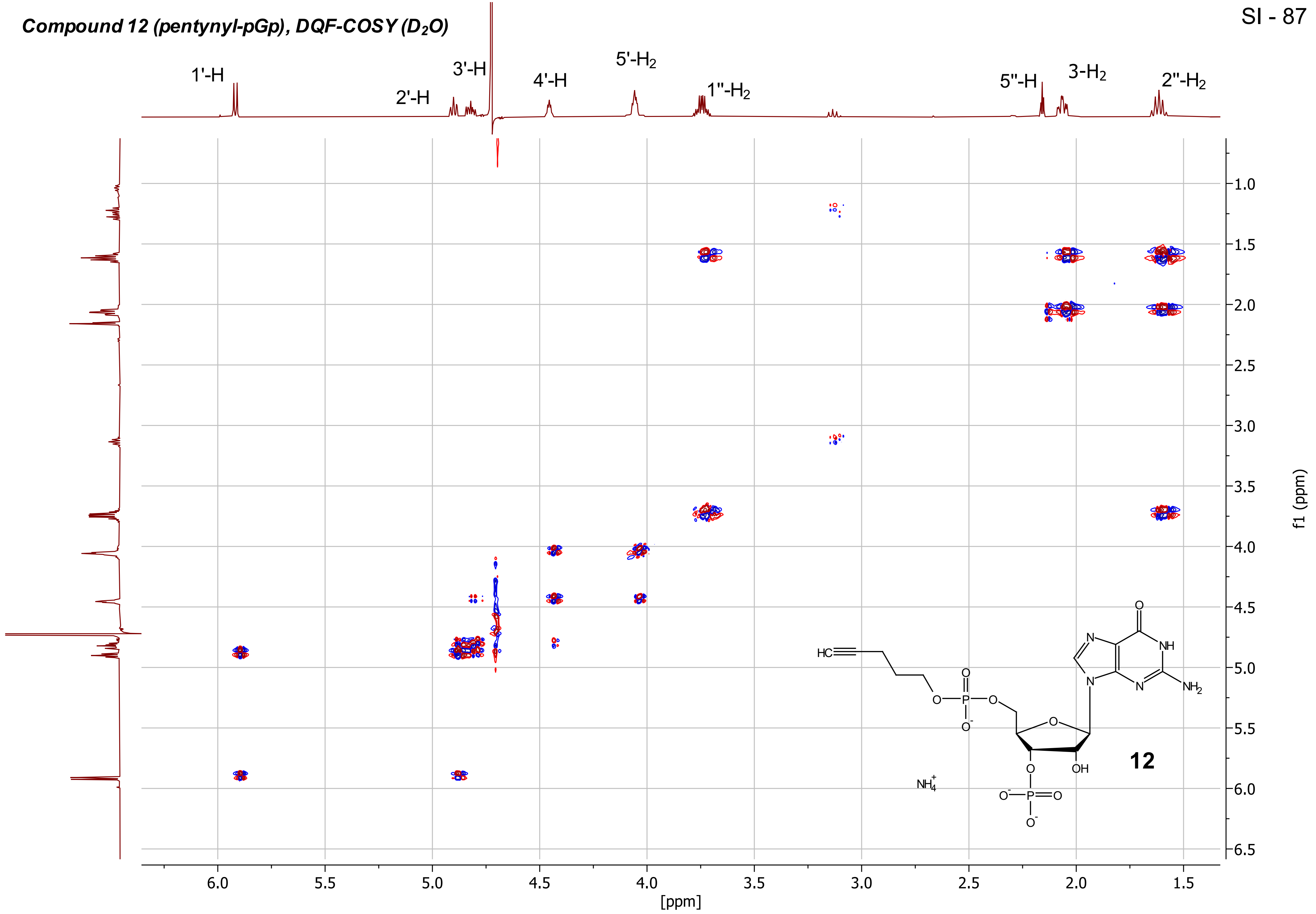

***Compound 12 (pentynyl-pGp), HSQC ($D_2O$)***



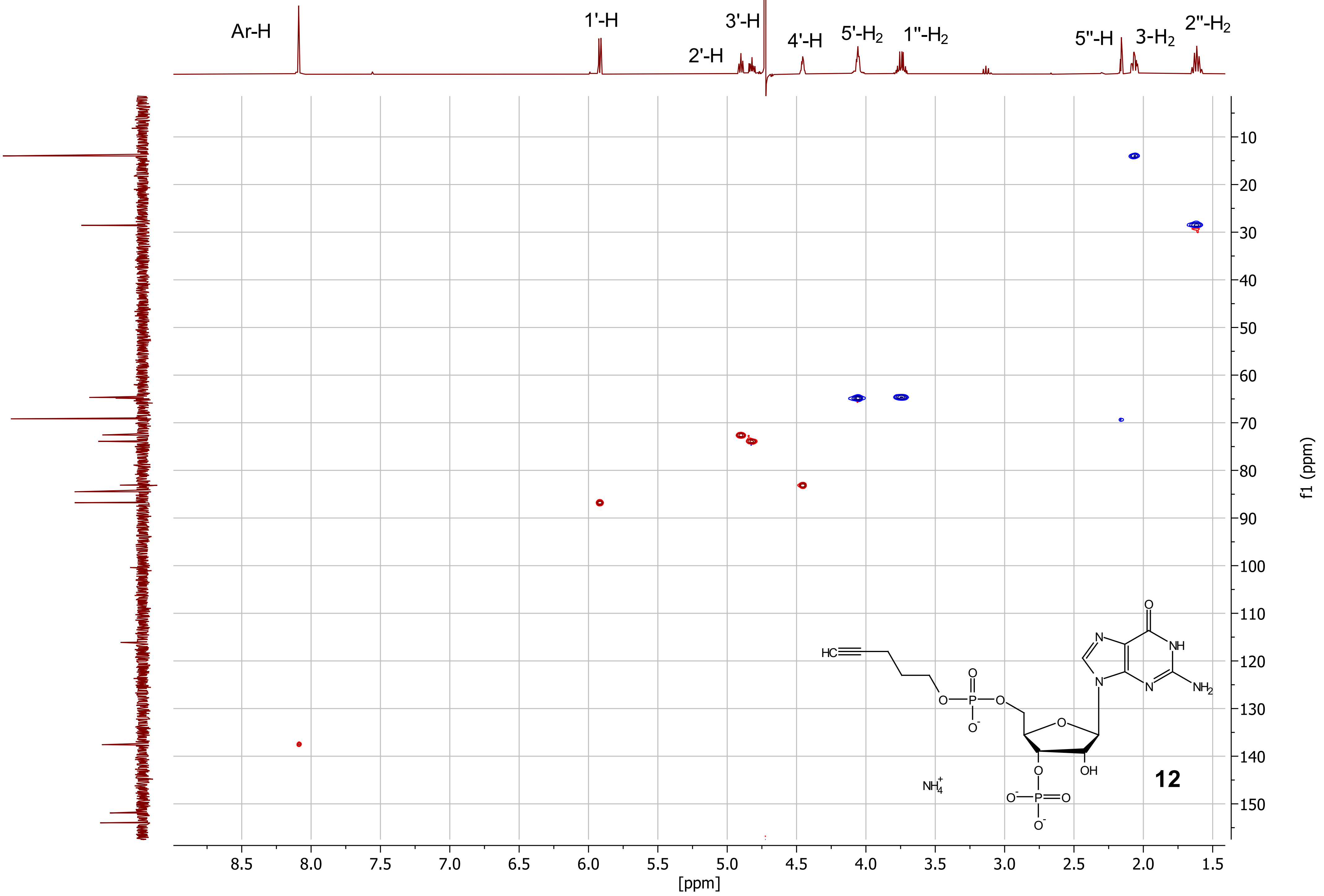


**12**

***Compound 12 (pentynyl-pGp), $^{1}H$-$^{31}P$-HMBC ($D_2O$)***

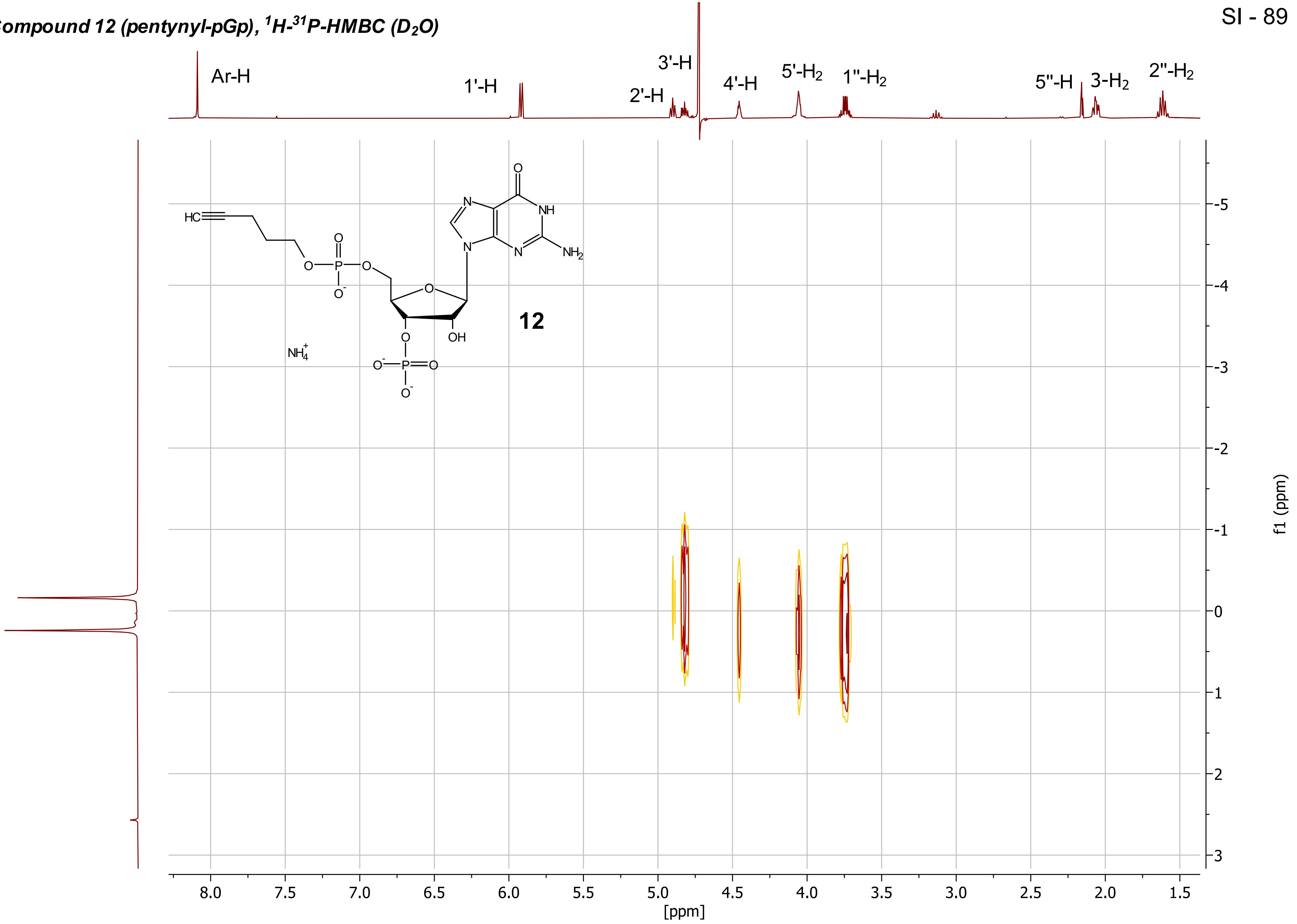

***Compound 12 (pentynyl-pGp), $^{13}C\{^{1}H\}$ - NMR (D2O, 101 MHz)***

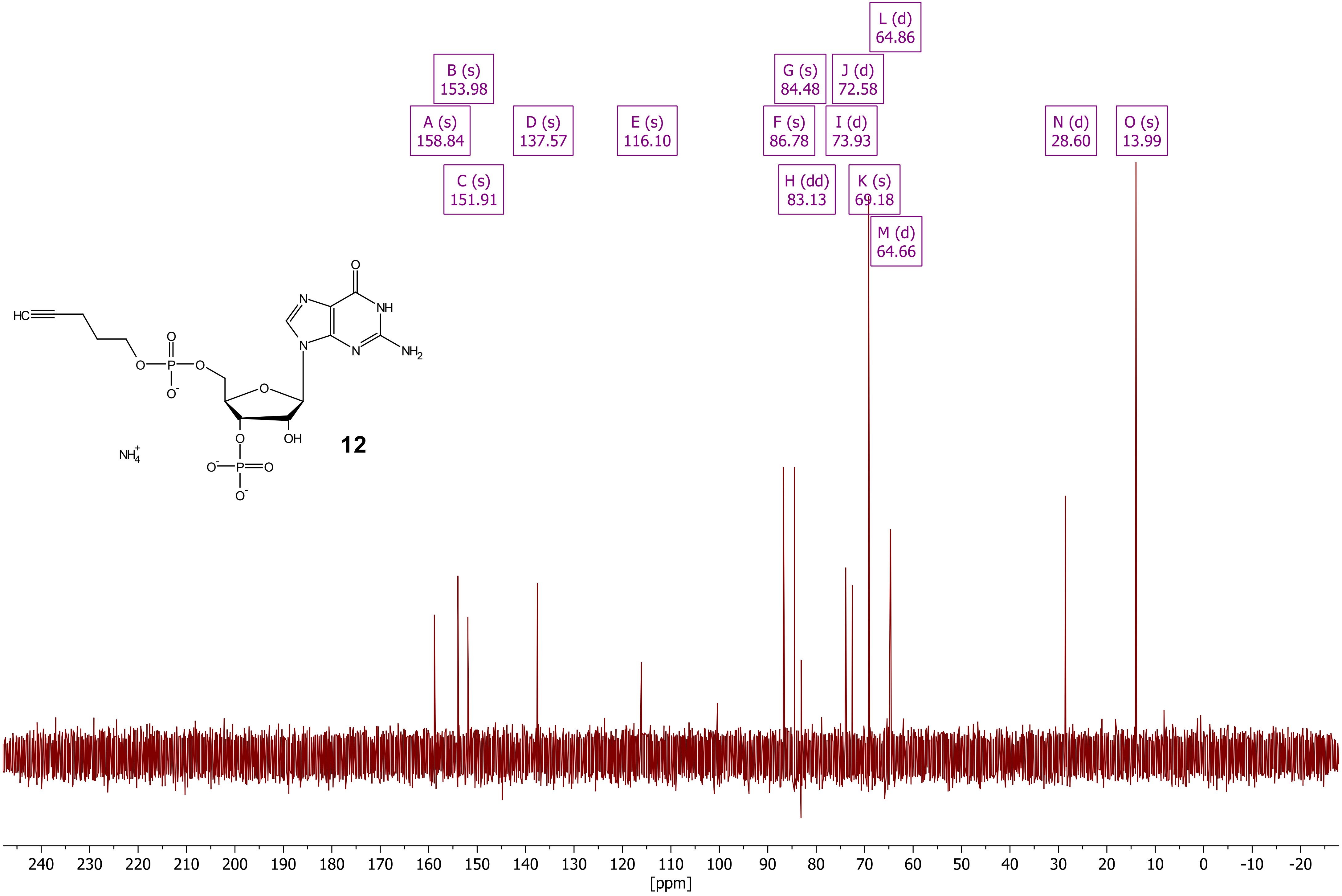

***Compound 13 (pentynyl-pGpp), $^{1}H$ - NMR ($D_2O$, 400 MHz)***

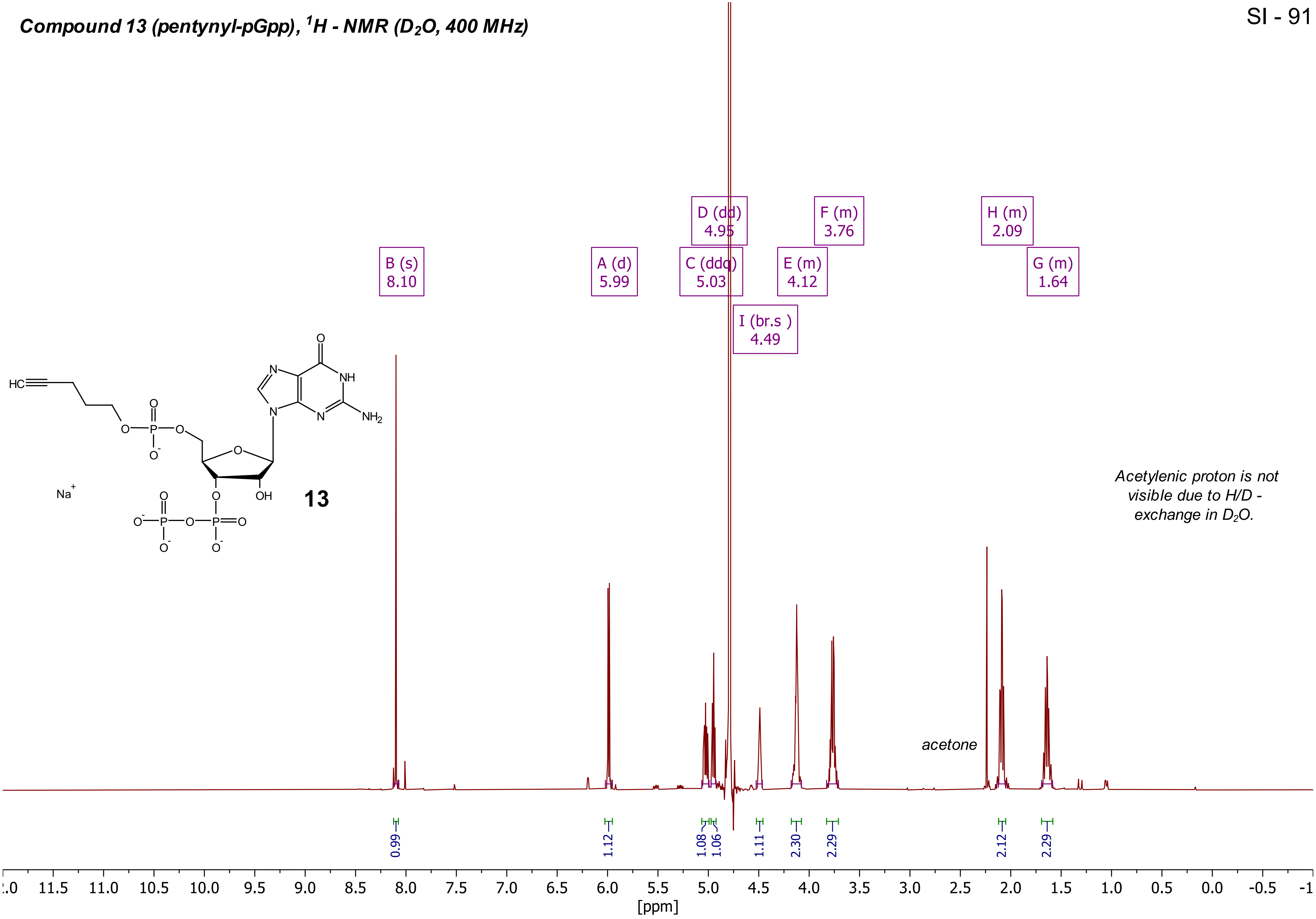

***Compound 13 (pentynyl-pGpp), $^{31}P\{^{1}H\}$ - NMR ($D_2O$, 162 MHz)***

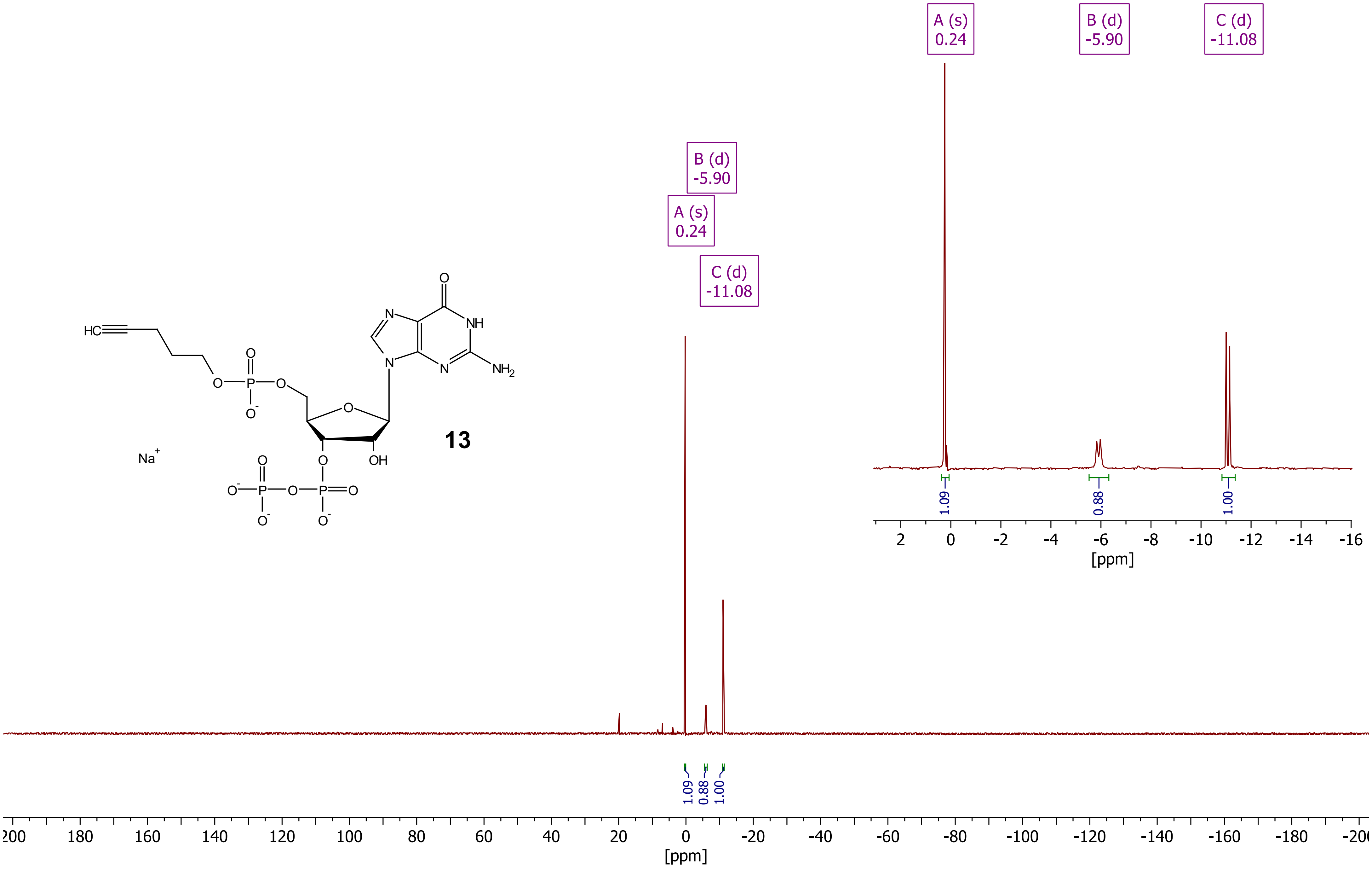

#### *Compound 13 (pentynyl-pGpp), DQF-COSY ($D_2O$)*



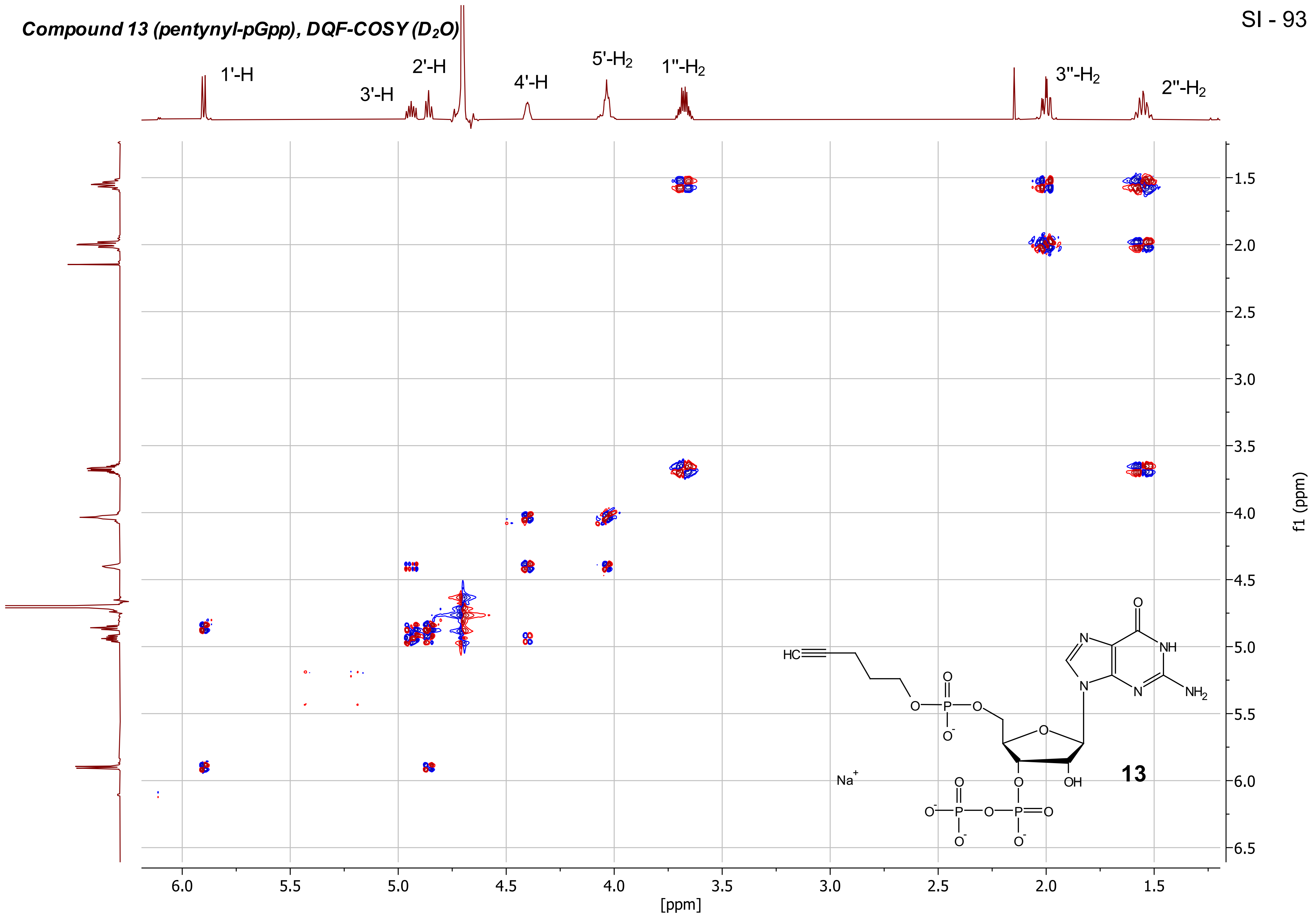

***Compound 13 (pentynyl-pGpp), HSQC ($D_2O$)***

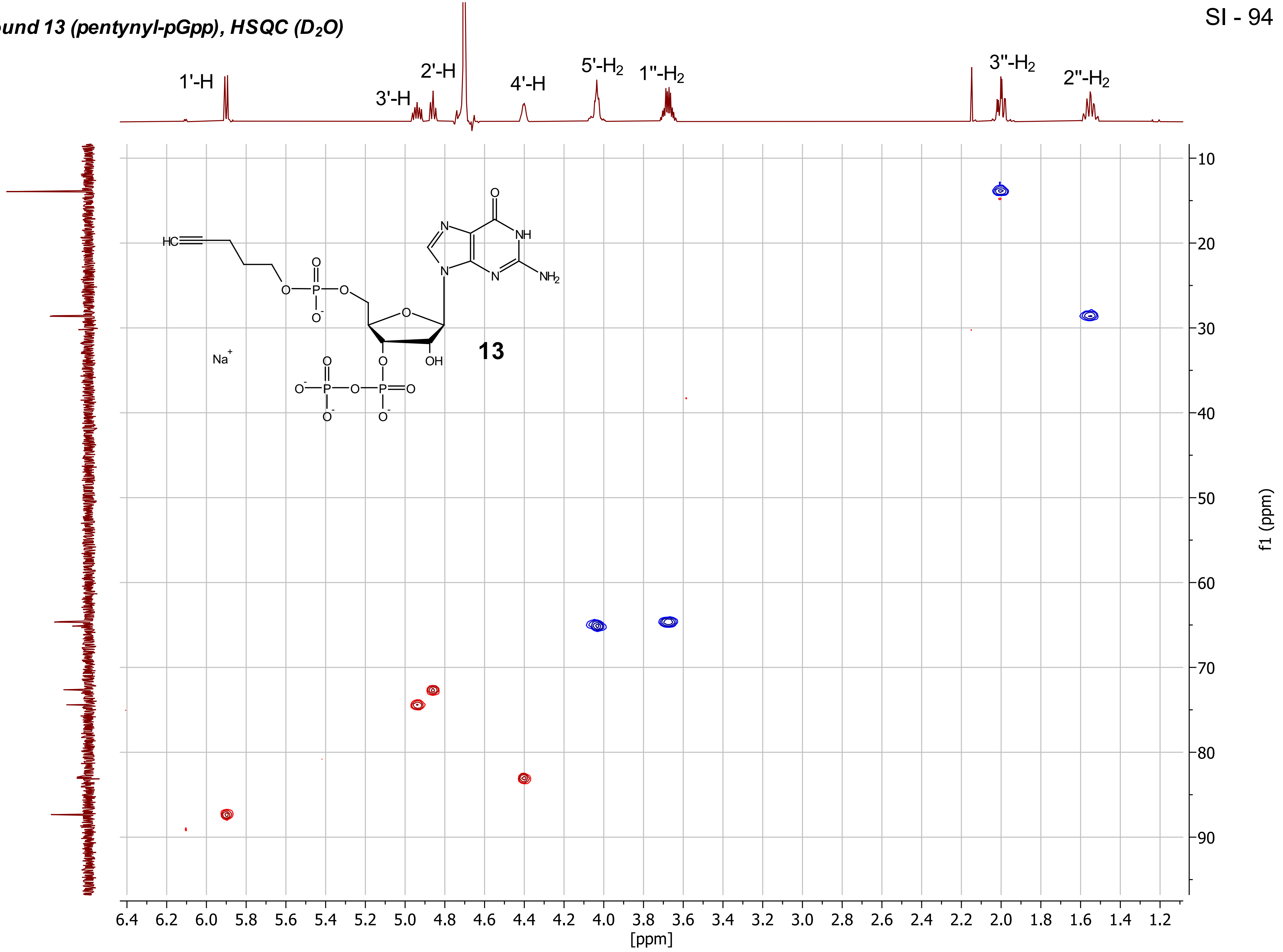

***Compound 13 (pentynyl-pGpp), 1H-31P-HMBC ($D_2O$)***

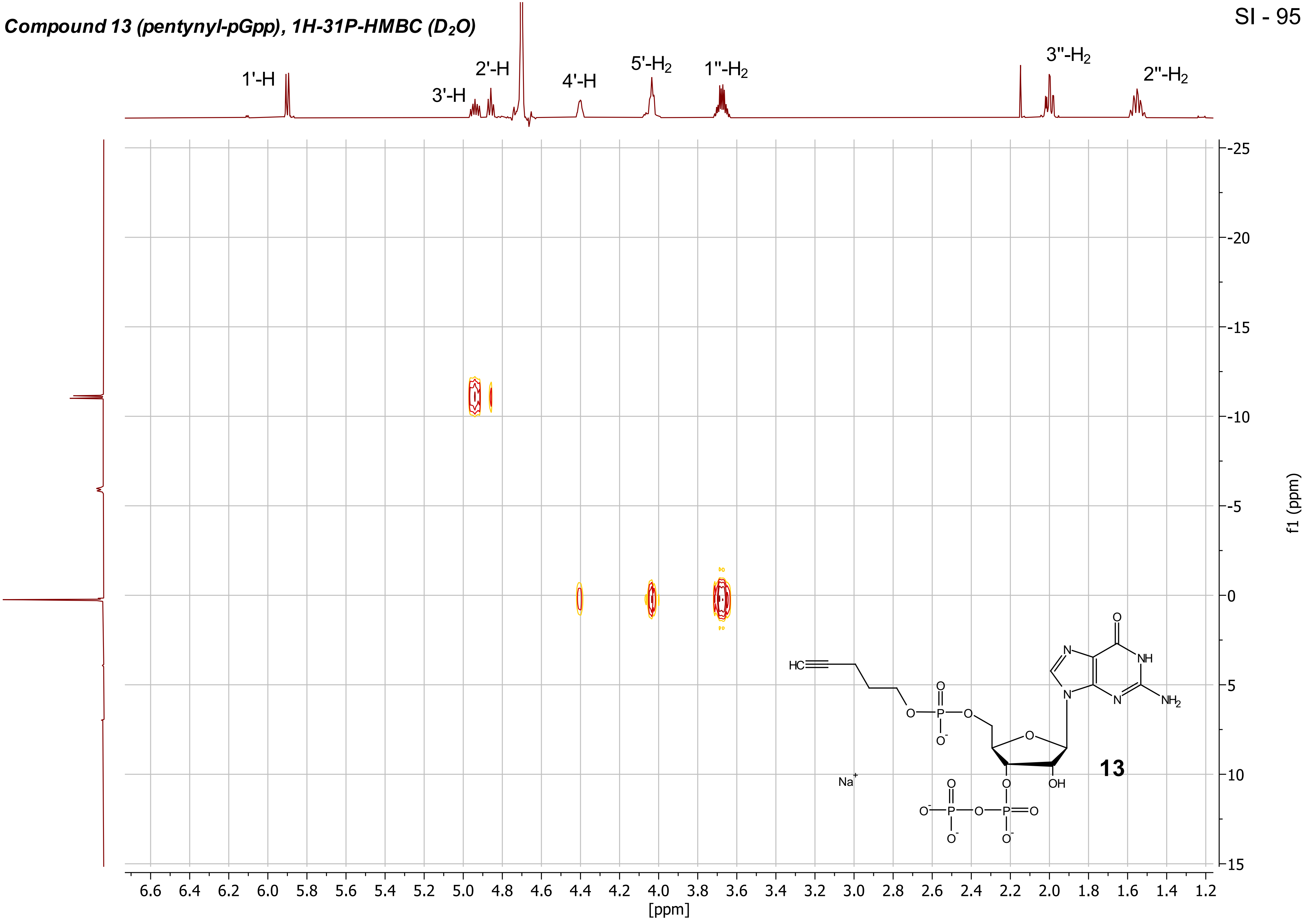

***Compound 13 (pentynyl-pGpp), $^{13}C\{^{1}H\}$ - NMR ($D_2O$, 101 MHz)***

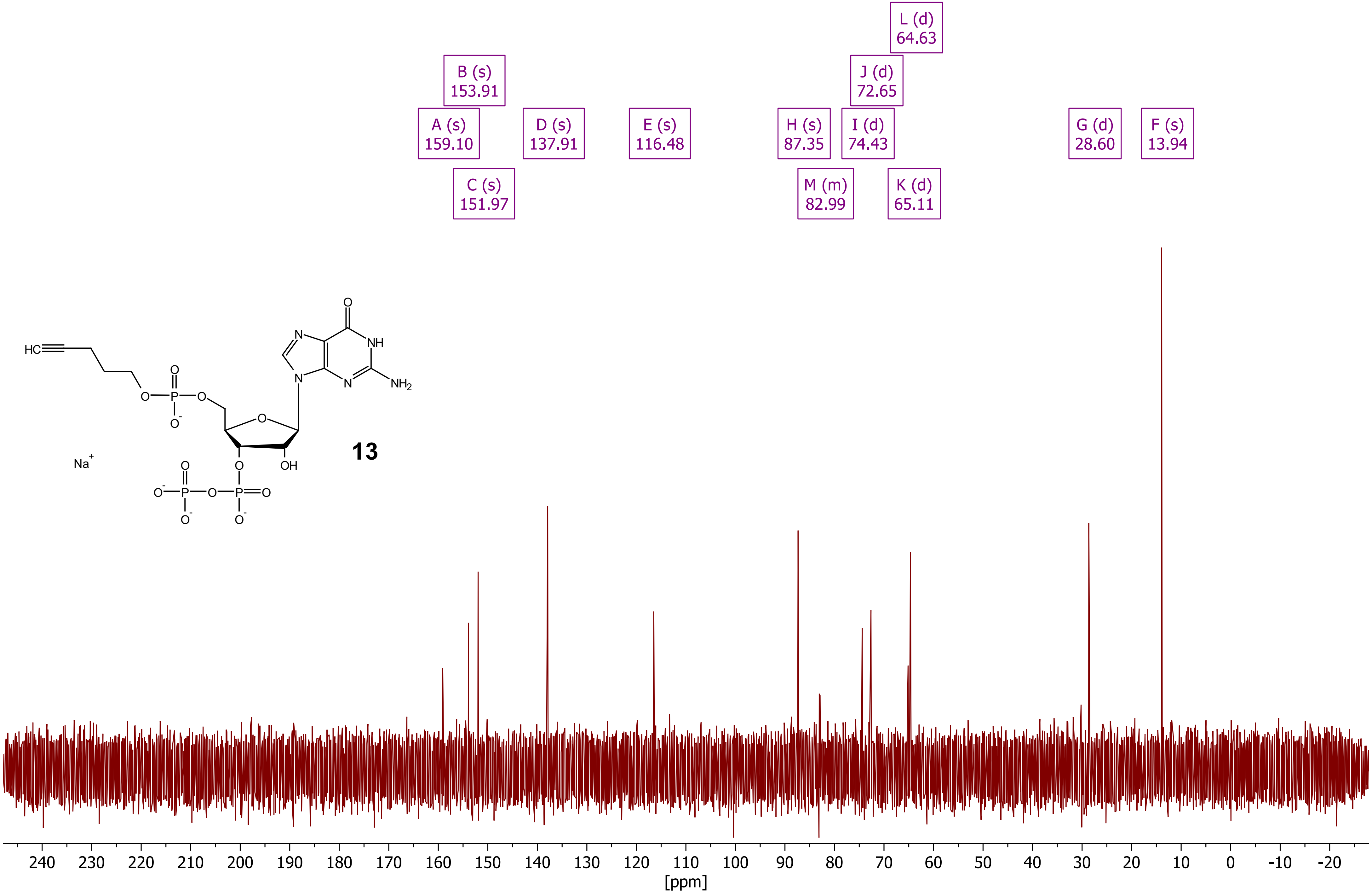

***Compound 15 (pentynyl-pppGpp), ¹H - NMR ($D_2O$, 400 MHz)***

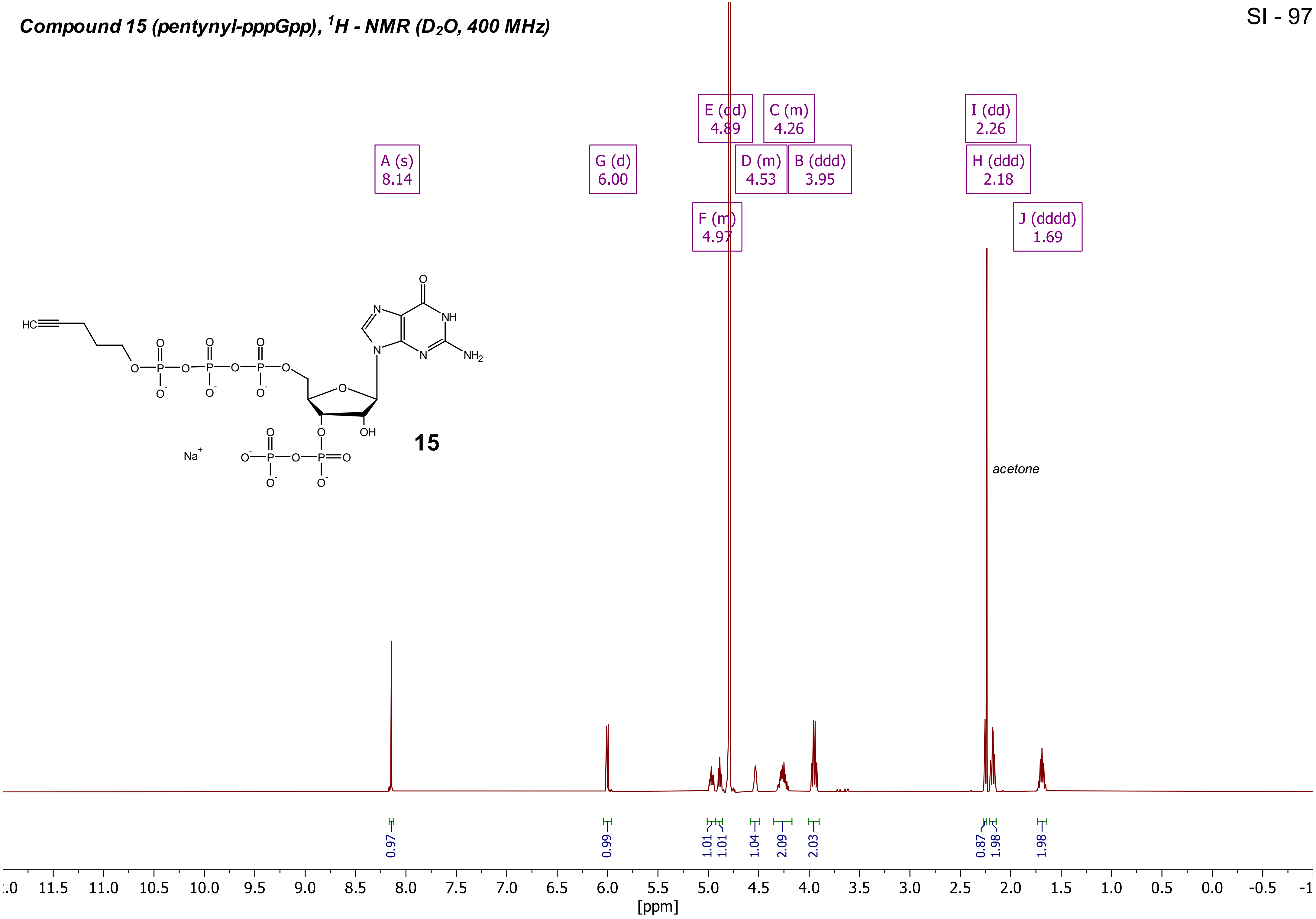

***Compound 15 (pentynyl-pppGpp), $^{31}P\{^{1}H\}$ - NMR ($D_2O$, 162 MHz)***

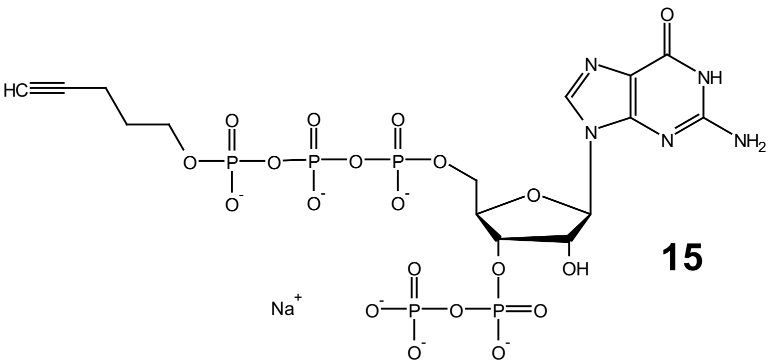

**15**

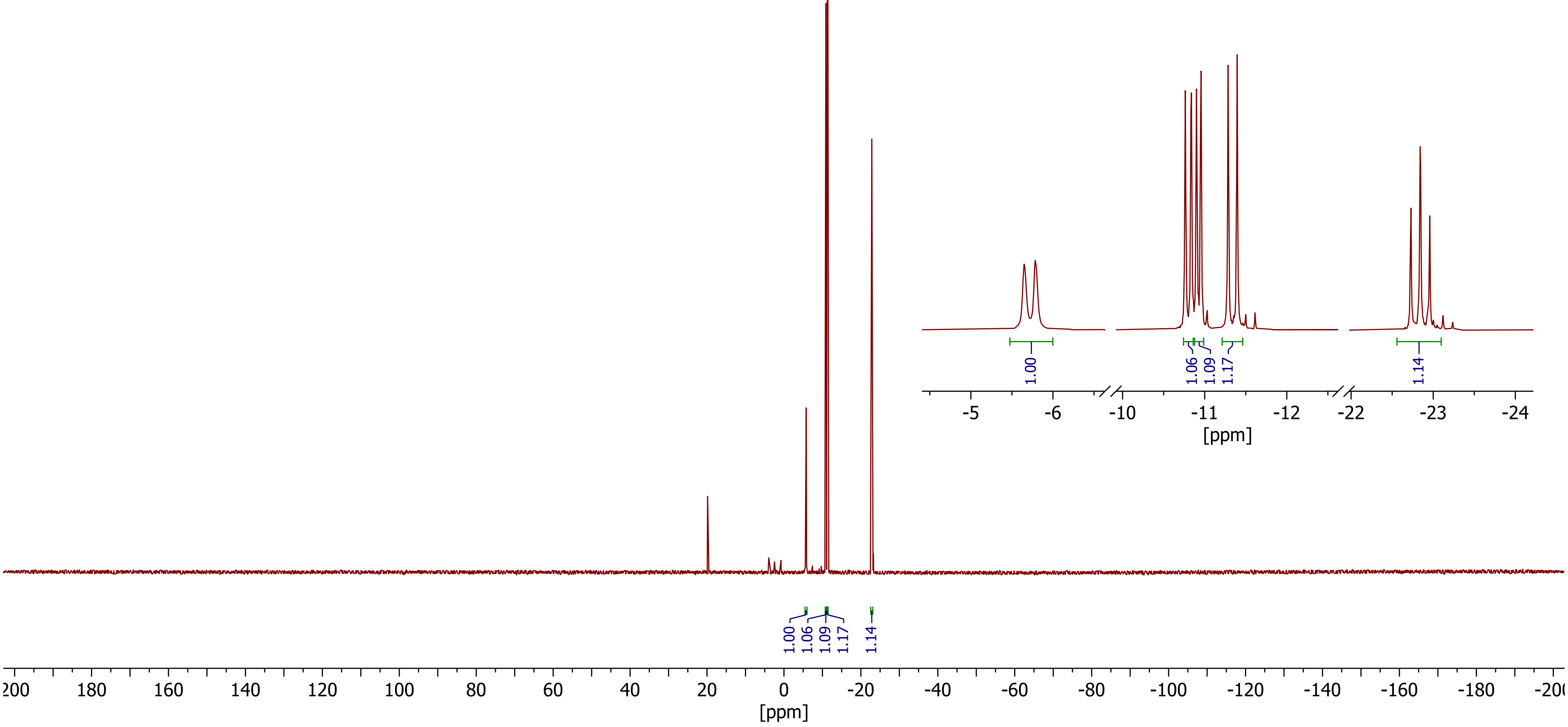

***Compound 15 (pentynyl-pppGpp), DQF-COSY ($D_2O$)***



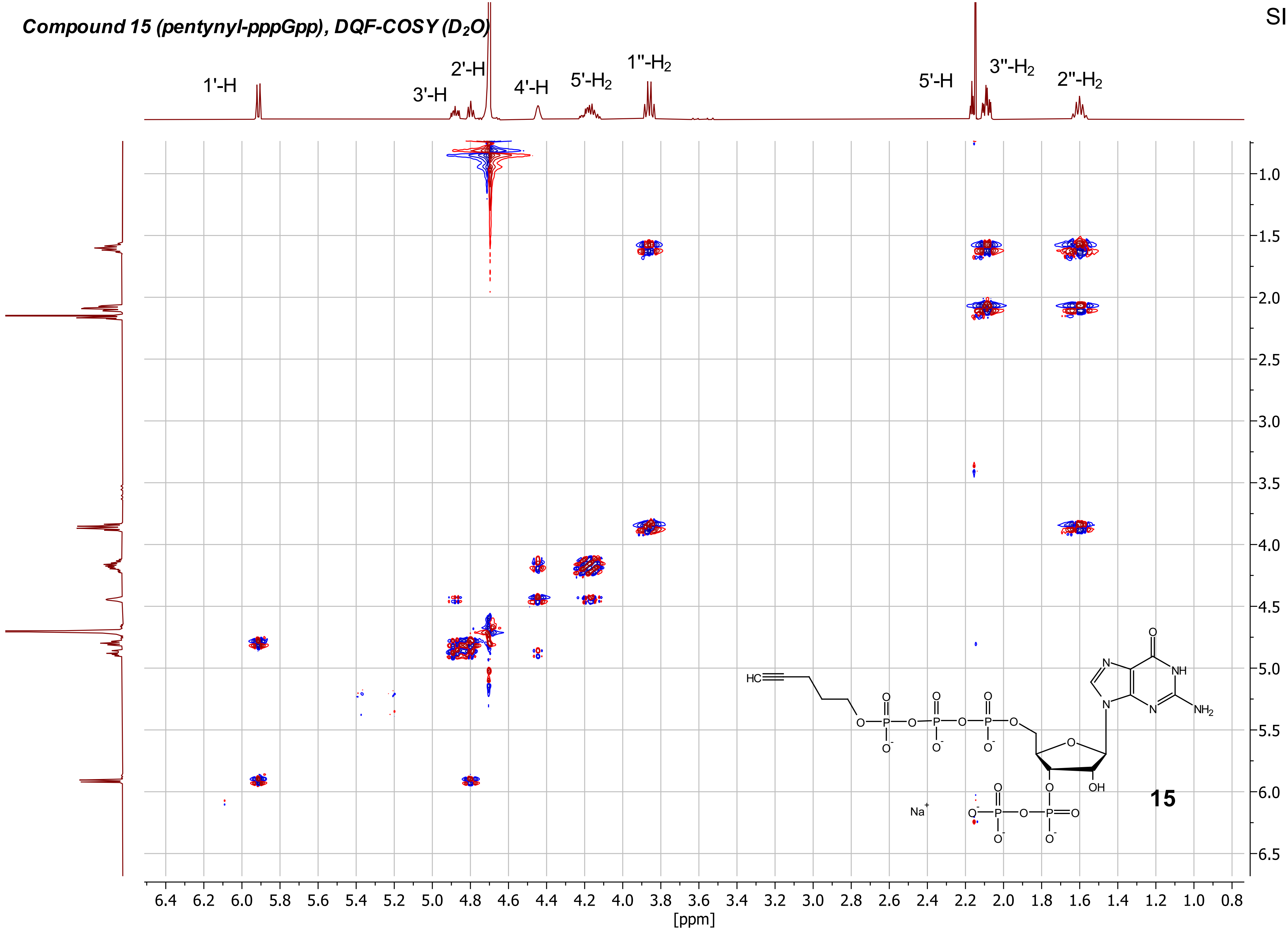

***Compound 15 (pentynyl-pppGpp), HSQC ($D_2O$)***

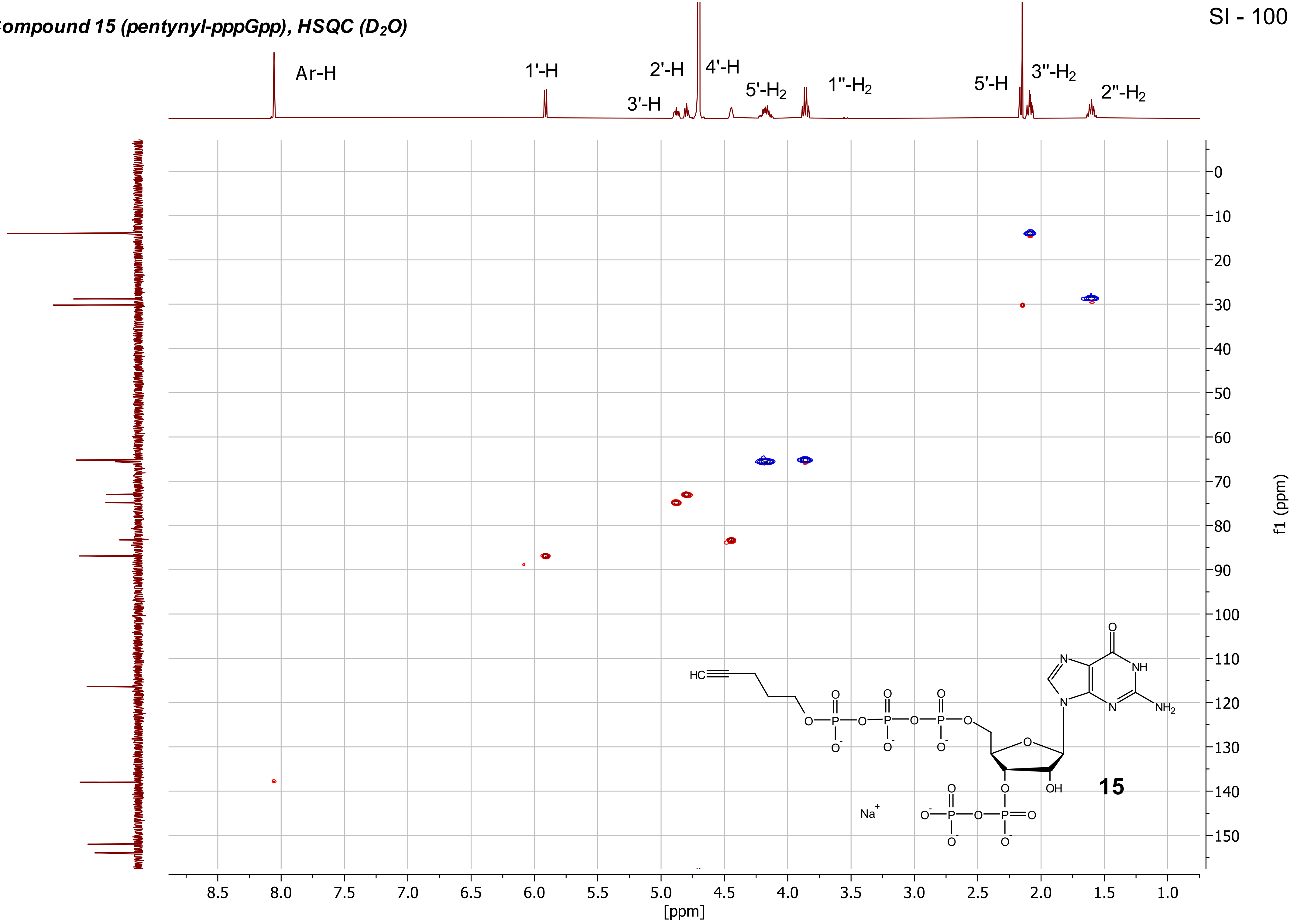

***Compound 15 (pentynyl-pppGpp), $^{1}H$-$^{31}P$-HMBC ($D_2O$)***

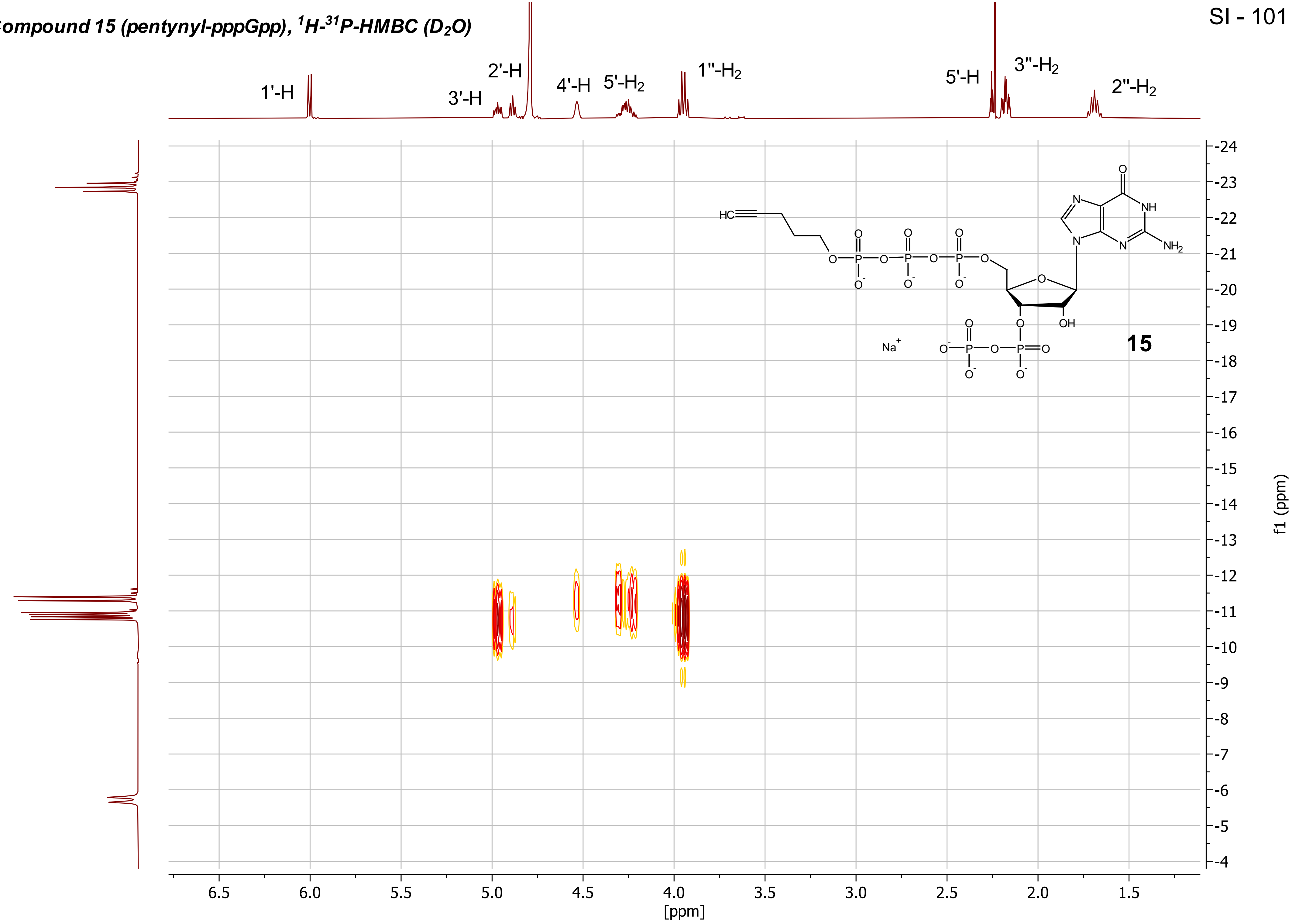

**Compound 15 (pentynyl-pppGpp), PP-COSY ($D_2O$)**



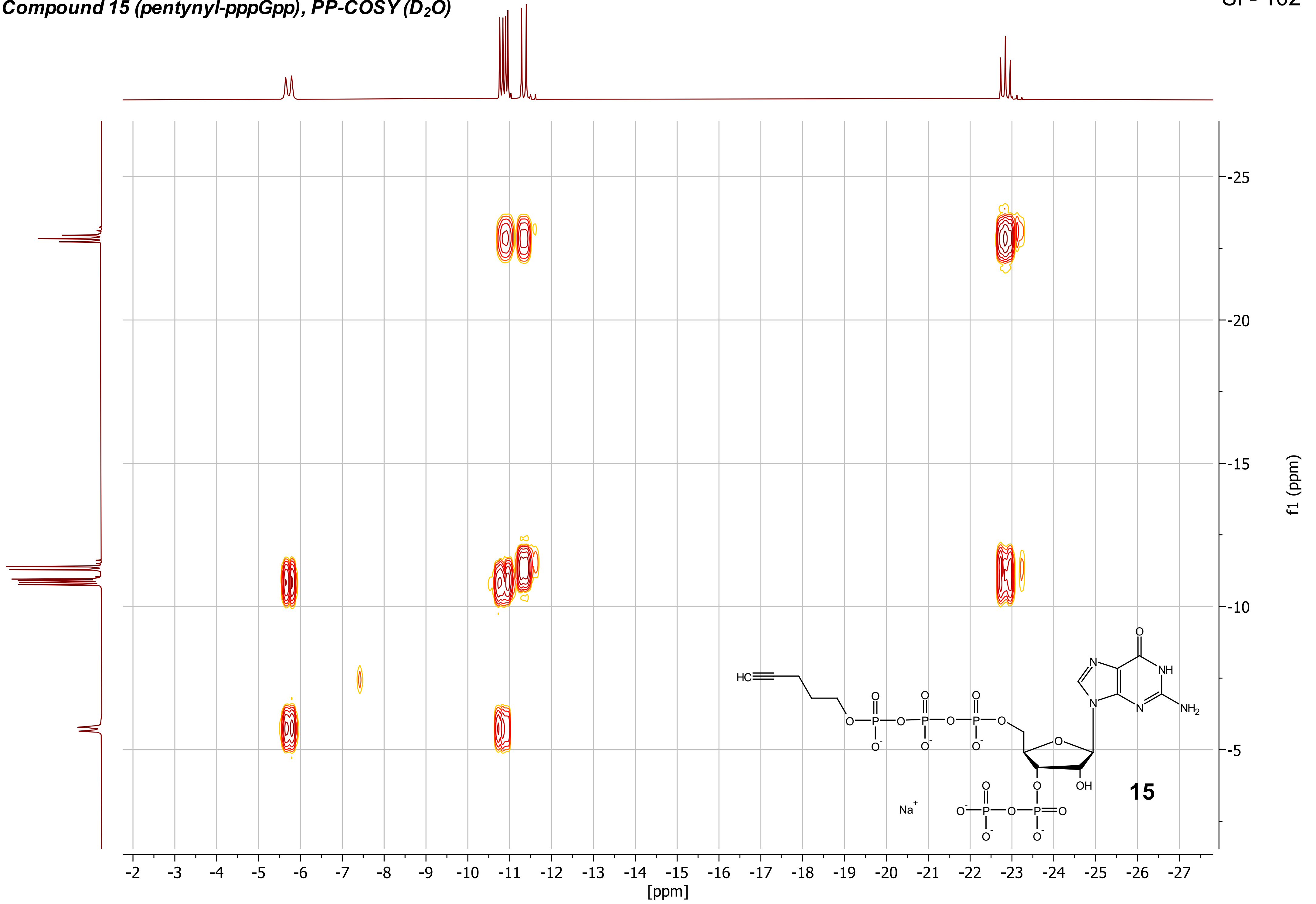


15

***Compound 15 (pentynyl-pppGpp), $^{13}C\{^1H\}$ - NMR ($D_2O$, 101 MHz)***

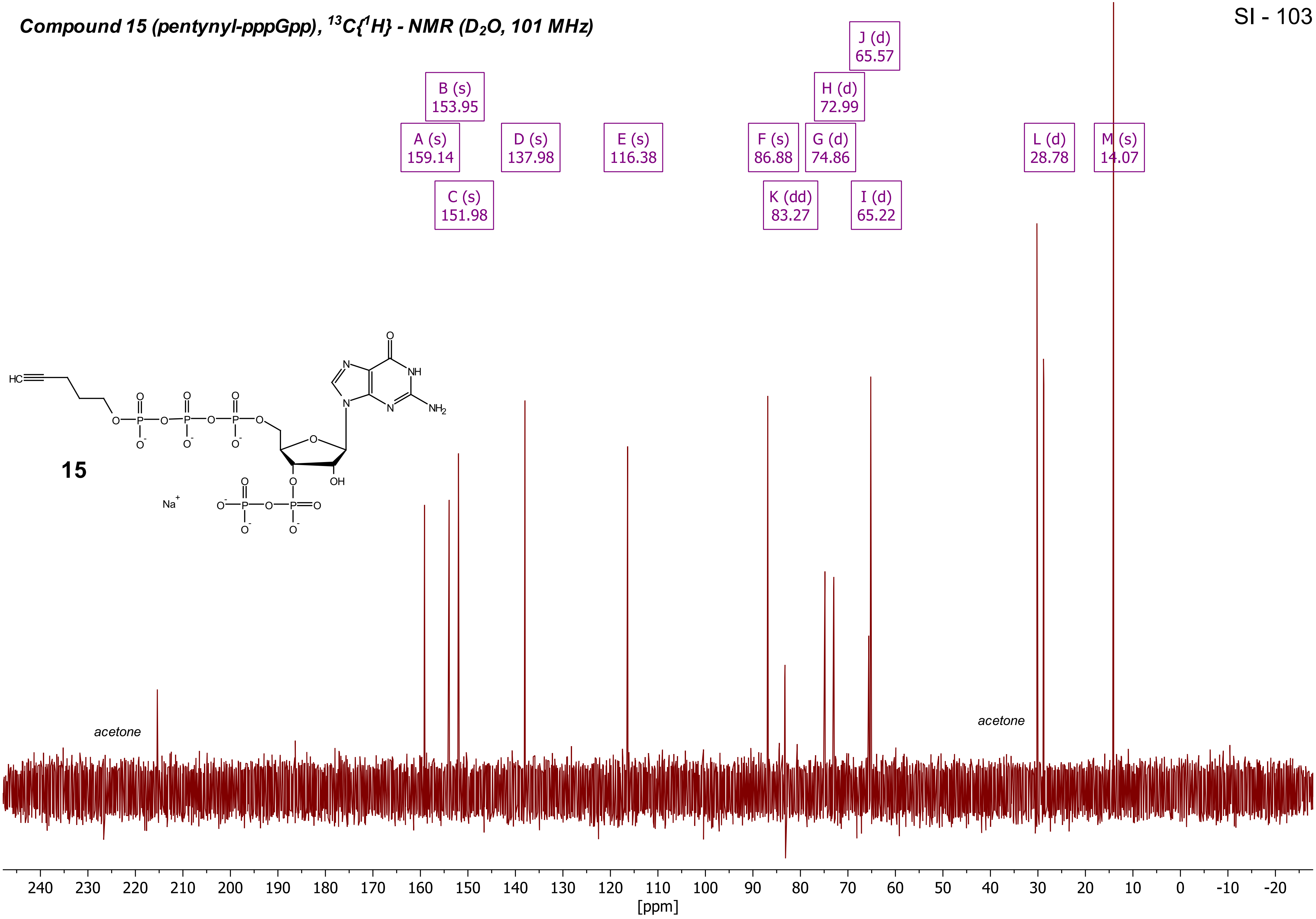

***Compound 16 (pentynyl-ppApp), $^1H$ - NMR ($D_2O$, 400 MHz)***

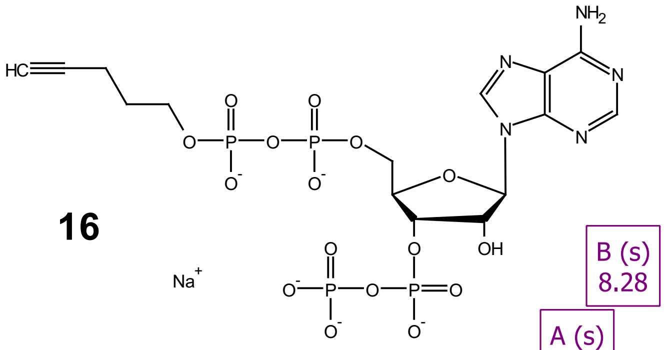


**16**

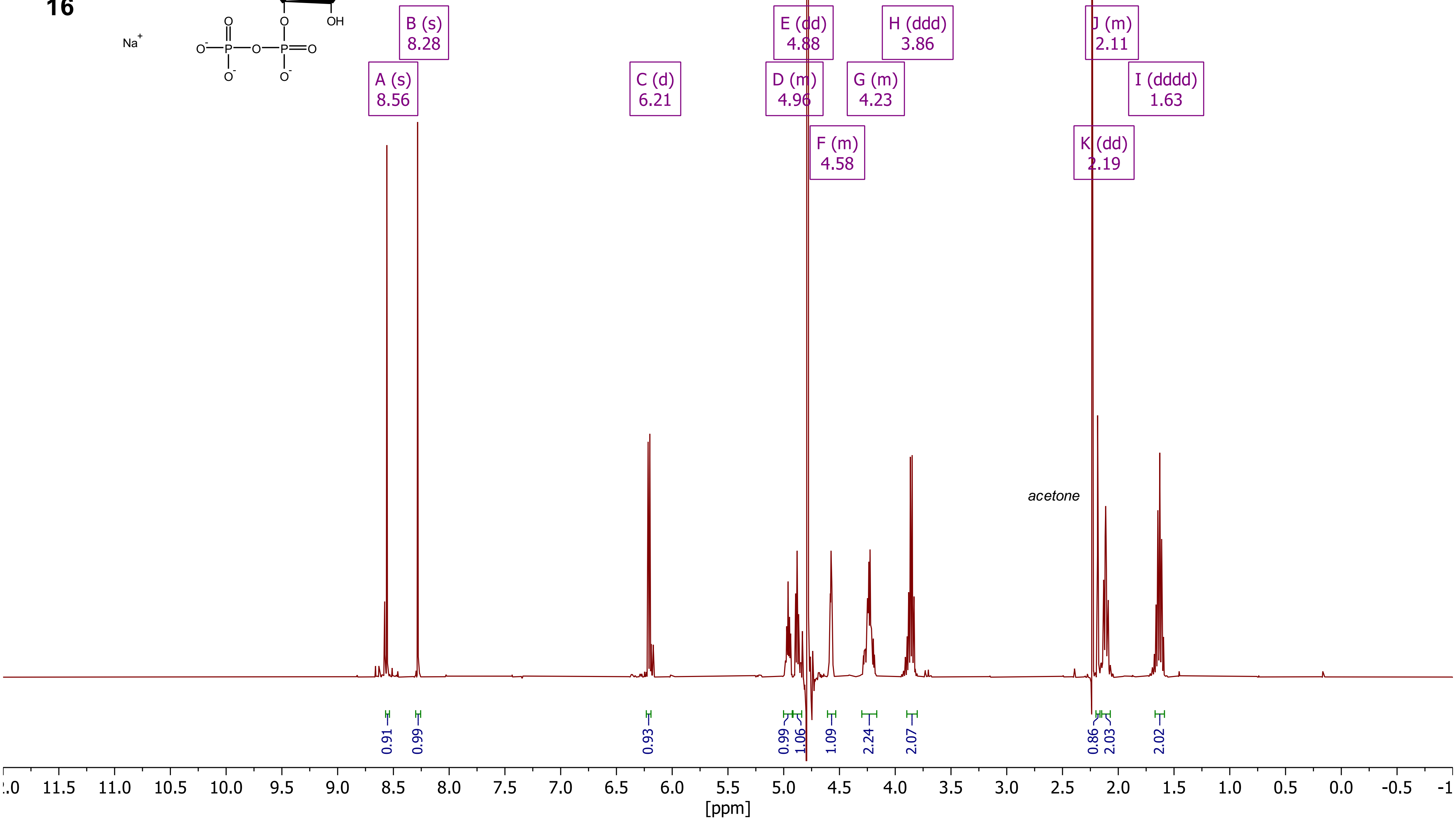

***Compound 16 (pentynyl-ppApp), $^{31}P\{^1H\}$ - NMR ($D_2O$, 162 MHz)***

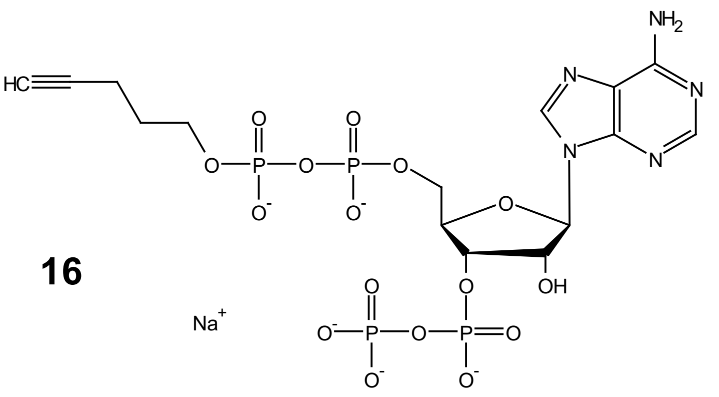

**16**

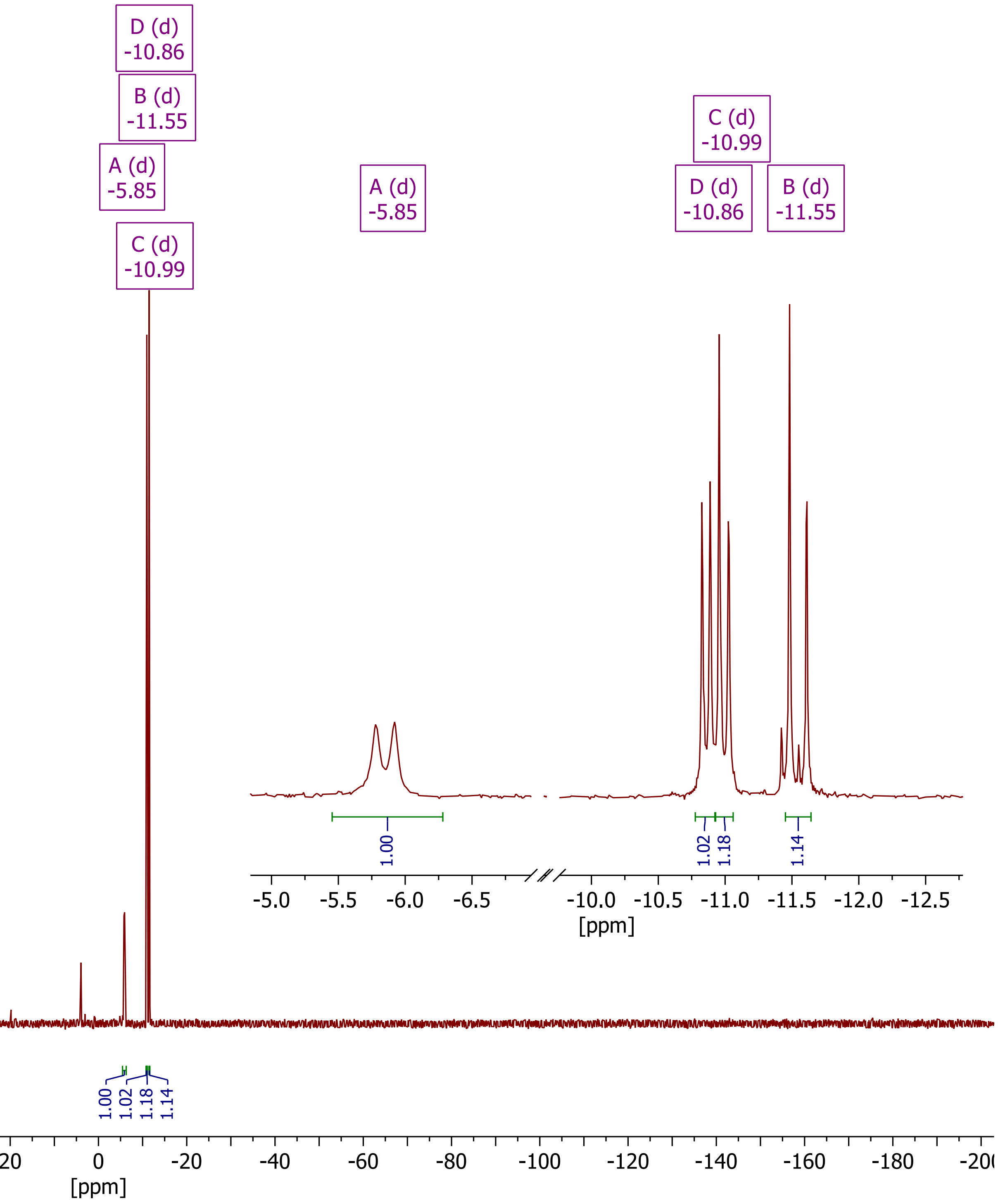

***Compound 16 (pentynyl-ppApp), DQF-COSY ($D_2O$, 162 MHz)***

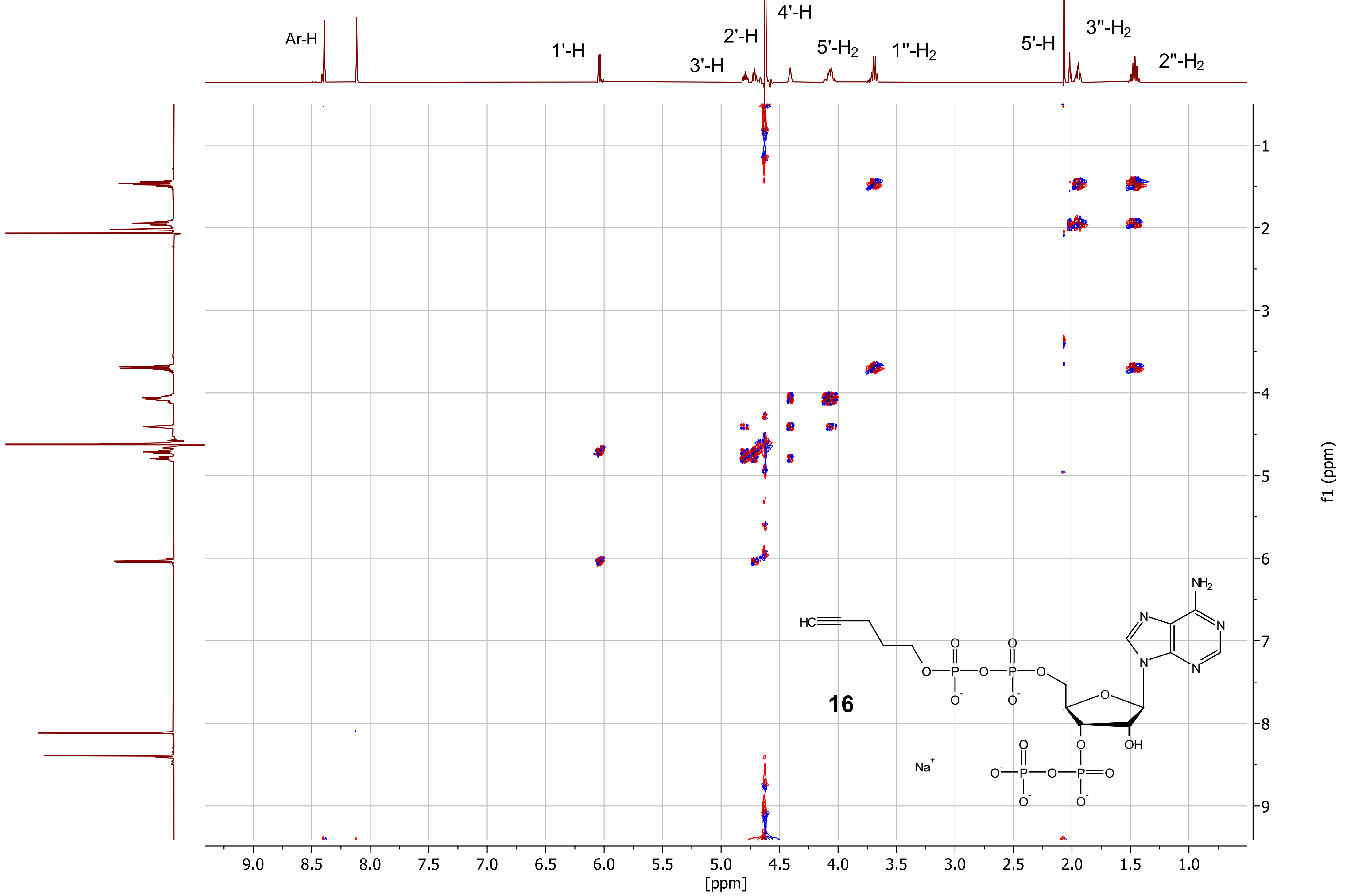

***Compound 16 (pentynyl-ppApp), HSQC ($D_2O$)***

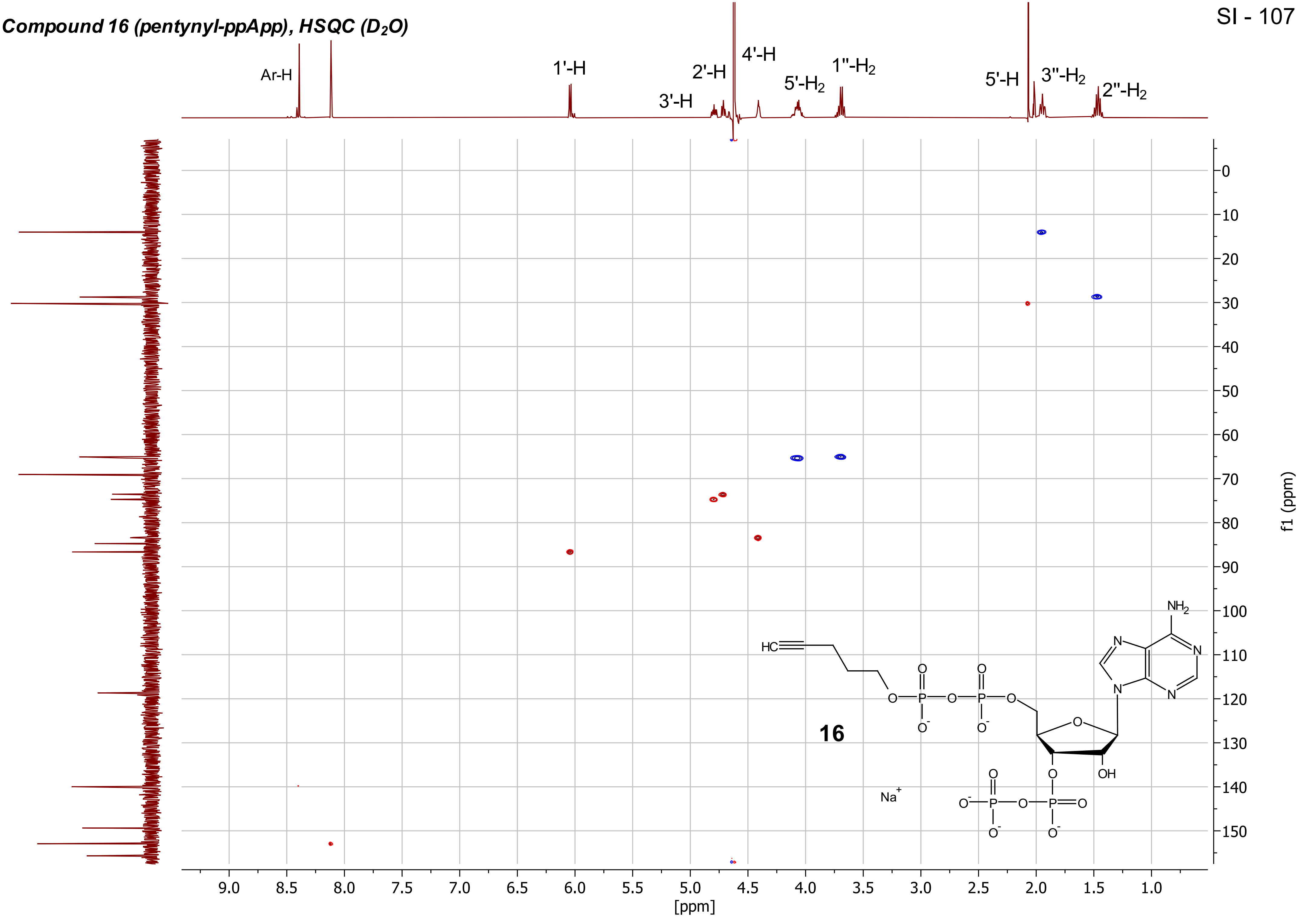

***Compound 16 (pentynyl-ppApp), $^{1}H$-$^{31}P$-HMBC ($D_2O$)***

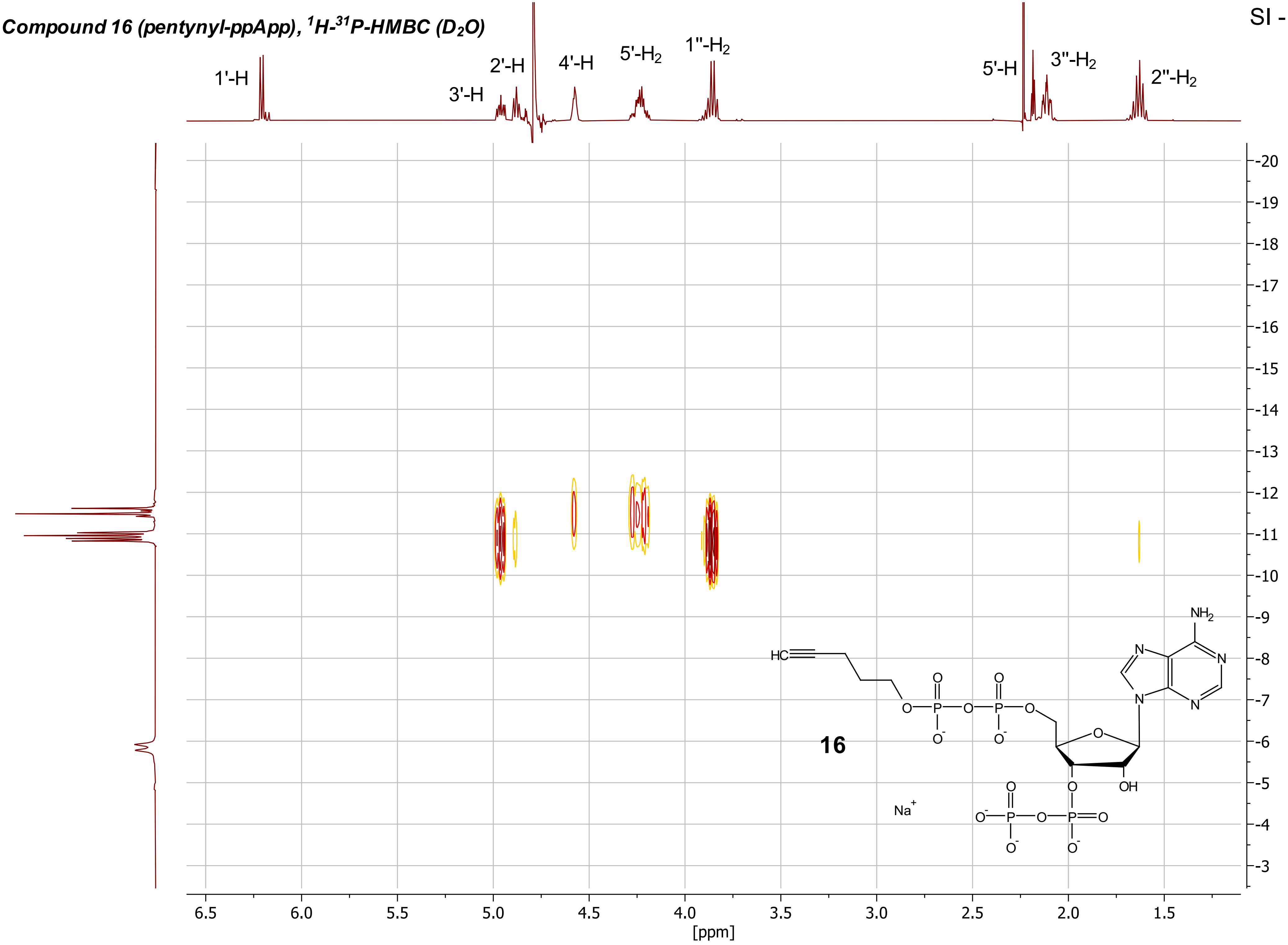

***Compound 16 (pentynyl-ppApp), PP - COSY ($D_2O$)***

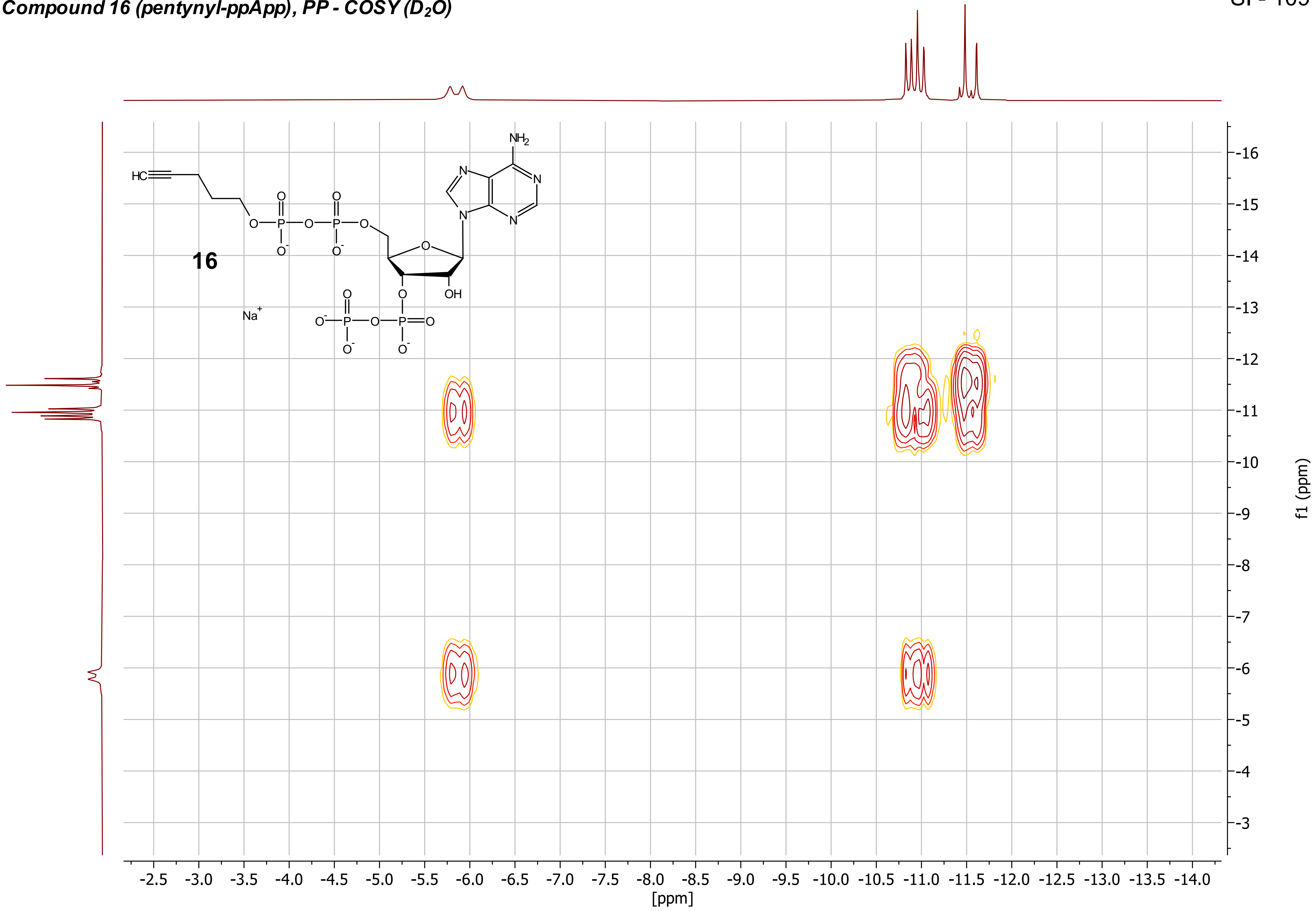

***Compound 16 (pentynyl-ppApp), $^{13}C\{^{1}H\}$ - NMR ($D_2O$, 101 MHz)***

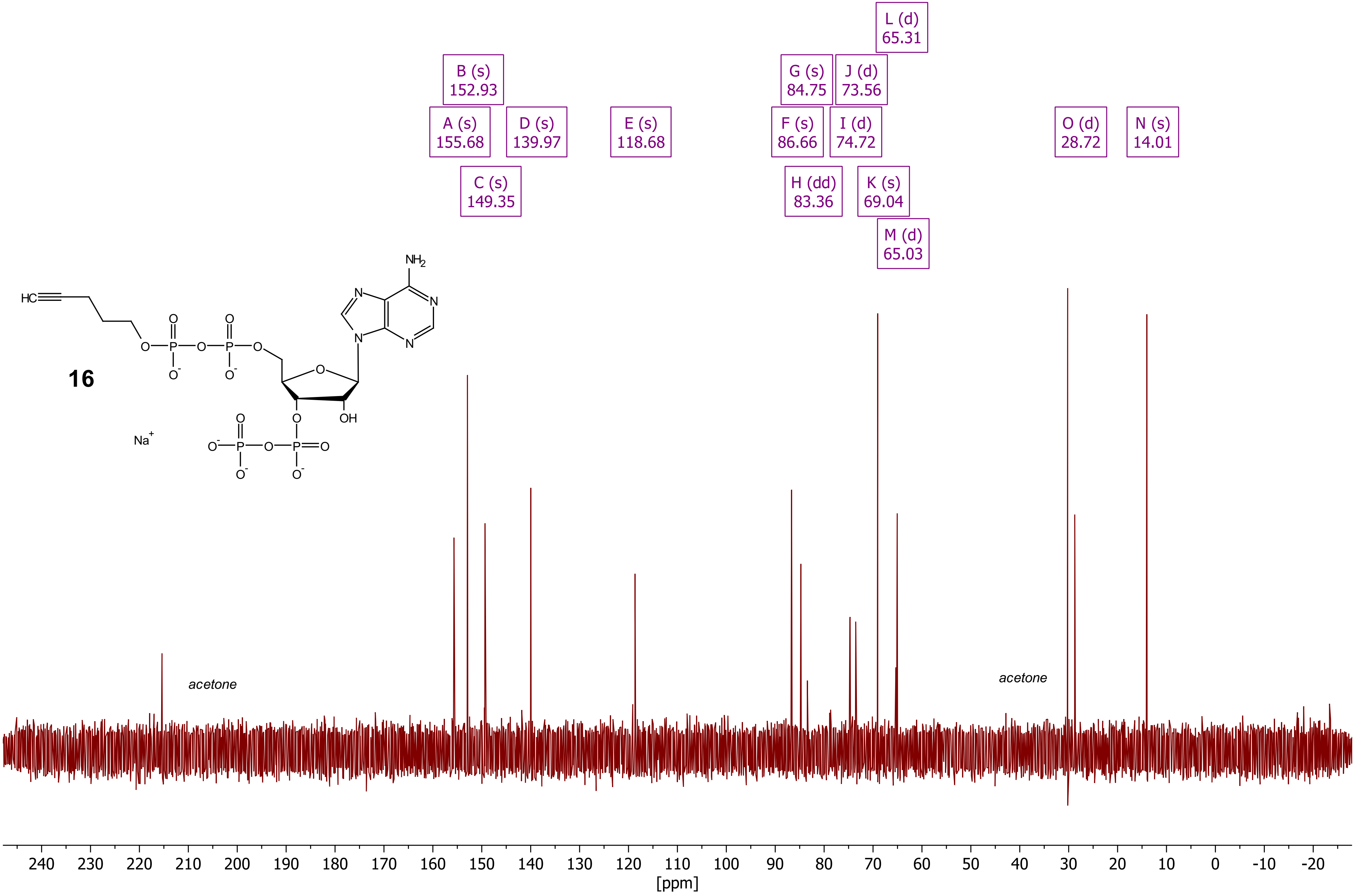

***Compound 21 (ppGpp - CC2), $^{1}H$ - NMR ($D_2O$, 400 MHz)***

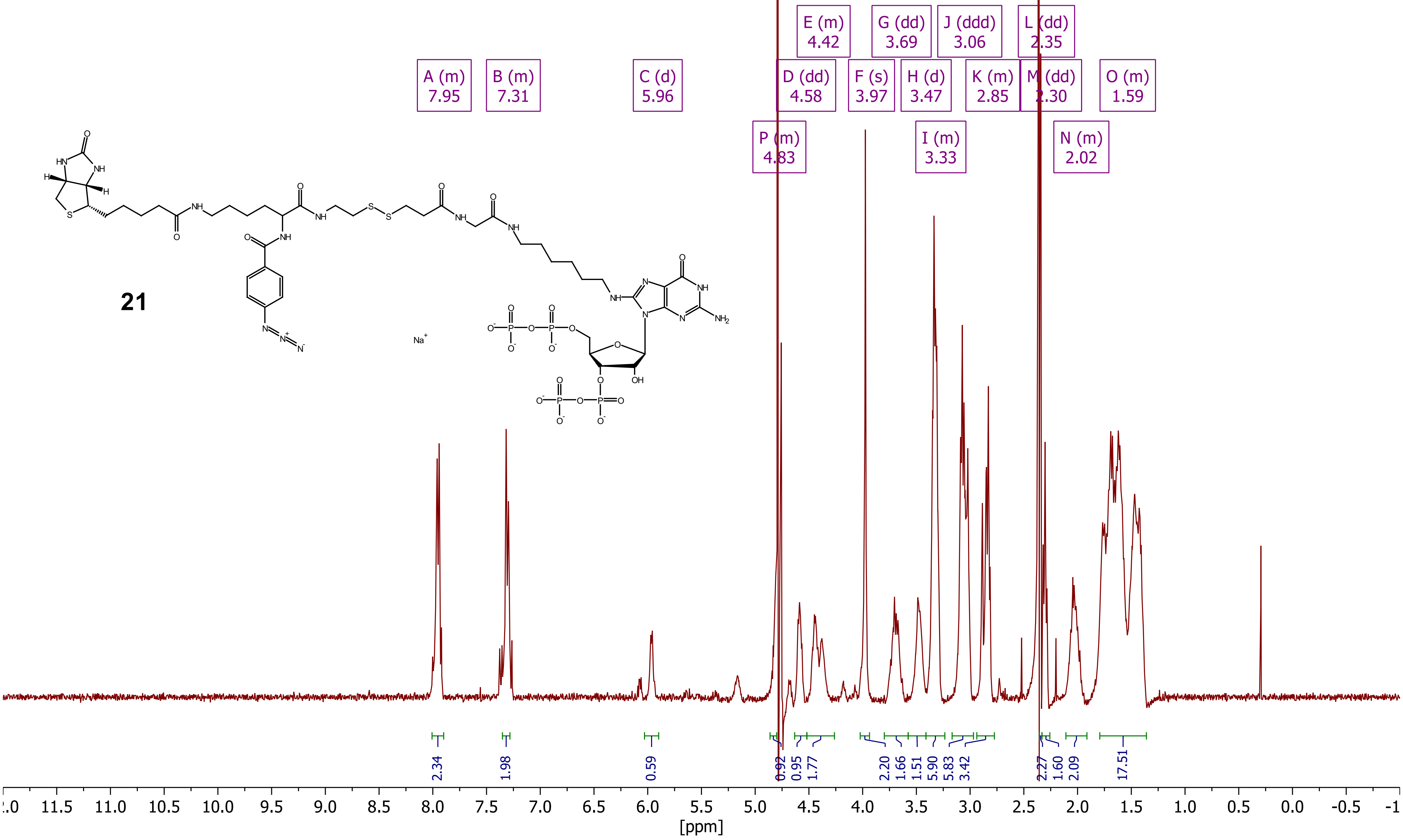

***Compound 21 (ppGpp - CC2), $^{31}$P - NMR ($D_2O$, 162 MHz)***

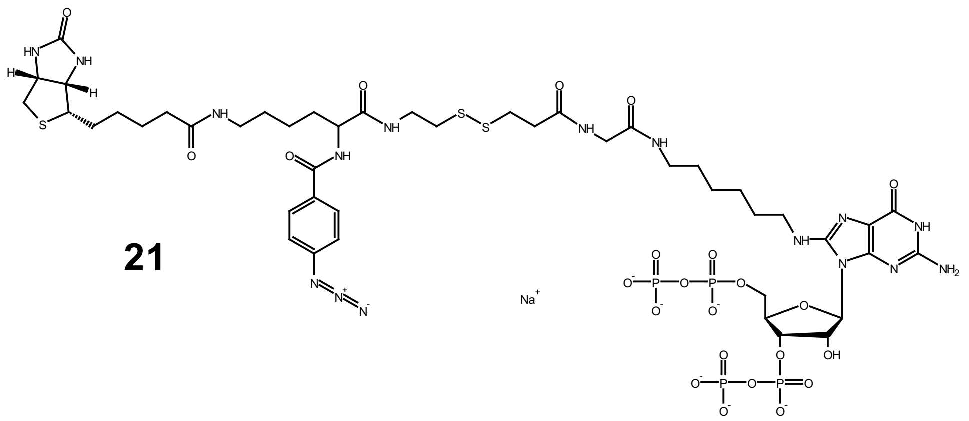


**21**

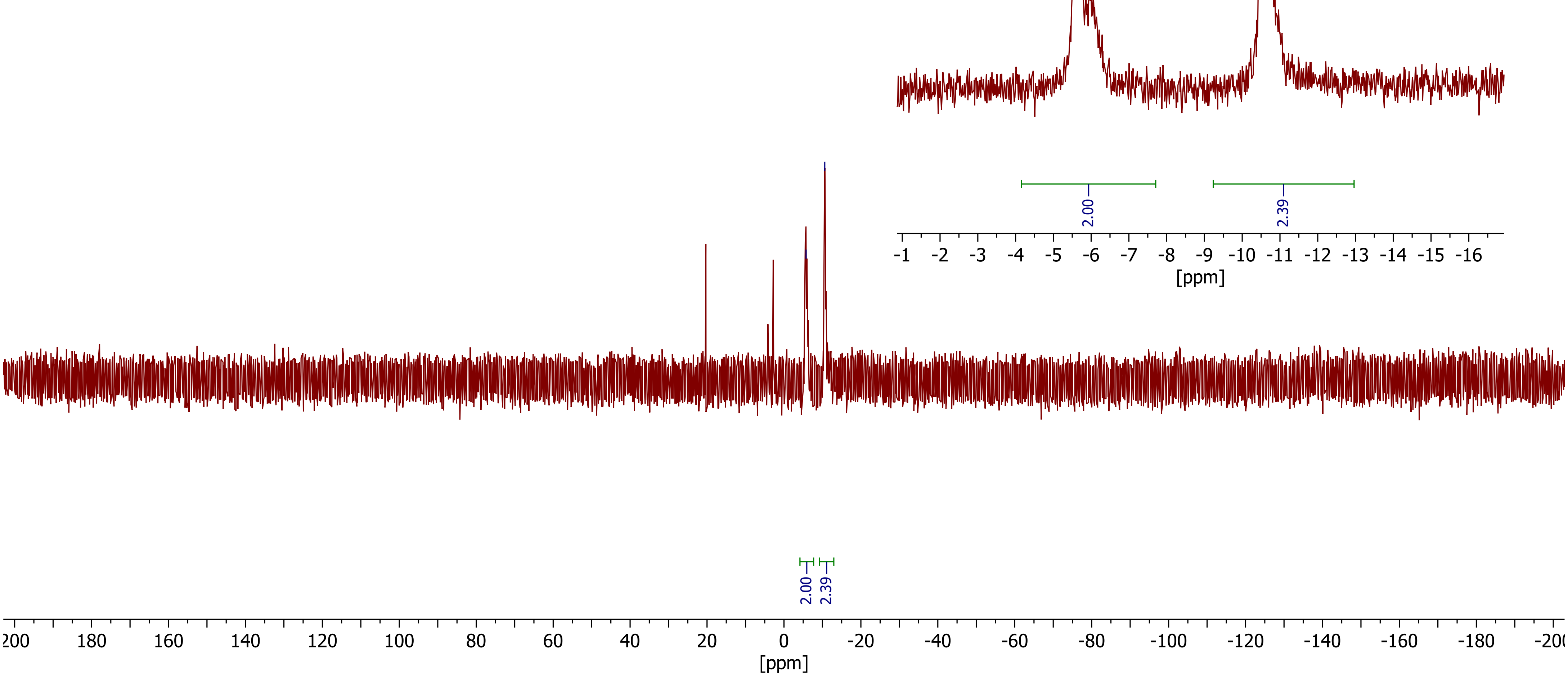

***Compound 22 (ppGpp - CC1), $^{1}H$ - NMR ($D_2O$, 400 MHz)***

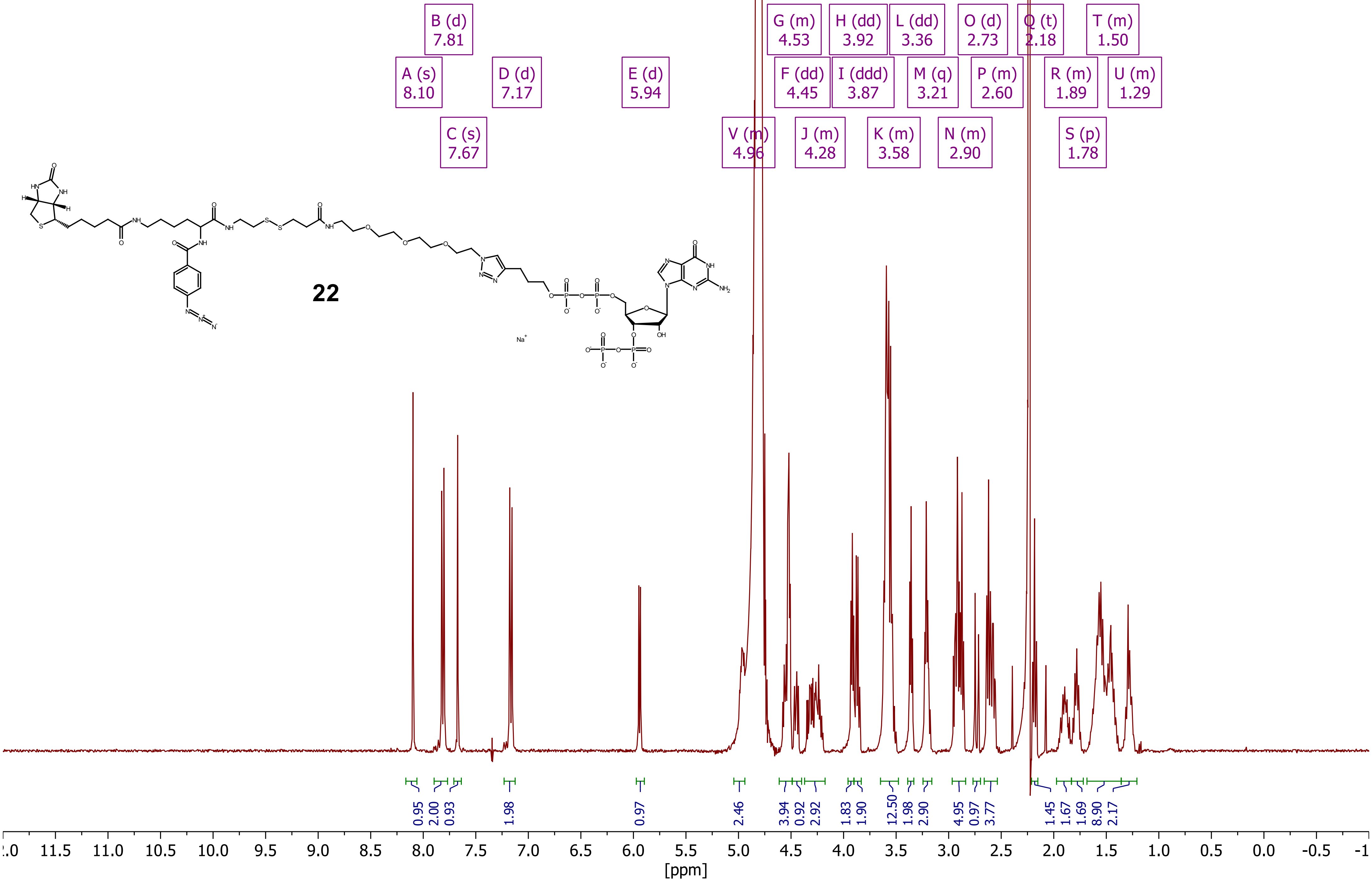

***Compound 22 (ppGpp - CC1), $^{31}P$ - NMR ($D_2O$, 162 MHz)***

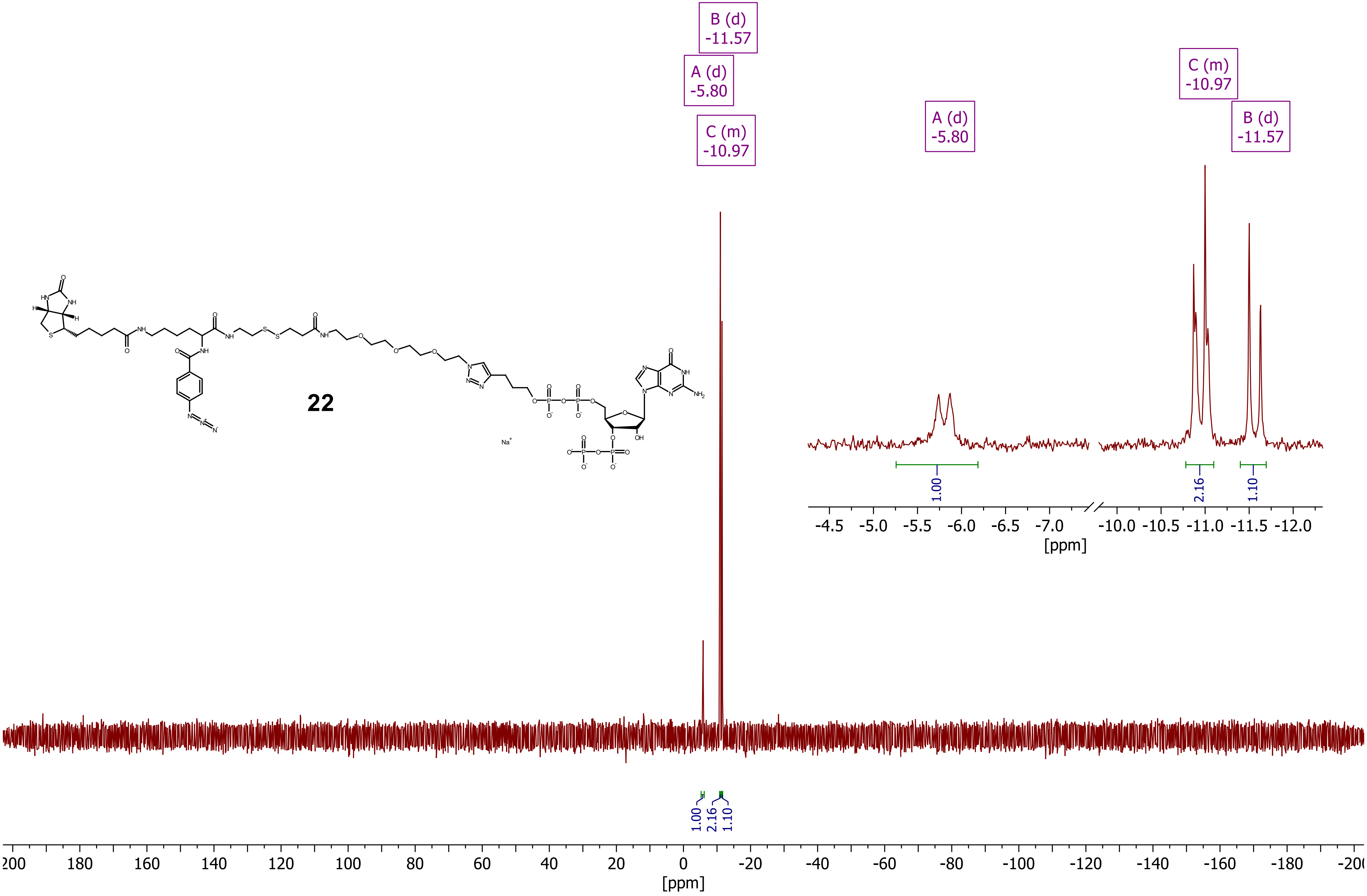

***Compound 23 (ppGpp - CC3), $^1$H - NMR ($D_2O$, 400 MHz)***

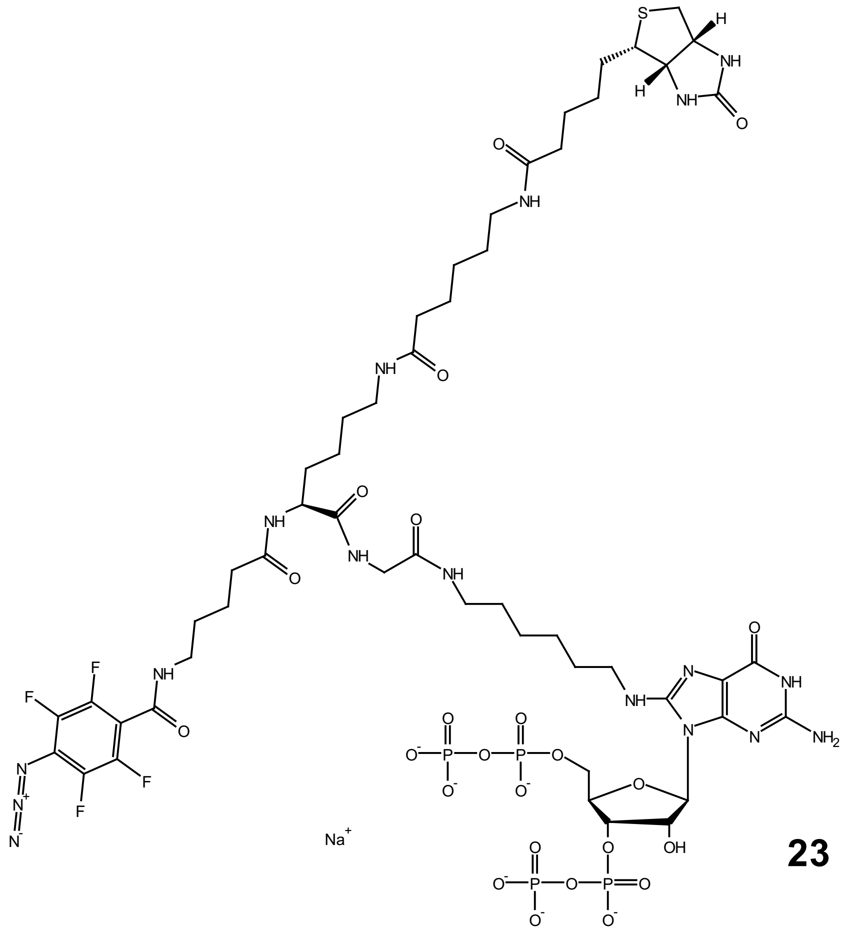

**23**

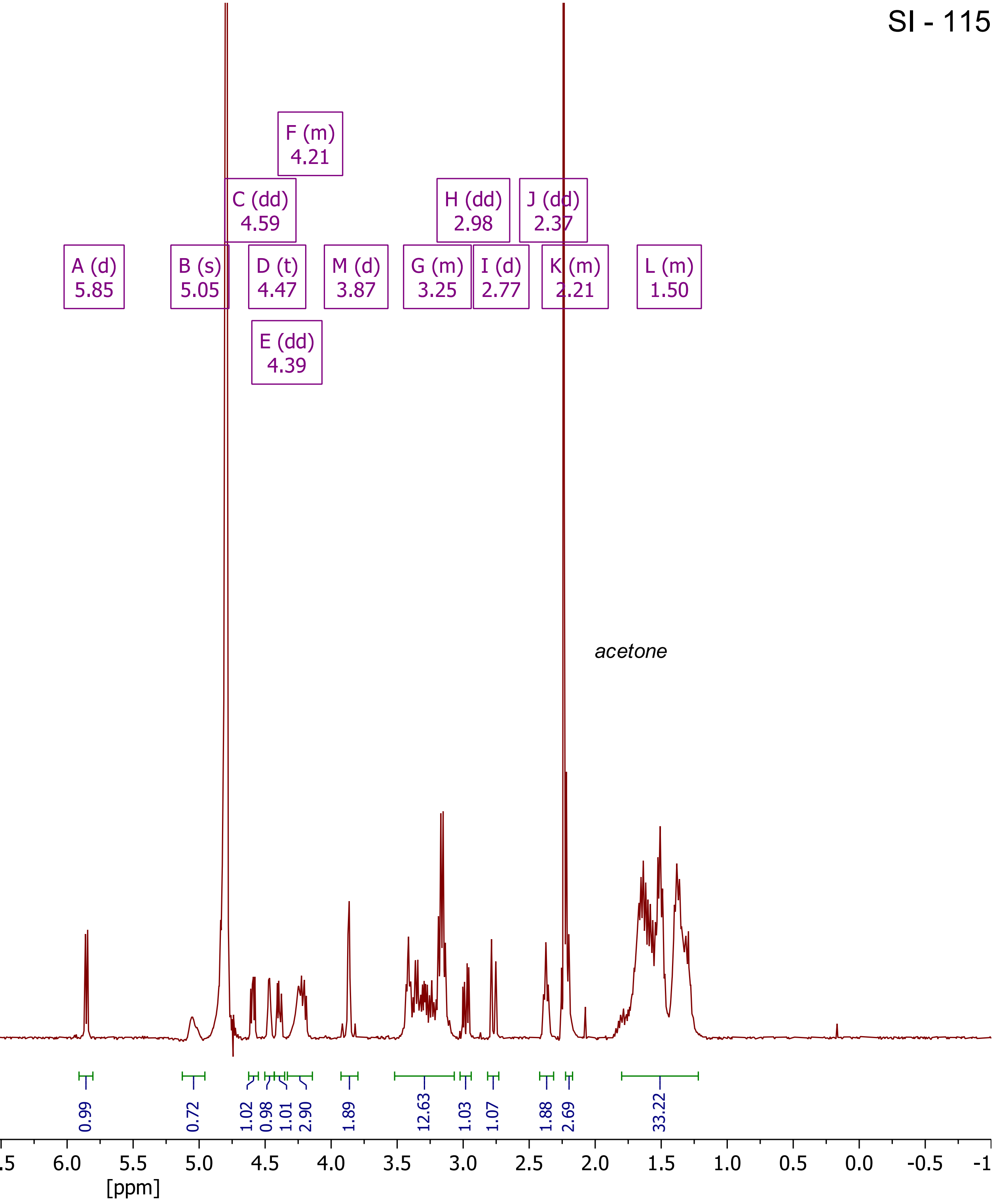

***Compound 23 (ppGpp - CC3), ${}^{19}F$ - NMR ($D_2O$, 377 MHz)***

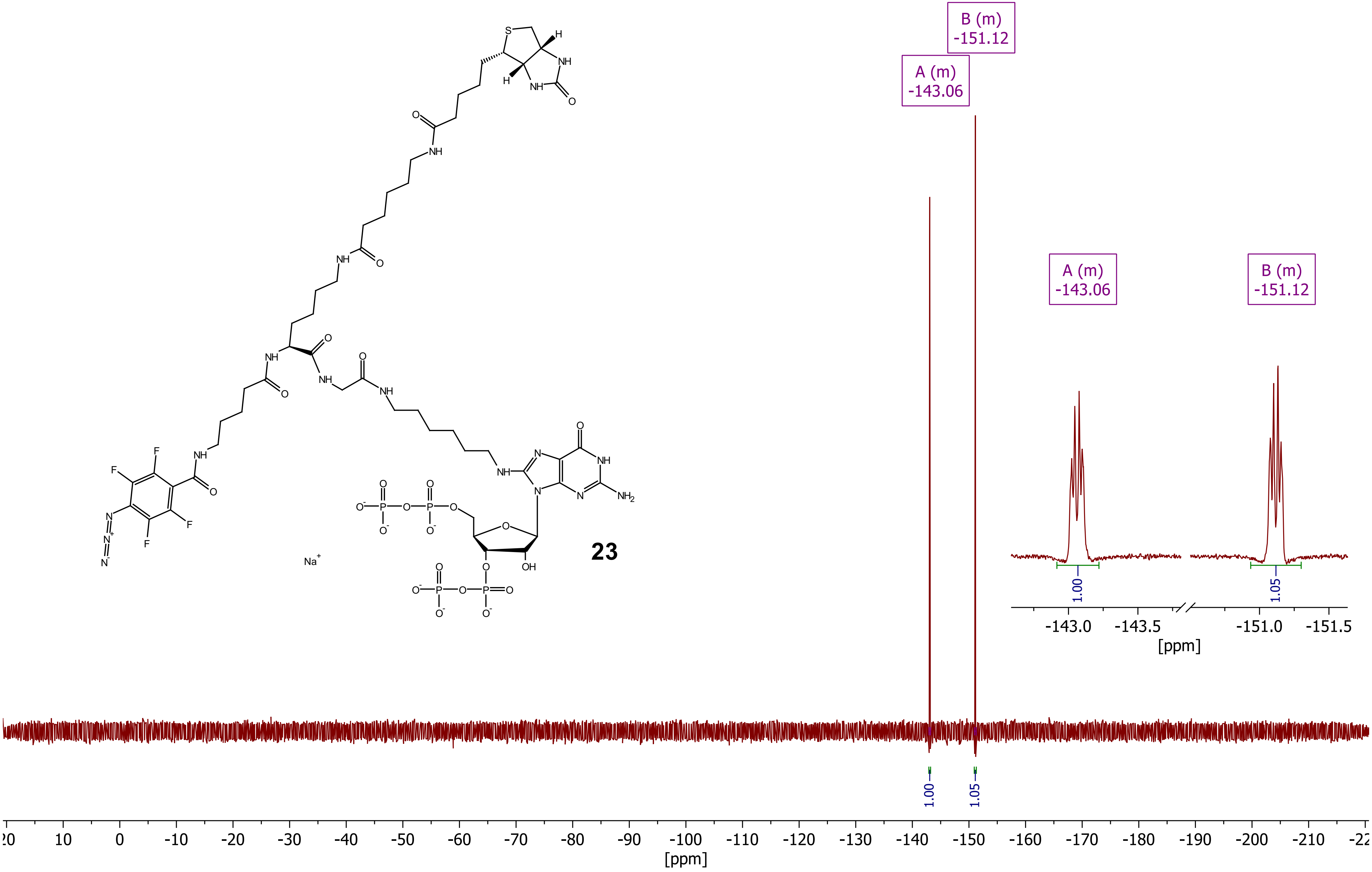

***Compound 23 (ppGpp - CC3), $^{31}P\{^{1}H\}$ - NMR ($D_2O$, 162 MHz)***

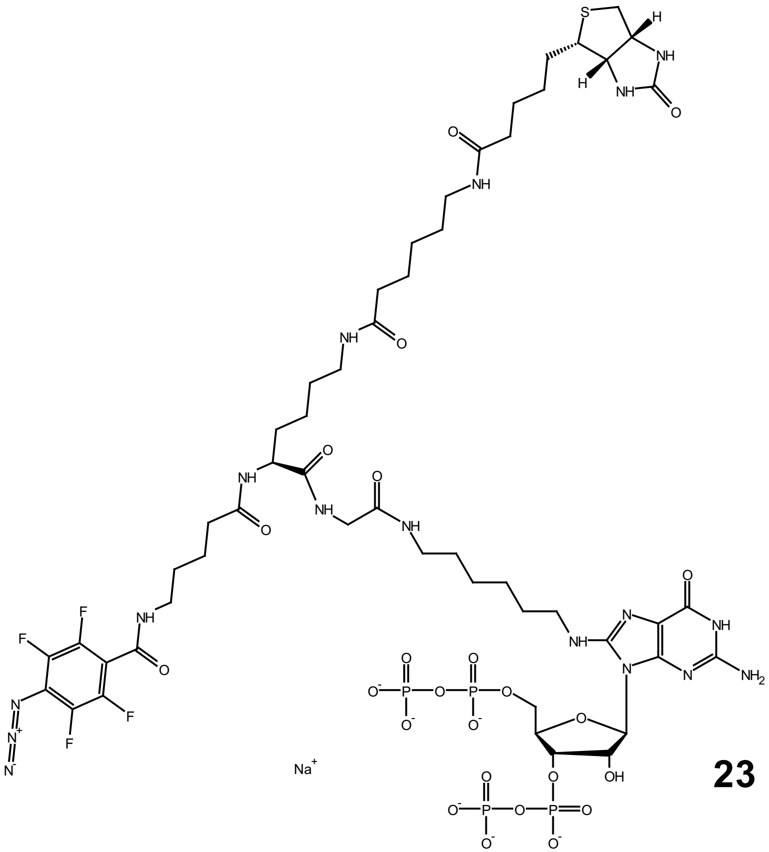

**23**

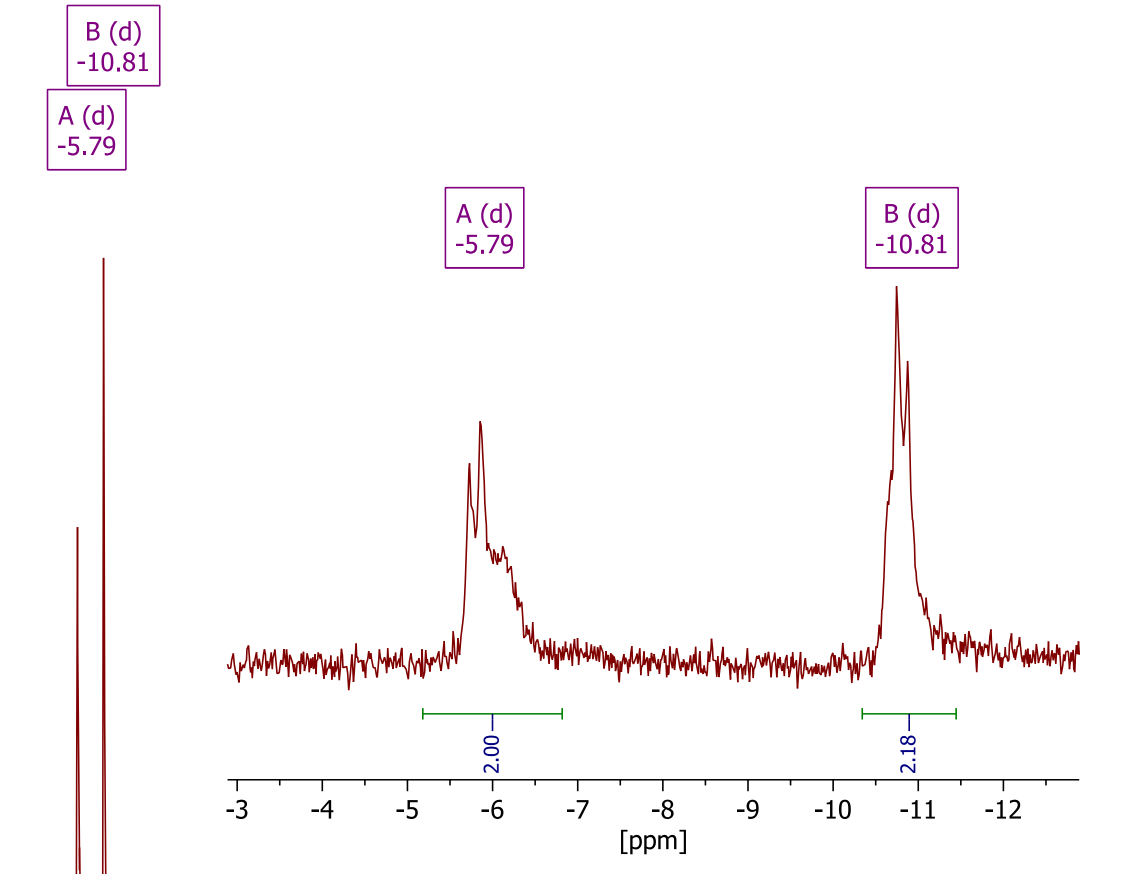


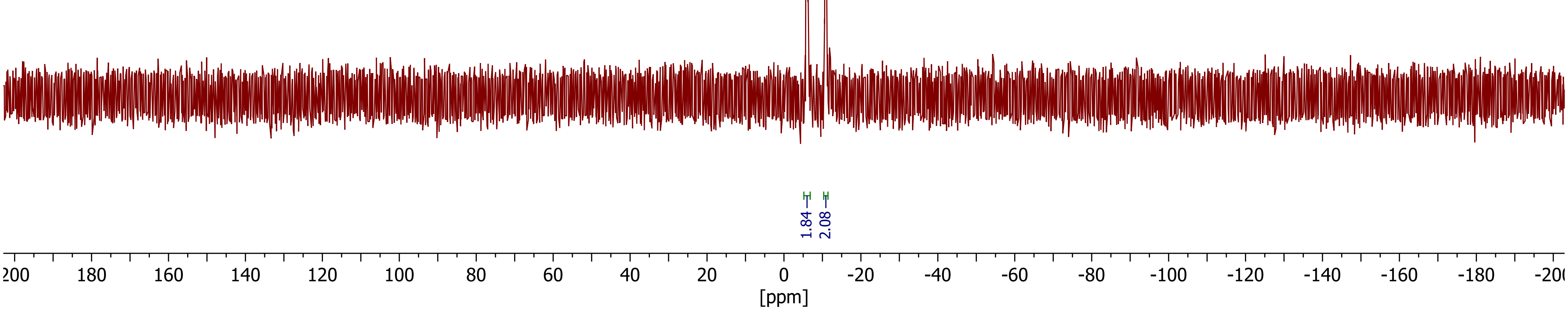

***Compound 24 (ppGpp - CC4), $^{1}H$ - NMR ($D_2O$, 400 MHz)***

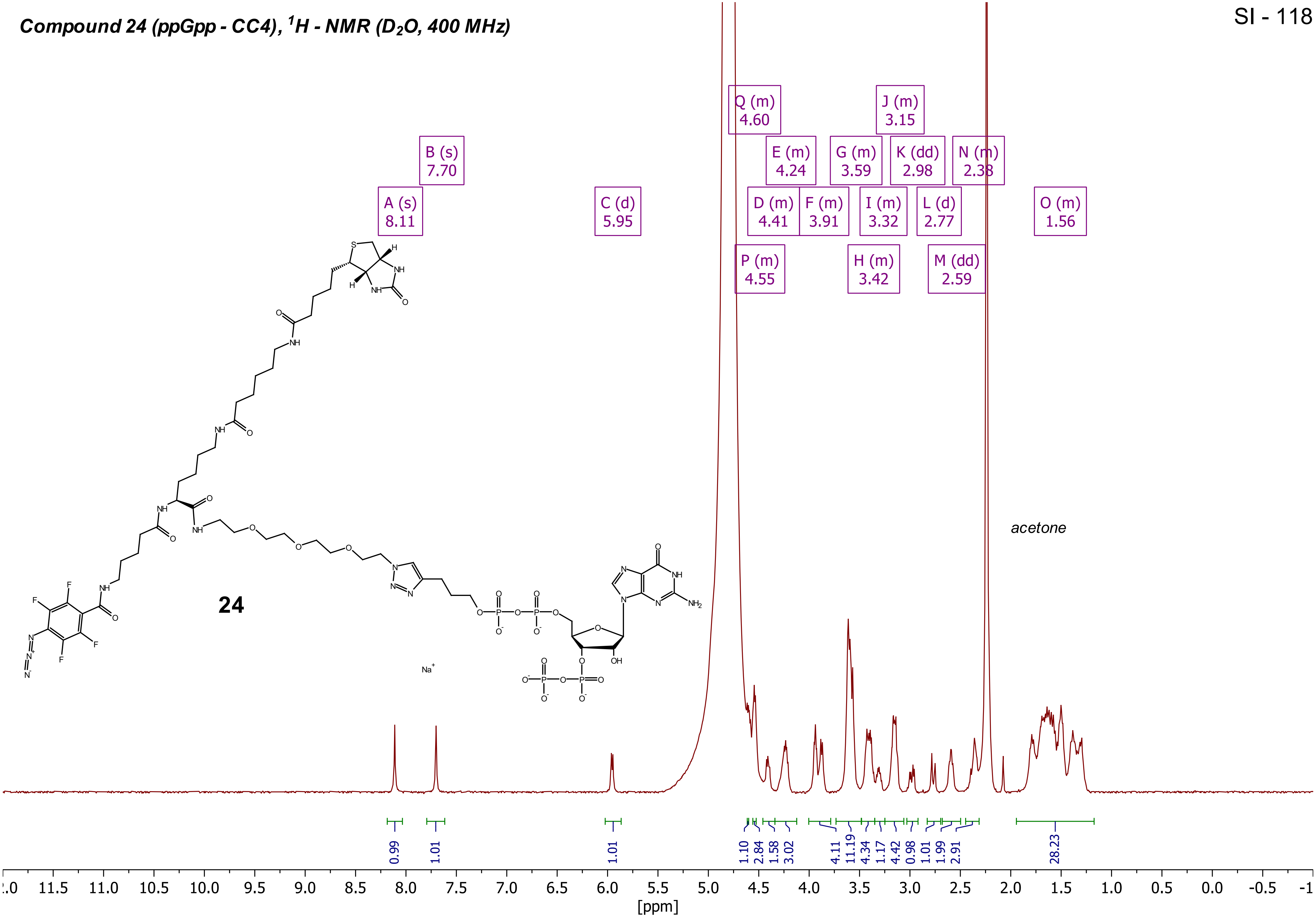

**Compound 24 (ppGpp - CC4), $^{19}F$ - NMR ($D_2O$, 377 MHz)**



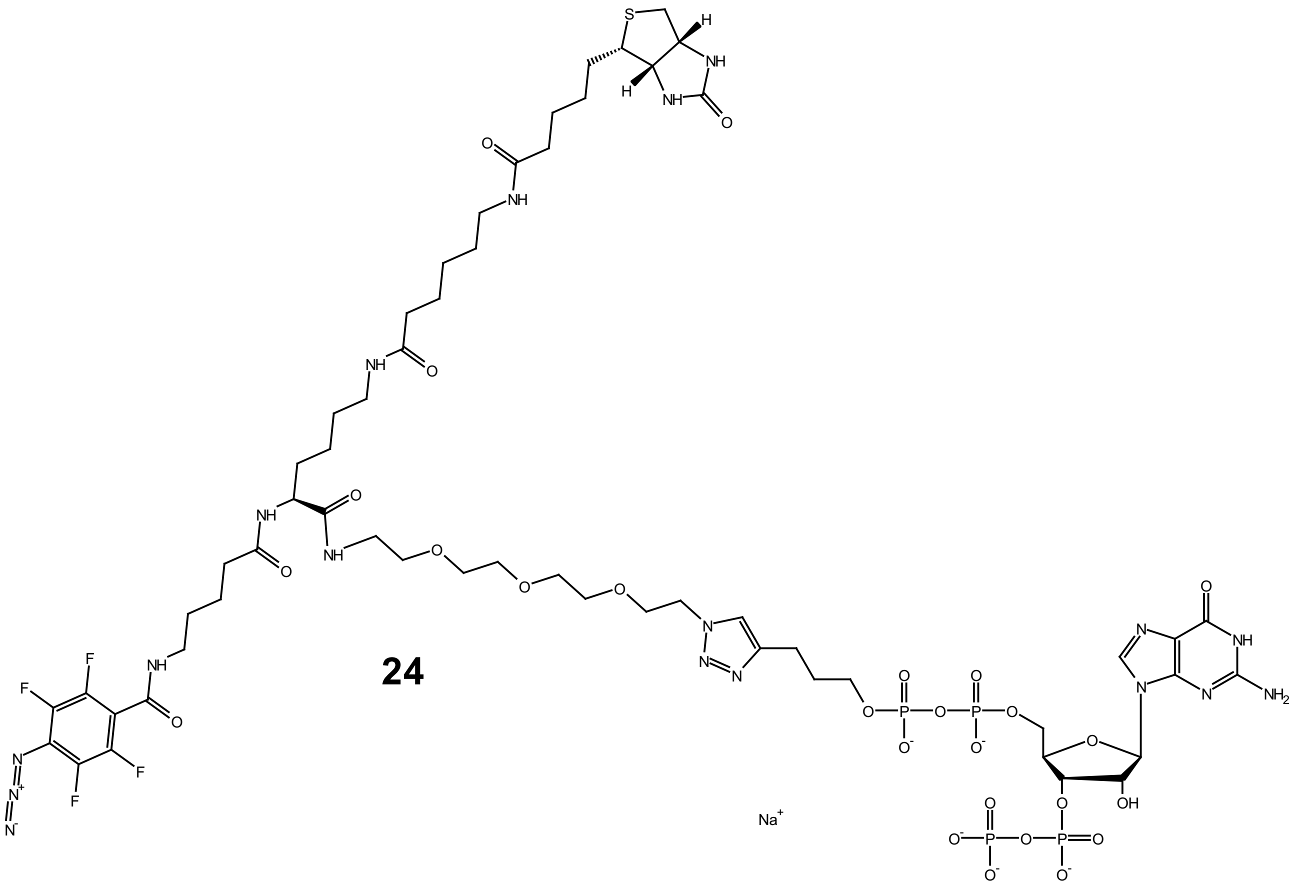

**24**

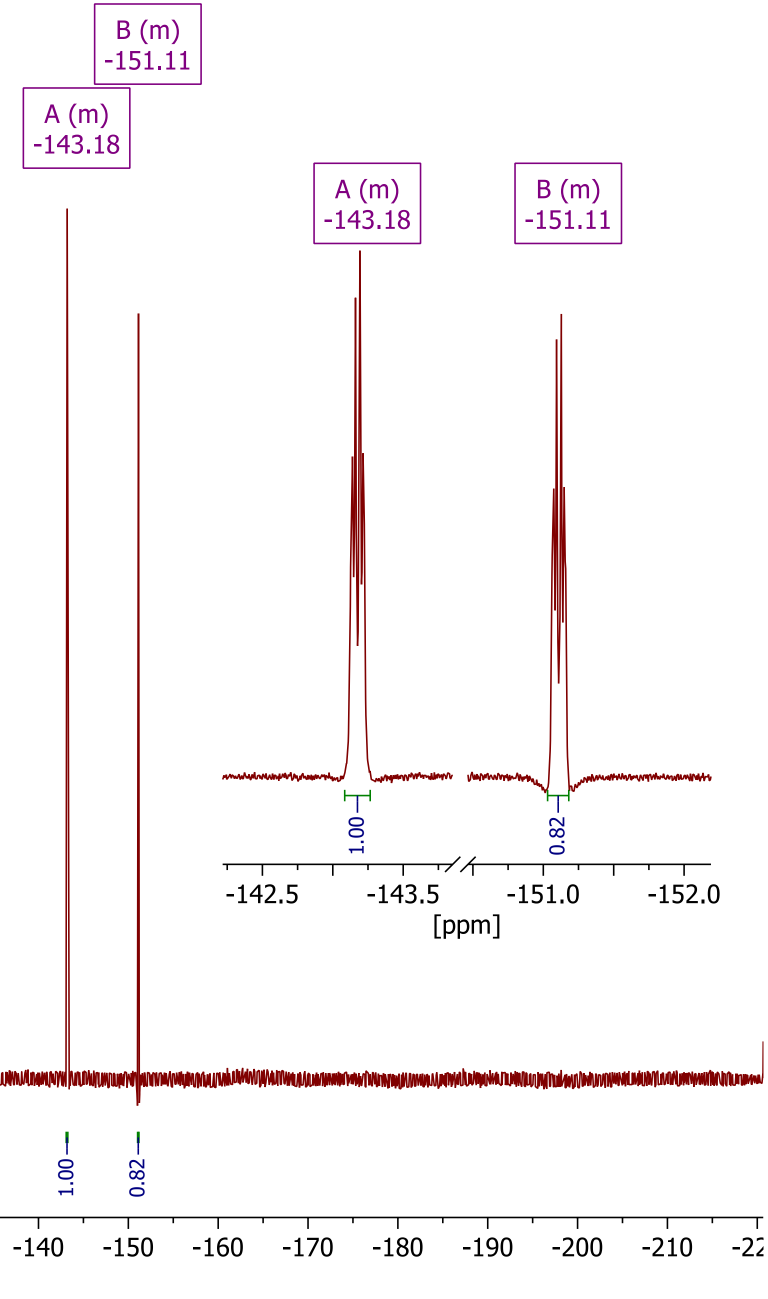

***Compound 24 (ppGpp - CC4), $^{31}P\{^1H\}$ - NMR ($D_2O$, 162 MHz)***

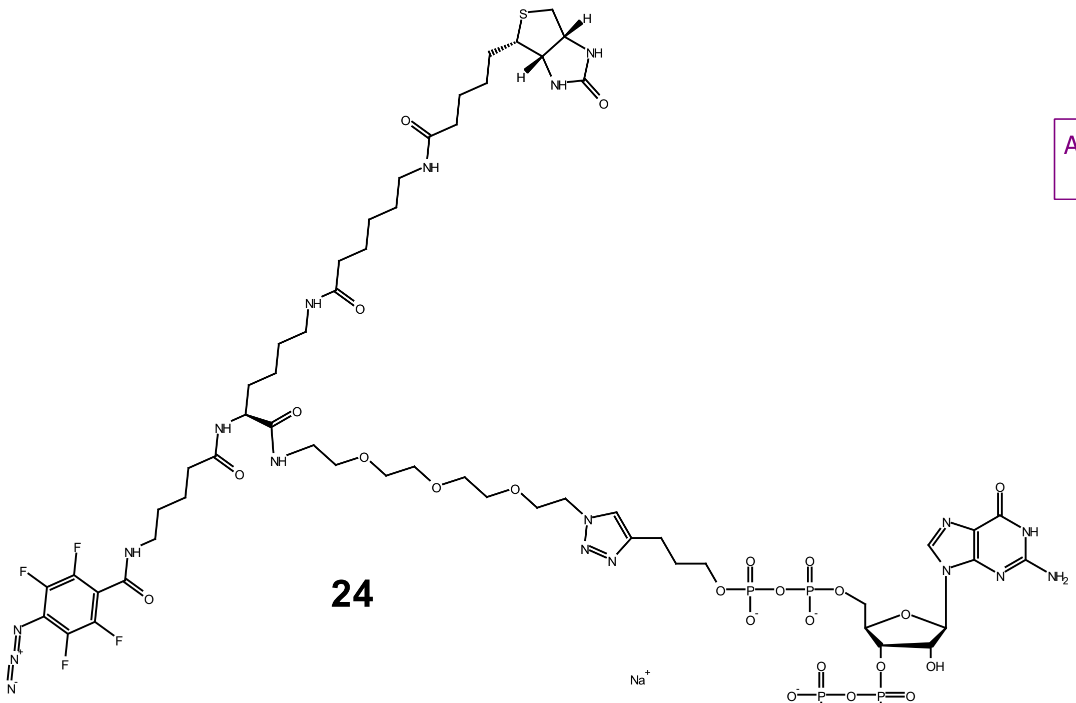

**24**

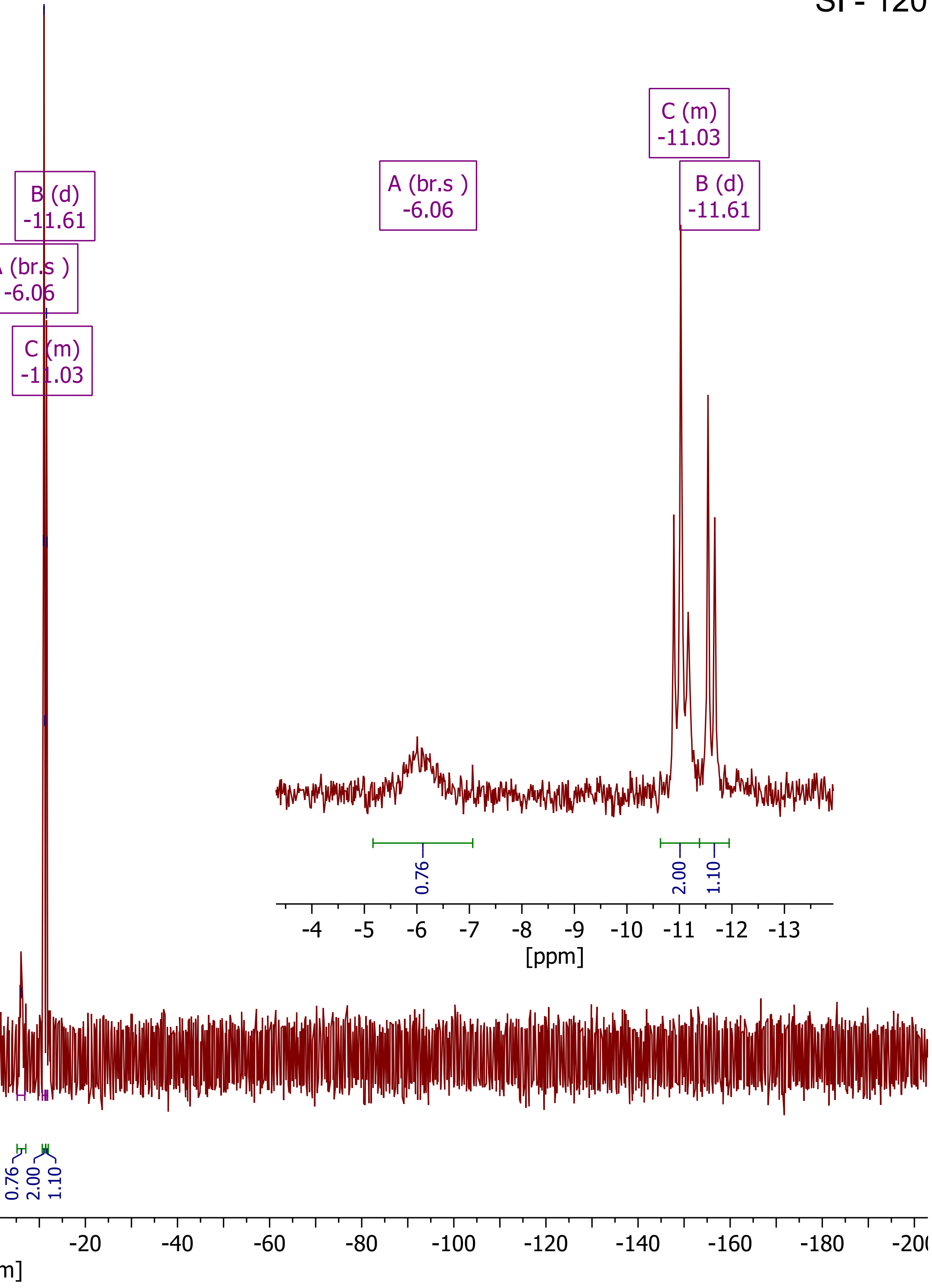

***Compound 25 (pGpp - CC5), $^1H$ - NMR ($D_2O$, 400 MHz)***

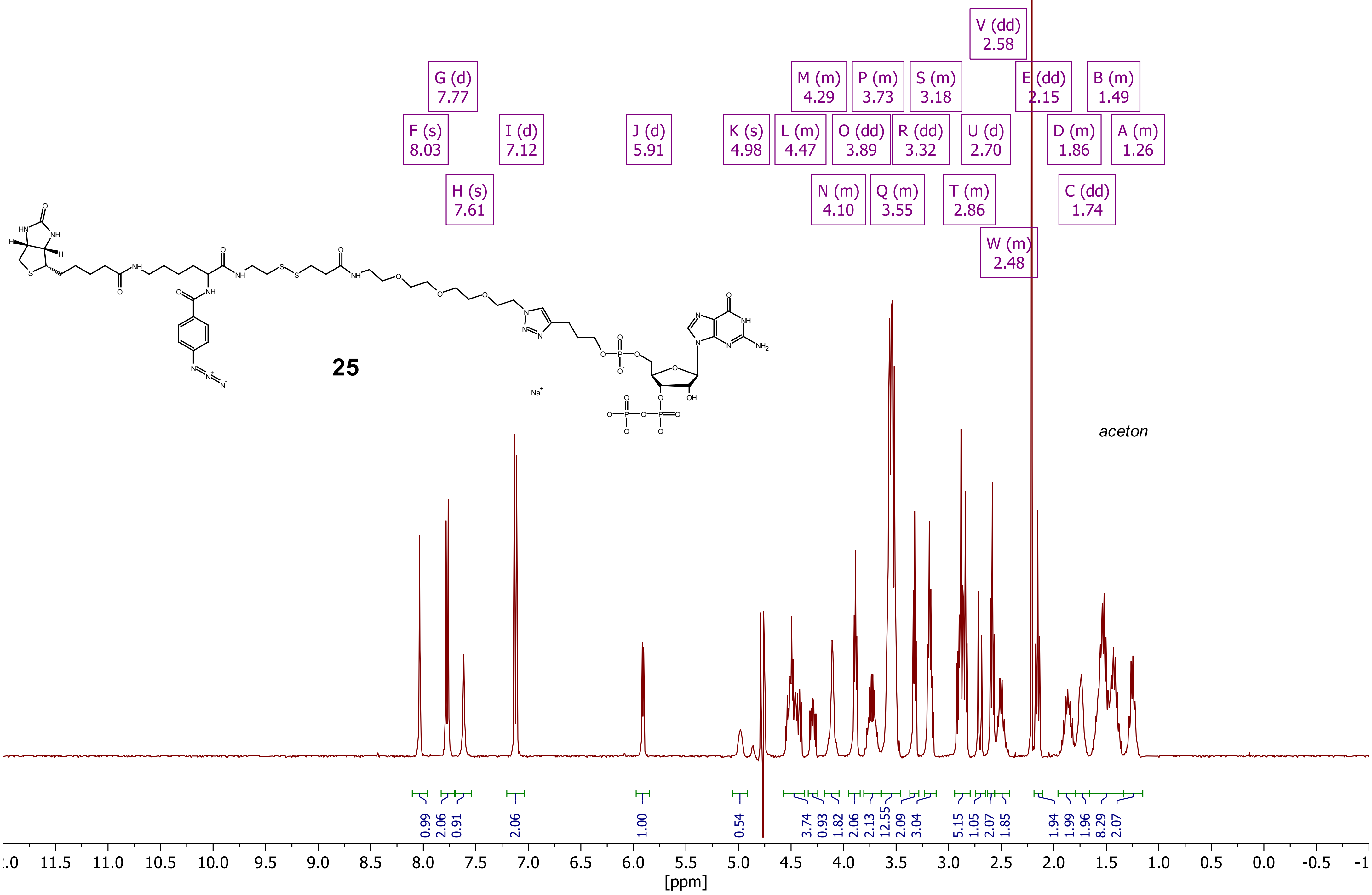

***Compound 25 (pGpp - CC5), $^{31}P\{^1H\}$ - NMR ($D_2O$, 162 MHz)***

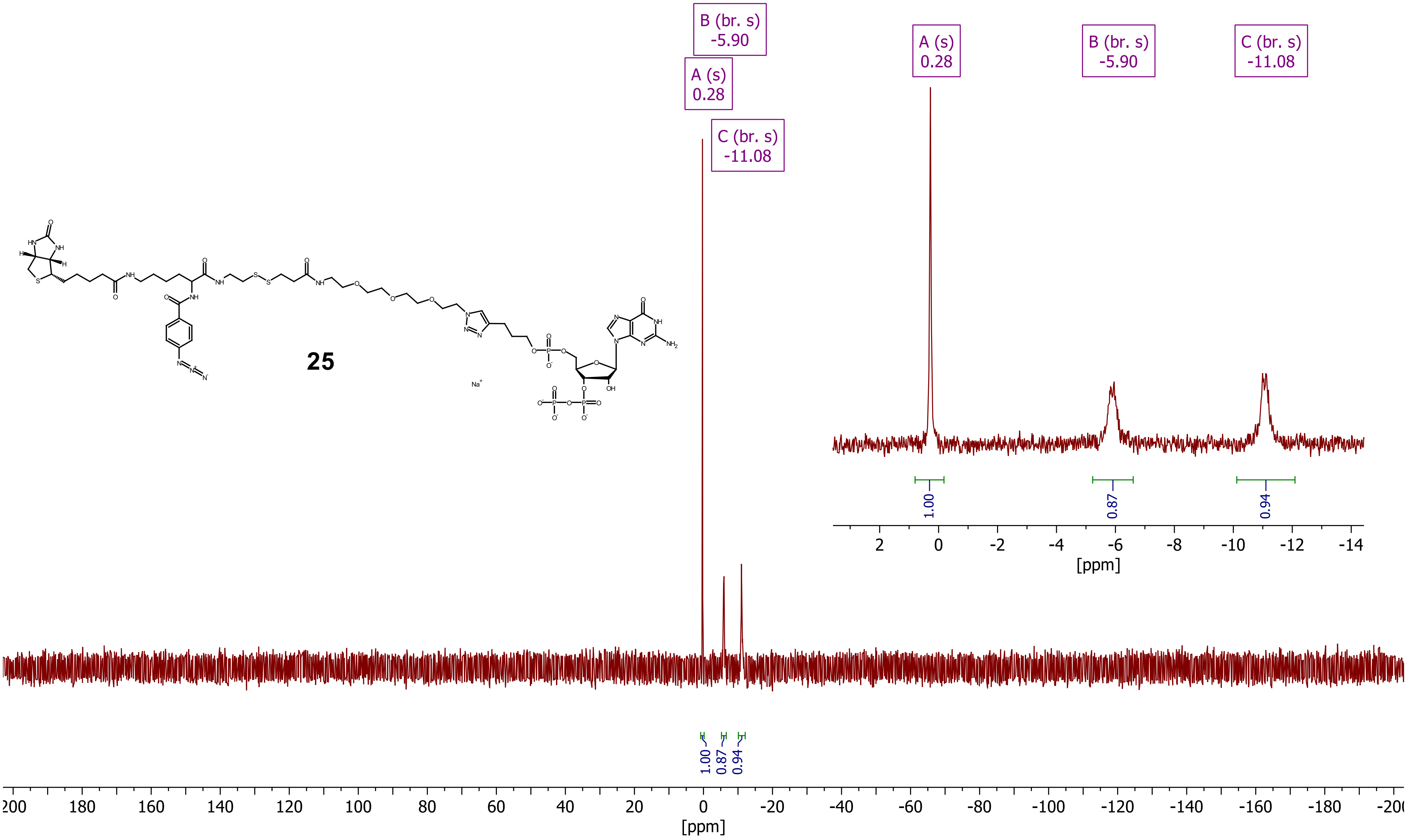


**25**

***Compound 26 (pppGpp - CC7), $^{1}H$ - NMR ($D_2O$, 400 MHz)***

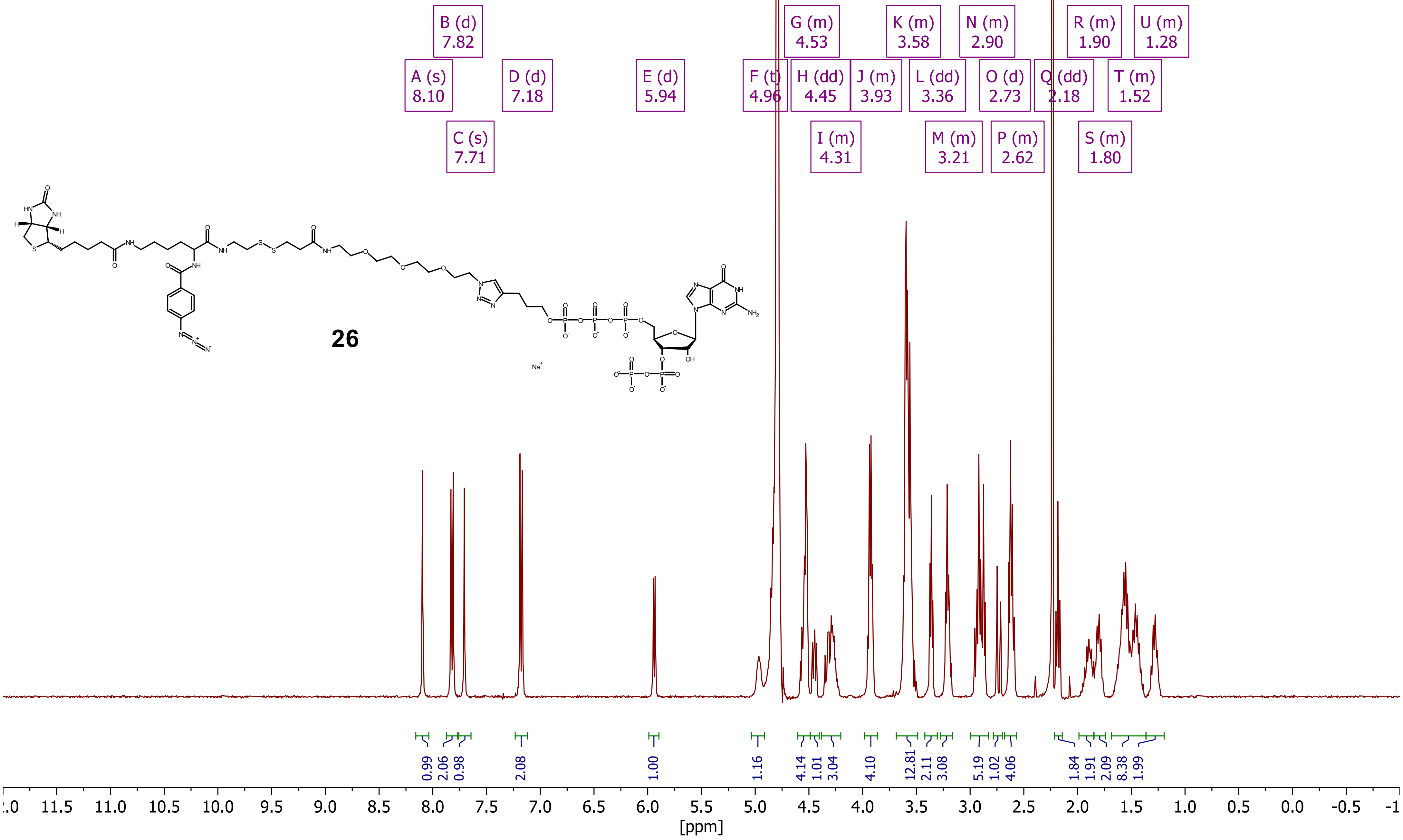

***Compound 26 (pppGpp - CC7), $^{31}P\{^1H\}$ - NMR ($D_2O$, 162 MHz)***

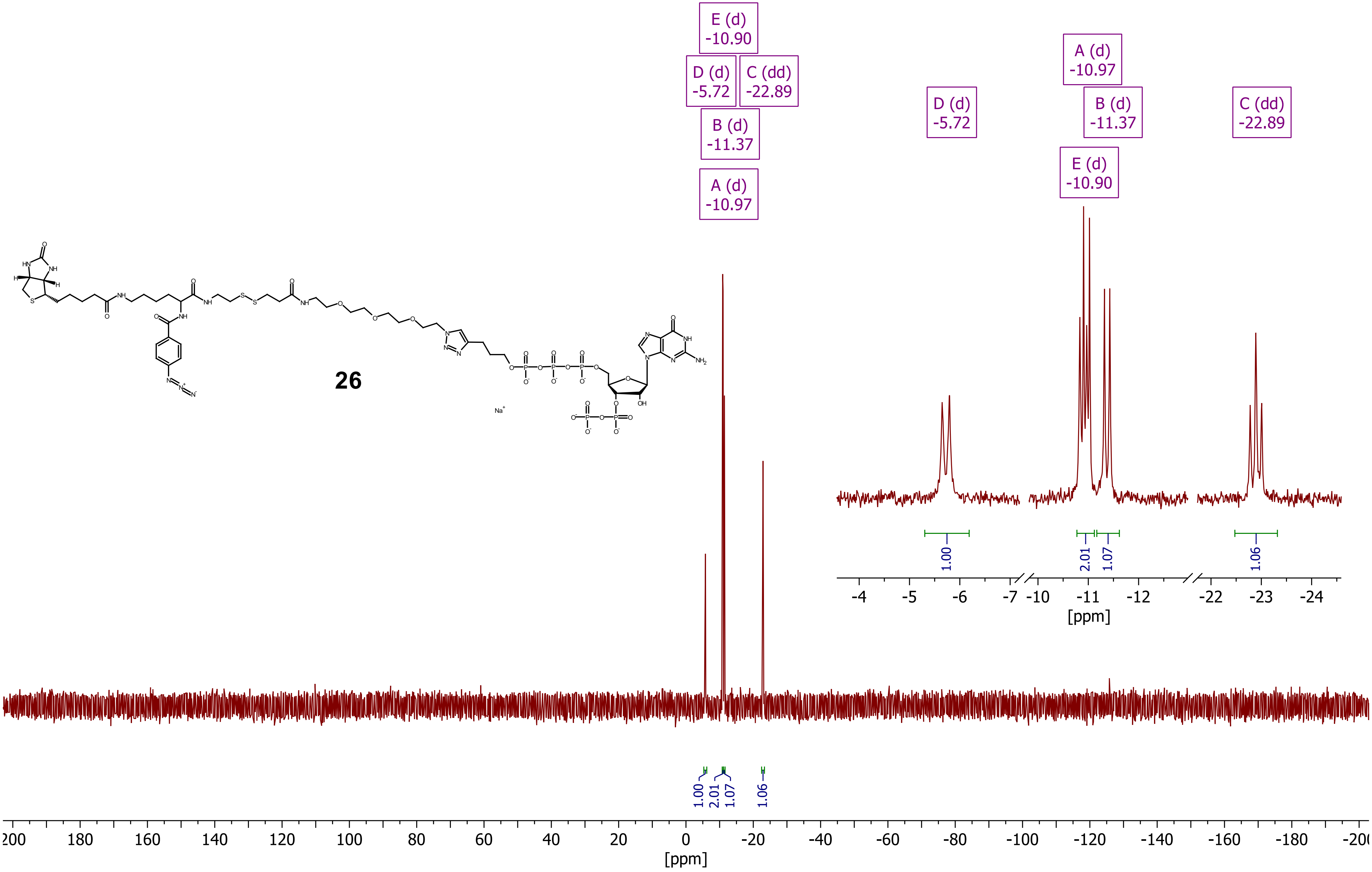

***Compound 27 (ppApp - CC8), ¹H - NMR ($D_2O$, 400 MHz)***

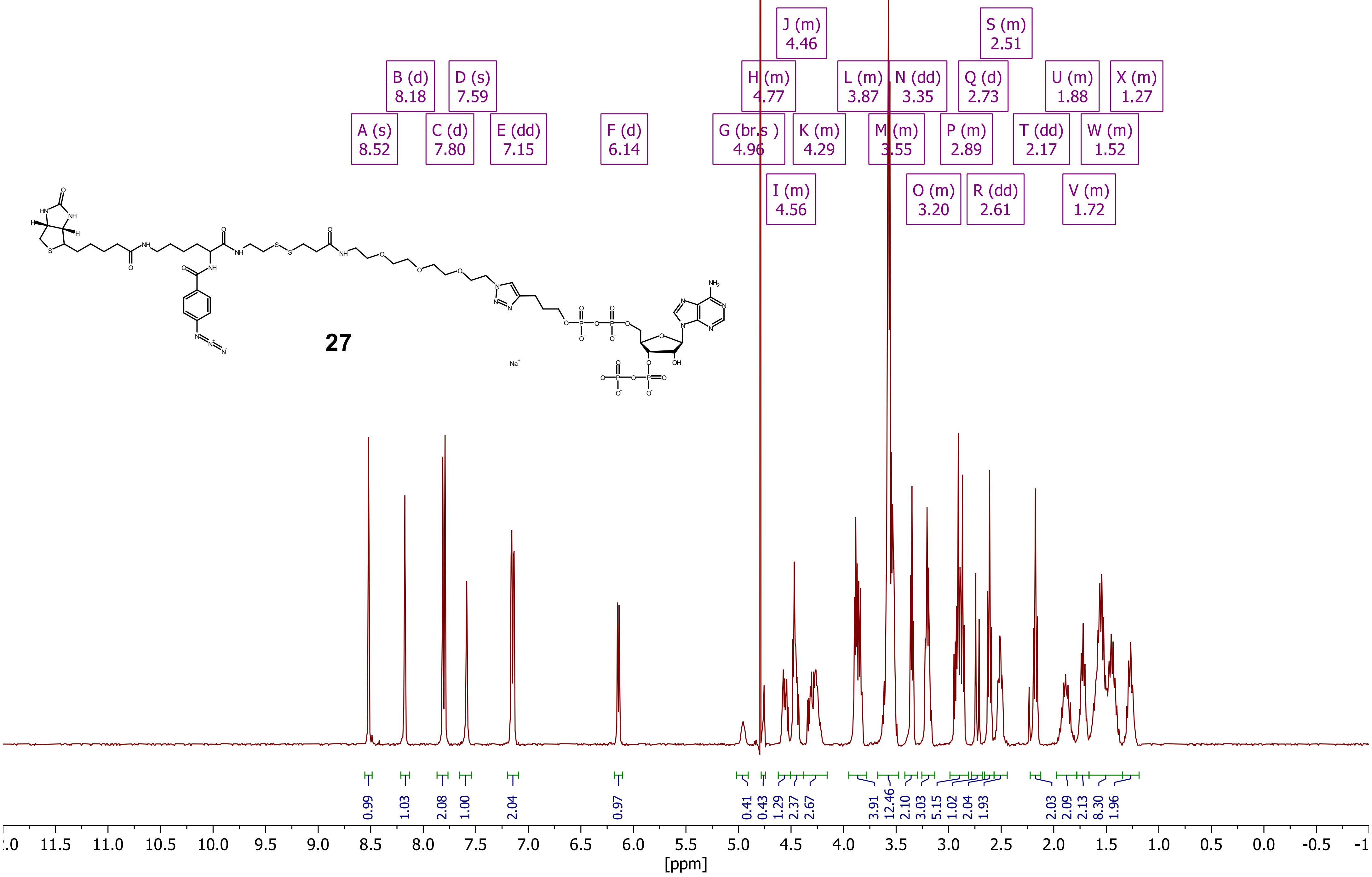

***Compound 27 (ppApp - CC8), $^{31}P\{^1H\}$ - NMR ($D_2O$, 162 MHz)***

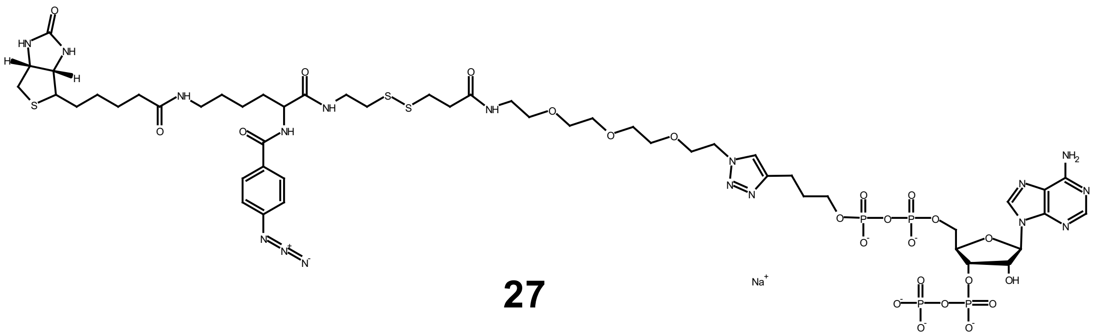

**27**

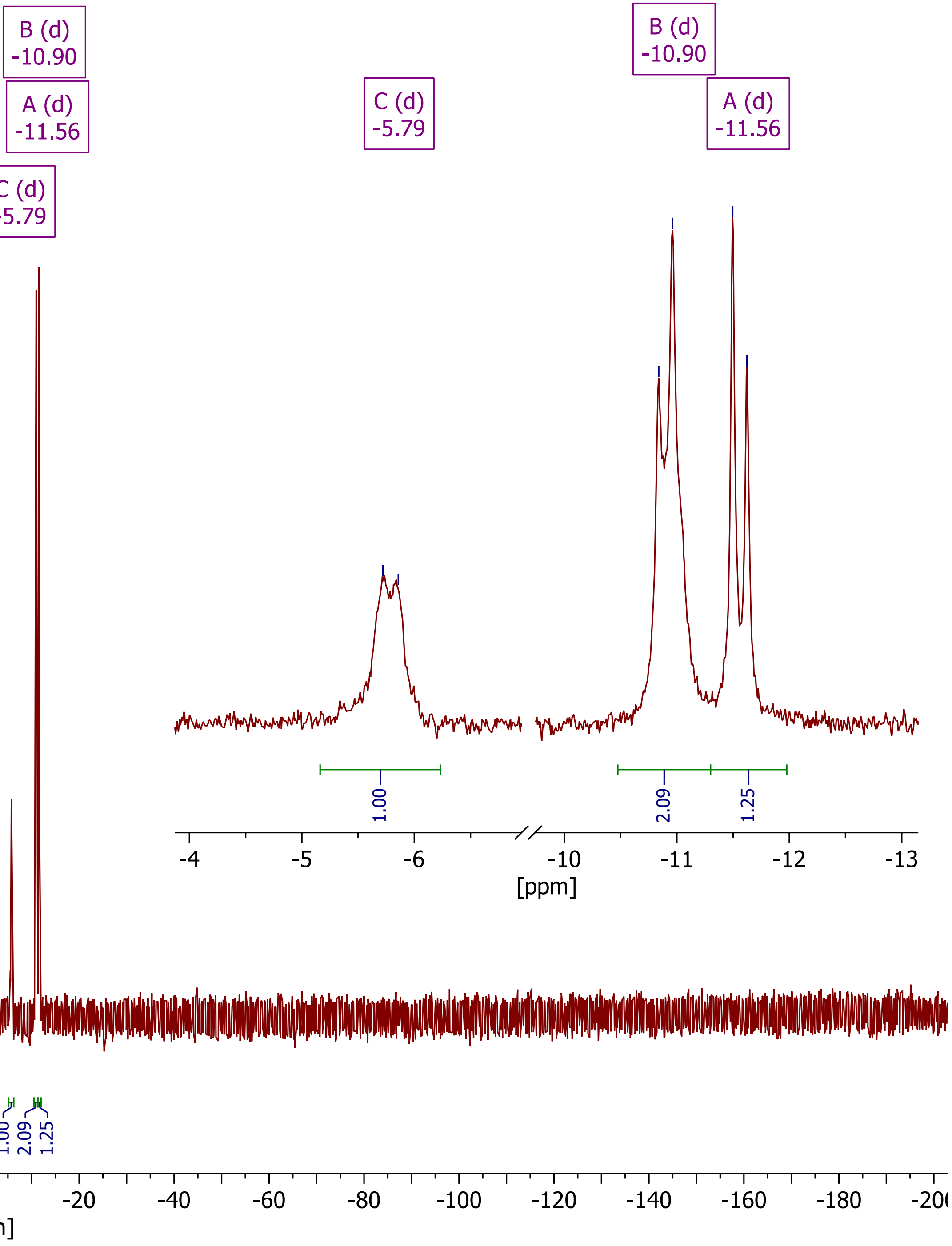

***Compound SI-13 (ppGp - CC6), $^{1}H$ - NMR ($D_2O$, 400 MHz)***

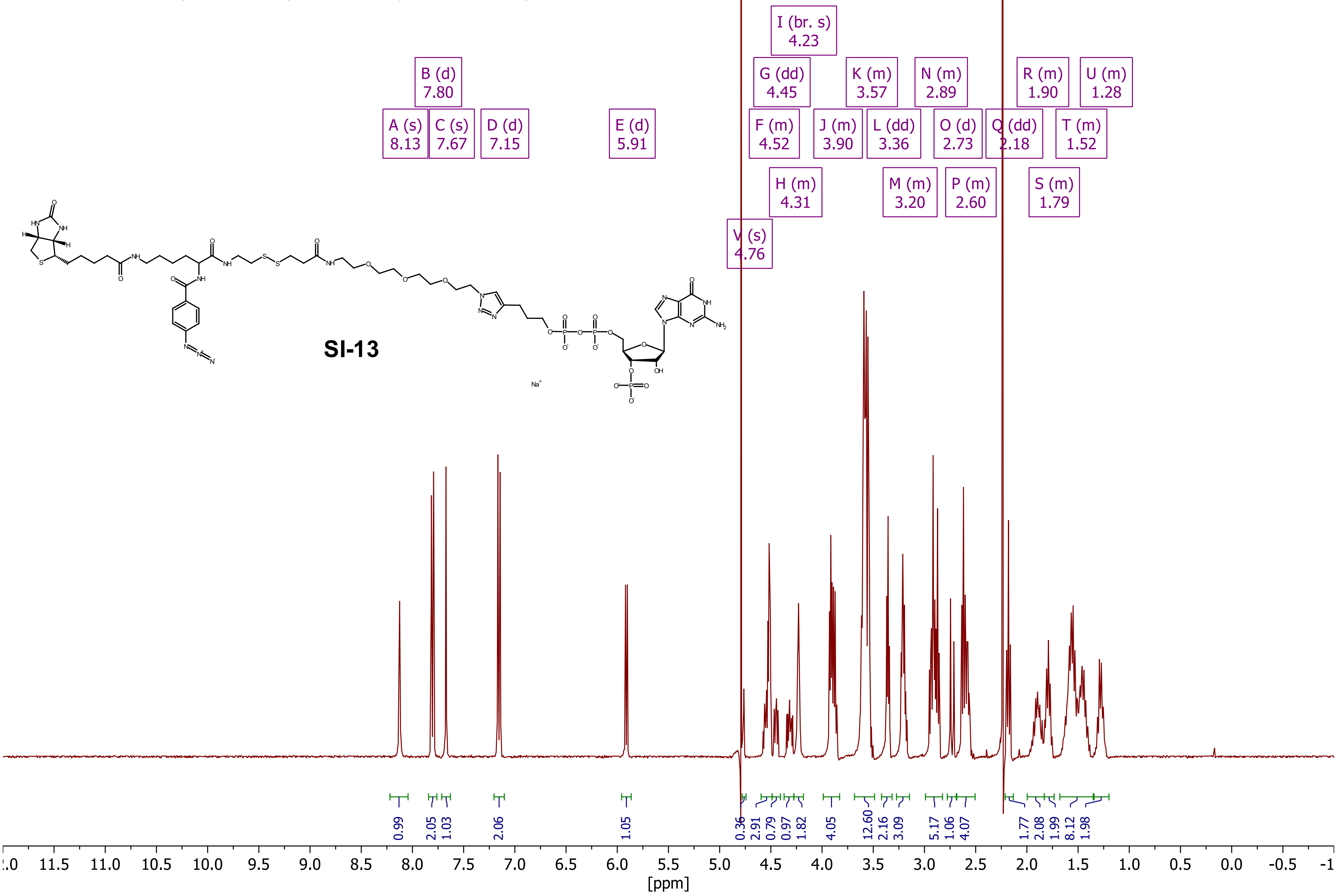

***Compound SI-13 (ppGp - CC6), $^{31}P\{^{1}H\}$ - NMR ($D_2O$, 162 MHz)***

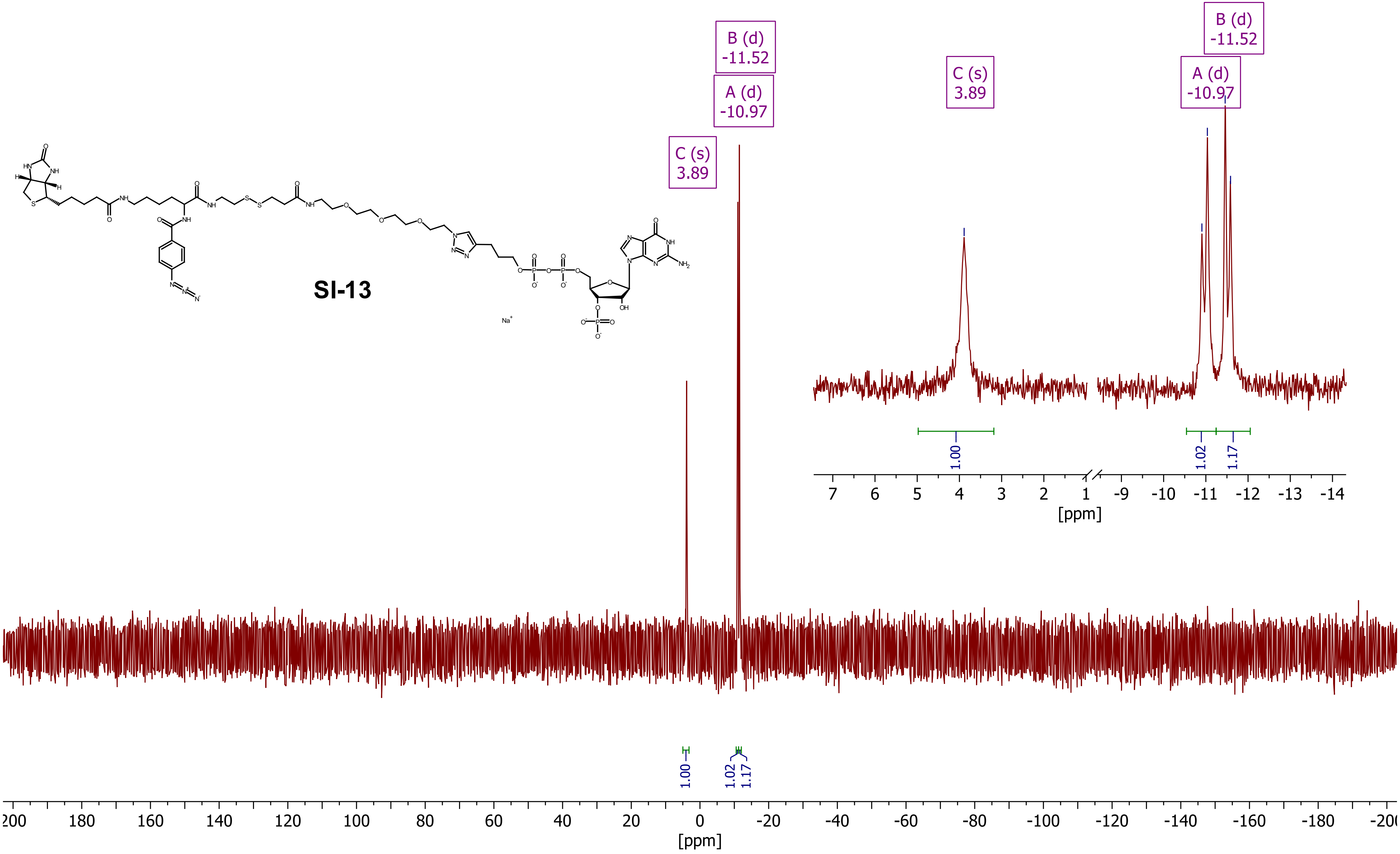

*Compound SI-5, $^{1}H$ - NMR (DMSO-$d_6$, 400 MHz)*

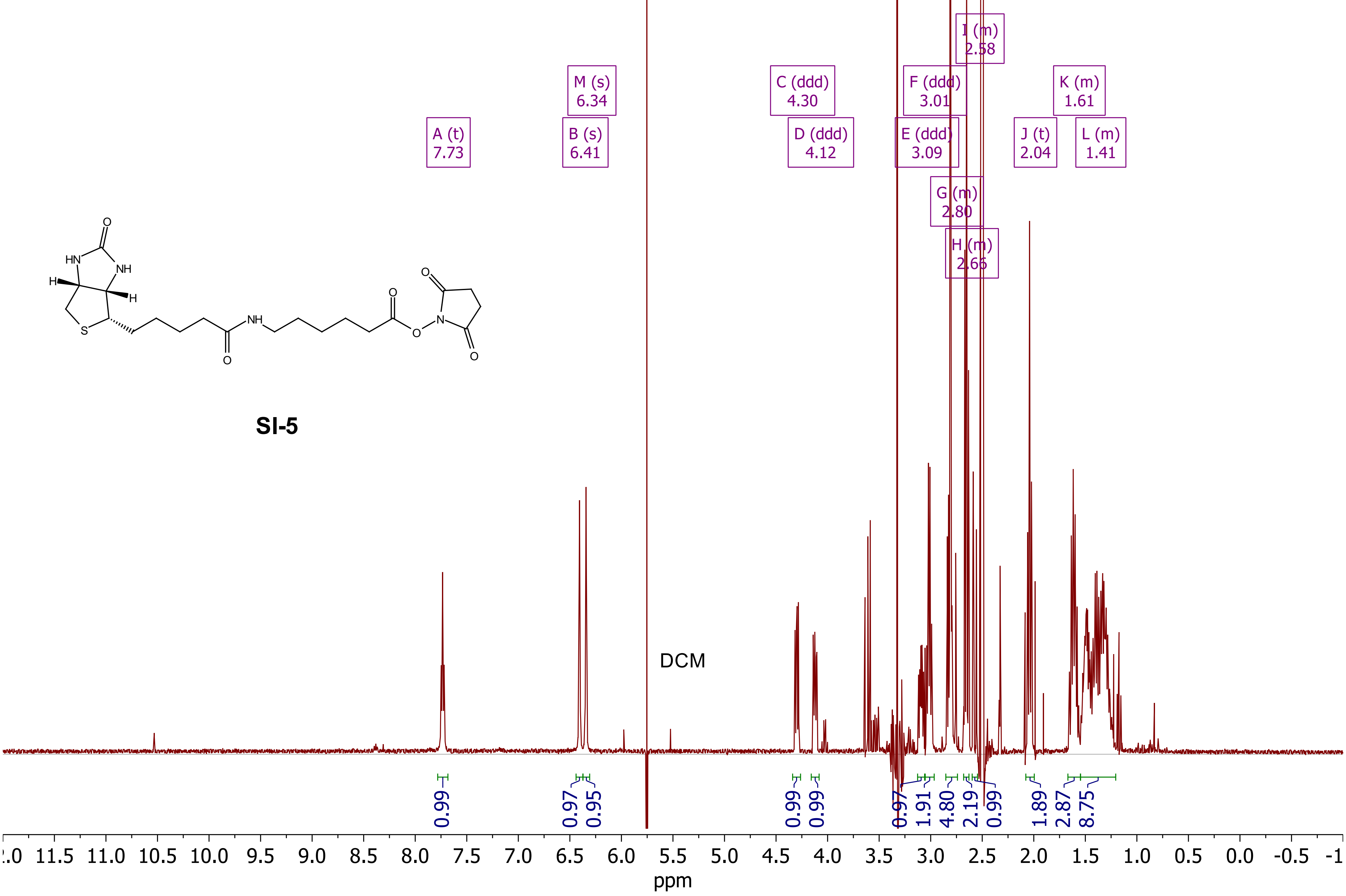

***Compound SI-5, $^{13}C\{^{1}H\}$ - NMR (DMSO-$d_6$, 101 MHz)***

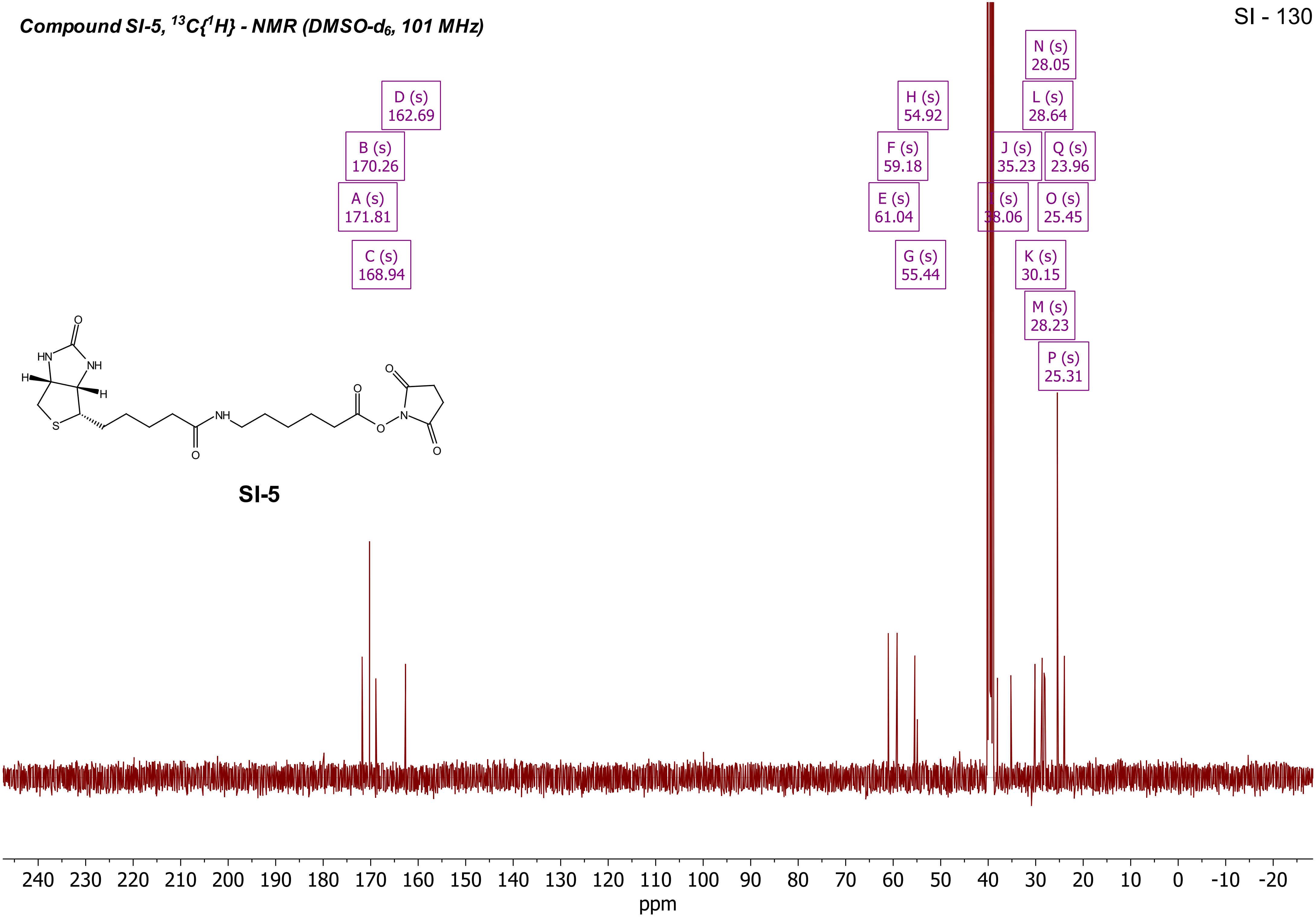


**SI-5**

*Compound SI-6, 1H - NMR ($D_2O$, 400 MHz)*

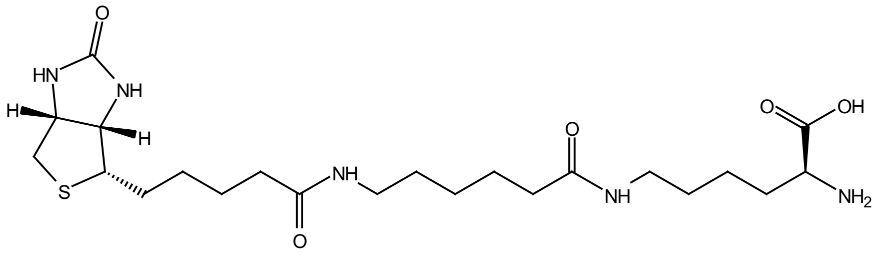

**SI-6**

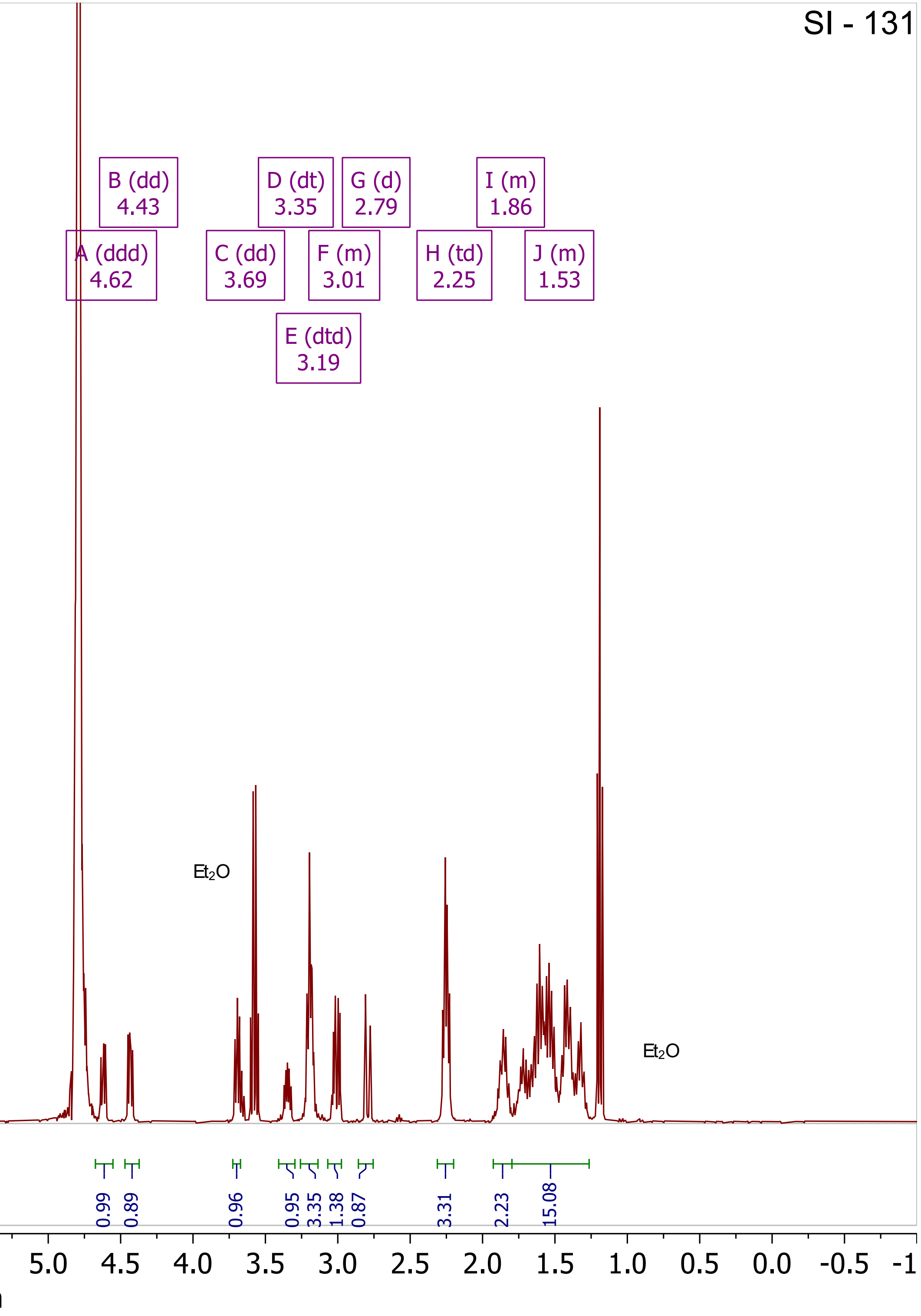

***Compound SI-6, $^{13}C\{^1H\}$ - NMR ($D_2O$, 101 MHz)***

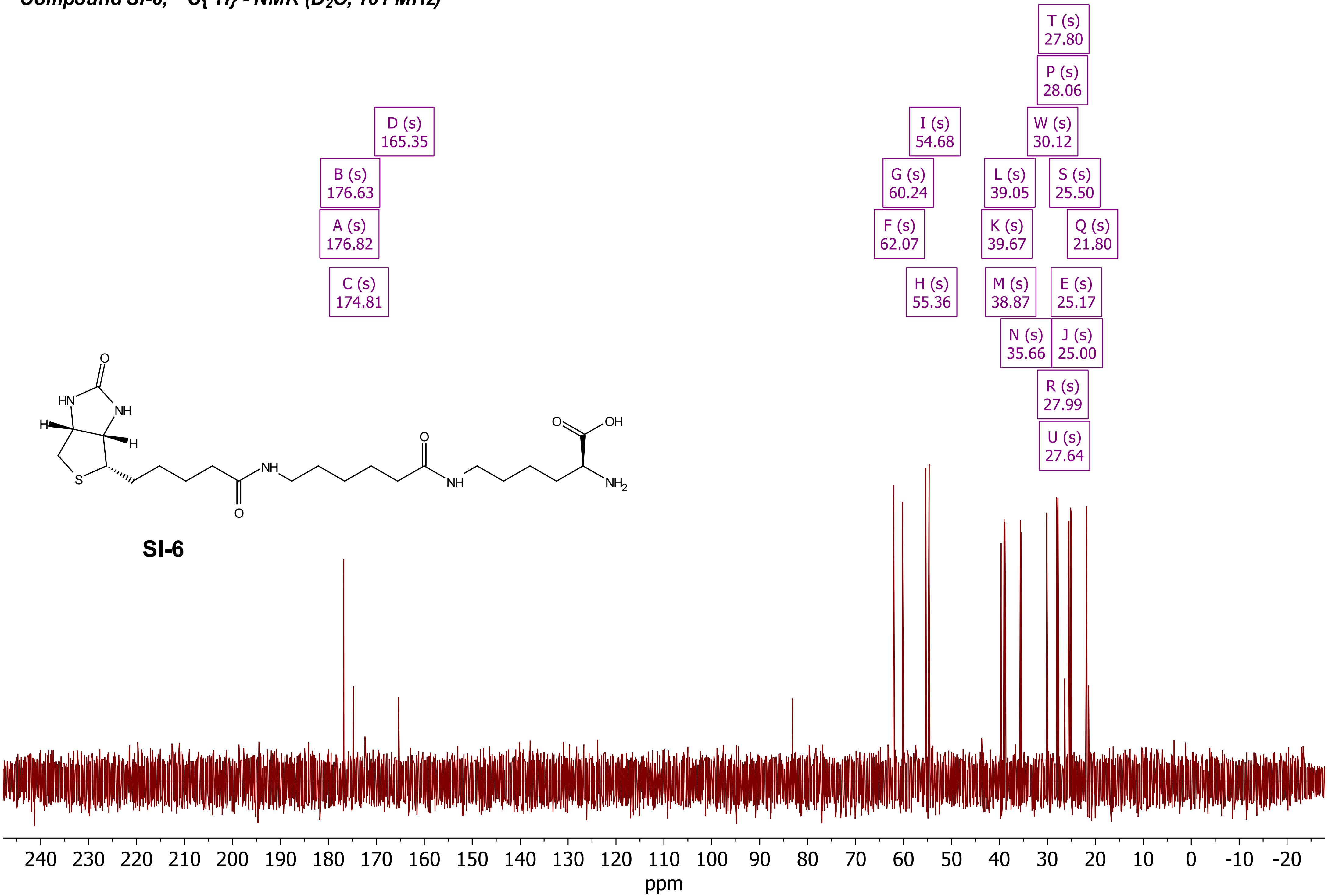

***Compound SI-10, [1]H - NMR (MeOD-d$_4$, 400 MHz)***

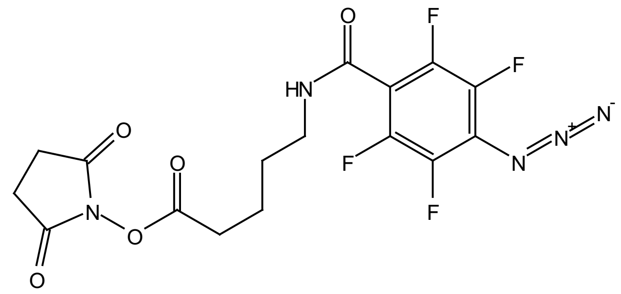

**SI-10**

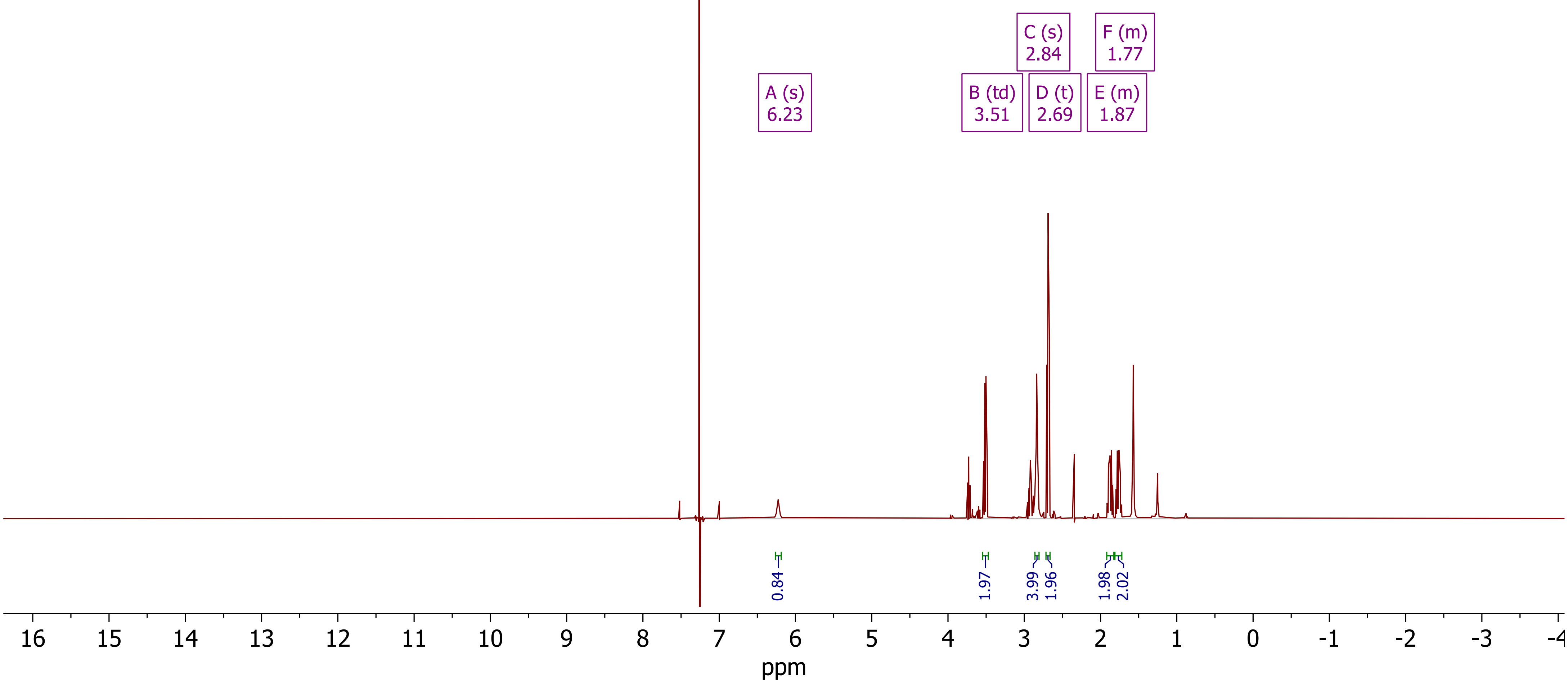

***Compound SI-10, $^{13}C[^{1}H]$ - NMR (MeOD-$d_4$, 101 MHz)***

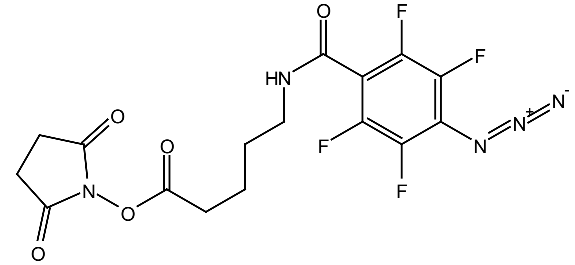


**SI-10**

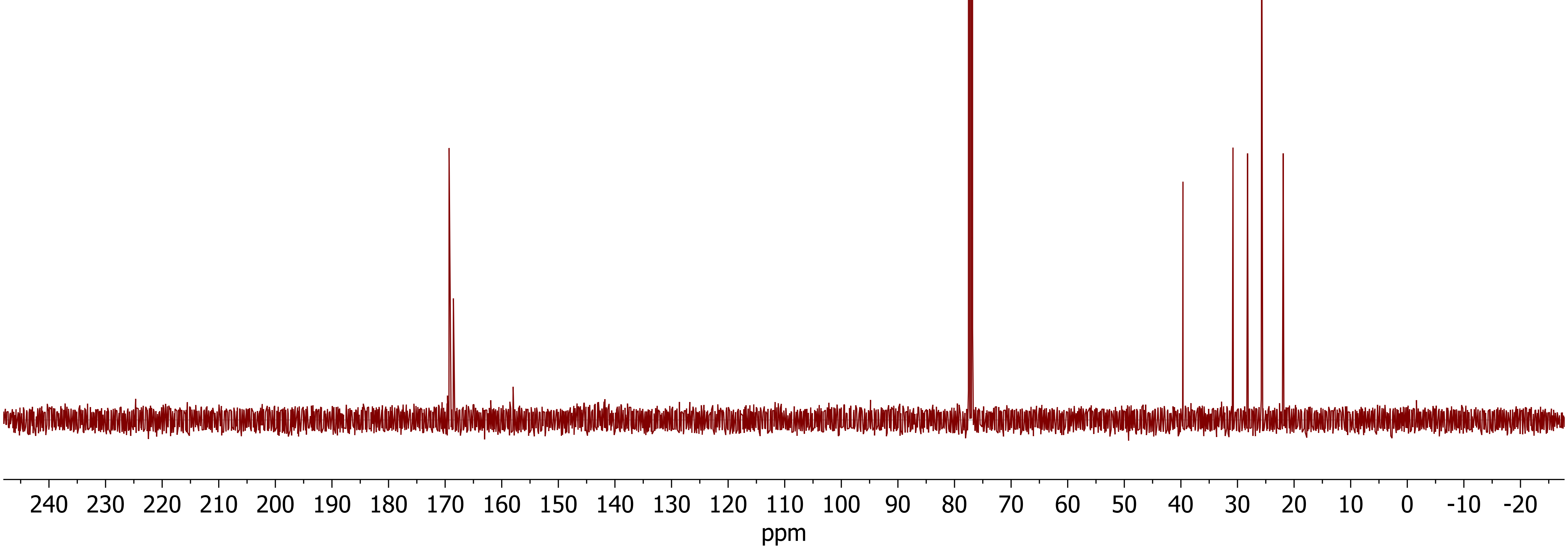

***Compound SI-10, $^{19}$F - NMR (MeOD-$d_4$, 377 MHz)***

**SI-10**

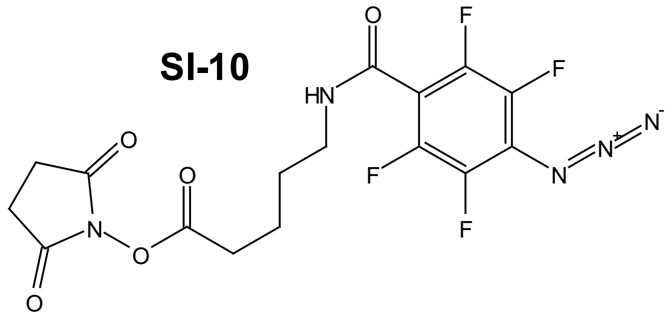

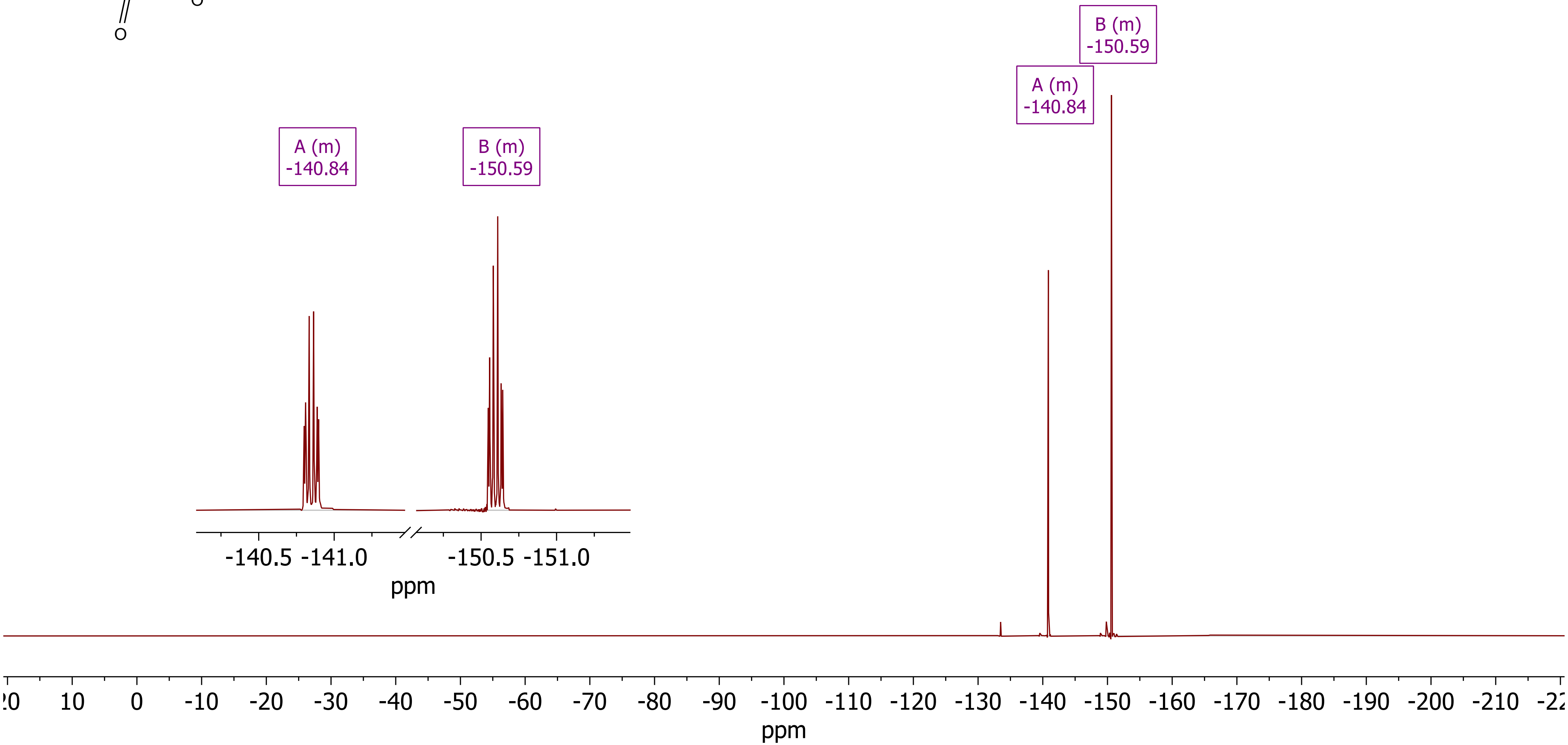

***Compound SI-11, 1H - NMR (DMSO-d6, 400 MHz)***

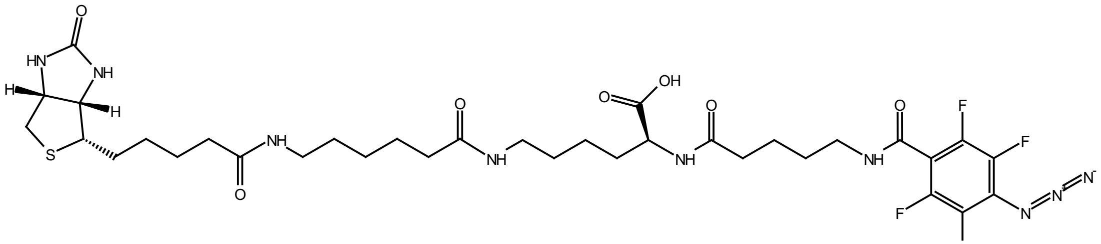

**SI-11**

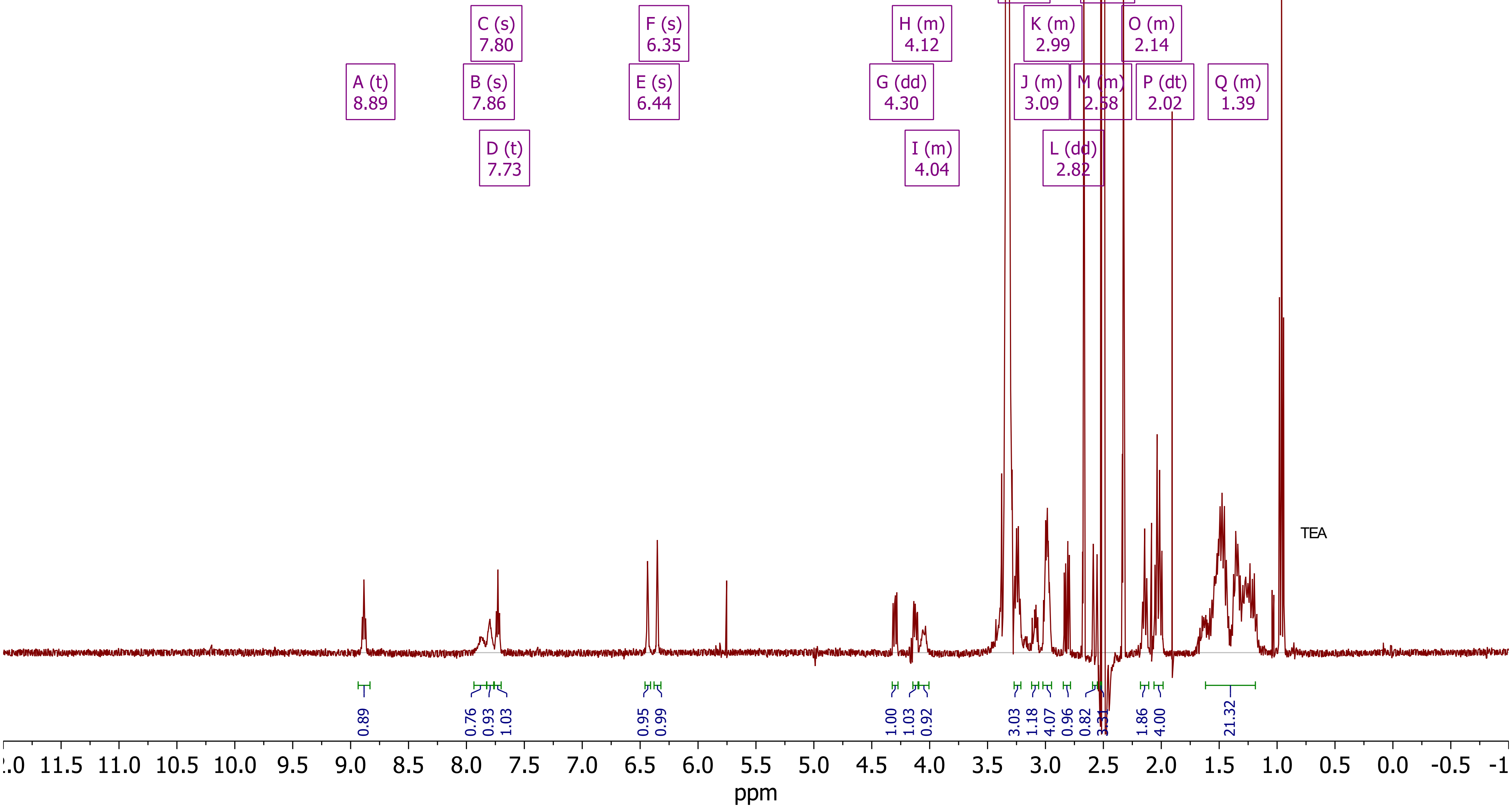

*Compound SI-11, ^19^F - NMR (DMSO-d$_6$, 377 MHz)*



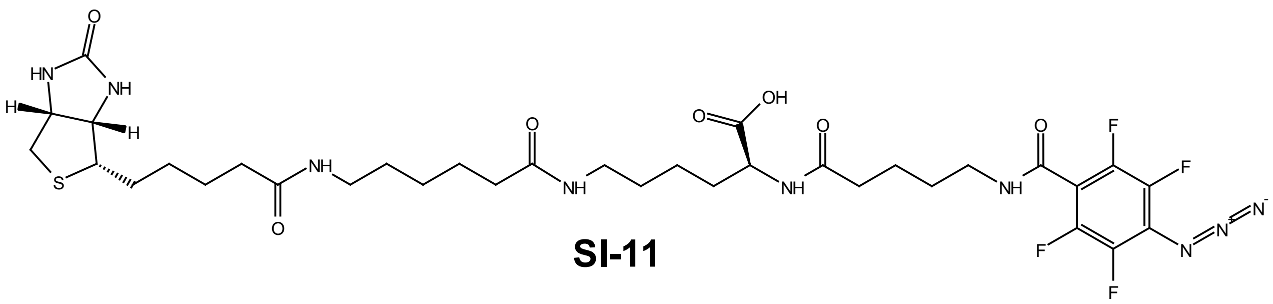

SI-11

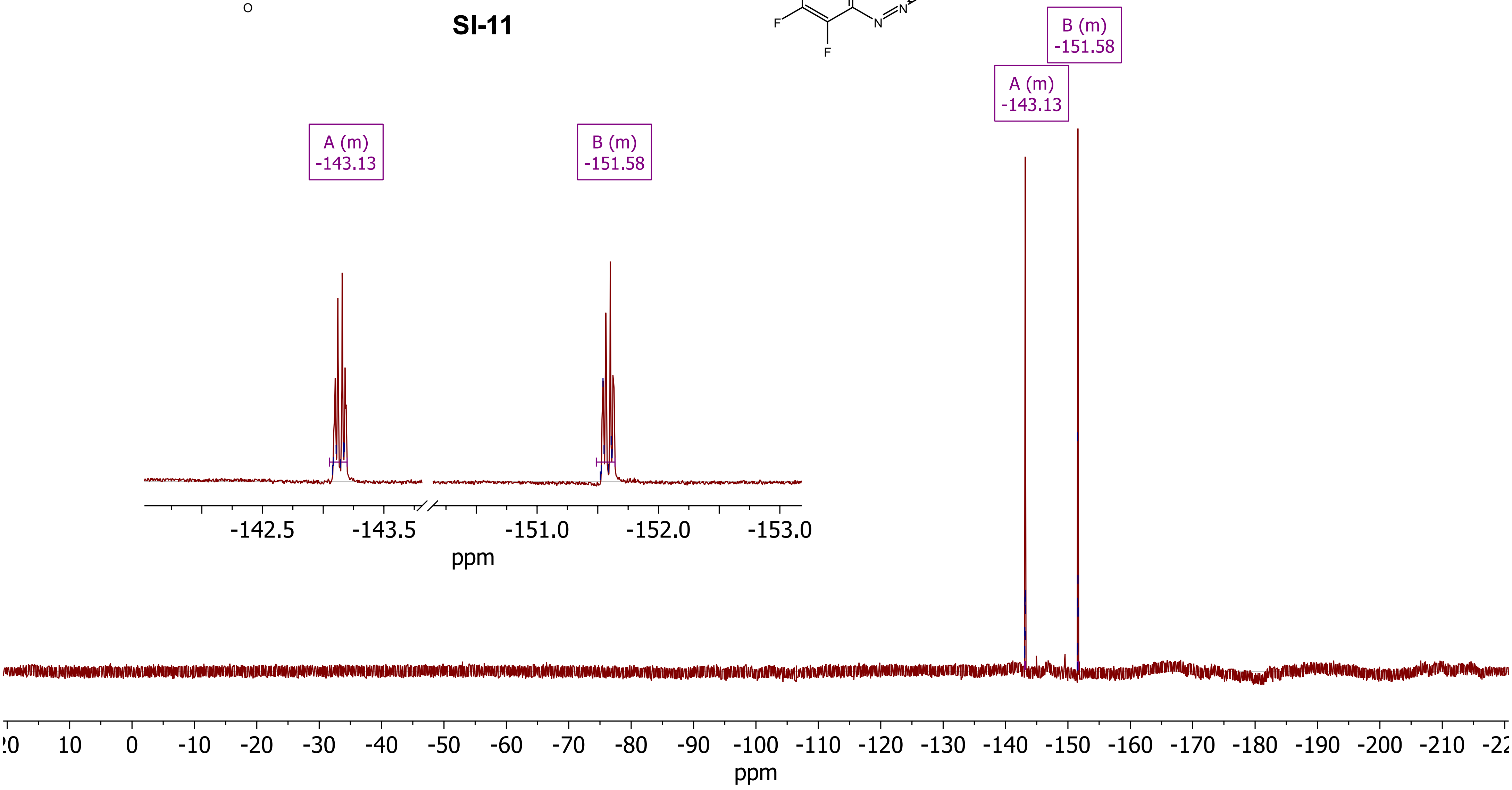

### *8. MS – spectra*

***(sorted according to molecule numbering)***

### *HRMS (ESI) Analysis of compound 5: HMDA – pGp*

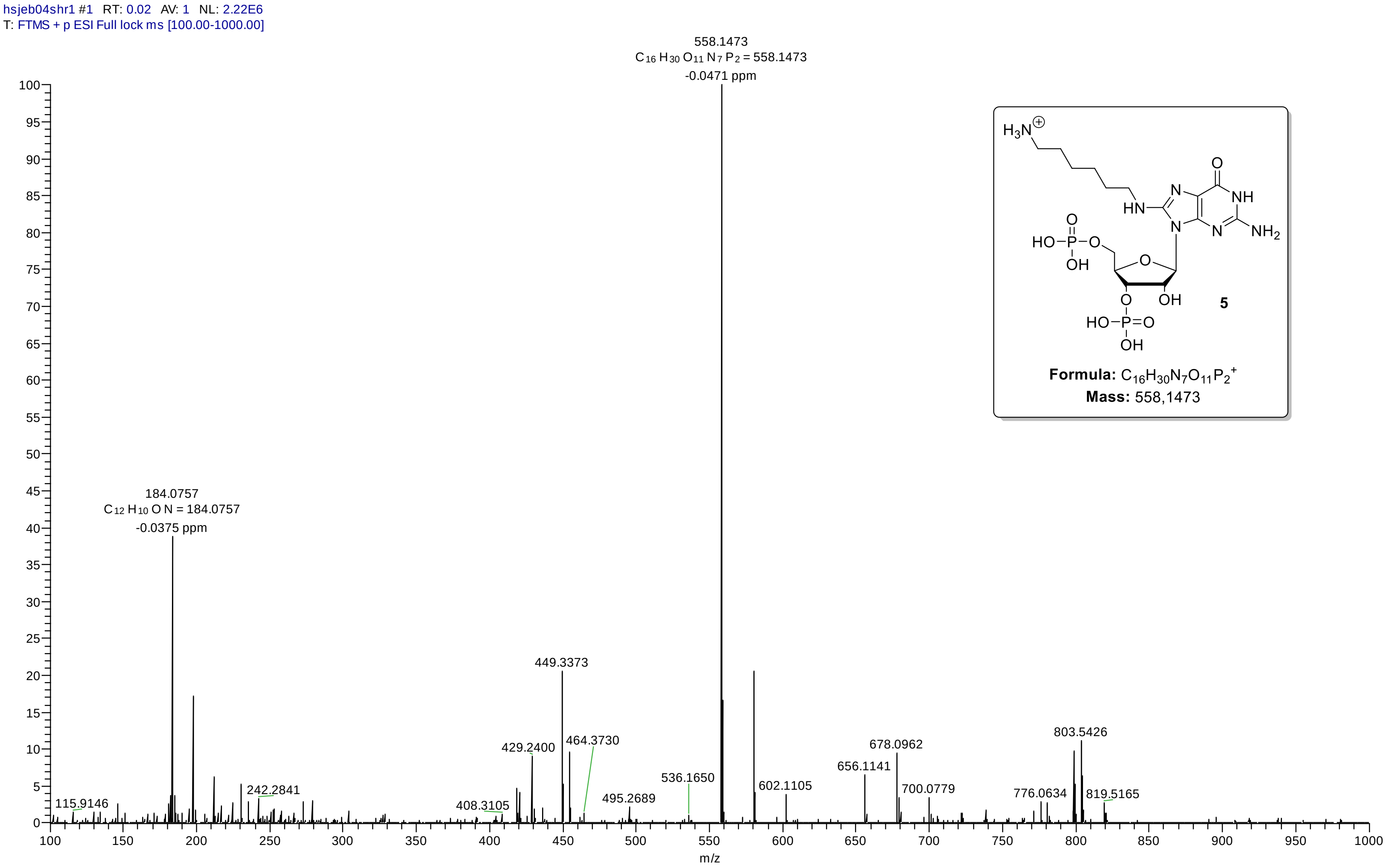

#### *HRMS (ESI) Analysis of compound 6: Fmoc-Glycyl-HMDA – pGp*

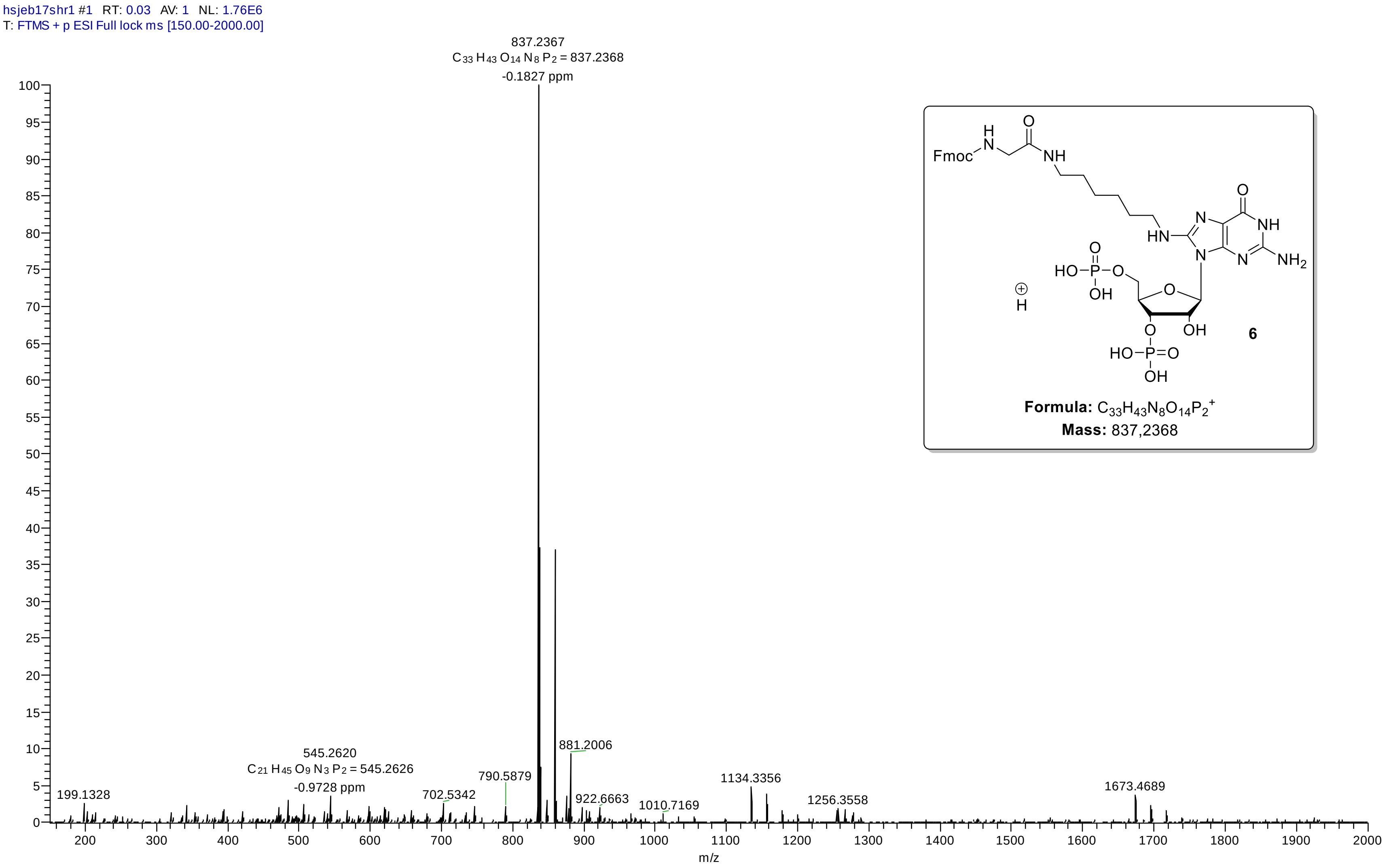

#### *HRMS (ESI) Analysis of compound 7: Glycyl-HMDA – ppGpp*

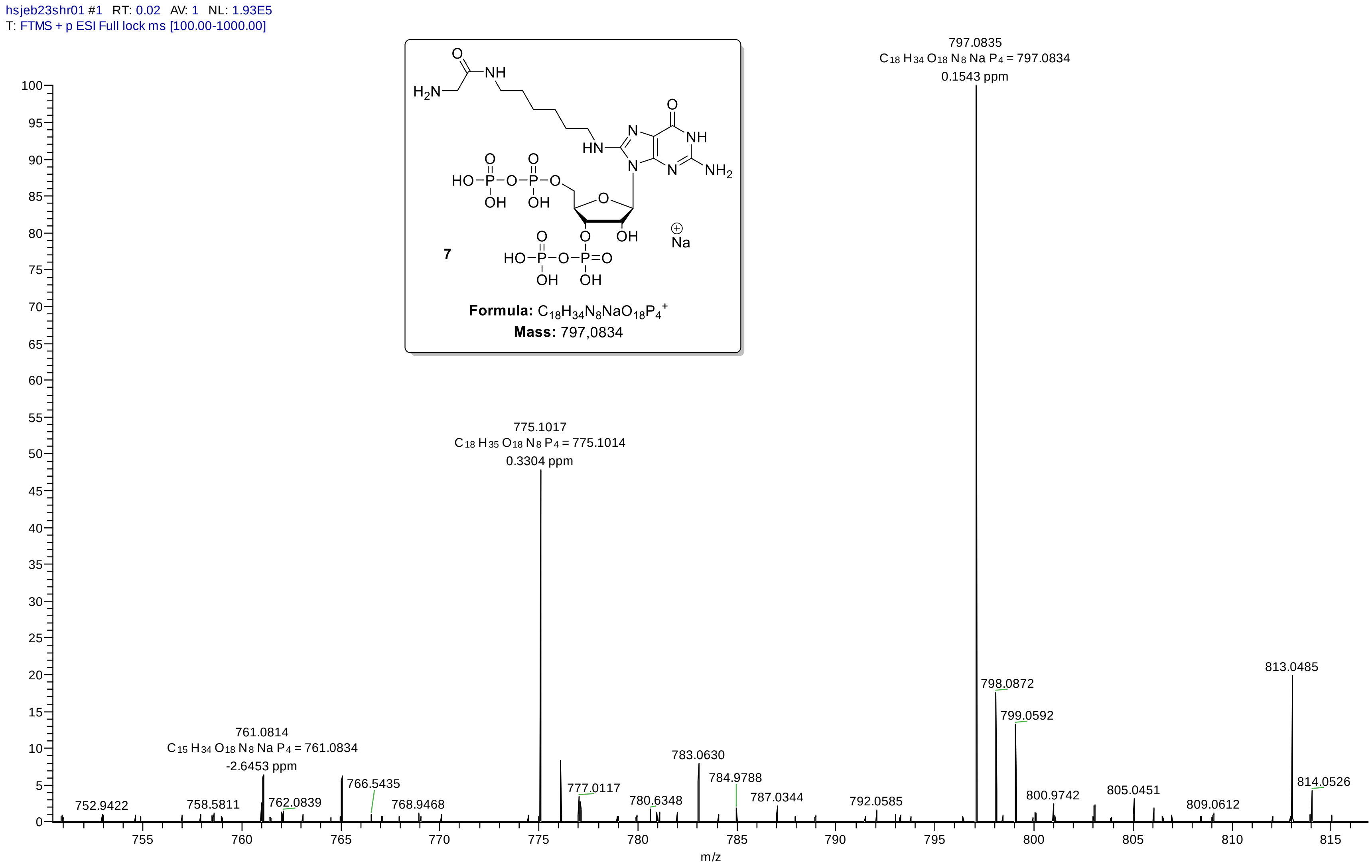

#### HRMS (ESI) Analysis of compound 9: pentynyl - pppGp

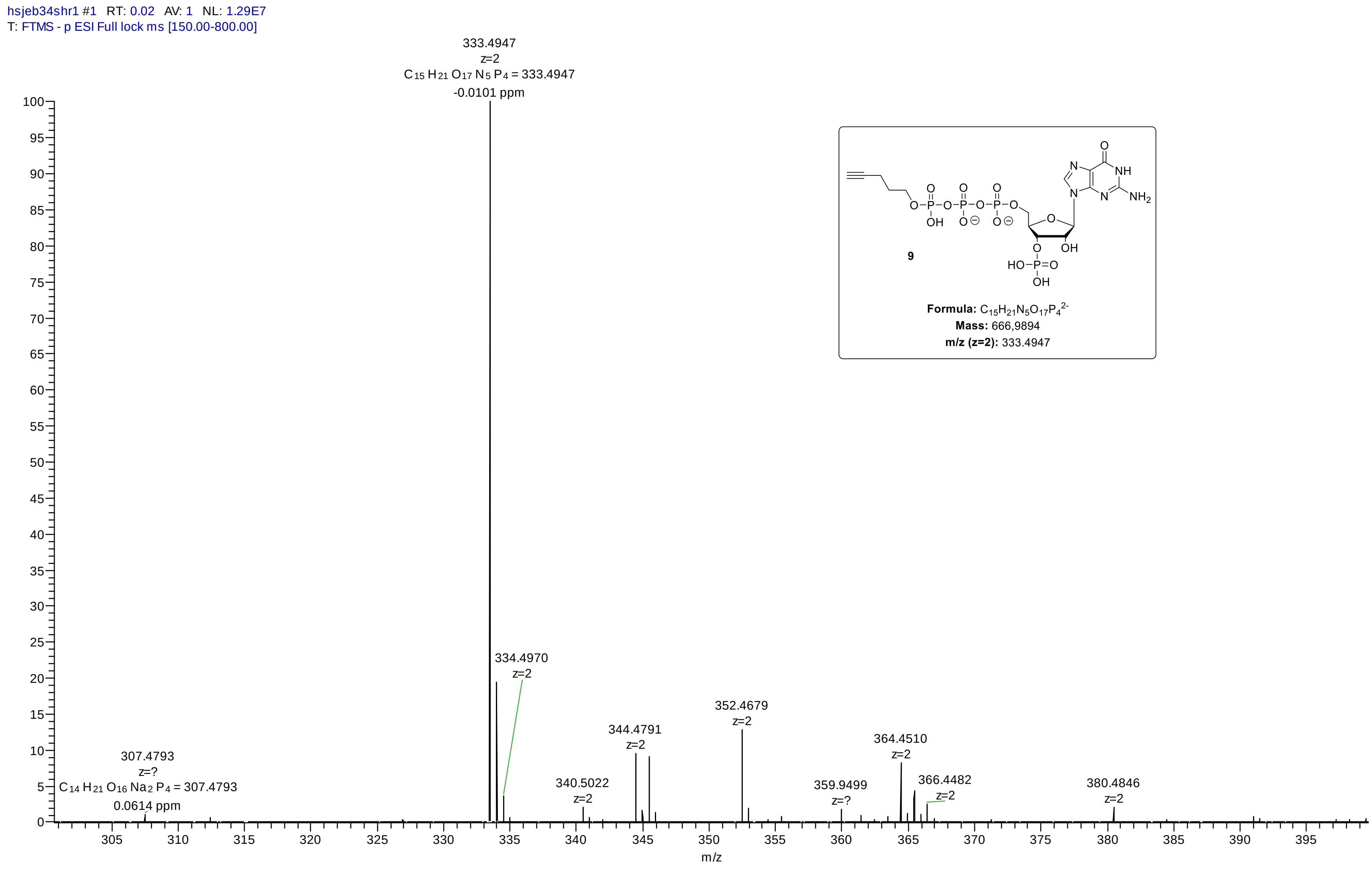

#### *HRMS (ESI) Analysis of compound 11: pentynyl - ppAp*

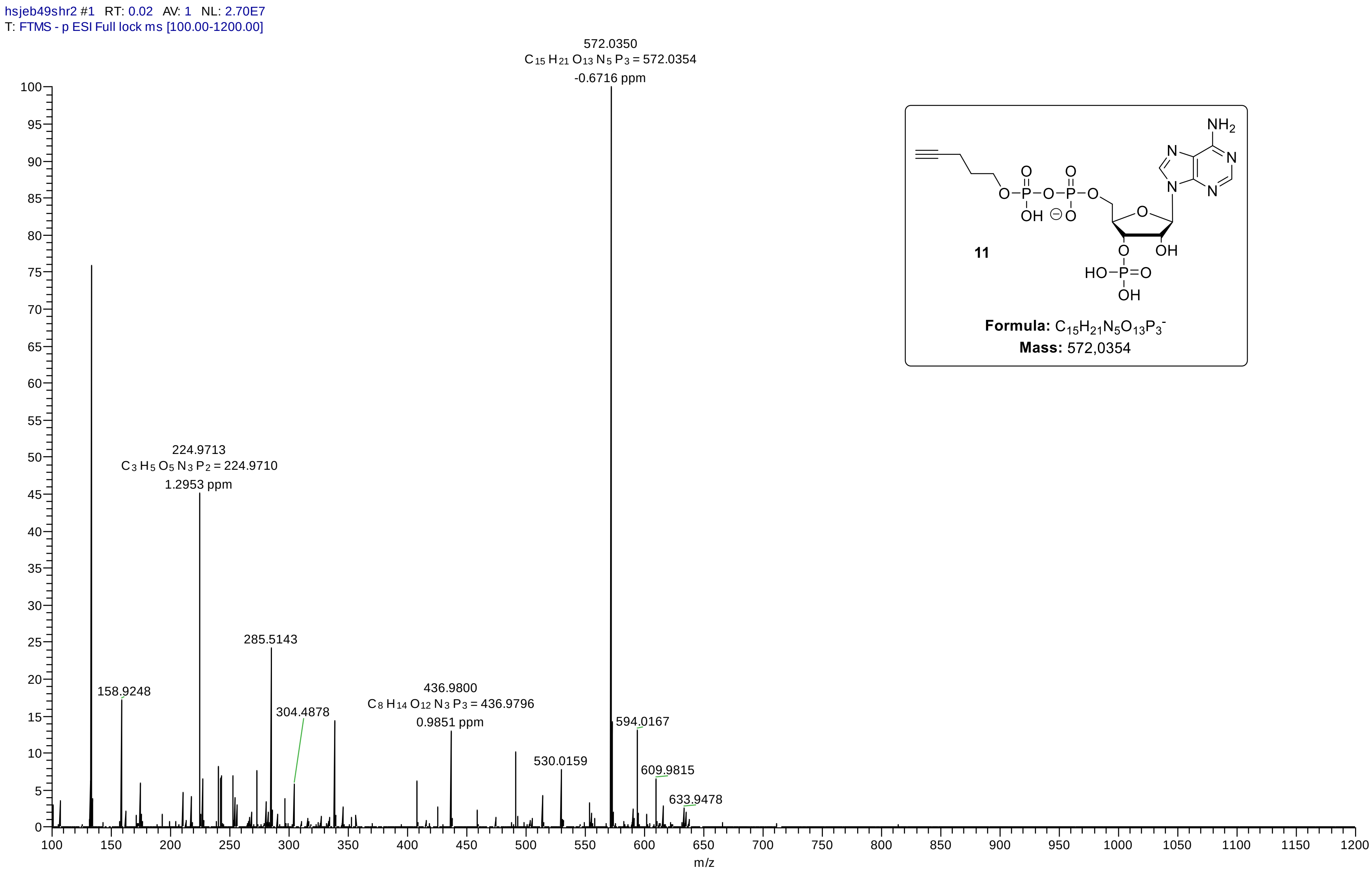


11

**Formula:** $C_{15}H_{21}N_5O_{13}P_3^-$
**Mass:** 572,0354

### *HRMS (ESI) Analysis of compound 12: pentynyl – pGp*

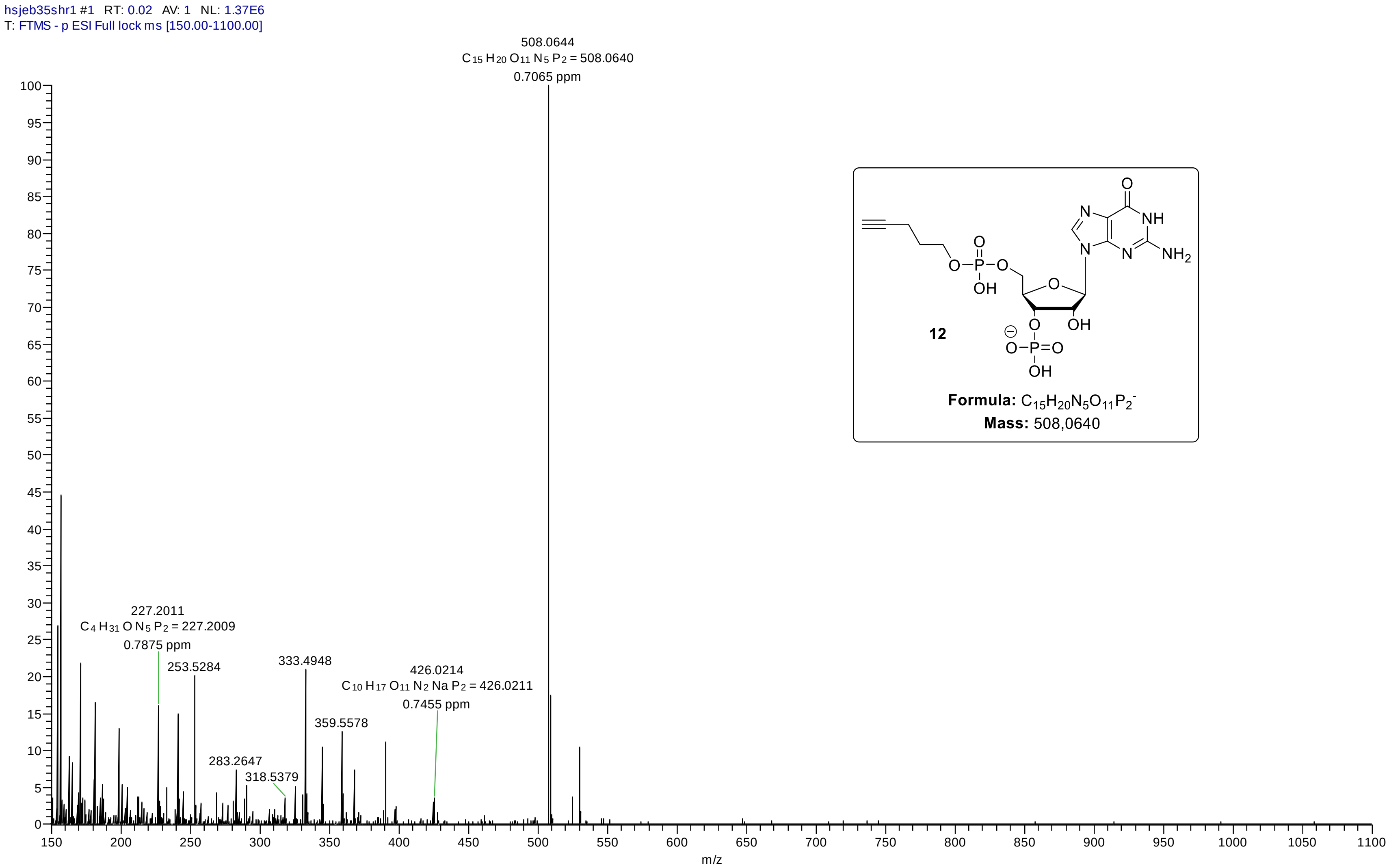

#### *HRMS (ESI) Analysis of compound 13: pentynyl – pGpp*

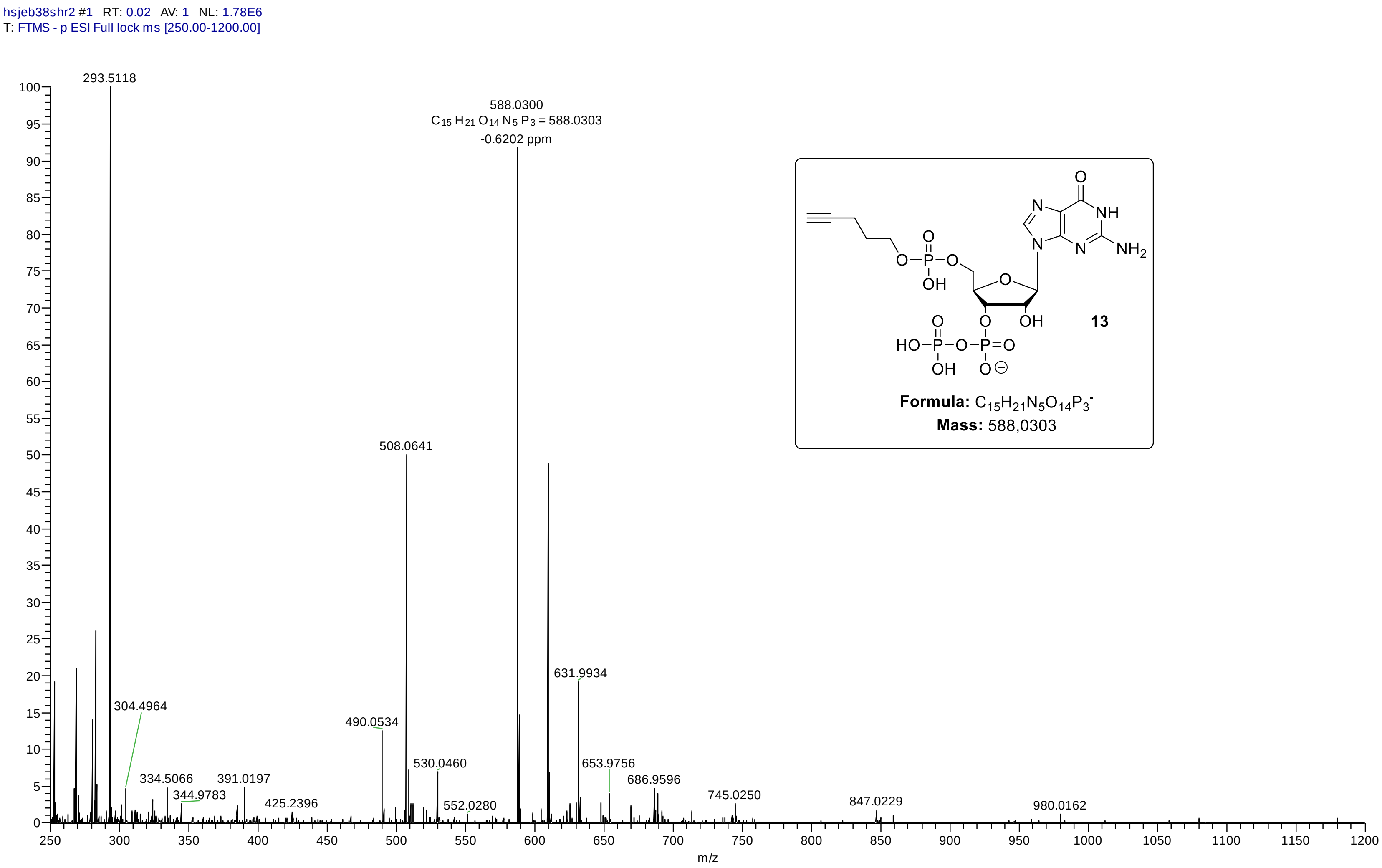

#### *HRMS (ESI) Analysis of compound 15: pentynyl – pppGpp*

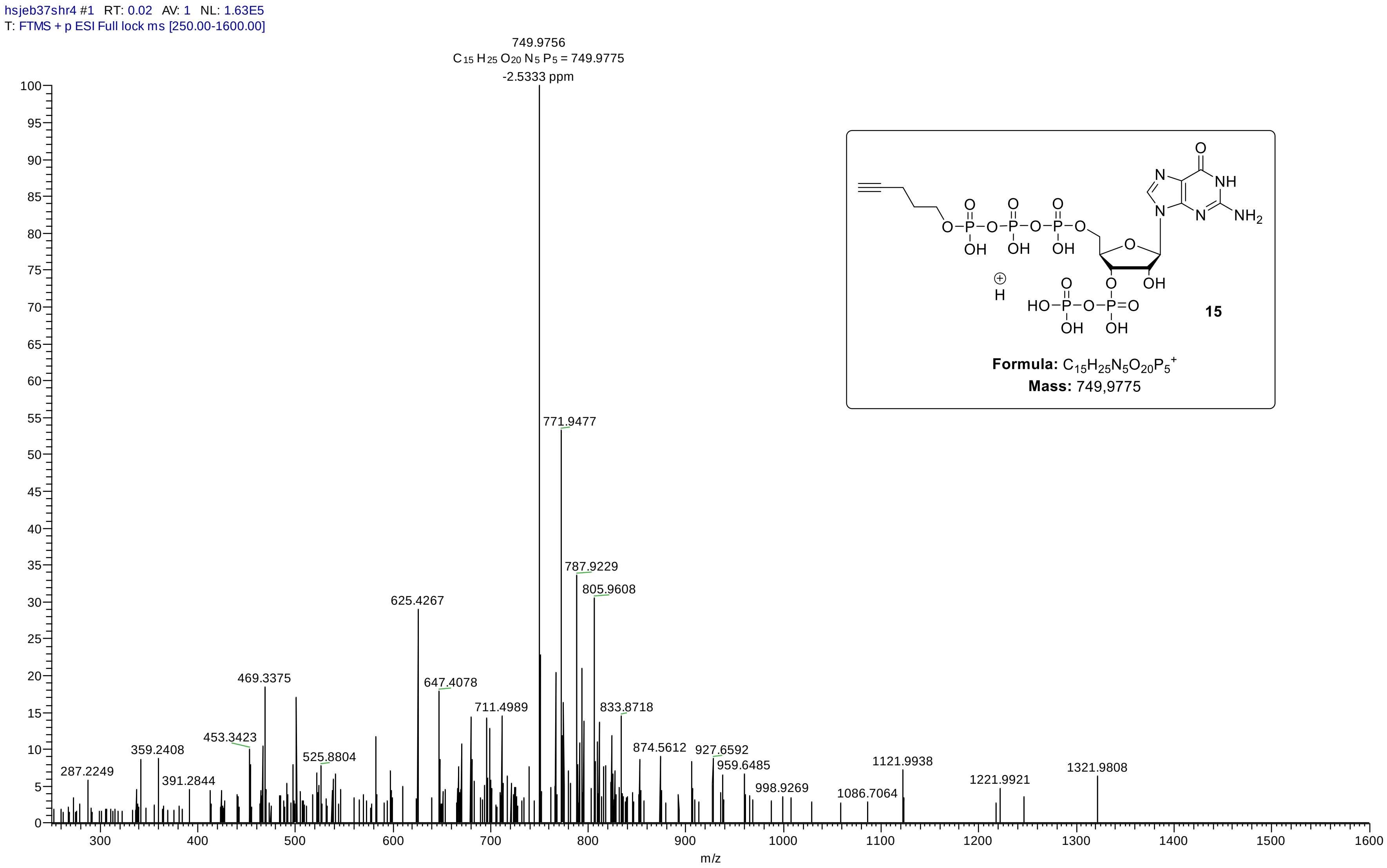

#### *HRMS (ESI) Analysis of compound 16: pentynyl – ppApp*

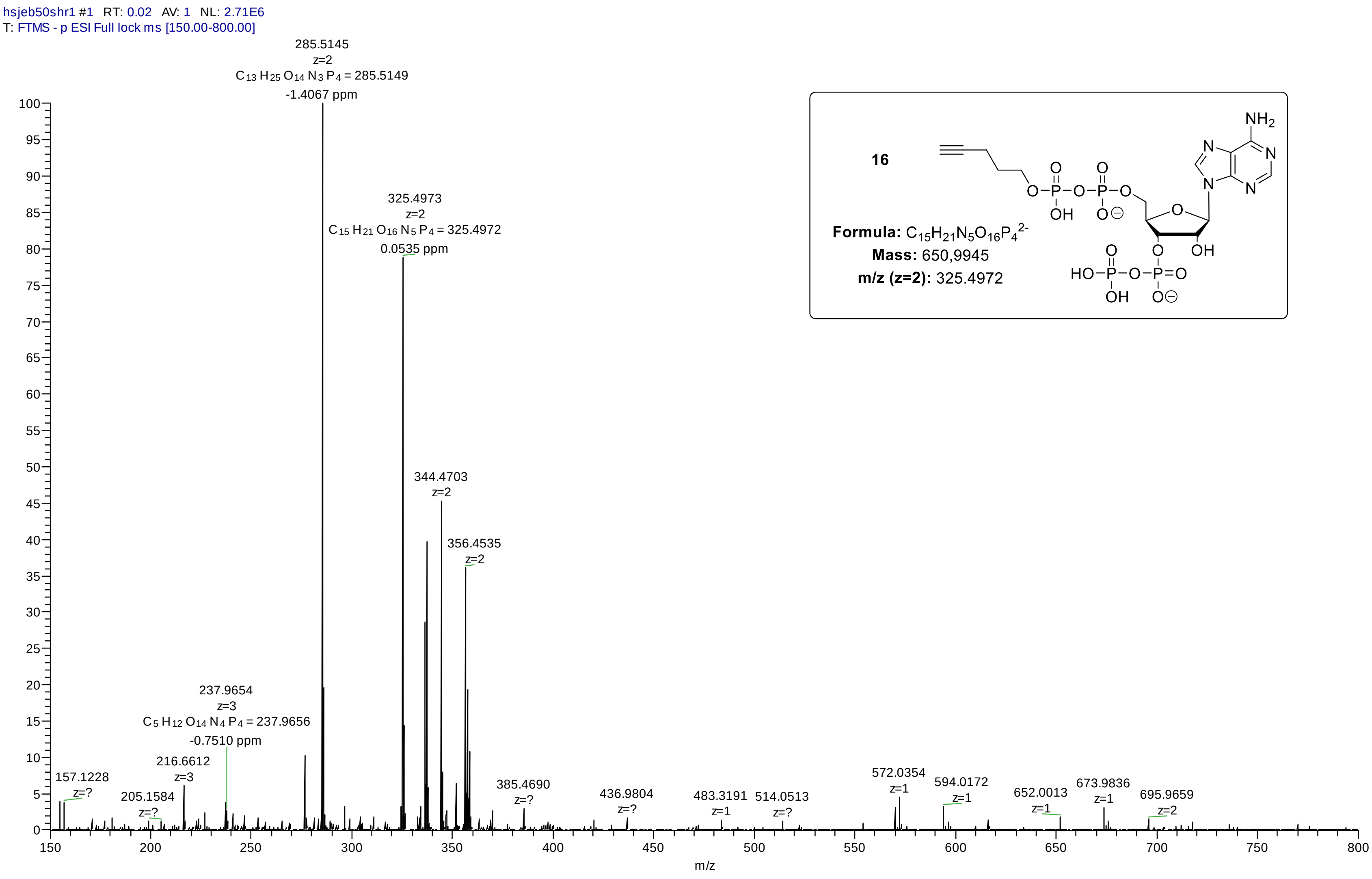


16

**Formula:** $C_{15}H_{21}N_5O_{16}P_4^{2-}$

**Mass:** 650,9945

**m/z (z=2):** 325.4972

#### *HRMS (ESI) Analysis of compound 17: amino – pGpp*

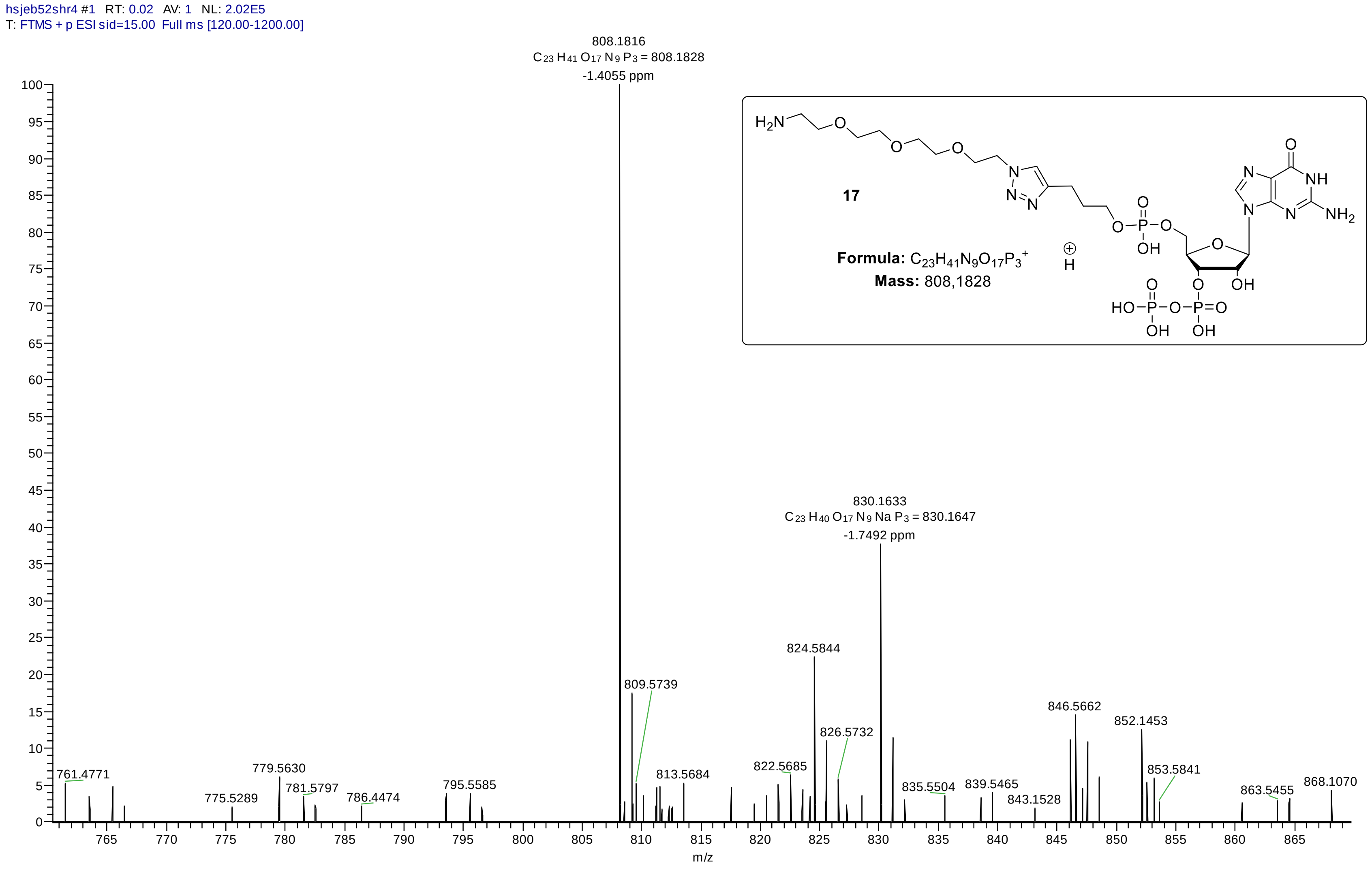

#### *HRMS (ESI) Analysis of compound 19: amino – pppGpp*

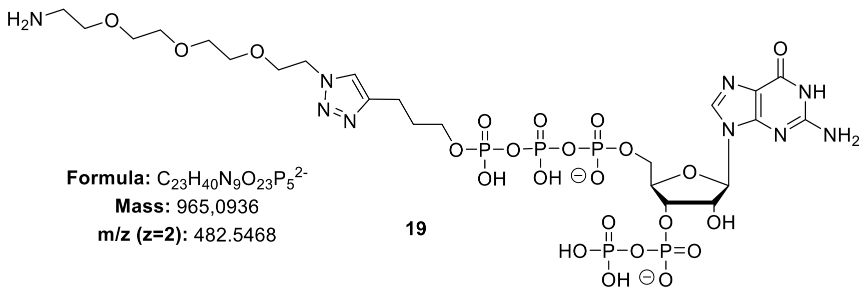

**Formula:** $C_{23}H_{40}N_9O_{23}P_5^{2-}$
**Mass:** 965,0936
**m/z (z=2):** 482.5468

**19**

##### Sample Spectra

**- Scan (rt: 0.090-0.166 min) Sub**

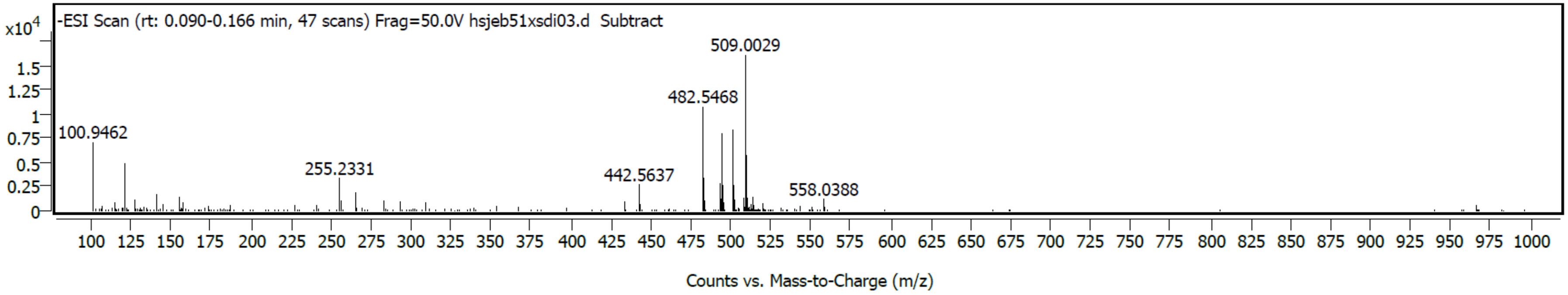

#### *HRMS (ESI) Analysis of compound 20: amino – ppApp*

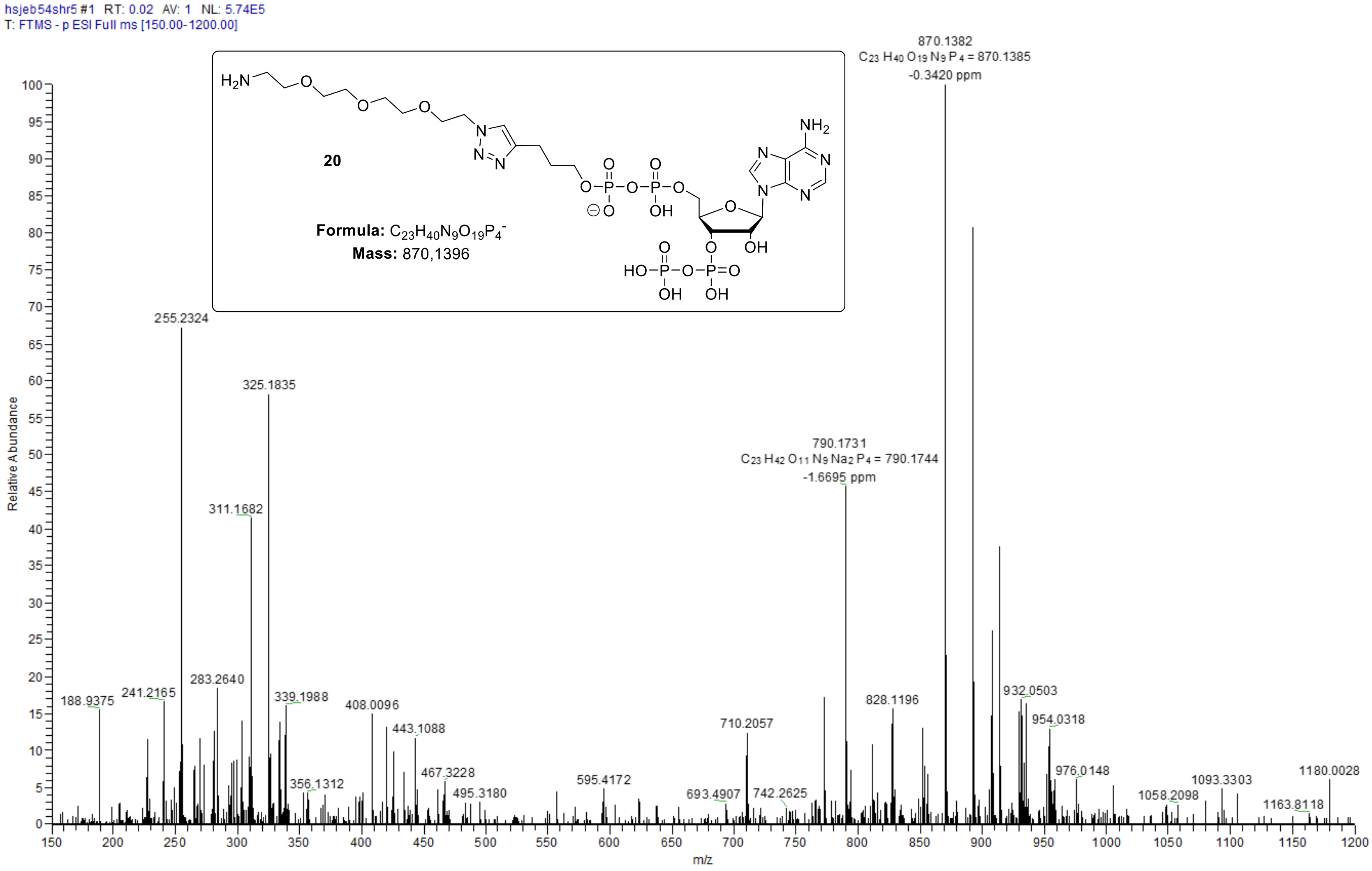

#### *HRMS (ESI) Analysis of compound SI-12: amino – ppGp*

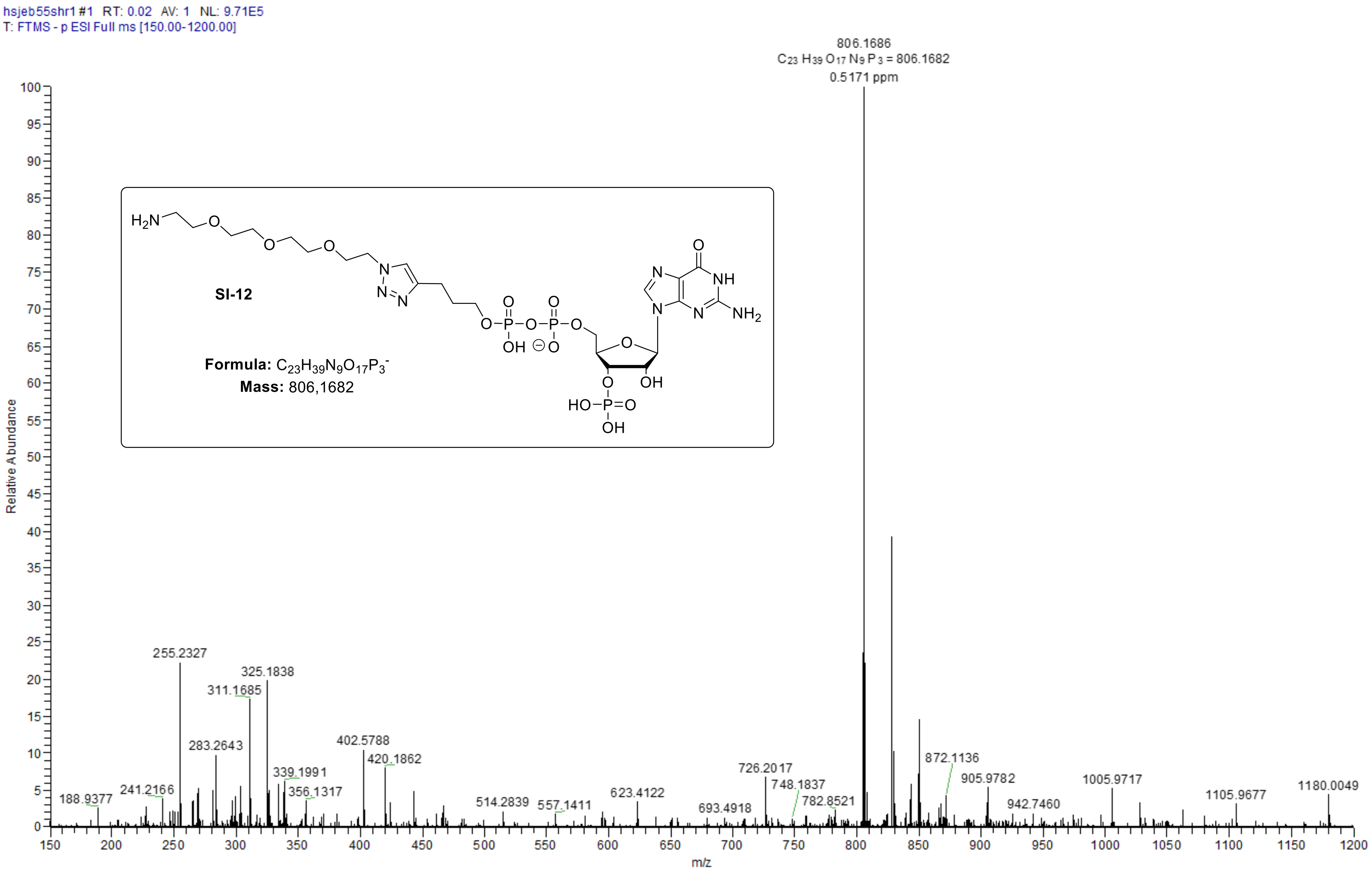

### *HRMS (ESI) Analysis of compound 21: ppGpp – CC1*

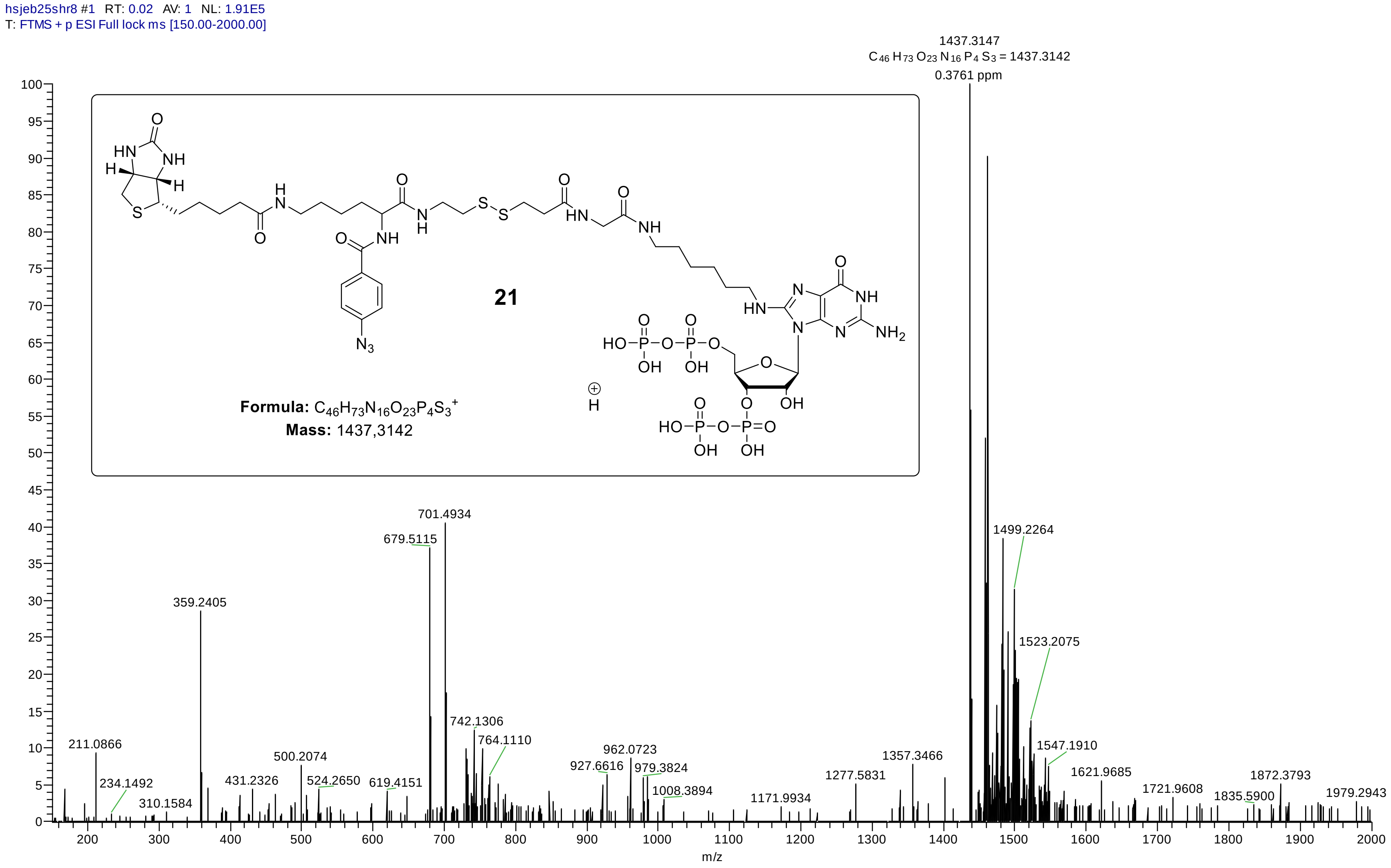

#### *HRMS (ESI) Analysis of compound 22: ppGpp – CC2*

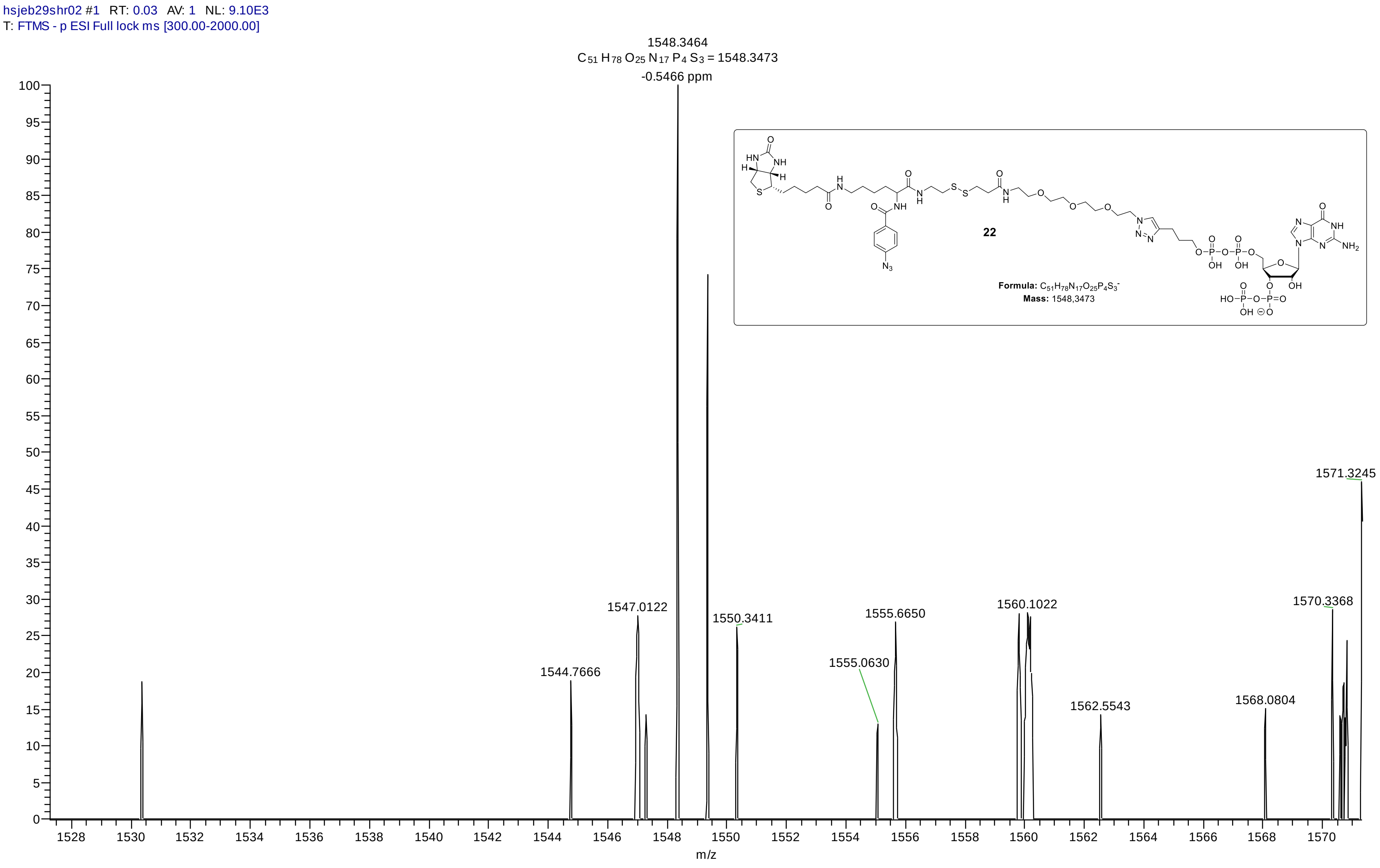

#### *HRMS (ESI) Analysis of compound 23: ppGpp – CC3*

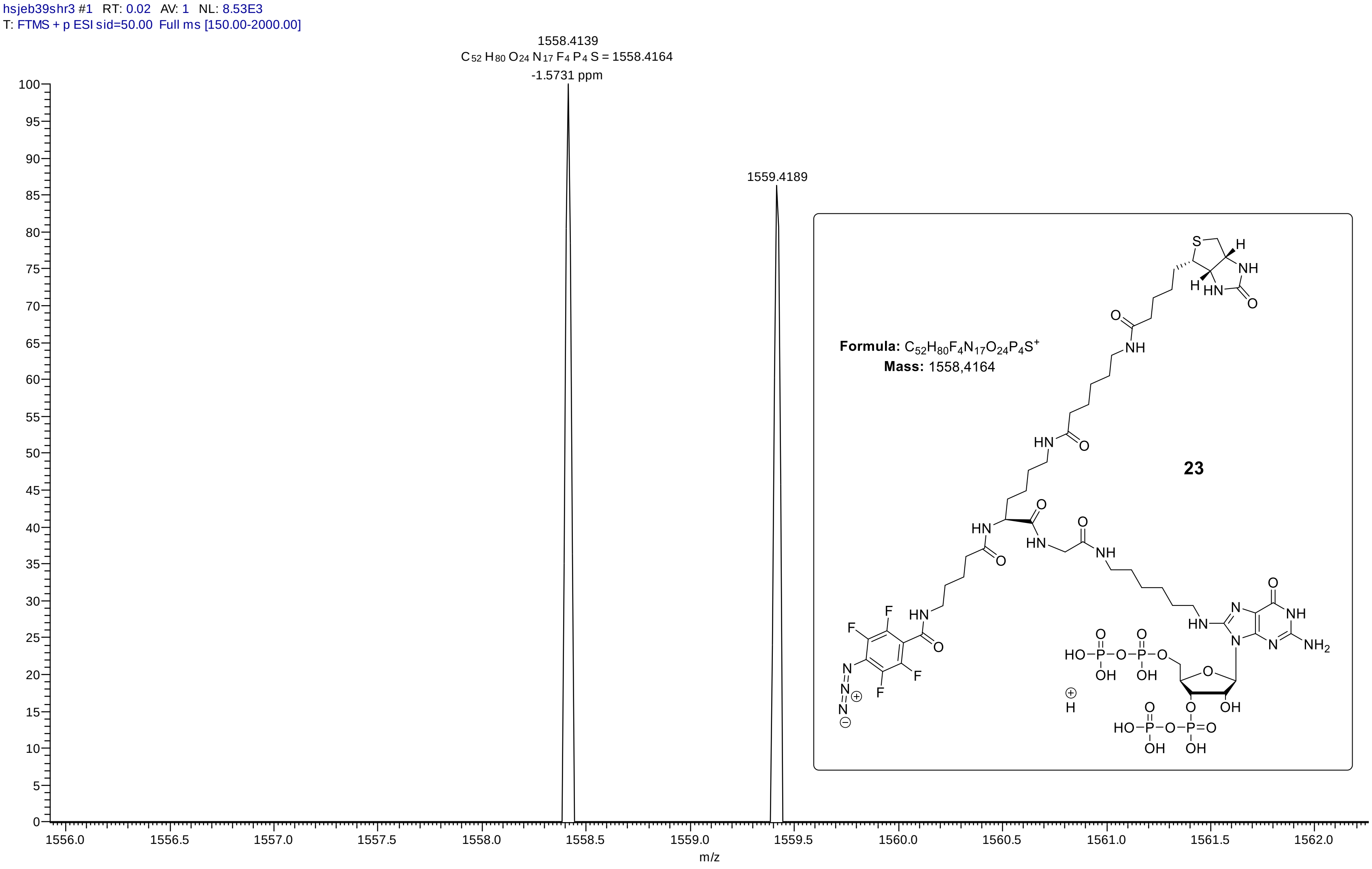

#### *HRMS (ESI) Analysis of compound XX: ppGpp – CC4*

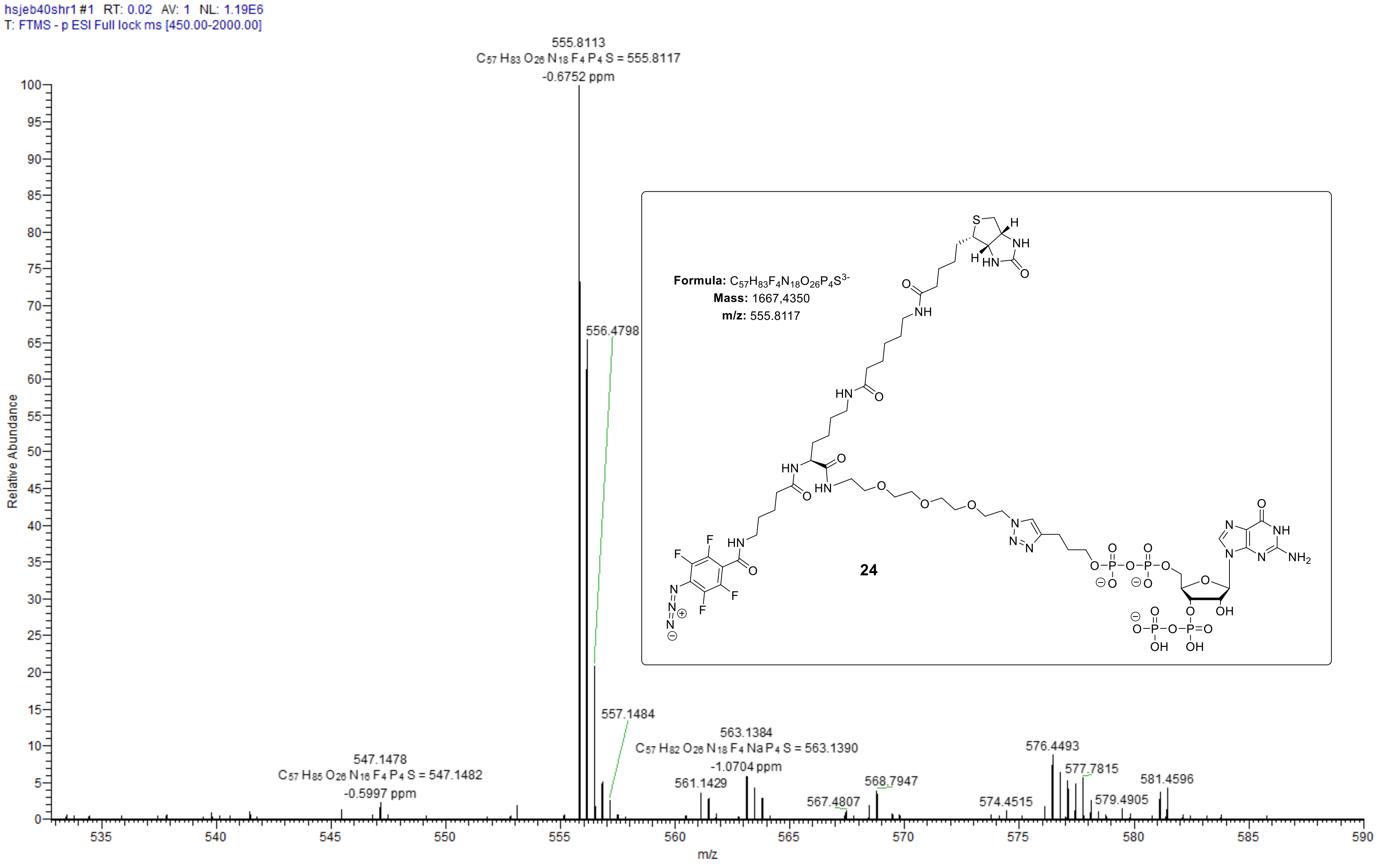

### *HRMS (ESI) Analysis of compound 25: pGpp - CC*

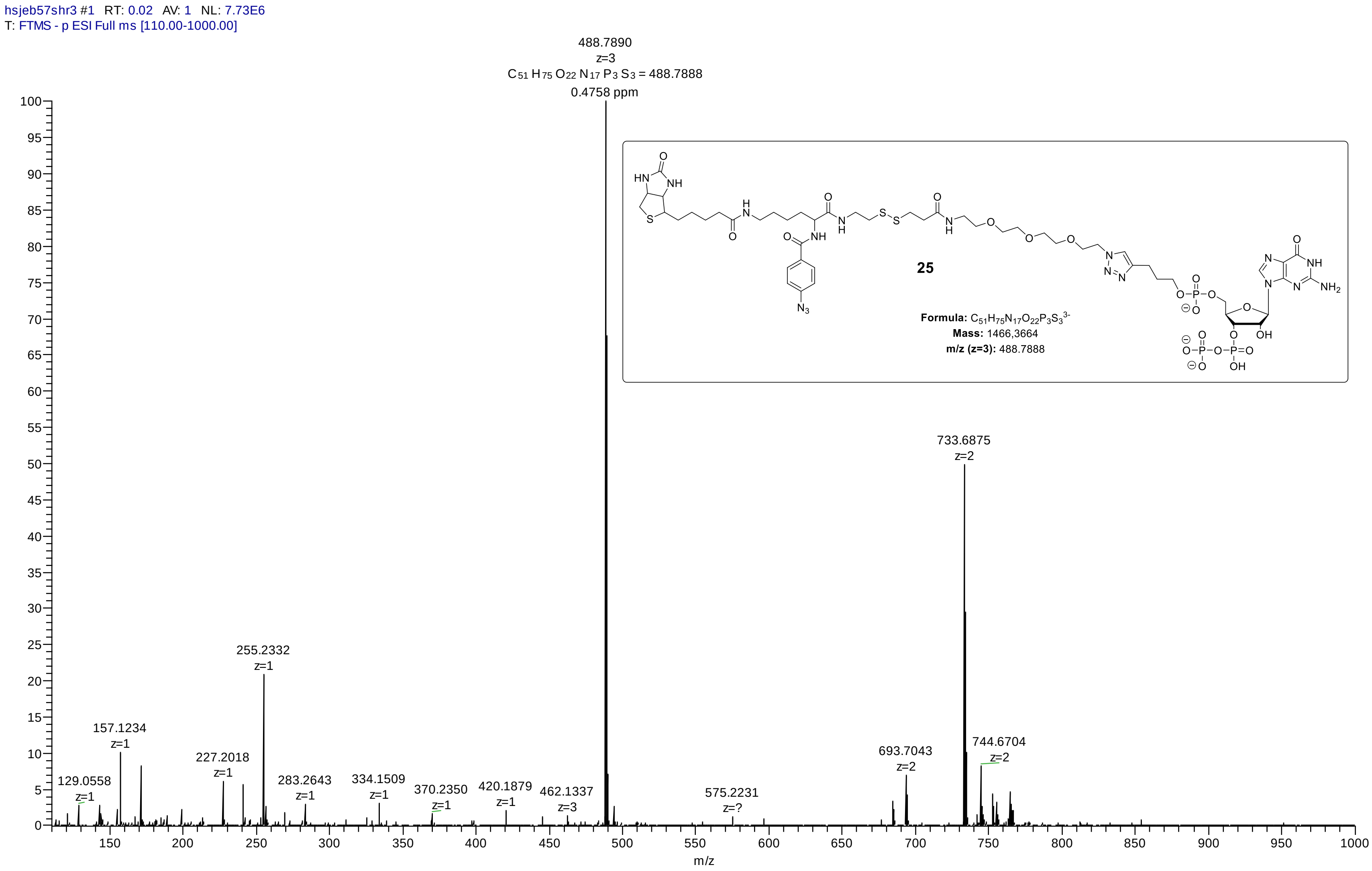

#### *HRMS (ESI) Analysis of compound 26: pppGpp - CC*

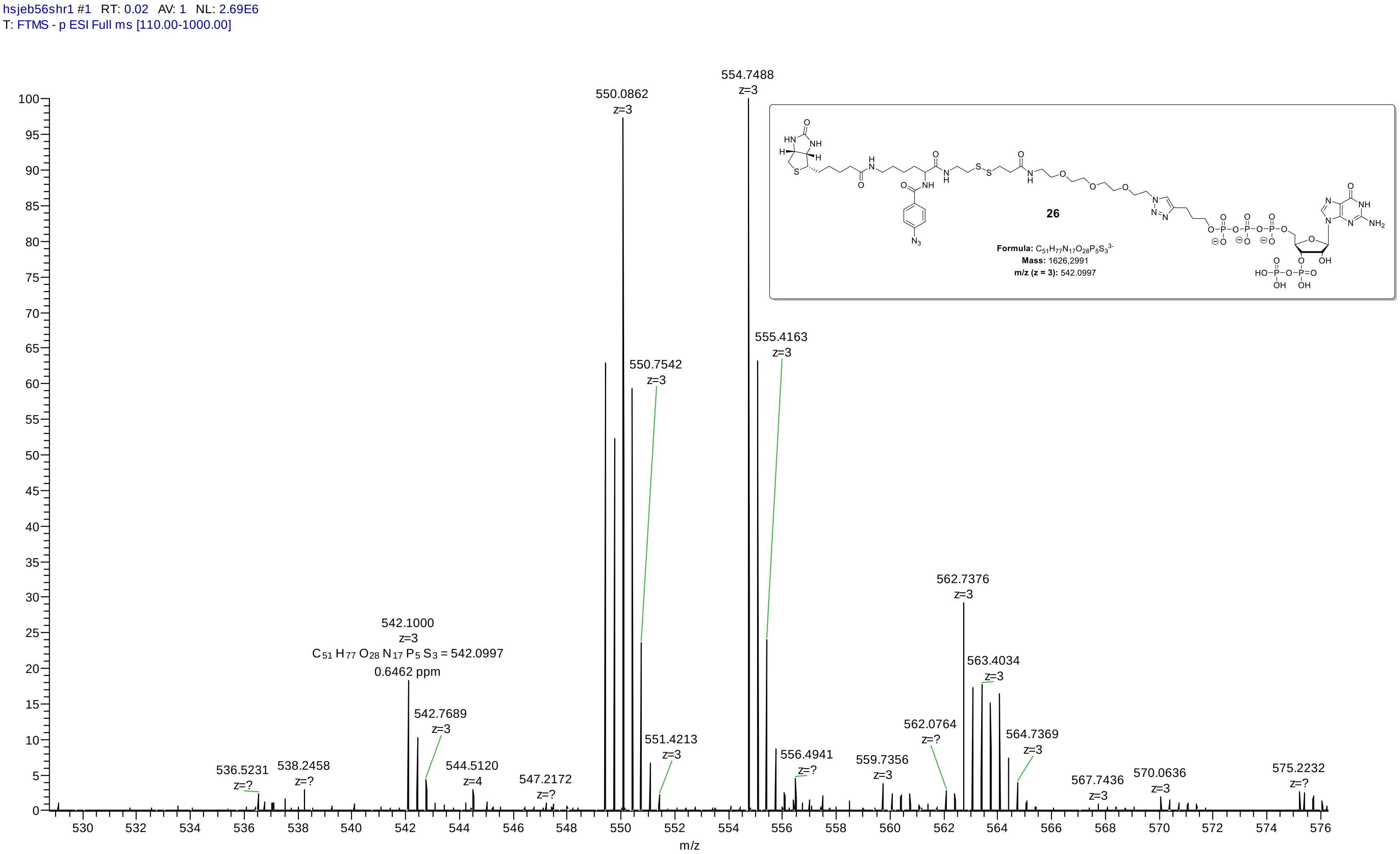

### *HRMS (ESI) Analysis of compound 27: ppApp - CC*

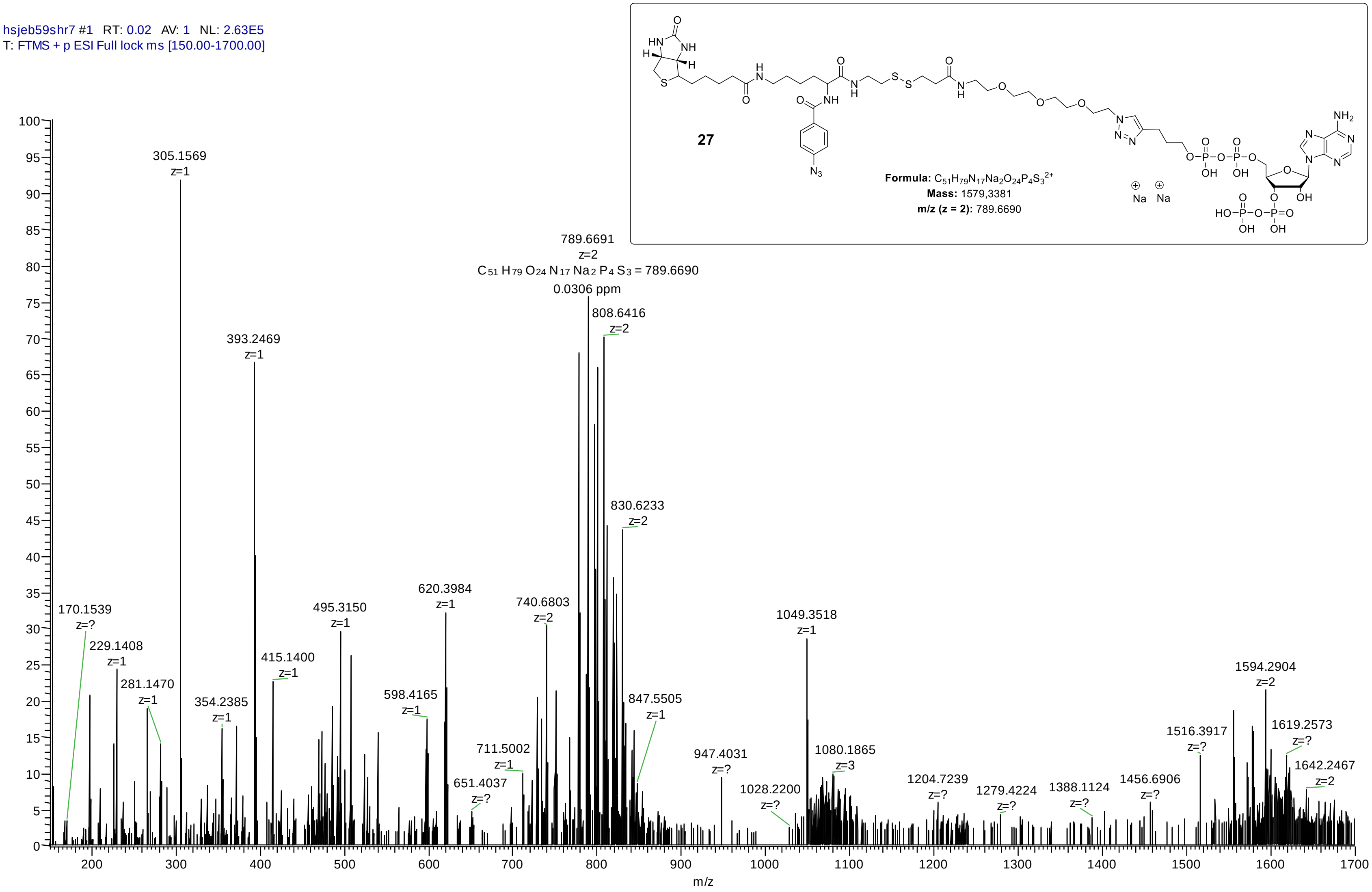

### *HRMS (ESI) Analysis of compound SI-13: ppGp - CC*

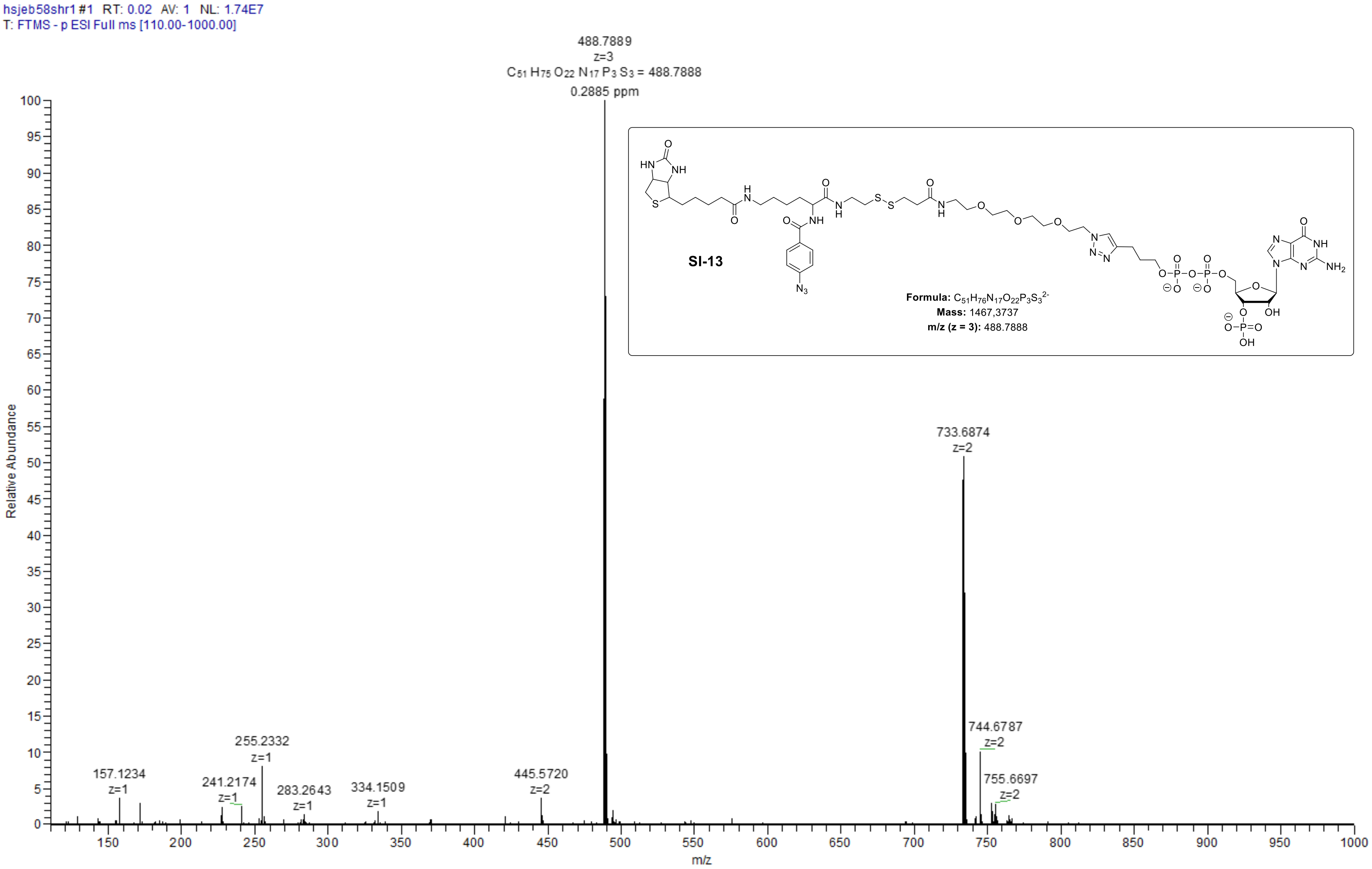

#### *HRMS (ESI) Analysis of compound SI-5: Biotin-NHS derivative*

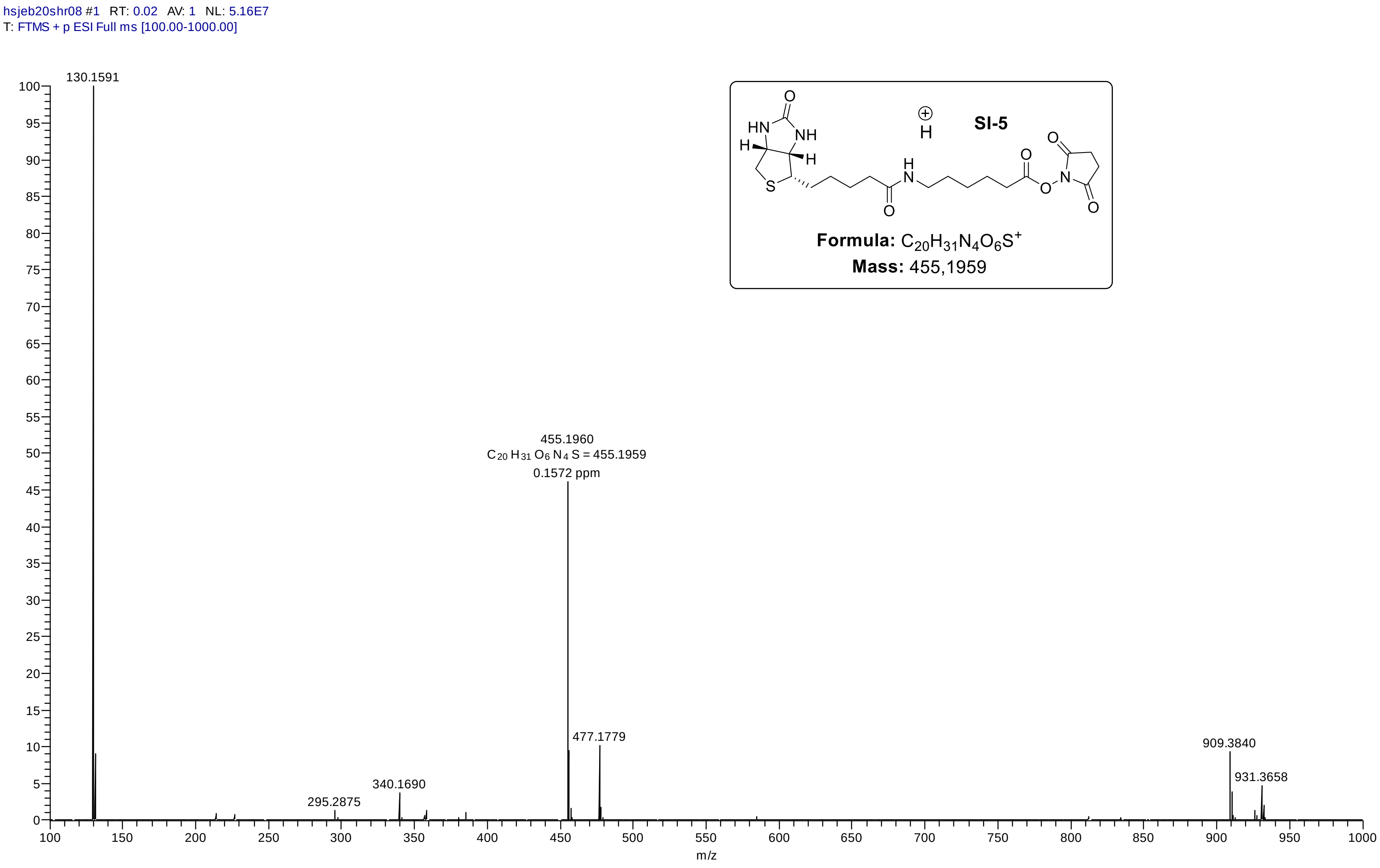

#### *HRMS (ESI) Analysis of compound SI-6: Biotin-lysine moiety*

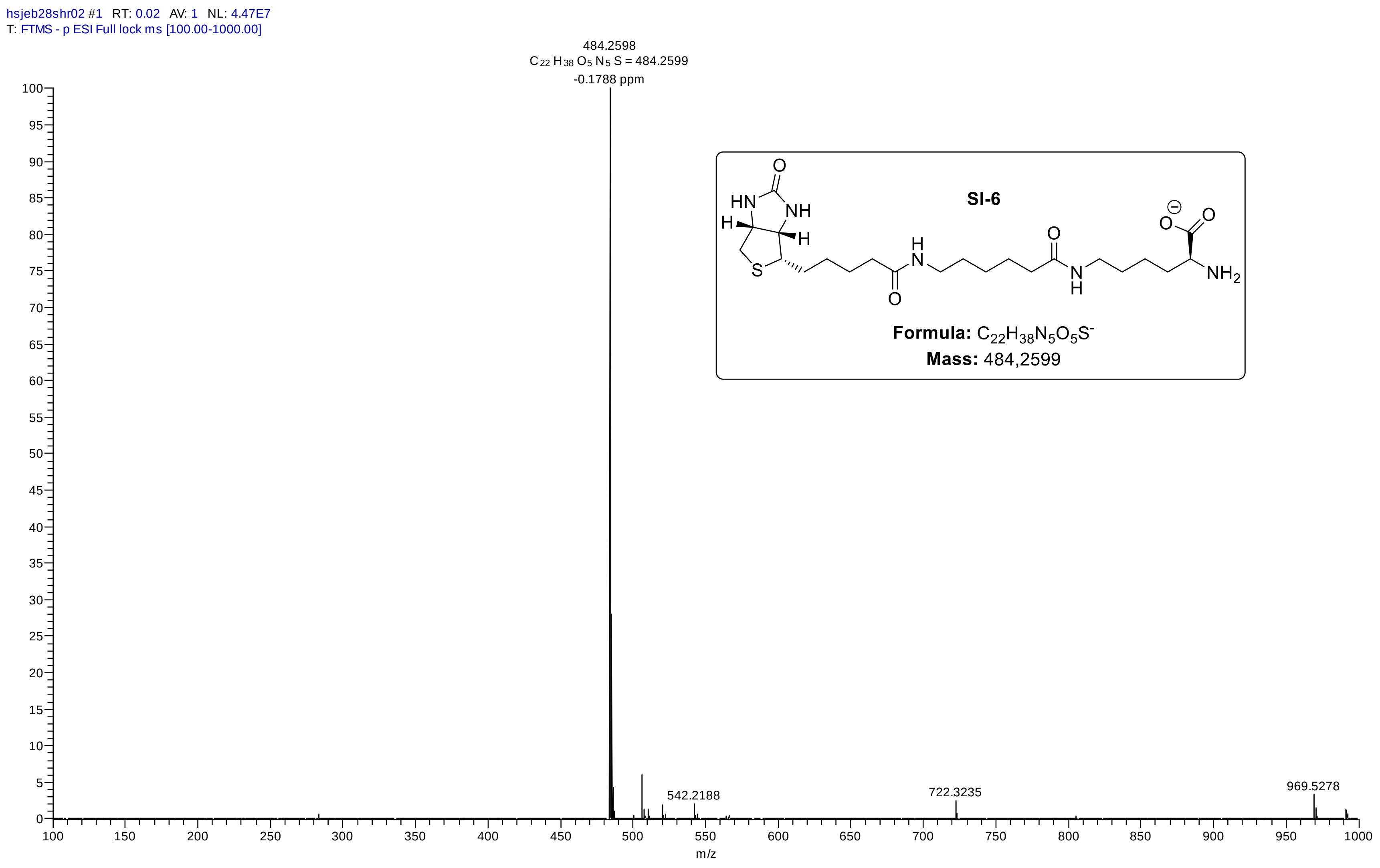

#### *HRMS (ESI) Analysis of compound SI-10: Fluorophenylazide derivative*

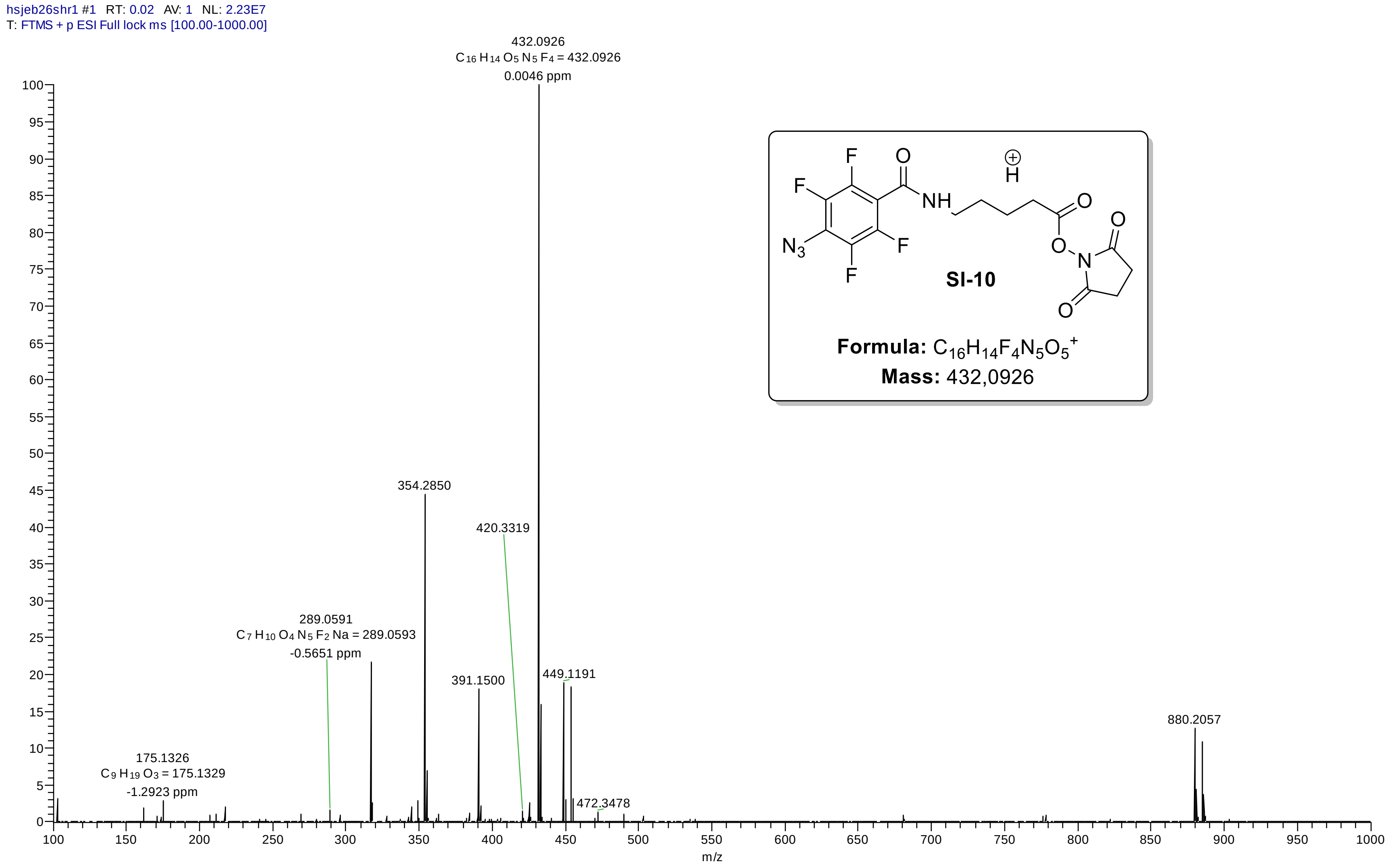

#### *HRMS (ESI) Analysis of compound SI-11: linker structure (carboxylic acid)*

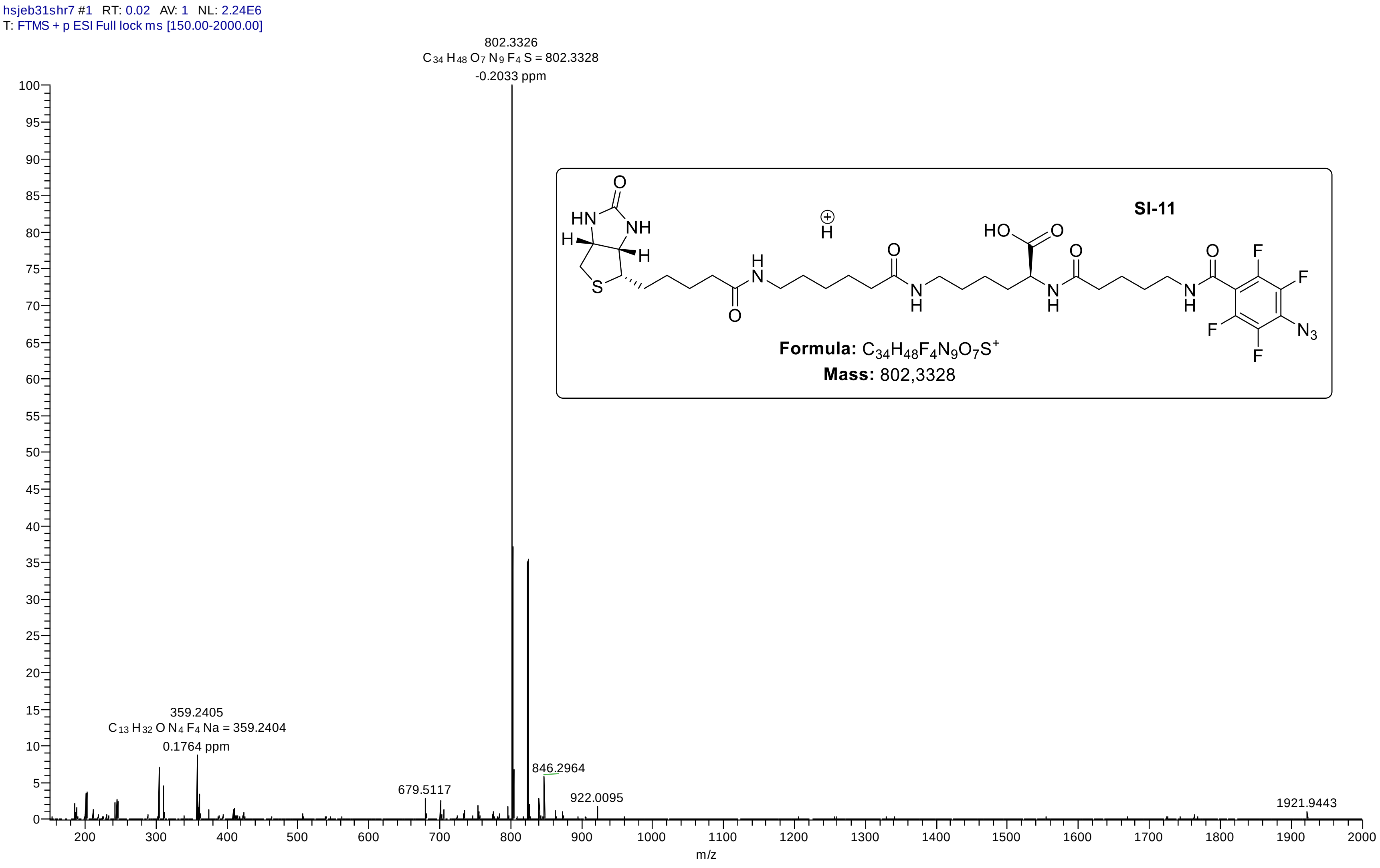

### *9. HPLC – analysis*

*HPLC-UV measurements were performed using a dionex ultimate 3000 system and a C18AQ – column. A gradient of $H_2O$/MeCN/TEAA(pH 7, 100 mM) was applied.*

#### *HPLC – UV – analysis of compound 7: amino – ppGpp 1*

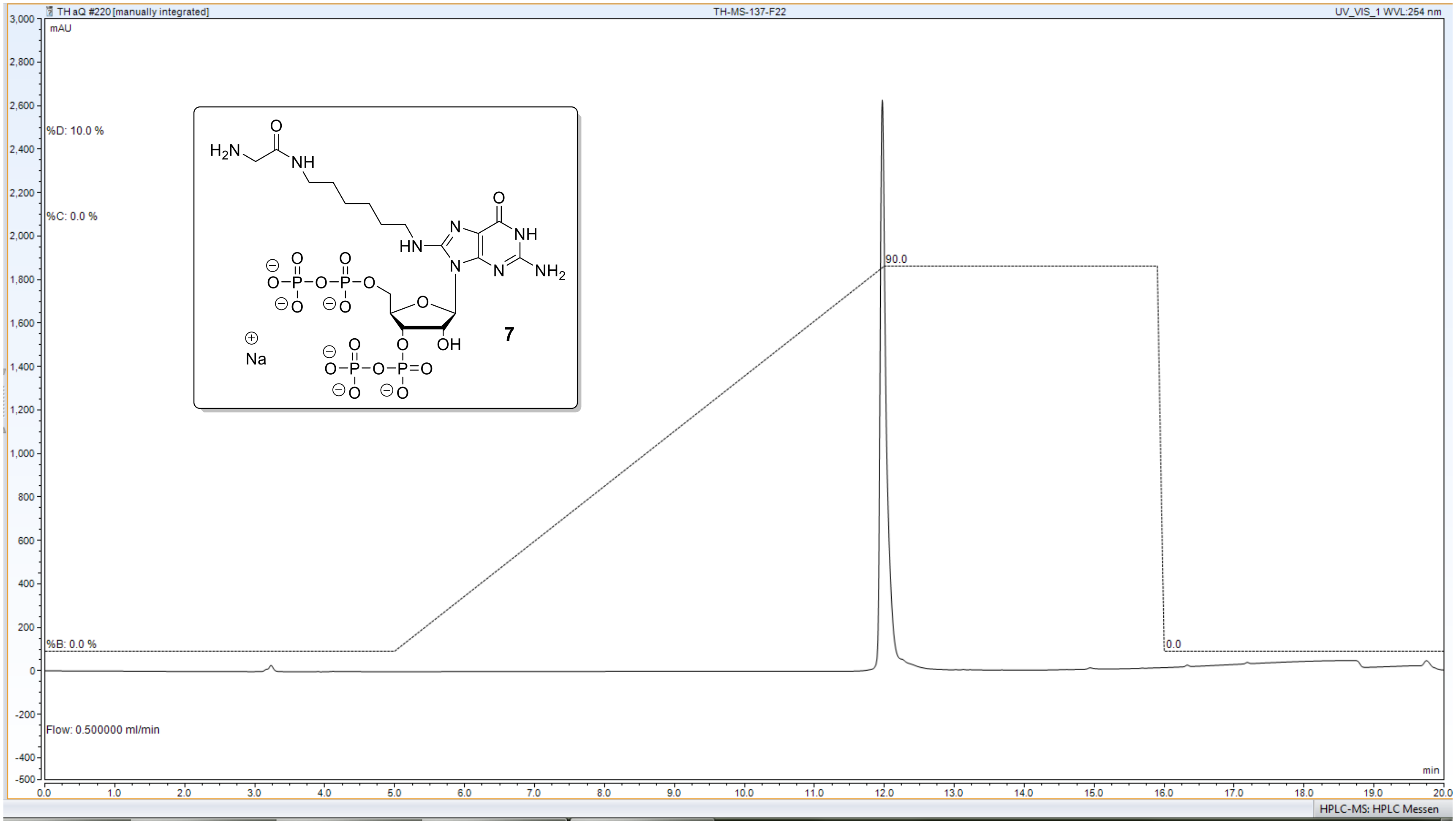


**C18 AQ – column** ($H_2O$/MeCN – gradient, 10 mM TEAA)

##### *HPLC – UV – analysis of compound 17: amino – pGpp*

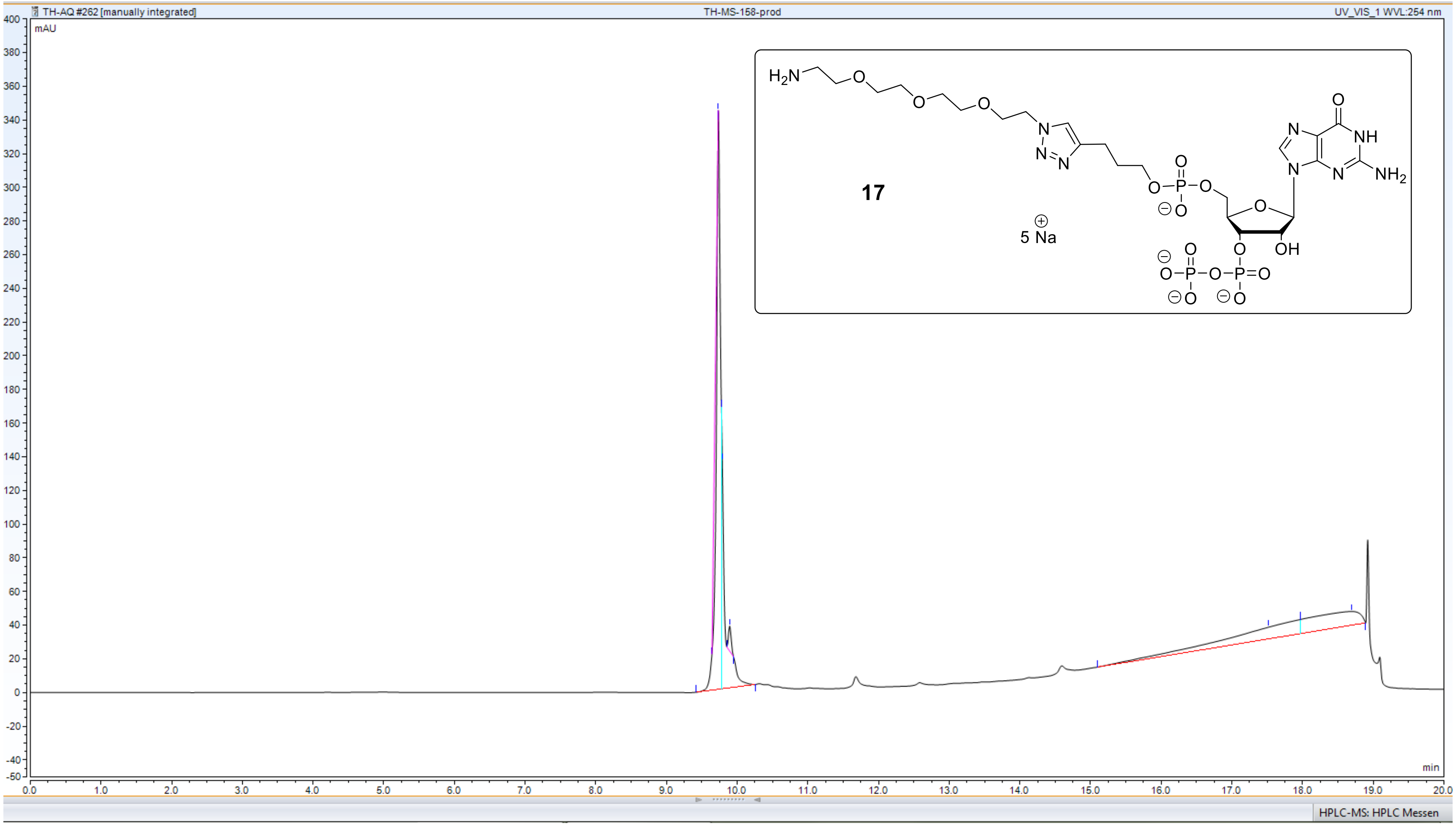


**C18 AQ – column** ($H_2O$/MeCN – gradient, 10 mM TEAA)

##### *HPLC – UV – analysis of compound 18: amino – ppGpp 2*

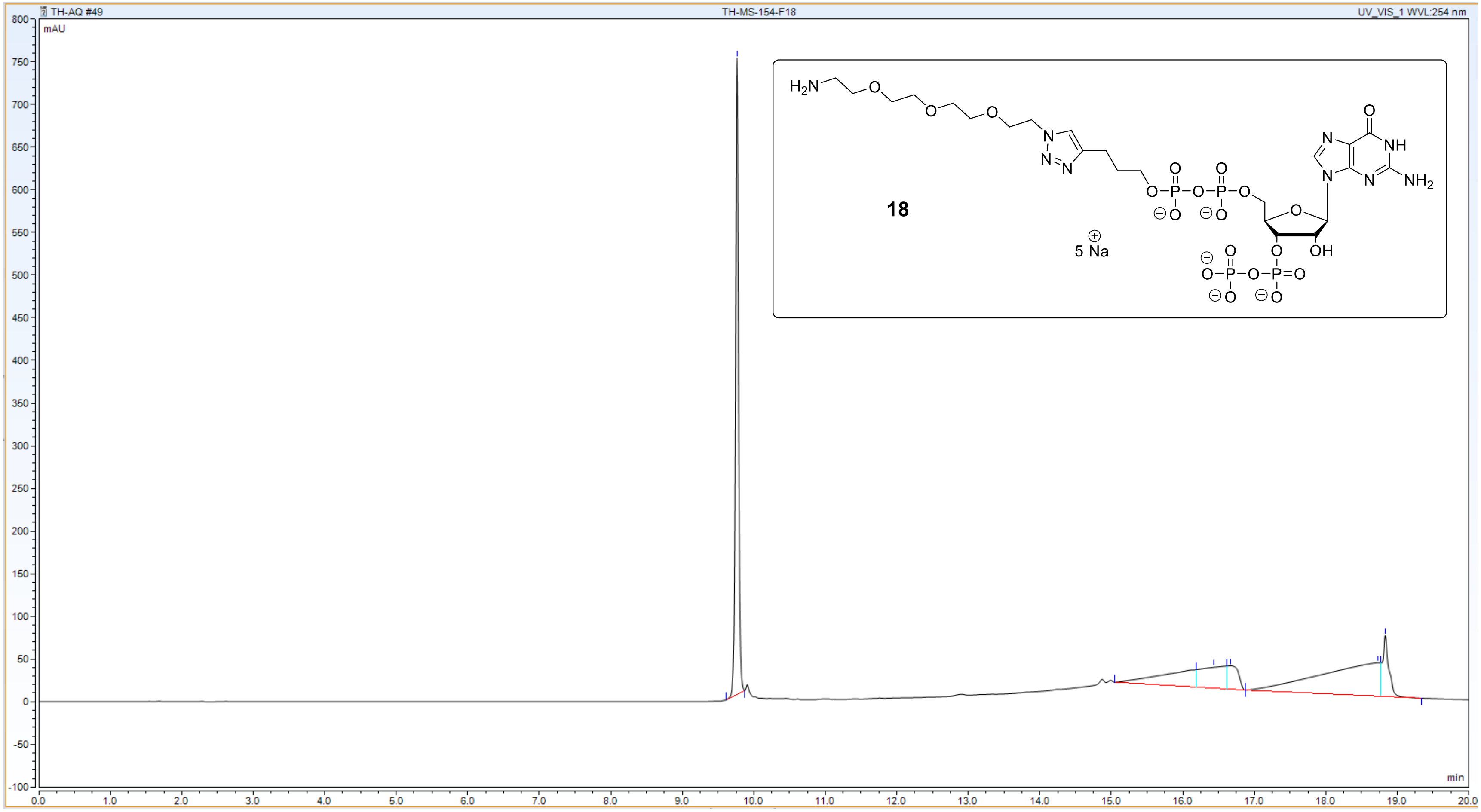


**C18 AQ – column** ($H_2O$/MeCN – gradient, 10 mM TEAA)

***HPLC – UV – analysis of compound 19: amino – pppGpp***

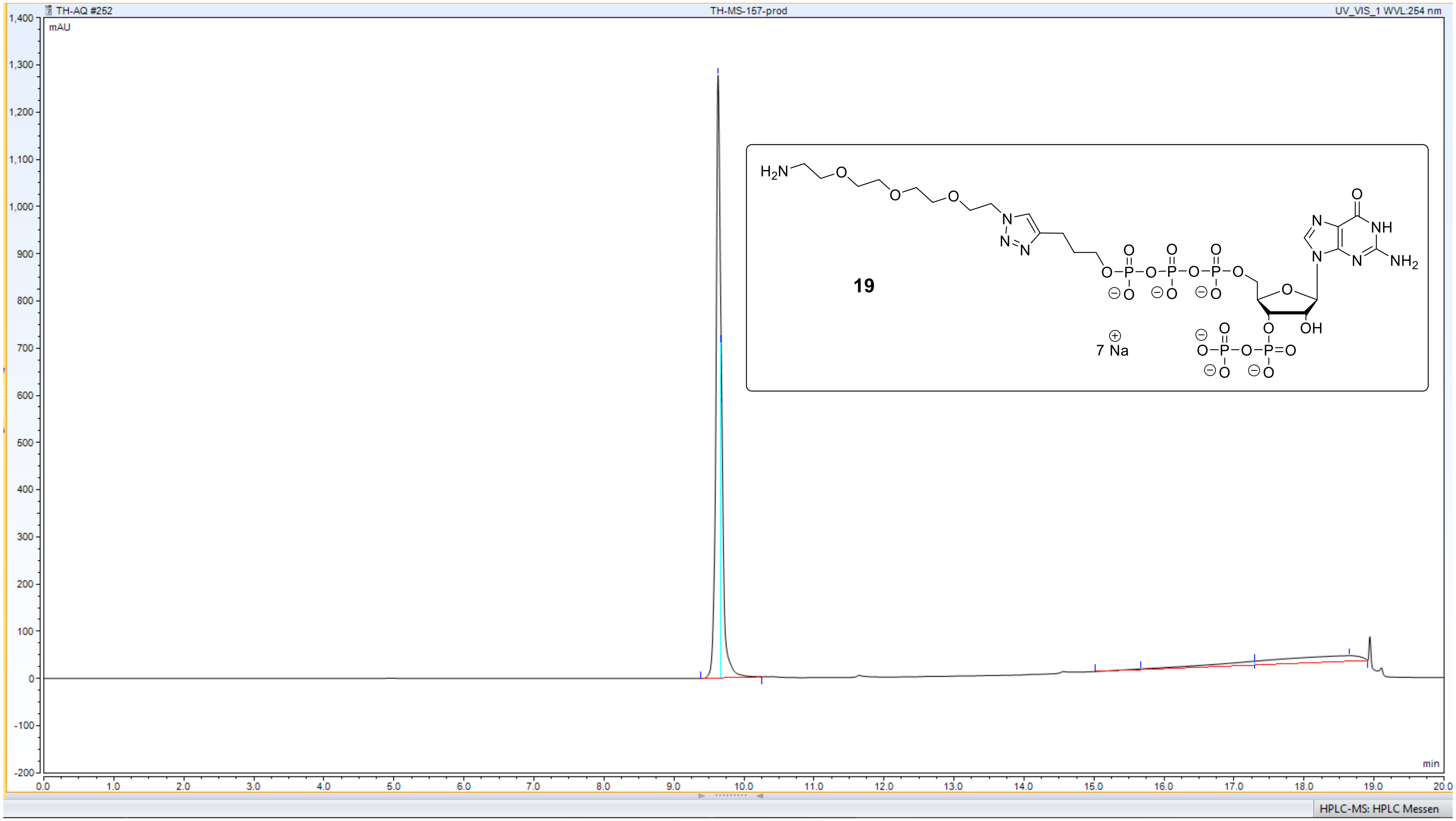


**C18 AQ – column** ($H_2O$/MeCN – gradient, 10 mM TEAA)

##### *HPLC – UV – analysis of compound 20: amino – ppApp*

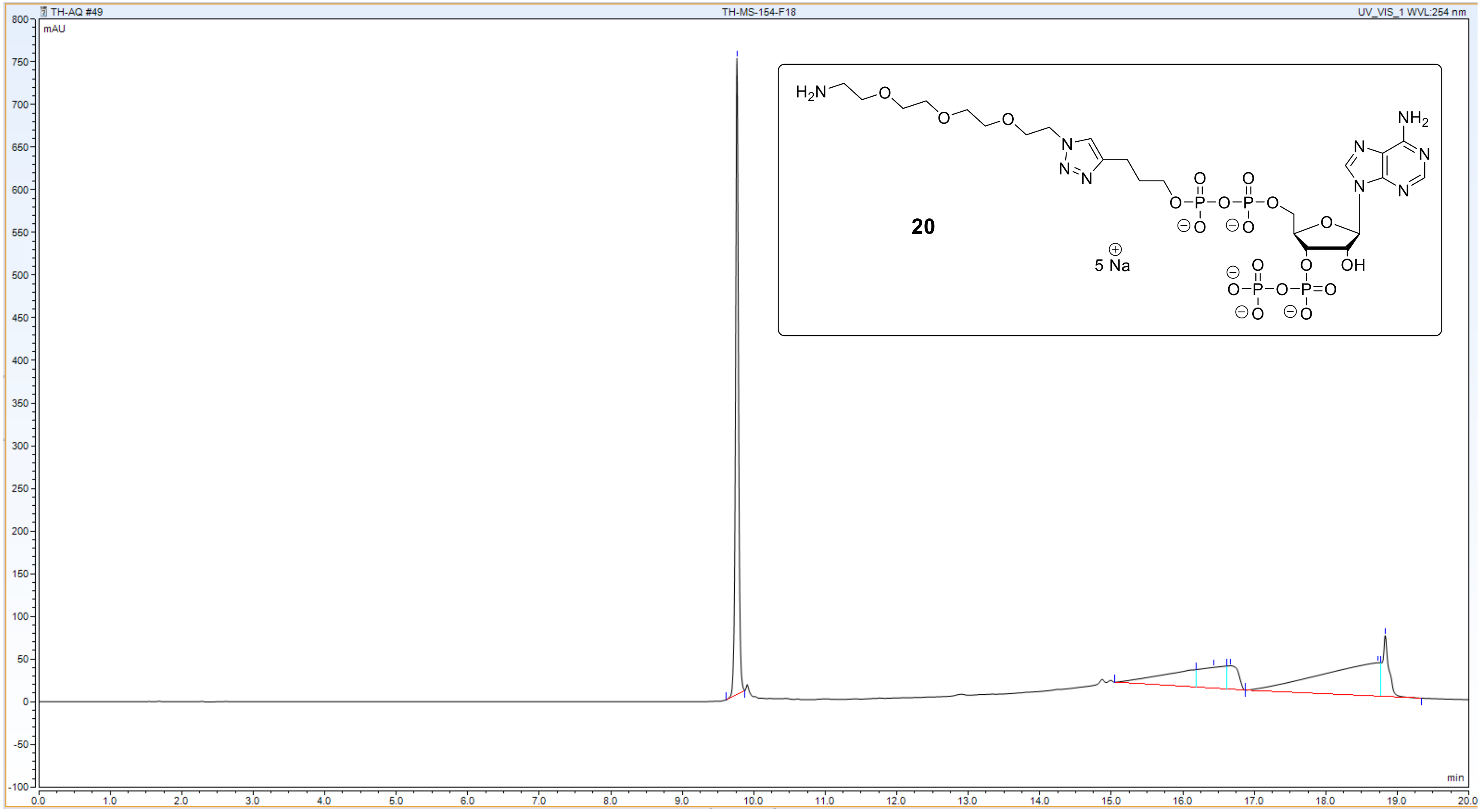


**C18 AQ – column** ($H_2O$/MeCN – gradient, 10 mM TEAA)

#### *HPLC – UV – analysis of compound SI-12: amino – ppGp*

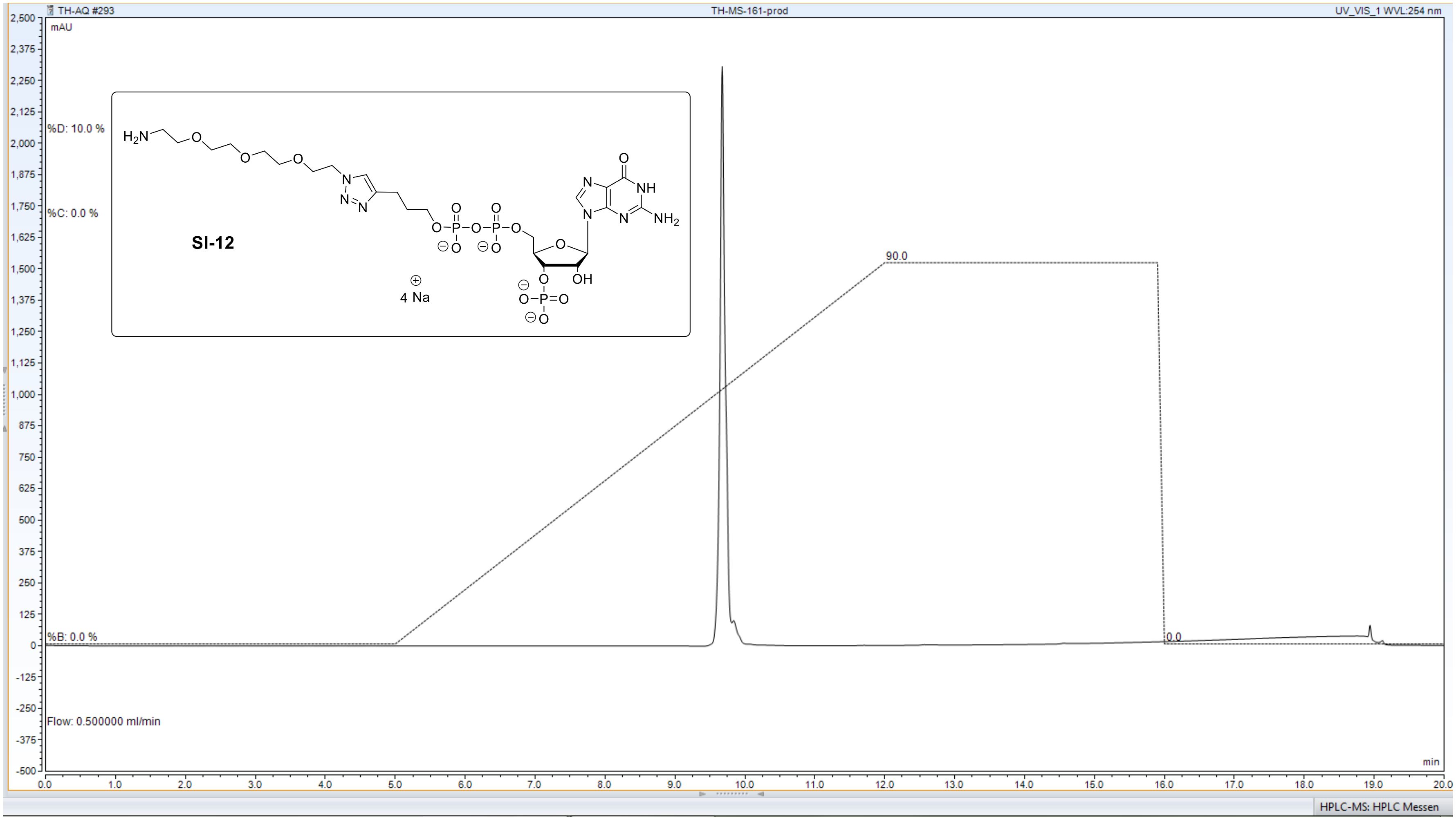


**C18 AQ – column** ($H_2O$/MeCN – gradient, 10 mM TEAA)

#### *HPLC – UV – analysis of compound 21: ppGpp – CC1*

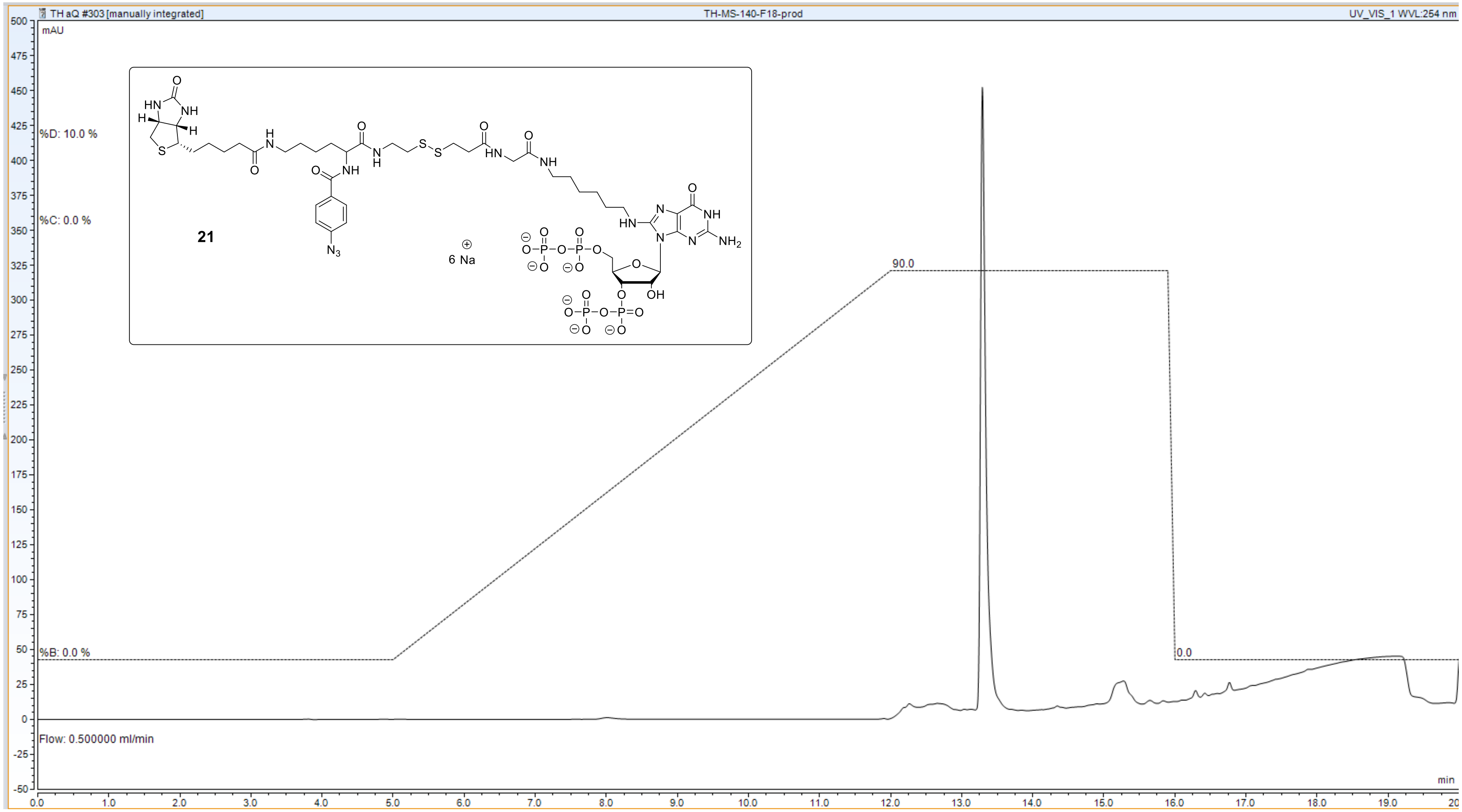


**C18 AQ – column** ($H_2O$/MeCN – gradient, 10 mM TEAA)

#### *HPLC – UV – analysis of compound 22: ppGpp – CC2*

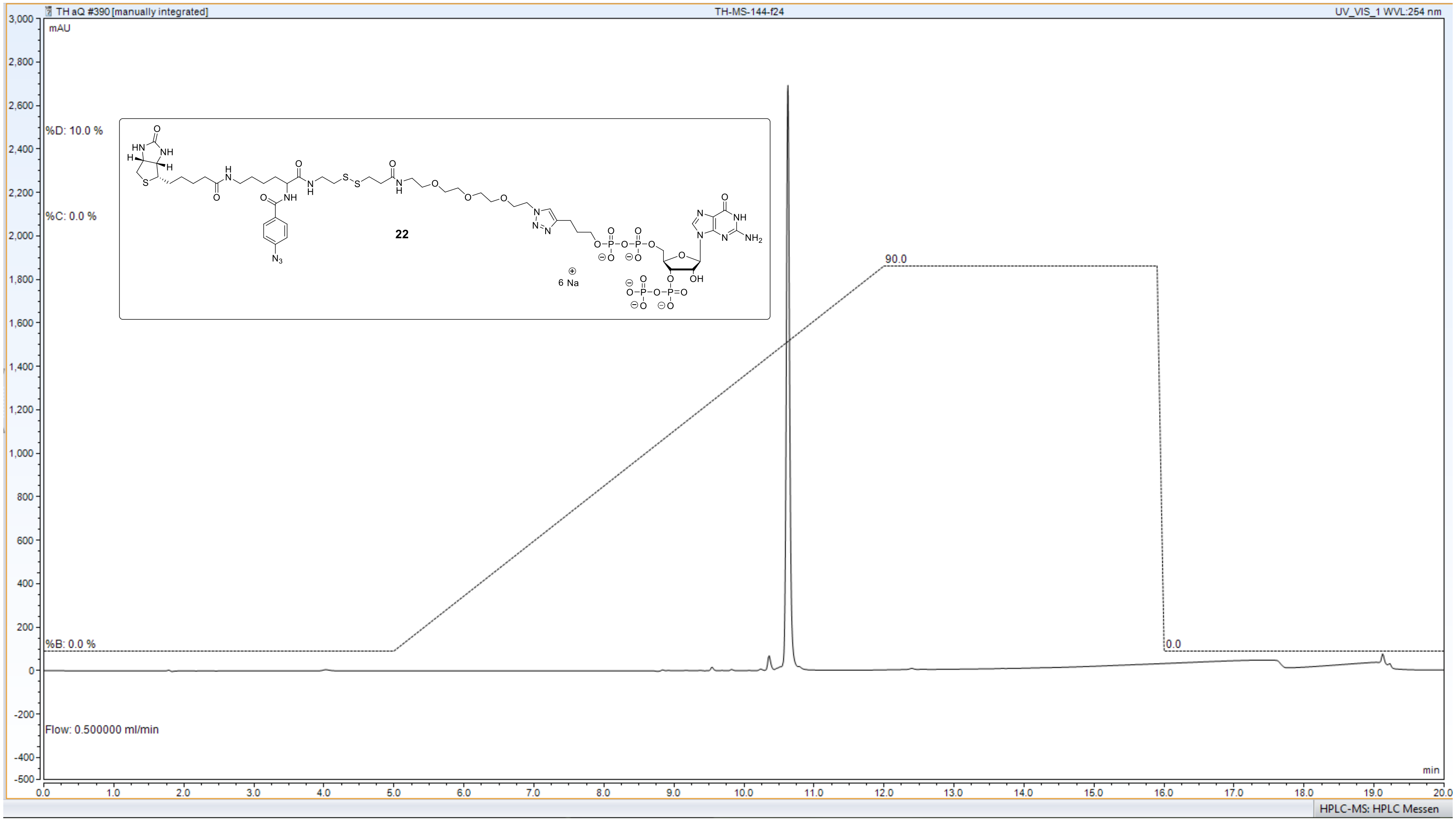


**C18 AQ – column** ($H_2O$/MeCN – gradient, 10 mM TEAA)

***HPLC – UV – analysis of compound 23: ppGpp – CC3***

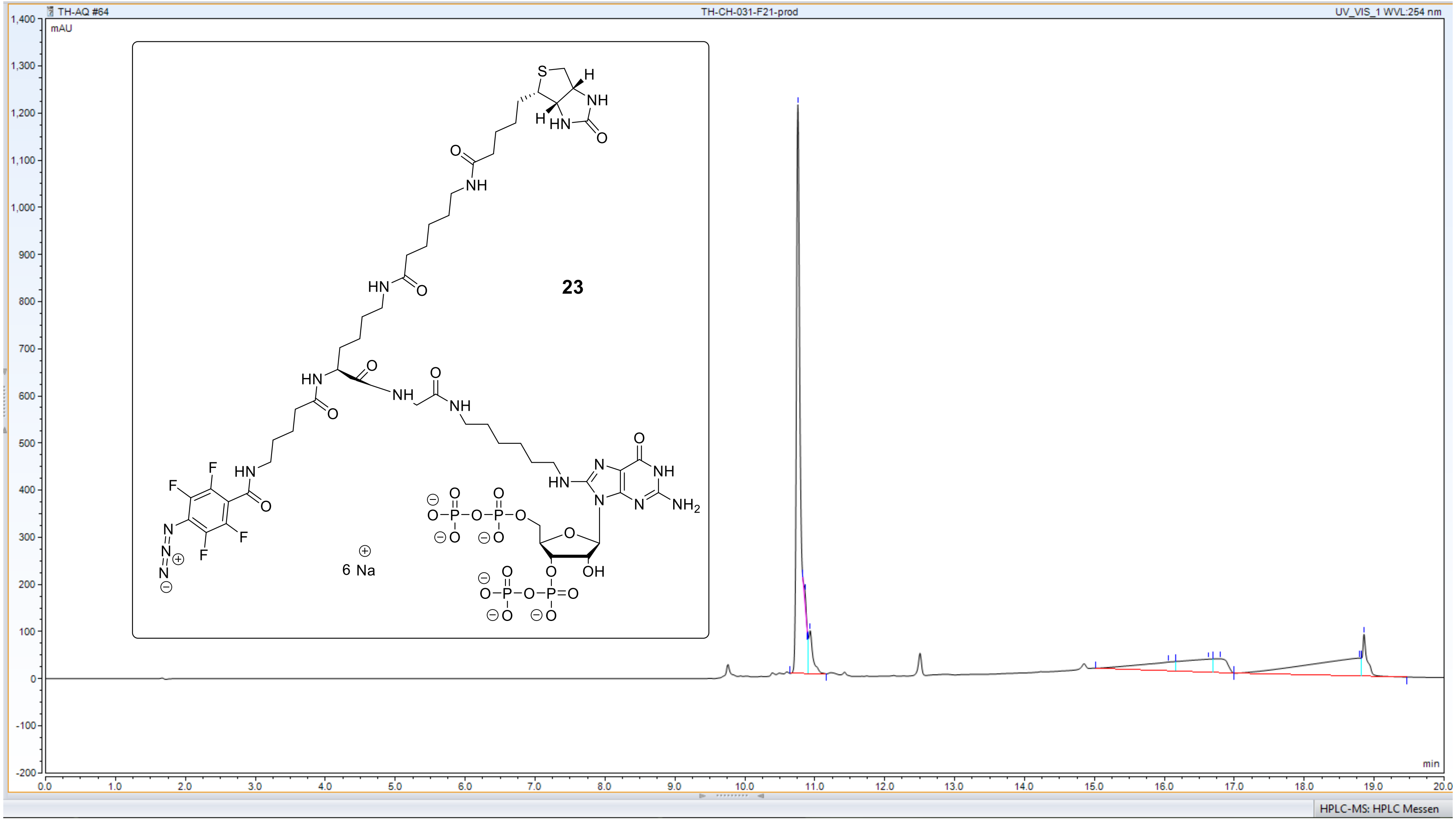


**C18 AQ – column** ($H_2O$/MeCN – gradient, 10 mM TEAA)

***HPLC – UV – analysis of compound 24: ppGpp – CC4***

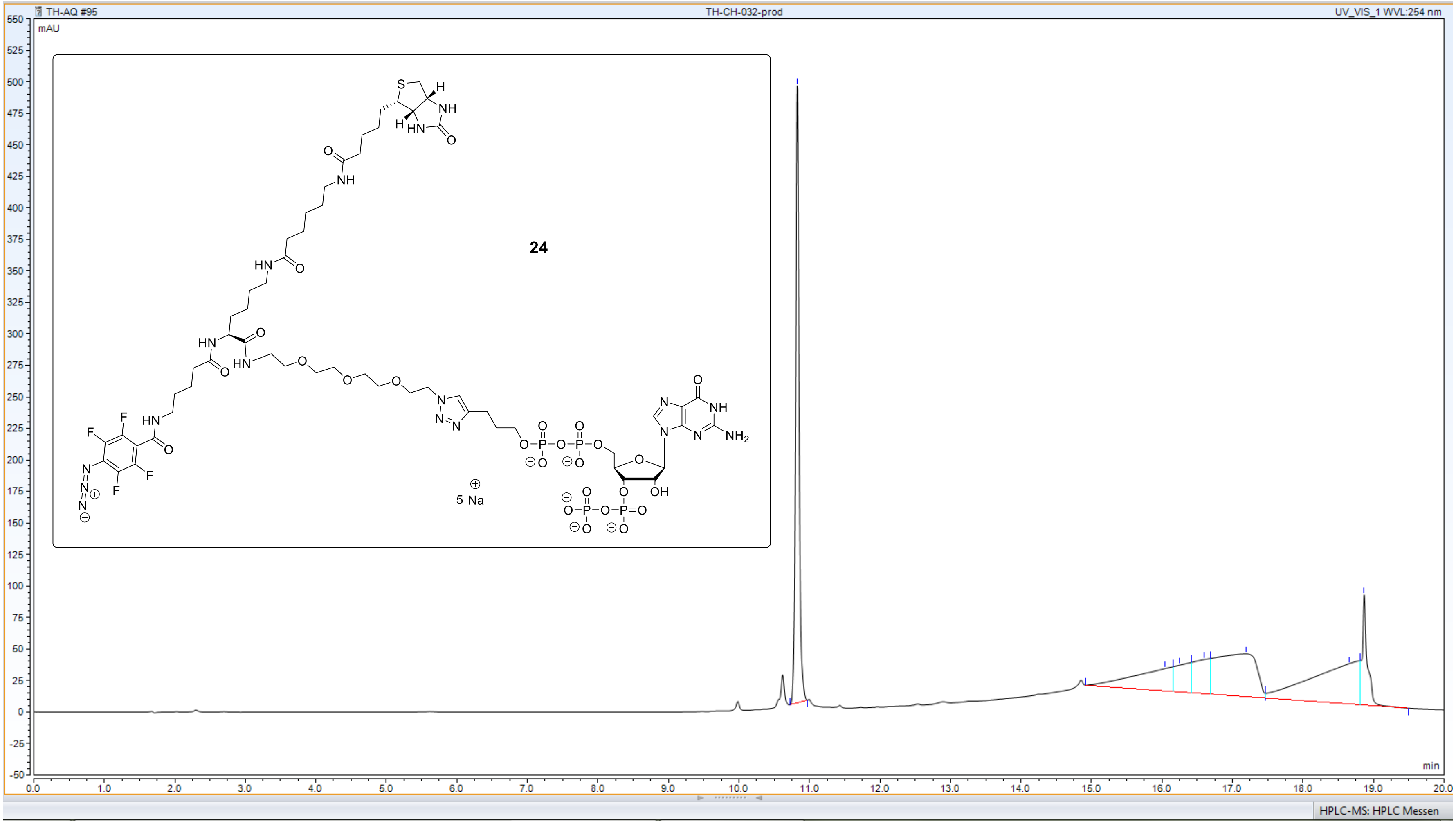


**C18 AQ – column** ($H_2O$/MeCN – gradient, 10 mM TEAA)

***HPLC – UV – analysis of compound 25: pGpp – CC***

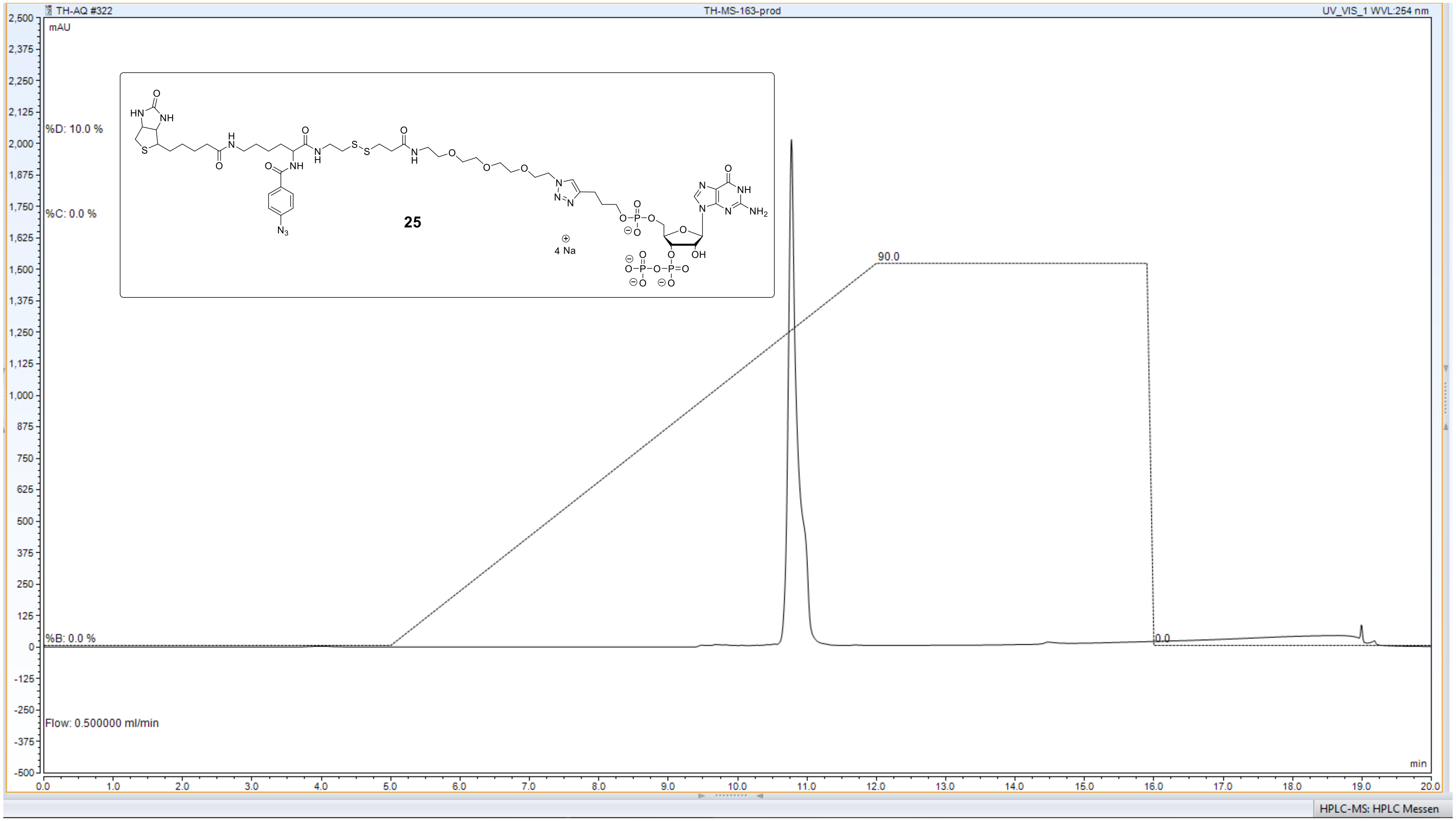


**C18 AQ – column** ($H_2O$/MeCN – gradient, 10 mM TEAA)

### *HPLC – UV – analysis of compound 26: pppGpp – CC*

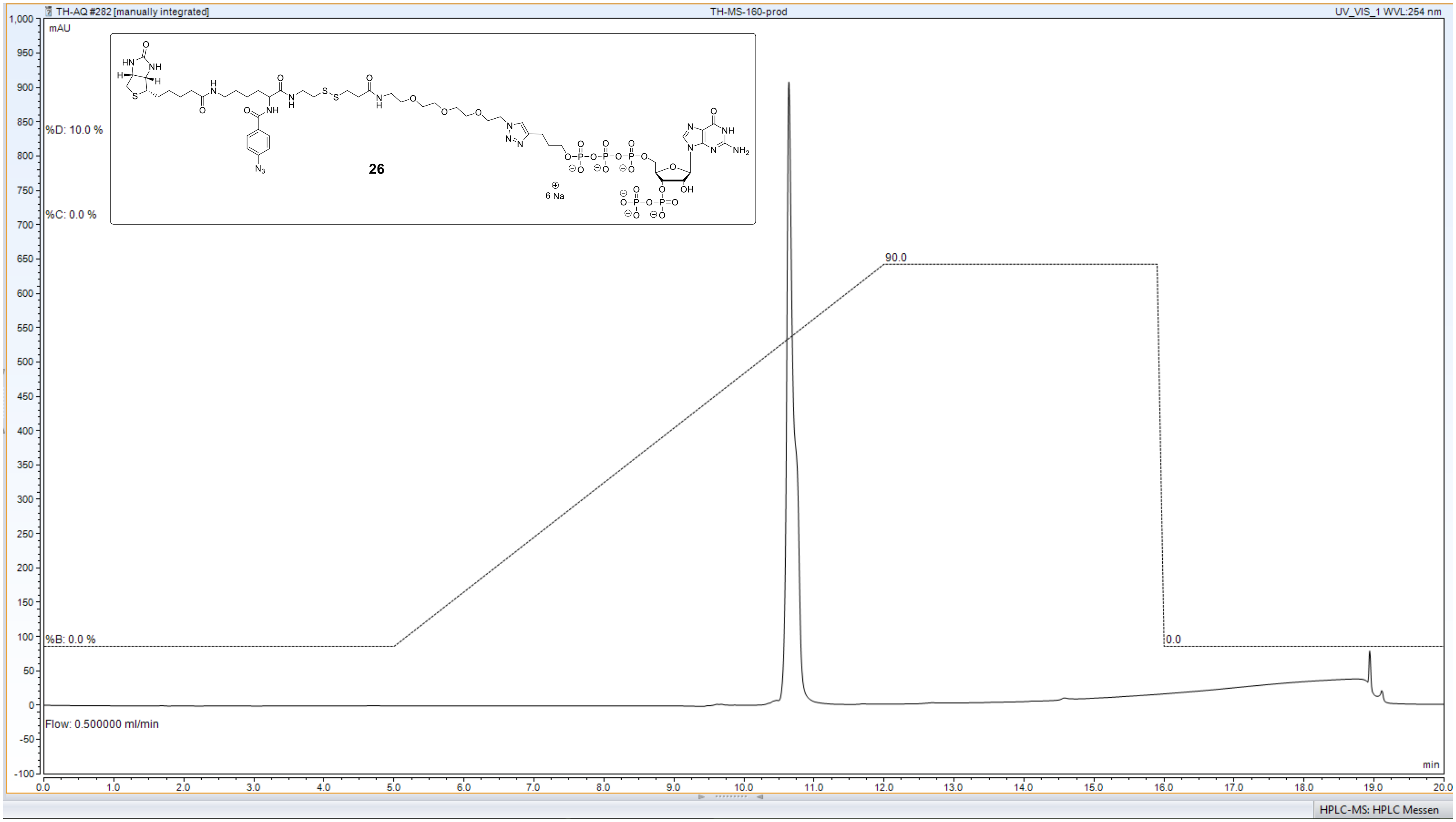


**C18 AQ – column** ($H_2O$/MeCN – gradient, 10 mM TEAA)

***HPLC – UV – analysis of compound 27: ppApp - CC***

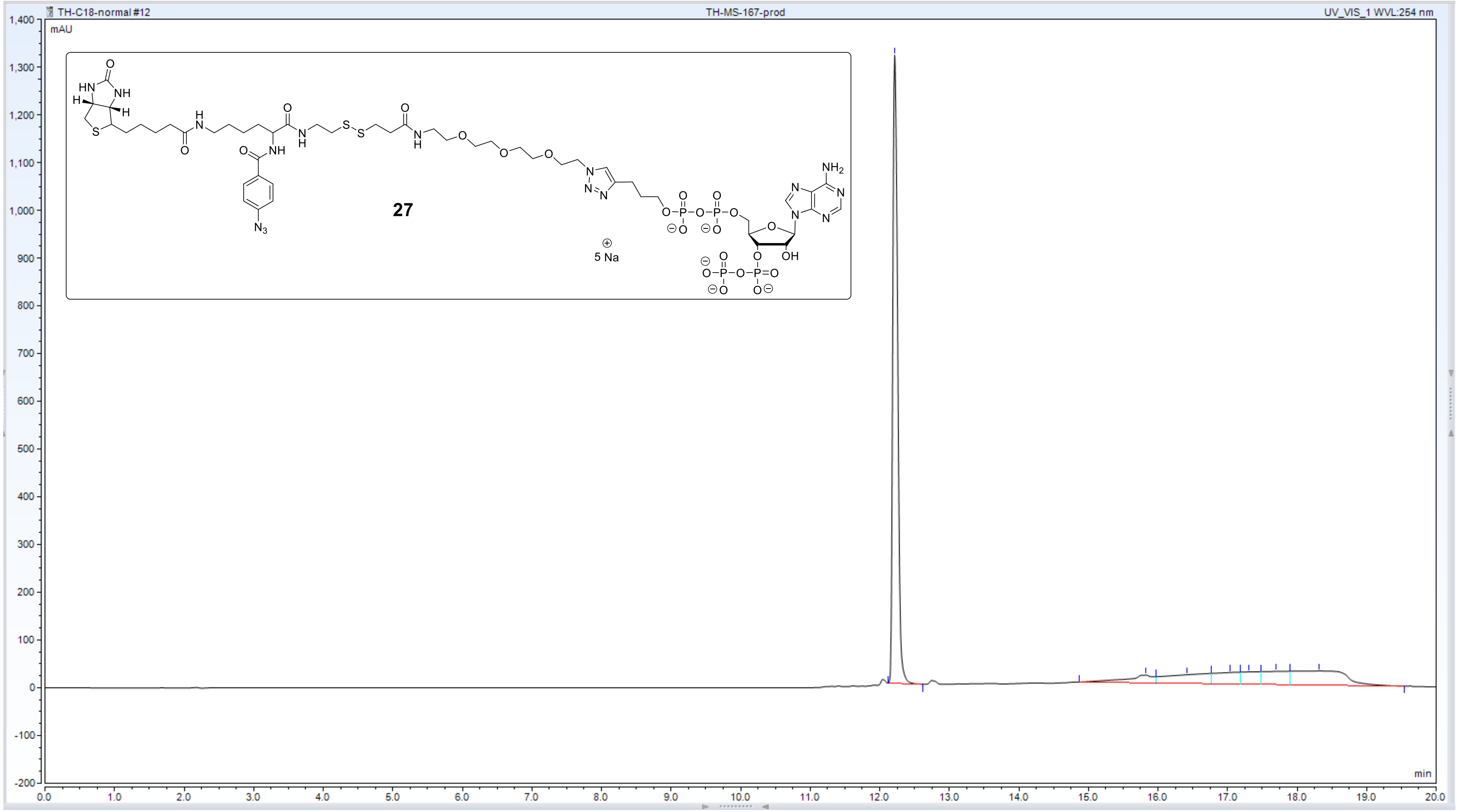


**C18 AQ – column** ($H_2O$/MeCN – gradient, 10 mM TEAA)

***HPLC – UV – analysis of compound SI-13: ppGp - CC***

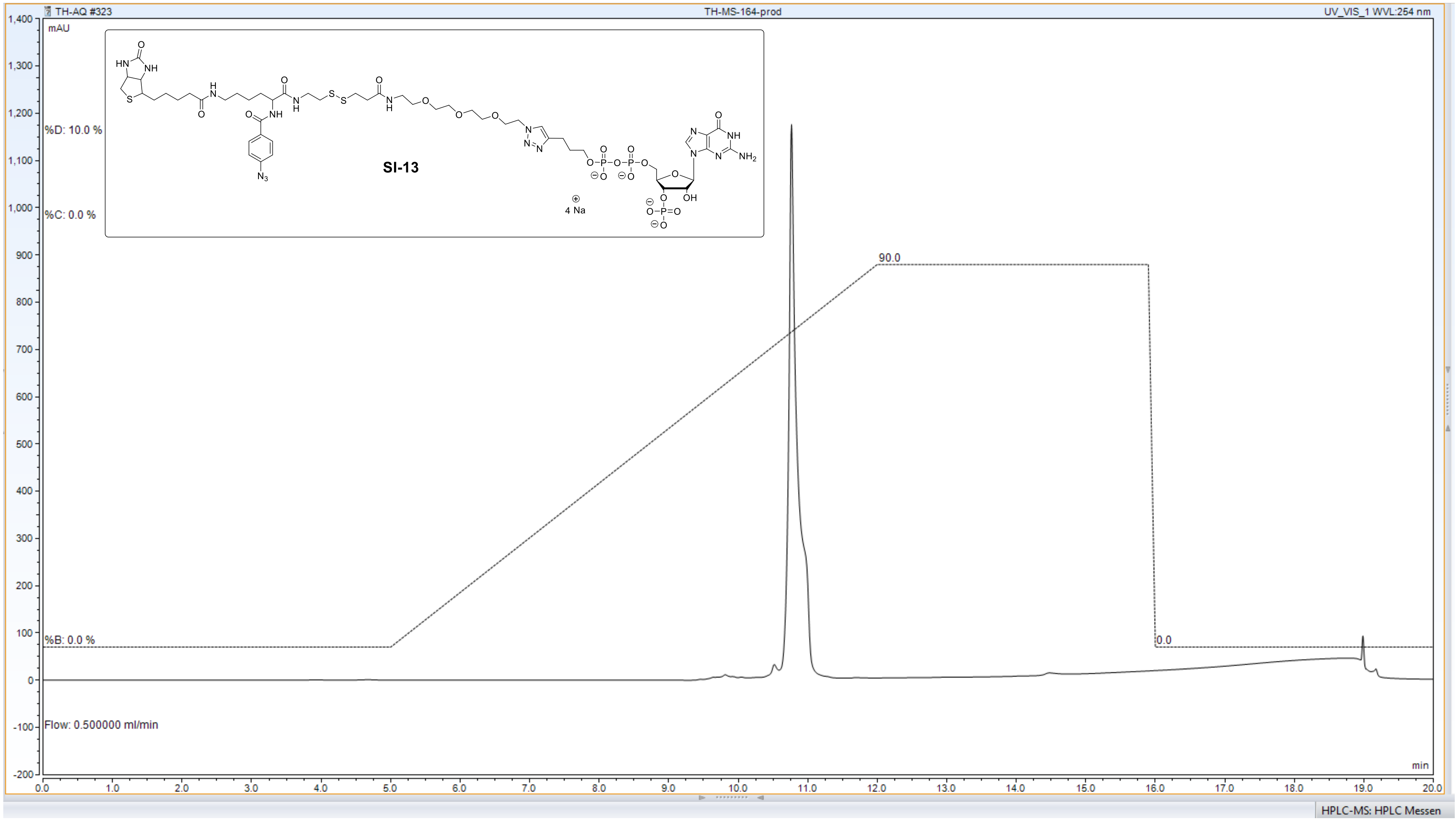


**C18 AQ – column** ($H_2O$/MeCN – gradient, 10 mM TEAA)
